# Complexoform-restricted covalent TRMT112 ligands that allosterically agonize METTL5

**DOI:** 10.1101/2025.05.25.655995

**Authors:** F. Wieland Goetzke, Steffen M. Bernard, Cheng-Wei Ju, Jonathan Pollock, Kristen E. DeMeester, Jacob Gross, Gabriel M. Simon, Chuan He, Bruno Melillo, Benjamin F. Cravatt

## Abstract

Adaptors serve as hubs to regulate diverse protein complexes in cells. This multitude of functions can complicate the study of adaptors, as their genetic disruption may simultaneously impair the activities of several compositionally distinct complexes (or adaptor ‘complexoforms’). Here we describe the chemical proteomic discovery of bicyclopyrrolidine acrylamide stereoprobes that react with cysteine-100 (C100) of the methyltransferase (MT) adaptor TRMT112 in human cells. Curiously, the stereoprobes showed negligible reactivity with uncomplexed recombinant TRMT112, and we found that this interaction was restored excluively in the presence of METTL5, but not other MTs. A co-crystal structure revealed stereoprobe binding to a composite pocket proximal to C100 of TRMT112 that is templated by METTL5 and absent in other TRMT112:MT complexes. Structural rearrangements promoted by stereoprobe binding in turn lead to allosteric agonism of METTL5, thus revealing how covalent ligands targeting a pleiotropic adaptor can confer partner-specific functional effects through reactivity with a single complexoform.

## Introduction

The ∼20,000 protein-encoding genes in the human genome yield a much larger number of proteoforms generated by myriad post-transcriptional and post-translational mechanisms^1–4^. Major categories of proteoforms include splice variants (spliceoforms), covalently modified proteins (e.g., phosphorylated or glycosylated proteins (phospho– or glyco-proteoforms)), and protein complexes of differing compositions (complexoforms^5,6^). Such diversification of the proteome provides important layers of control over dynamic metabolic and signaling pathways in the cell^1–4^.

Small molecules serve as central tools for perturbing the functions of proteins in cells, and there is growing interest in the discovery of chemical probes that target specific proteoforms of proteins^7–12^. Such proteoform-restricted probes have the potential to modulate biology with more precision than that achieved by the complete pharmacological or genetic disruption of proteins. Experimental approaches for identifying ligands that bind specific proteoforms, however, remain poorly defined and are challenged by the diverse and dynamic states of proteins in cells, only a small subset of which are typically accounted for in conventional screening assays performed with recombinant proteins.

We and others have shown that activity-based protein profiling (ABPP) offers a versatile chemical proteomic approach for globally mapping small molecule-protein interactions in human cells^13–16^. ABPP of focused libraries of stereochemically defined covalent (electrophilic or photoreactive) small molecules (or ‘stereoprobes’) has proven particularly useful for identifying cryptic sites of ligandability on diverse classes of proteins, including adaptors, RNA-binding proteins, and transcriptional regulators^17–22^. Some of these small molecule-protein interactions have been further found to require additional factors for recapitulation in recombinant systems (e.g., the presence of DNA for stereoprobe binding to a transcription factor^19^), thus underscoring the potential for ABPP to illuminate state-dependent protein liganding events in cells.

The stereoprobes studied to date by chemical proteomics include both fragments^17,22^ and more elaborated compounds bearing sp³-rich, entropically constrained cores^18,20,21,23,24^. This work has underscored the importance of scaffold diversity for expanding the ligandability of the human proteome, as different classes of acrylamide stereoprobes (azetidines^20,24^, tryptolines^18,20,23^, spirocycles^21^) show only limited overlap in their protein interactions in cells. Here, we describe the synthesis of a structurally distinct class of bicyclopyrrolidine acrylamide stereoprobes and their analysis by cysteine– and protein-directed ABPP^23^. We show that the bicyclopyrrolidine acrylamides, despite having substantially attenuated intrinsic and proteomic reactivity compared to other classes of acrylamide stereoprobes (azetidines, tryptolines), stereoselectively engage a unique set of proteins in human cancer cells. Integration of our cysteine– and protein-directed ABPP data revealed a surprising inconsistency in the apparent reactivity of bicyclopyrrolidine acrylamides with the methytransferase (MT) adaptor protein TRMT112 that we determined to reflect the exclusive engagement of cysteine-100 (C100) of TRMT112 when this adaptor is bound to METTL5, but not other MT partners. Structural studies reveal that the bicyclopyrrolidine acrylamides bind to a composite pocket at the TRMT112:METTL5 interface in close proximity to C100 that is not found in other TRMT112:MT complexes. We finally show that bicyclopyrrolidine acrylamides promote allosteric changes in the TRMT112:METTL5 complex that correlate with enhanced RNA methyltransferase activity. These findings provide a chemical proteomic roadmap for the discovery of complexoform-restricted covalent liganding events and reveal how such interactions can yield selective agonists of one partner (METTL5) of a pleiotropic adaptor protein (TRMT112).

## Results

### Design and synthesis of bicyclopyrrolidine acrylamide stereoprobes

Bicyclic and spirocyclic saturated heterocycles have emerged as building blocks of interest for modern medicinal chemistry because they give access to less explored chemical space and their high sp^3^-content and conformational restriction can offer beneficial (physio)chemical properties for binding to historically challenging protein pockets^25,26^. Here we designed and synthesized two diastereoisomeric pairs of enantiomers of acrylamide stereoprobes based on an octahydrocyclopenta[c]pyrrole core scaffold (henceforth referred to as ‘bicyclopyrrolidines’) (**Fig. 1a**). The structures feature two diversifiable appendages and an endocyclic nitrogen as an attachment point for a cysteine-reactive acrylamide. The synthesis relies on an enantioselective (>95% ee) rhodium-catalyzed arylation of a racemic allyl chloride precursor ((±)-**1**)^27^ followed by a subsequent diastereo– and regio-selective rhodium-catalyzed hydroboration/oxidation sequence (**5**, **6**)^28^ with an optional Mitsunobu stereoinversion of the secondary alcohol to access the alternative diastereomer (**7**, **8**) (**Extended Data Fig. 1**). Each stereoisomer can then be elaborated into an acrylamide in a synthetic sequence including S_N_Ar reactions with electron-deficient aryl halides^29^, additional functionalization reactions, and a final deprotection/acryloylation sequence. However, it should be noted that only four of the possible isomers can be synthesized via this route.

**Fig. 1.**
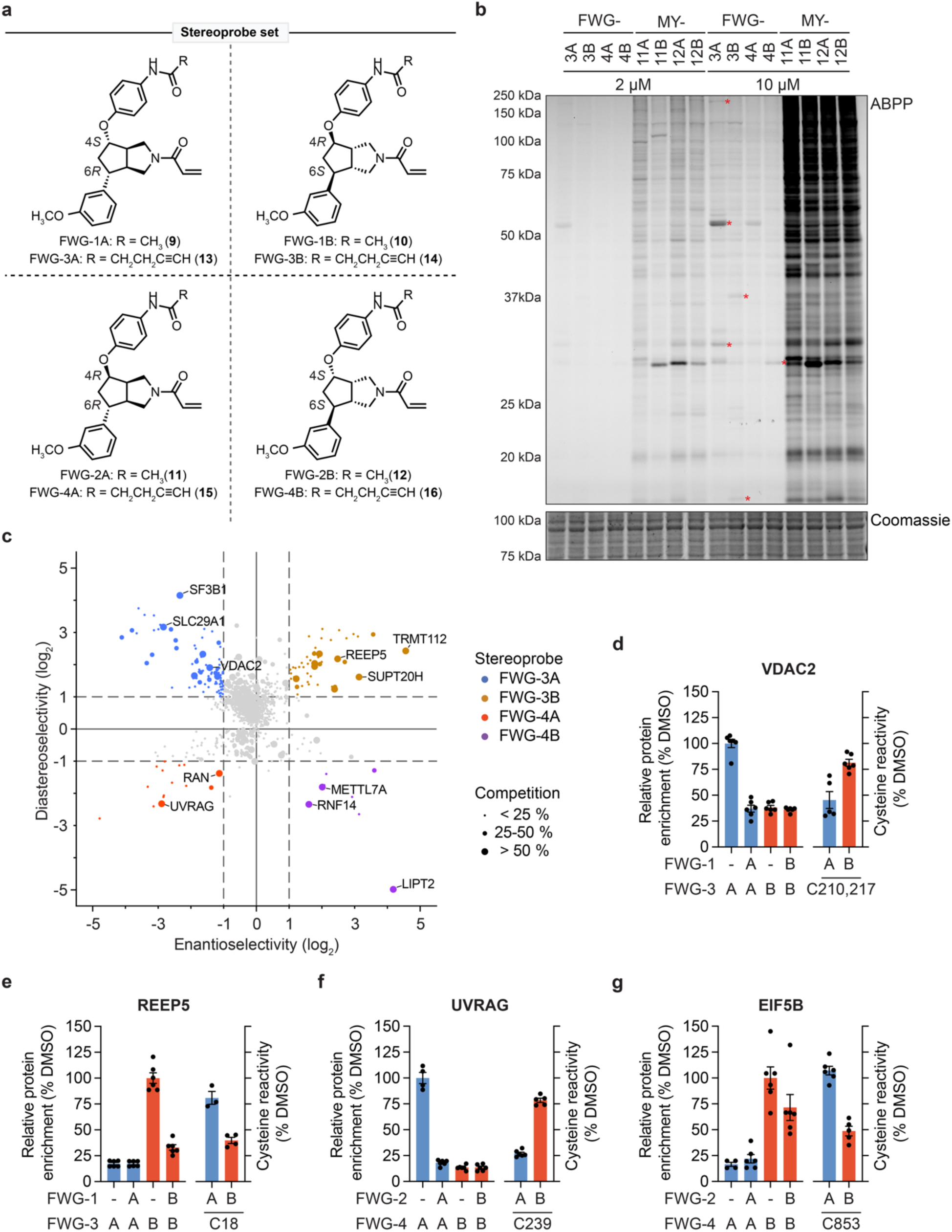
Structures and chemical proteomic analysis of bicyclopyrrolidine acrylamide stereoprobes. **a**, Structures of non-alkyne (FWG-1A (**9**), 1B (**10**), 2A (**11**), and 2B (**12**)) and alkyne (FWG-3A (**13**), 3B (**14**), 4A (**15**), and 4B (**16**)) bicyclopyrrolidine acrylamide stereoprobes. **b**, Gel-ABPP data for Ramos cells treated with the indicated concentrations of alkyne bicyclopyrrolidine acrylamide or azetidine acrylamide^24,30^ (MY-11A, 11B, 12A, and 12B; see Supplementary Information) stereoprobes for 1 h. Stereoprobe-reactive proteins were visualized by CuAAC conjugation to an azide-rhodamine reporter group, SDS-PAGE, and in gel fluorescence scanning. Upper and lower gels show ABPP data and Coomassie blue staining, respectively. Red asterisks mark representative proteins that were stereoselectively engaged by bicyclopyrrolidine acrylamide acrylamides (shown for 10 µM condition). Data are from a single experiment representative of two independent experiments. **c**, Quadrant plot showing stereoselectively liganded proteins for each stereoconfiguration of the bicyclopyrrolidine acrylamides in protein-directed ABPP experiments performed as follows: Ramos cells treated with non-alkyne competitors FWG-1A, 1B, 2A, and 2B (50 μM, 3 h), followed by stereomatched alkynes FWG-3A, 3B, 4A, and 4B (10 μM,1 h) and protein-directed ABPP analysis. Enantioselectivity (*x* axis) is the ratio of enrichment for one stereoisomer versus its enantiomer, and diastereoselectivity (*y* axis) is the ratio of enrichment of one stereoisomer versus the average of its two diastereomers. Data are average values for four-six independent biological experiments. **d-g**, Bar graphs comparing cysteine– and protein-directed ABPP for representative proteins stereoselectively liganded by bicyclopyrrolidine acrylamides (see **Table 1** for more complete list of liganded proteins). Data are average values ± s.e.m. for four-six independent biological experiments.

**Table 1.**
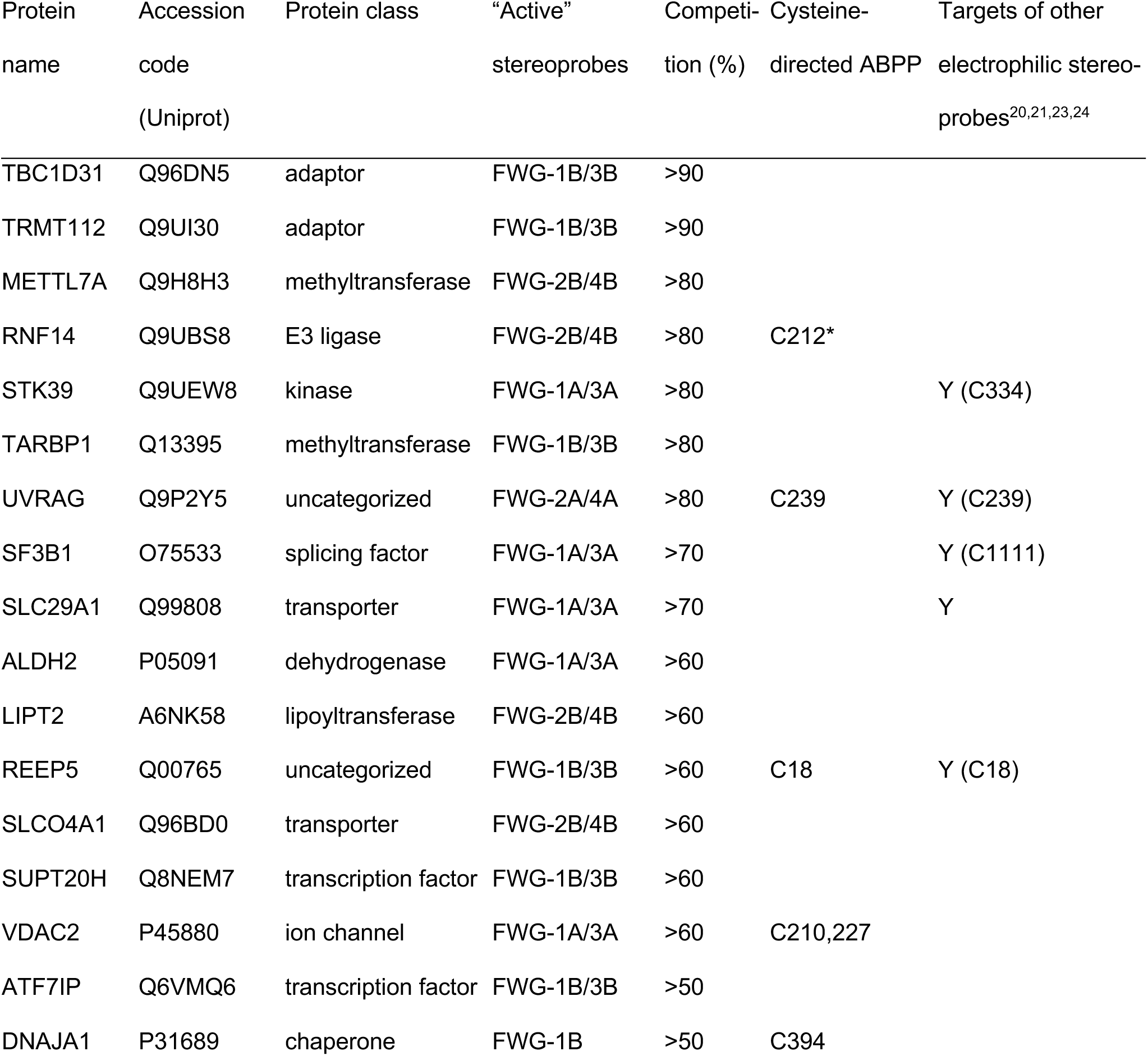
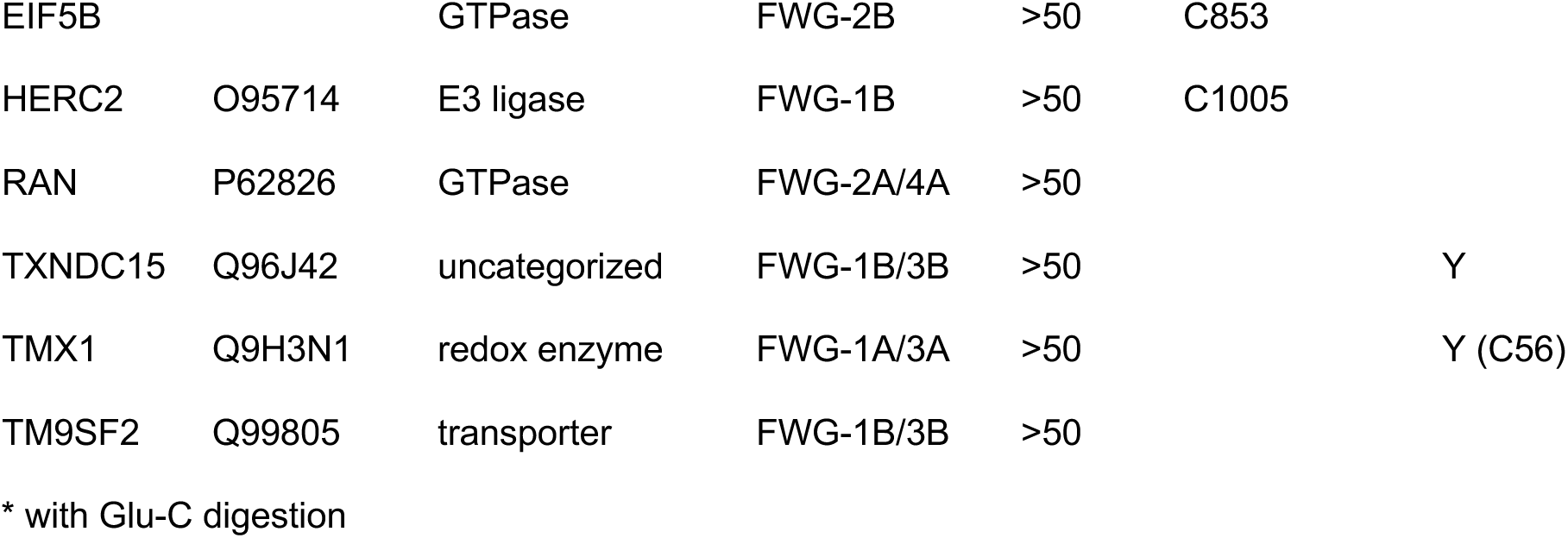
List of proteins that were stereoselectively liganded by bicyclopyrrolidine acrylamide stereoprobes as determined by cysteine– and protein-directed ABPP of Ramos cells (a protein was considered liganded if it displayed > 2-fold enantioselective enrichment by one or more stereoprobe in protein-directed ABPP experiments; and >50% competition of this enrichment by non-alkyne stereoprobes in cysteine and/or protein-directed ABPP experiments. Protein class assignments are from GO (Panther), KEGG BRITE, and UniProt.

| Protein name | Accession code (Uniprot) | Protein class | “Active” stereoprobes | Competition (%) | Cysteine-directed ABPP | Targets of other electrophilic stereoprobes<sup>20,21,23,24</sup> |
| --- | --- | --- | --- | --- | --- | --- |
| TBC1D31 | Q96DN5 | adaptor | FWG-1B/3B | >90 |  |  |
| TRMT112 | Q9UI30 | adaptor | FWG-1B/3B | >90 |  |  |
| METTL7A | Q9H8H3 | methyltransferase | FWG-2B/4B | >80 |  |  |
| RNF14 | Q9UBS8 | E3 ligase | FWG-2B/4B | >80 | C212* |  |
| STK39 | Q9UEW8 | kinase | FWG-1A/3A | >80 |  | Y (C334) |
| TARBP1 | Q13395 | methyltransferase | FWG-1B/3B | >80 |  |  |
| UVRAG | Q9P2Y5 | uncategorized | FWG-2A/4A | >80 | C239 | Y (C239) |
| SF3B1 | O75533 | splicing factor | FWG-1A/3A | >70 |  | Y (C1111) |
| SLC29A1 | Q99808 | transporter | FWG-1A/3A | >70 |  | Y |
| ALDH2 | P05091 | dehydrogenase | FWG-1A/3A | >60 |  |  |
| LIPT2 | A6NK58 | lipoyltransferase | FWG-2B/4B | >60 |  |  |
| REEP5 | Q00765 | uncategorized | FWG-1B/3B | >60 | C18 | Y (C18) |
| SLCO4A1 | Q96BD0 | transporter | FWG-2B/4B | >60 |  |  |
| SUPT20H | Q8NEM7 | transcription factor | FWG-1B/3B | >60 |  |  |
| VDAC2 | P45880 | ion channel | FWG-1A/3A | >60 | C210,227 |  |
| ATF7IP | Q6VMQ6 | transcription factor | FWG-1B/3B | >50 |  |  |
| DNAJA1 | P31689 | chaperone | FWG-1B | >50 | C394 |  |
| EIF5B |  | GTPase | FWG-2B | >50 | C853 |  |
| HERC2 | O95714 | E3 ligase | FWG-1B | >50 | C1005 |  |
| RAN | P62826 | GTPase | FWG-2A/4A | >50 |  |  |
| TXNDC15 | Q96J42 | uncategorized | FWG-1B/3B | >50 |  | Y |
| TMX1 | Q9H3N1 | redox enzyme | FWG-1A/3A | >50 |  | Y (C56) |
| TM9SF2 | Q99805 | transporter | FWG-1B/3B | >50 |  |  |

\* with Glu-C digestion

### ABPP of bicyclopyrrolidine acrylamide stereoprobes in human cancer cells

Each pair of enantiomeric bicyclopyrrolidine acrylamide stereoprobes was synthesized in non-alkyne (FWG-1A (**9**), 1B (**10**), 2A (**11**), 2B (**12**)) and alkyne (FWG-3A (**13**), 3B (**14**), 4A (**15**), 4B (**16**)) form (**Fig. 1a**) for use in gel– and mass spectrometry (MS)-ABPP experiments in the Ramos B-lymphocyte human cancer cell line. Gel-ABPP experiments revealed that the four alkynylated bicyclopyrrolidine acrylamides – FWG-3A/B and FWG-4A/B – showed much lower proteomic reactivity in Ramos cells compared to previously reported azetidine acrylamides (MY-11A/B and MY-12A/B)^24,30^ (**Fig. 1b**), and these differential profiles matched the respective glutathione reactivities of each stereoprobe class (**Extended Data Fig. 2**). Despite the lower overall reactivity of the bicyclopyrrolidine acrylamides, several stereoselective liganding events were observed for these compounds in Ramos cells (**Fig. 1b**, red asterisks).

We next analyzed the bicyclopyrrolidine acrylamides by protein-directed MS-ABPP^23^ (**Extended Data Fig. 3**), where Ramos cells were first treated with non-alkyne ‘competitors’ (FWG-1A, 1B, 2A, 2B; 50 µM, 3 h; or DMSO control), followed by stereochemically matched alkynes (FWG-3A, 3B, 4A, 4B; 10 µM, 1 h). Cells were then harvested, lysed, and the alkyne-modified proteins conjugated to biotin-azide by copper-catalyzed azide-alkyne cycloaddition (CuAAC or click) chemistry^31,32^, enriched using streptavidin beads, digested with trypsin, and analyzed by multiplexed (tandem mass tag (TMT)) MS-based proteomics. 20 proteins in total were stereoselectively liganded by the bicyclopyrrolidine acrylamides based on the following criteria: i) > 2-fold enantioselective enrichment by an alkyne bicyclopyrrolidine acrylamide; and ii) > 50% competitive blockade of this enrichment by the corresponding non-alkyne competitor (**Table 1** and **Supplementary Dataset 1**). As visualized in a quadrant plot display (**Fig. 1c**), the stereoselectively liganded proteins were distributed across the four bicyclopyrrolidine acrylamides with a slight bias in relative number toward the stereoprobes with a cis display of appendages (FWG-1A/3A (4*S*,6*R*) and FWG-1B/3B (4*R*,6*S*)).

We next performed cysteine-directed ABPP experiments that quantified, in total, > 18,000 cysteines in Ramos cells, of which six showed stereoselective liganding by the bicyclopyrrolidine acrylamides (defined as > 50% enantioselective decreases in iodoacetamide-desthiobiotin reactivity (**Table 1** and **Supplementary Dataset 1**). These events included cysteines in several proteins also assigned as liganded by protein-directed ABPP (REEP5_C18, UVRAG_C239, VDAC2_C210, C227), as well as a cysteine in EIF5B (C853), a protein with stereoselective enrichment and competition values that fell just below the cutoff for liganding in protein-directed ABPP experiments. In general, cysteine– and protein-directed ABPP measured similar extents of bicyclopyrrolidine acrylamide engagement for each protein target quantified by both methods (**Fig. 1d-g**).

The 23 total protein targets of the bicyclopyrrolidine acrylamides originated from diverse structural and functional classes and included several proteins that have not yet been liganded by other classes of acrylamide stereoprobes^20,21,23,24^ and, for which, chemical tools are more generally lacking (**Table 1**). Our ABPP data thus indicate that bicyclopyrrolidine acrylamides engage a unique set of proteins in human cells and do so against a backdrop of reduced overall proteomic reactivity compared to other classes of electrophilic stereoprobes.

### Initial characterization of protein targets of bicyclopyrrolidine acrylamides

A substantial number of bicyclopyrrolidine acrylamide-liganded proteins were identified by protein-, but not cysteine-directed ABPP. As described previously^23^, such stereoprobe liganding events mapped exclusively by protein-directed ABPP often occur at cysteines on non-proteotypic peptides that are difficult to detect by MS in cysteine-directed ABPP experiments. For the E3 ubiquitin ligase RNF14 – a target of the FWG-2B/4B stereoprobes (**Extended Data Fig. 4a**) – we succeeded in mapping the site of liganding as C212 by a combination of cysteine– and protein-directed ABPP experiments performed on cells recombinantly expressing this protein. For the cysteine-directed ABPP experiments, we used an alternative Glu-C protease digest, which quantified six cysteine-containing peptides in RNF14, two of which showed stereoselective decreases in IA-DTB reactivity in FWG-2B-treated cells – amino acids 65-78 (containing C68) and 201-226 (containing C212, C220, C223, C225) (**Extended Data Fig. 4b** and **Supplementary Dataset 1**). In each case, the Glu-C cleavage sites furnished peptides of substantially altered length compared to the corresponding tryptic peptides (**Extended Fig. 4c**), which may have resulted in improved proteotypicity for detection in cysteine-directed ABPP experiments. Complementary protein-directed ABPP experiments performed with trypsin digests identified a stereoselectively enriched and unmodified peptide containing C220, C223 and C225 in FWG-4B-treated cells (**Extended Data Fig. 4d**, **e** and **Supplementary Dataset 1**), indicating that these cysteines are not the site of bicyclopyrrolidine acrylamide reactivity. Subsequent gel-ABPP experiments revealed that the C68A-RNF14 mutant maintained stereoselective reactivity with FWG-4B, while the C212A-RNF14 mutant lost reactivity (**Extended Data Fig. 4f**), supporting that C212 is the site of engagement by bicyclopyrrolidine acrylamides. Interestingly, the AlphaFold^33,34^-predicted structure of RNF14 places C68 and C212 in close proximity (**Extended Data Fig. 4g**), which suggests that FWG-2B reactivity with C212 may sterically impair IA-DTB reactivity with C68.

A deeper analysis of our chemical proteomic data suggested a different mechanism may underlie the exclusive mapping of TRMT112 as a target of bicyclopyrrolidine acrylamides by protein, but not cysteine-directed ABPP. TRMT112 is an essential 15 kDa adaptor protein that binds to and regulates the function of a diverse set of S-adenosylmethionine (SAM)-dependent methyltransferases (MTs) involved in RNA, DNA, and protein methylation^35–42^. Protein-directed ABPP revealed that each pair of bicyclopyrrolidine acrylamides showed enantioselective reactivity with TRMT112, with FWG-1B/3B showing the strongest engagement profile followed by FWG-2B/4B (**Fig. 2a** and **Extended Data Fig. 5a**). Both cysteines in TRMT112 – C33 and C100 – were quantified in our cysteine-directed ABPP experiments, and C100 showed a modest (∼40%) stereoselective decrease in IA-DTB reactivity in FWG-1B and FWG-2B treated cells that fell just below our two-fold cutoff for liganding (**Fig. 2a** and **Extended Data Fig. 5b**). This change, however, was much smaller in magnitude compared to the > 90% blockade of TRMT112 enrichment caused by FWG-1B in protein-directed ABPP experiments (**Fig. 2a**). Additional evidence that C100 was the likely site of bicyclopyrrolidine acrylamide engagement in TRMT112, the tryptic peptide containing this cysteine was not observed in our protein-directed ABPP experiments, which otherwise showed good sequence coverage of TRMT112 (**Extended Data Fig. 5c**), with all quantified peptides, including the tryptic peptide containing C33, showing clear stereoselective enrichment in FWG-3B-treated cells (**Extended Data Fig. 5d**).

**Fig. 2.**
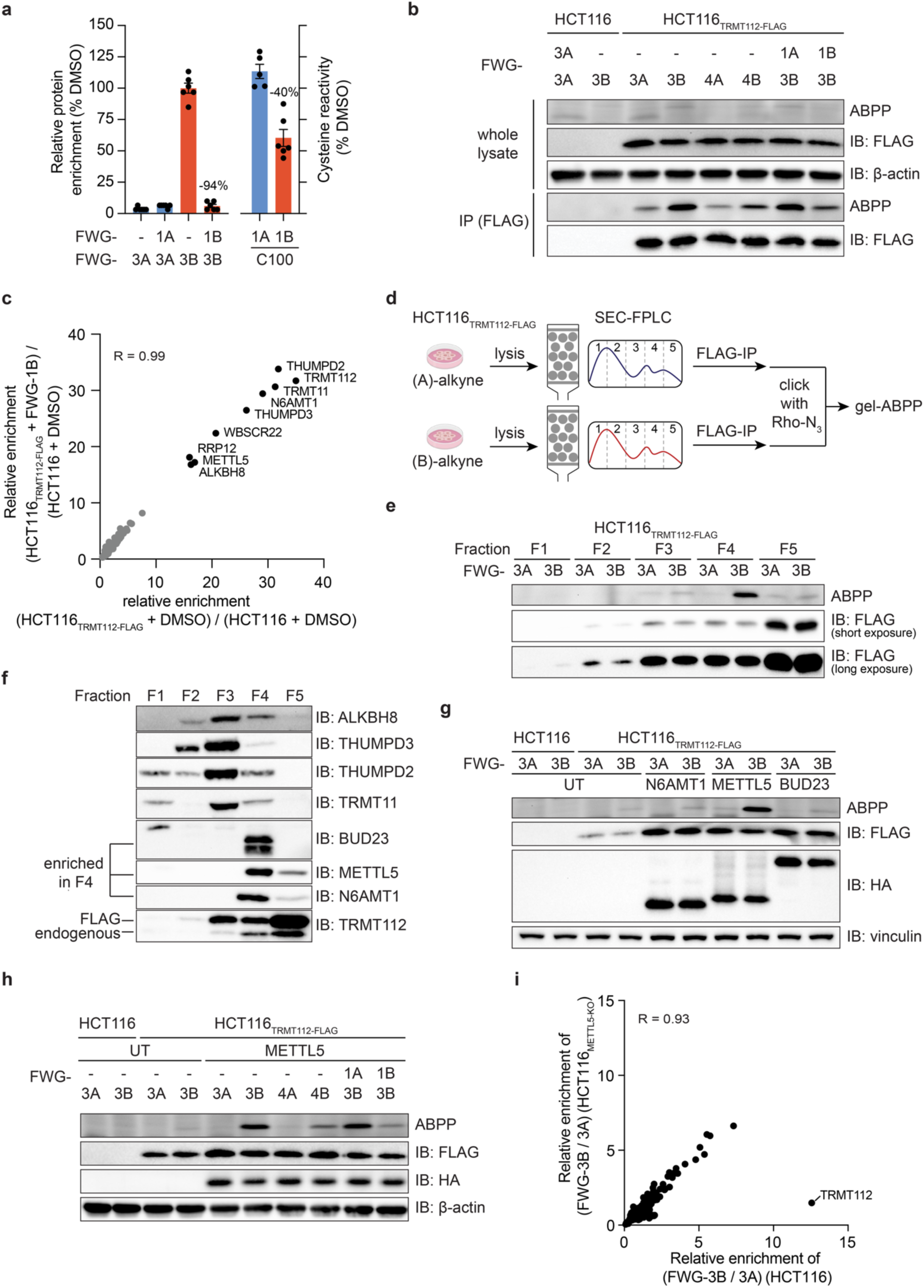
Biyclopyrrolidine acrylamides engage a TRMT112:METTL5 complexoform in cells. **a**, Bar graphs comparing cysteine– and protein-directed ABPP for TRMT112 (see **Extended Data Fig. 4a**, b for bar graphs including all stereoisomers). Left *y-axis*: Protein-directed ABPP data for Ramos cells treated with DMSO, and non-alkyne competitors FWG-1A and FWG-1B (50 μM, 3 h), followed by stereomatched alkynes FWG-3A and 3B (10 μM, 1 h). Right *y-axis*: Cysteine-directed ABPP data for Ramos cells treated with FWG-1A and FWG-1B (50 μM, 3 h), followed by lysis and treatment with IA-DTB (100 μM, 1 h). Data are average values ± s.e.m. for five-six independent biological experiments. **b**, Gel-ABPP data demonstrating stereoselective engagement of recombinant TRMT112-FLAG by FWG-3B that is only visible after anti-FLAG immunoprecipitation. HCT116 (parental) and HCT116 cells with stable expression of recombinant TRMT112-FLAG (HCT116_TRMT112-FLAG_ cells) were treated with FWG-1A and 1B (50 μM, 3 h), followed by treatment with FWG-3A, 3B, 4A, and 4B (10 μM, 1 h). TRMT112-FLAG was enriched by immunoprecipitation with anti-FLAG magnetic beads and analyzed by gel-ABPP (as described in **Fig. 1b**). Data are from a single experiment representative of two independent experiments. **c**, Proteins co-enriched with TRMT112 in anti-FLAG IP-MS experiments performed with parental HCT116 cells (control) or HCT116_TRMT112-FLAG_ cells treated with DMSO of FWG-1B (50 µM, 4 h). Scatter-plot showing strong correlation (R = 0.99) for co-enriched proteins in DMSO-(x axis) vs FWG-1B-(y-axis) treated HCT116_TRMT112-FLAG_ cells. Data are average values for eight independent biological experiments. **d**, Workflow of the SEC-ABPP experiments. Cells with expression of WT-TRMT112-FLAG were treated with alkyne probes. Clarified lysates were injected into a Superdex 200 Increase 10/300 GL column attached to an ÄKTA pure FPLC system and fractionated into five fractions. WT-TRMT112-FLAG was enriched by immunoprecipitation with anti-FLAG agarose beads and analyzed by gel-ABPP (as described in **Fig. 1b**). **e**, Gel-ABPP (top) and western blotting (middle and bottom) data of SEC fractions from HCT116_TRMT112-FLAG_ cells treated with FWG-3A or FWG-3B (10 µM, 1 h) showing that the FWG-3B-reactive proteoform of TRMT112 elutes in fraction 4 while the majority of the TRMT112 protein is found in fraction 5. Data are from a single experiment representative of two independent experiments. **f**, Western blots showing SEC elution profiles of TRMT112-associated MTs and TRMT112 from SEC-fractionated lysates of HCT116_TRMT112-FLAG_ cells. Data are from a single experiment representative of two independent experiments. **g**, Gel-ABPP data demonstrating stereoselective engagement of TRMT112 by FWG-3B in HCT116_TRMT112-FLAG_ cells co-expressing METTL5, but not N6AMT1 or BUD23. HCT116_TRMT112-FLAG_ cells were transiently transfected with cDNAs for HA-tagged N6AMT1, METTL5, and BUD23, followed by treatment with FWG-3A and 3B (10 μM, 1 h), and gel-ABPP analysis. Data are from a single experiment representative of two independent experiments. **h**, Gel-ABPP data demonstrating blockade of the FWG-3B-WT-TRMT112 interaction in METTL5-co-expressing HCT116_TRMT112-FLAG_ cells by FWG-1B, but not FWG-1A. HCT116_TRMT112-FLAG_ cells were transiently transfected with a cDNA for HA-METTL5 and pre-treated with FWG-1A or 1B (50 μM, 3 h), followed by FWG-3A, 3B, 4A, and 4B (10 μM, 1 h), and analyzed by gel-ABPP (as described in **Fig. 1b**). Data are from a single experiment representative of two independent experiments. **i**, Protein-directed ABPP experiments demonstrating loss of stereoselective enrichment of TRMT112 by FWG-3B in METTL5-KO cells. Scatter-plot comparing the stereoselective enrichment profiles (FWG-3B/FWG-3A; 10 µM, 1 h) of proteins in parental (*x* axis) vs METTL5-KO (*y* axis) HCT116 cells. Data are average values for four independent biological experiments. The linear regression analysis was performed excluding the TRMT112 data.

Curious about the quantitative discrepancy in liganding of TRMT112 as measured by cysteine-(∼40%) and protein-(>90%) directed ABPP, we assessed bicyclopyrrolidine acrylamide reactivity with HCT116 cells stably expressing recombinant FLAG epitope-tagged TRMT112 (HCT116_TRMT112-FLAG_). Despite strong expression of recombinant TRMT112 as determined by anti-FLAG immunoblotting, we observed negligible signals for reactivity with the preferred stereoprobe FWG-3B by gel-ABPP (**Fig. 2b**). Following anti-FLAG immunoprecipitation (IP), the stereoselective engagement of TRMT112 by FWG-3B and the stereoselective blockade of this engagement by FWG-1B could be observed, indicating that at least a portion of the recombinant protein retained the reactivity profile observed for endogenous TRMT112 (**Fig. 2b**). Consistent with a small fraction of recombinant TRMT112 existing in a ligandable state, anti-FLAG IP-MS experiments failed to detect substantial reductions in the unmodified C100-containing tryptic peptide in FWG-1B-treated cells (**Extended Data Fig. 5e**). We also attempted to express and characterize C100A and C100S mutants of TRMT112, but these variants were produced at very low levels in HCT116 cells (**Extended Data Fig. 5f**).

Our ABPP data for both endogenous and recombinant TRMT112 suggested that the bicyclopyrrolidine acrylamide stereoprobes may react with a specific proteoform of this protein. We next set out to identify the stereoprobe-liganded TRMT112 proteoform.

### Bicyclopyrrolidine acrylamides engage a TRMT112:METTL5 complexoform in cells

The anti-FLAG IP-MS experiments described above also identified substantial co-enrichment of several previously reported MT interaction partners of TRMT112^35^ – THUMPD2, TRMT11, N6AMT1, THUMPD3, BUD23, METTL5, and ALKBH8 (**Fig. 2c**), and FWG-1B did not alter these interactions (**Fig. 2c**). Considering that TRMT112 participates in such a diverse array of MT complexes in human cells, we contemplated whether the bicyclopyrrolidine acrylamides might preferentially react with one or a subset of these TRMT112:MT complexoforms. We designed a workflow to test this hypothesis wherein HCT116_TRMT112-FLAG_ cells were treated with FWG-3A or FWG-3B, lysed, and then fractionated by size exclusion chromatography (SEC). Each SEC fraction was then subject to anti-FLAG IP, click conjugation of an azide-rhodamine tag, and analysis by gel-ABPP (**Fig. 2d**). This experiment revealed that FWG-3B predominantly reacted with the proteoform of TRMT112-FLAG migrating in fraction 4 of the SEC profile, whereas the vast majority of the TRMT112-FLAG protein, as determined by anti-FLAG immunoblotting, was found in fraction 5 with lower quantities spread across fractions 2-4 (**Fig. 2e**). The reaction of FWG-3B with TRMT112 in fraction 4 was stereoselective (not observed with FWG-3A) (**Fig. 2e**) and blocked by pre-treatment of cells with FWG-1B (**Extended Data Fig. 5g**), matching the properties of endogenous TRMT112 characterized in Ramos cells.

Additional immunoblotting experiments of the SEC fractions revealed that three TRMT112 interacting MTs – BUD23, METTL5, and N6AMT1 – were enriched in fraction 4, while other MTs, such as THUMPD2, THUMPD3, and TRMT11, were found in higher molecular weight (MW) fractions (F2-F3) (**Fig. 2f**). These data suggested that the bicyclopyrrolidine acrylamides may preferentially react with TRMT112 when this adaptor is bound to one of the three co-migrating MTs in fraction 4. We transiently co-expressed HA-tagged BUD23, METTL5, and N6AMT1 in HCT116_TRMT112-FLAG_ cells and then treated these cells with FWG-3A or FWG-3B followed by gel-ABPP analysis. This experiment revealed that FWG-3B reacted with TRMT112 exclusively in cells co-expressing the RNA methyltransferase METTL5 (**Fig. 2g**). The FWG-3B-TRMT112 interaction in METTL5-co-expressing cells was stereoselective (**Fig. 2g, h**) and blocked by pre-treatment with FWG-1B (**Fig. 2h**). Finally, to determine whether METTL5 was also required for bicyclopyrrolidine acrylamide reactivity with endogenous TRMT112, we generated METTL5-knockout (KO) HCT116 cells using CRISPR-Cas9 methods. Protein-directed ABPP experiments revealed a complete loss of stereoselective enrichment of TRMT112 by FWG-3B in METTL5-KO cells compared to parental HCT116 cells, while other FWG-3B-enriched proteins were generally unaffected in METTL5-KO cells (**Fig. 2i** and **Supplementary Dataset 1**). Additionally, the METTL5-KO cells maintained similar expression levels of TRMT112 compared to parental HCT116 cells as determined by immunoblotting (**Extended Data Fig. 5h**).

Taken together, our results support that bicyclopyrrolidine acrylamide stereoprobes react with TRMT112 exclusively when this pleiotropic adaptor is bound to METTL5. We next set out to further characterize the structure-activity relationship and mechanism of this complexoform-restricted liganding event.

### Optimization of covalent ligands for the TRMT112:METTL5 complex

While our initial bicyclopyrrolidine acrylamide ligand FWG-1B engaged TRMT112 in cells with high stereoselectivity, this interaction showed modest potency as determined by gel-ABPP (IC_50_ value = 12 µM; 95% confidence intervals (C.I.) of 11 – 14 μΜ; **Extended Data Fig 6a, b**). We therefore performed an initial structure-activity analysis to identify higher potency ligands for the TRMT112:METTL5 complex. We selected three positions on the bicyclopyrrolidine scaffold that could be diversified using robust chemical transformations, including amide coupling, S_N_Ar/Mitsunobu reaction, and rhodium-catalyzed (Rh-cat.) arylation (**Fig. 3a**). Analogs at the C4 substitution (FWG-9B, 11B, 19B, 21B, and 23B; **Fig. 3b**) produced only subtle differences in potency as determined by gel-ABPP experiments performed in HCT116_TRMT112-FLAG:HA-METTL5_ cells, with the exception of FWG-23B, which featured an *N*-(2-phenyl)acetamide substitution and exhibited a marked decrease in potency (**Fig. 3c**).

**Fig. 3.**
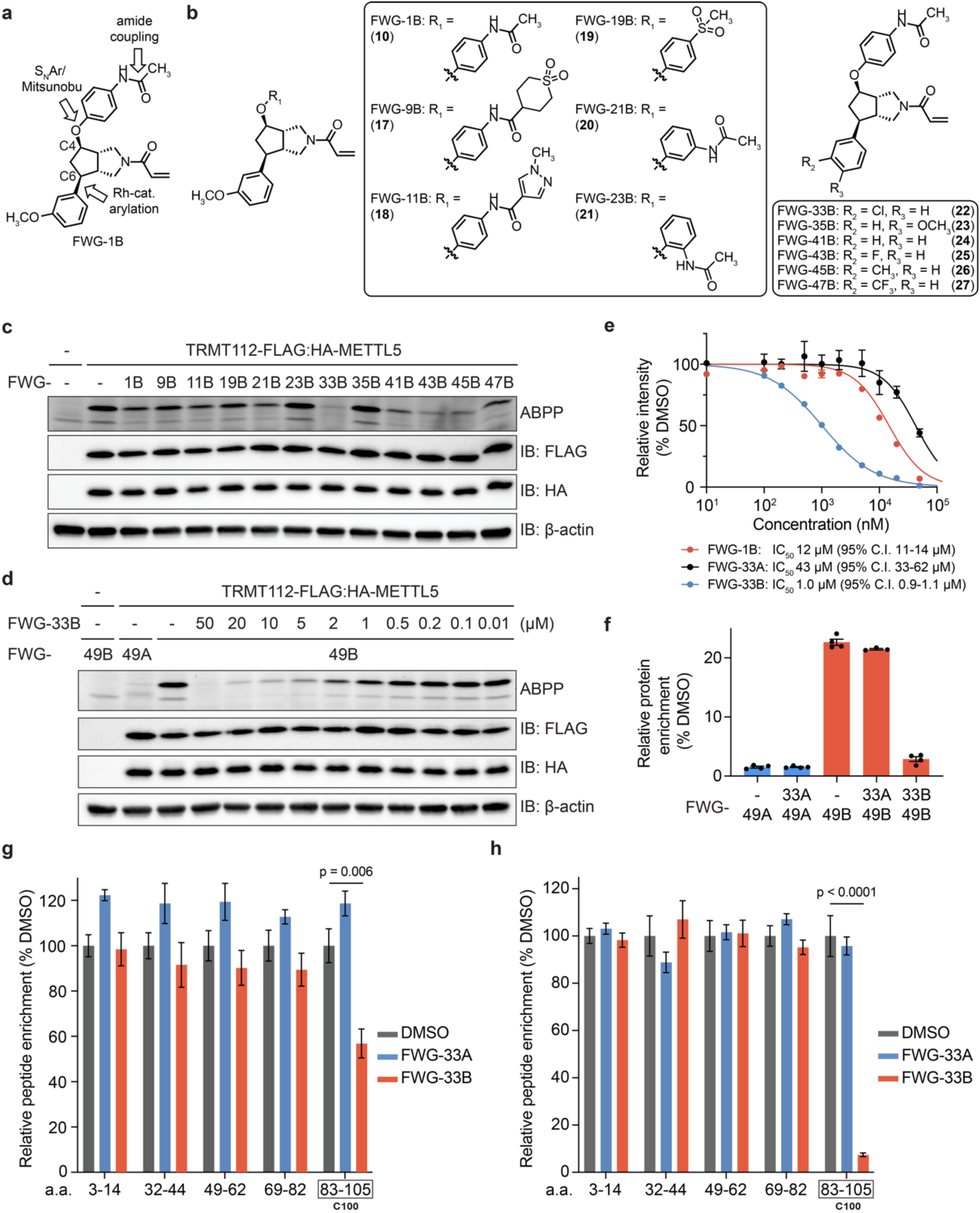
Optimization of bicyclopyrrolidine acrylamide stereoprobes targeting the TRMT112:METTL5 complex. **a**, Diversification points for bicyclopyrrolidine acrylamide FWG-1B. **b**, Structures of bicyclopyrrolidine acrylamide analogs tested for interactions with the TRMT112:METTL5 complex. **c**, Gel-ABPP data showing engagement of TRMT112 by bicyclopyrrolidine acrylamide analogs in comparison to original ligand FWG-1B. HCT116 cells stably expressing TRMT112-FLAG and HA-METTL5 (HCT116_TRMT112-FLAG:HA-METTL5_ cells) were pre-treated with the indicated bicyclopyrrolidine acrylamide analogs (20 μM, 1 h), followed by FWG-3B (10 μM, 1 h), and gel-ABPP analysis. Data are from a single experiment representative of two independent experiments. **d**, Gel-ABPP data showing concentration-dependent blockade of the FWG-49B-TRMT112 interaction by FWG-33B. HCT116_TRMT112-FLAG:HA-METTL5_ cells were pre-treated with the indicated concentrations of FWG-33B for 3 h, followed by FWG-49A and 49B (2 μM, 1 h), and gel-ABPP analysis. Data are from a single experiment representative of two independent experiments. **e**, Quantification of gel-ABPP data for engagement of TRMT112 by FWG-1B, FWG-33A, and FWG-33B in HCT116_TRMT112-FLAG:HA-METTL5_ cells (see **Fig. 3d**, **Extended Data Fig. 6a** and **Extended Data Fig. 6d**). Data are average values ± s.e.m. for two independent biological experiments. **f**, Bar graph showing protein-directed ABPP for endogenous TRMT112 from Ramos cells pre-treated with DMSO or FWG-33A or FWG-33B (5 μM, 3 h), followed by FWG-49A and FWG-49B (2 μM,1 h) and protein-directed ABPP analysis. Data are average values ± s.e.m. for three-four independent biological experiments. **g**, **h**, Quantification of engagement of TRMT112_C100 by FWG-33B (5 µM, 4 h) as measured by monitoring the C100-containing tryptic peptide in anti-FLAG (**g**) or anti-HA (**h**) IP-MS experiments from HCT116_TRMT112-FLAG:HA-METTL5_ cells. Data are average values ± s.e.m. normalized to DMSO for six-eight independent biological experiments. Statistical significance between the DMSO and the FWG-33B treated peptides was assessed using unpaired-multiple *t*-tests (Holm-Šídák approach to multiple comparisons; reported p values are adjusted p values).

In contrast, modifications at the C6 position (FWG-33B, 35B, 41B, 43B, 45B, 47B; **Fig. 3b**) resulted in much more pronounced differences in reactivity with TRMT112, with the 3-chlorophenyl analogue FWG-33B showing the greatest apparent improvement in potency (**Fig. 3c**). The greater impact of the C6 over C4 position on bicyclopyrrolidine acrylamide reactivity with TRMT112 is consistent with our original protein-directed ABPP data, where the C4 epimer of FWG-3B – FWG-4B – also enantioselectively enriched TRMT112 (**Extended Data Fig. 5a**).

We next generated alkyne analogues of FWG-33B and its enantiomer FWG-33A (FWG-49B and 49A; (**Extended Data Fig. 6c**)) and used these compounds as target engagement probes in subsequent gel-ABPP experiments measuring the concentration-dependent reactivity of FWG-1B, FWG-33A, and FWG-33B with TRMT112 in HCT116_TRMT112-FLAG:HA-METTL5_ cells (**Fig. 3d, e** and **Extended Data Fig. 6a, d**). These experiments revealed an ∼10-fold improvement in potency for FWG-33B compared to FWG-1B (IC_50_ values: FWG-1B: 12 μΜ; FWG-33B: 1.0 μΜ (95% C.I. 0.9 – 1.1 μΜ)) with maintenance of stereoselective reactivity (FWG-33A: 43 μΜ (95% C.I. 33 – 62 μΜ)). FWG-49B also appeared to share this increase in potency, as this alkyne stereoprobe proved capable of visualizing endogenous TRMT112 at 2 µM test concentrations in HCT116 cells (**Extended Data Fig. 6e, f**). We next evaluated the proteome-wide reactivity of FWG-33B and alkyne analog FWG-49B by protein-directed ABPP in Ramos cells. These experiments confirmed robust stereoselective enrichment of TRMT112 in FWG-49B vs FWG-49A-treated cells and the near-complete (∼90%) blockade of this enrichment by pre-treatment with FWG-33B (5 μM, 3 h) (**Fig. 3f**). In contrast, pre-treatment with the enantiomer FWG-33A did not perturb enrichment of TRMT112 by FWG-49B (**Fig. 3f**). One additional stereoselective target of FWG-33B was identified in this experiment – TBC1D31 (**Extended Data Fig. 6g** and **Supplementary Dataset 1**) – which is an adaptor protein involved in cilium biogenesis^43^.

We next assessed the fraction of total and METTL5-complexed TRMT112 that reacted with FWG-33B in HCT116_TRMT112-FLAG:HA-METTL5_ cells as measured by IP-MS using anti-FLAG and anti-HA antibodies, respectively. We surmised that, in these experiments, signals for the tryptic peptide containing C100 of TRMT112 (a.a. 83-105: R.TMHLLLEVEVIEGTLQCPESGR.M) would provide an estimate of FWG-33B engagement of total (anti-FLAG) vs METTL5-complexed (anti-HA) TRMT112. The anti-FLAG IP-MS revealed an ∼ 40% reduction in unmodified a.a. 83-105 peptide in FWG-33B-treated cells (**Fig. 3g**), suggesting that, in cells recombinantly expressing both TRMT112 and METTL5, only ∼40% of the total TRMT112 was engaged by FWG-33B. In contrast, the signals for the unmodified a.a. 83-105 peptide were decreased > 90% in anti-HA IP-MS experiments from these same cells (**Fig. 3h**), supporting that the fraction of TRMT112 interacting with METTL5 is fully engaged by FWG-33B in cells.

### Crystal structure of a TRMT112:METTL5 complex bound to FWG-33B

We expressed WT-TRMT112:METTL5 and C100A-TRMT112:METTL5 complexes in *E. coli* (BL21) and purified these complexes by metal affinity chromatography leveraging an N-terminal His-tag incorporated into METTL5, followed by anion-exchange and size-exclusion chromatography. We found that FWG-49B reacted in a stereoselective manner with the purified WT-TRMT112:METTL5 complex, but not the C100A-TRMT112:METTL5 complex, and pre-treatment with FWG-33B, but not FWG-33A (5 µM each, 1 h), fully blocked FWG-49B reactivity with WT-TRMT112 (**Fig. 4a**). We also confirmed the reactivity of FWG-33B with purified WT-TRMT112 by intact MS analysis (**Extended Data Fig. 7a**) and used this method to measure a *k*_obs_/[I] of 160 M^-1^s^-1^ (C.I. 150 – 180 M^-1^s^-1^) for FWG-33B.

**Fig. 4.**
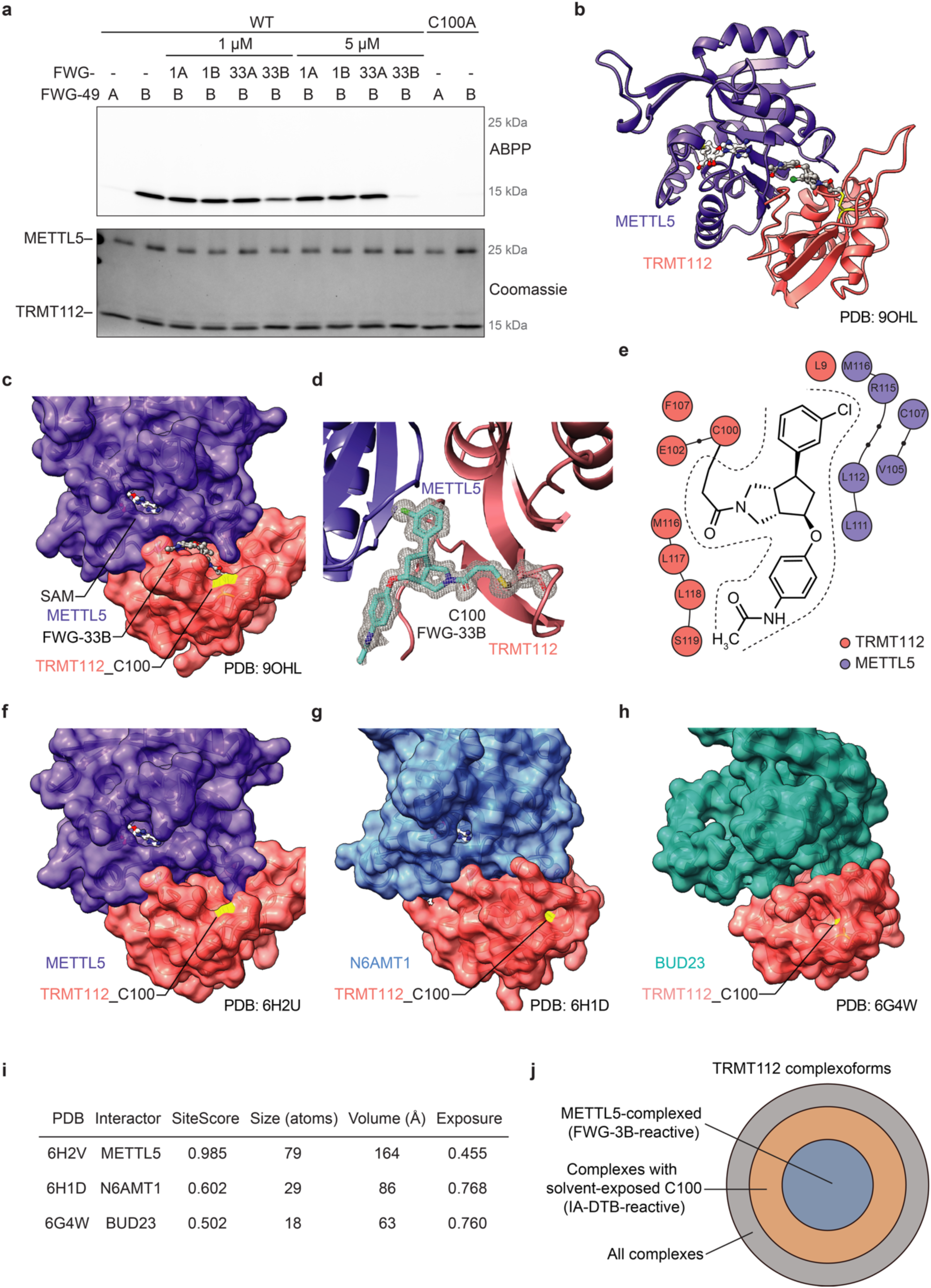
Crystal structure of a FWG-33B-modified TRMT112:METTL5 complex. **a**, Gel-ABPP data showing stereoselective and site-specific liganding of purified WT-TRMT112:WT-His-METTL5 with FWG-49B. Purified WT-TRMT112:WT-His-METTL5 complex (1 μM) was pre-treated with FWG-1A, 1B, 33A, or 33B (1 or 5 μM, 1 h), followed by FWG-49A or 49B (1 μM, 1 h), and analyzed by gel-ABPP (as described in **Fig. 1b**). Data are from a single experiment representative of two independent experiments. **b**, Co-crystal structure of the TRMT112-METTL5 complex with FWG-33B and SAM (PDB: 9OHL, 1.3 Å). **c**, Surface area representation of the TRMT112-METTL5 complex showing locating of FWG-33B and SAM (C100 shown in yellow)**. d**, *F*_O_*-F*_C_ omit map contoured at 3σ showing clear density for FWG-33B continuous with TRMT112_C100. **e**, Protein-ligand interaction diagram highlighting residues in TRMT112 and METTL5 that interact with FWG-33B. Amino acids within 4 Å of FWG-33B are shown (analyzed with Schrödinger Maestro). Small black dots represent amino acids in the chain that are not within 4 Å of FWG-33B. **f-h**, Crystal structures of an apo TRMT112:METTL5 complex (**f**; PDB: 6H2U), a TRMT112:N6AMT1 complex (**g**; PDB: 6H1D), and a TRMT112:BUD23 complex (**h**; PDB: 6G4W). **i**, Site mapping with Schrödinger Maestro near TRMT112_C100 reveals large differences in apparent solvent accessibility for the TRMT112:METTL5 complex compared to the TRMT112:N6AMT1 and TRMT112:BUD23 complexes. **j**, Concentric circle diagram showing TRMT112 complexoforms as relates to their potential reactivity with bicyclopyrrolidine acrylamides (e.g., FWG-3B) and/or IA-DTB.

We next determined the structure of the FWG-33B-liganded TRMT112:METTL5 complex in the presence of the SAM cofactor by X-ray crystallography. This 1.3 Å crystal structure (PDB: 9OHL; **Supplementary Table 1**) revealed that FWG-33B binds in a well-formed pocket residing at the interface of TRMT112 and METTL5 that is fully distinct from the SAM-binding pocket of METTL5 (∼22.5 Å distance between SAM (CH_3_) and FWG-33B (C6a)) (**Fig. 4b, c**). Clear electron density confirmed a covalent adduct between the acrylamide of FWG-33B and C100 of TRMT112 (**Fig. 4d**). FWG-33B interacts extensively with both TRMT112 (301 Å^2^ total surface area of interaction) and METTL5 (235 Å^2^ total surface area of interaction) that includes contacts with several residues from each protein (**Fig. 4e**), reflecting the composite nature of the pocket. A tight fit is observed for the 3-chlorophenyl group (C6) of FWG-33B, which interacts with hydrophobic residues V105, C107, L112, R115, and M116 of METTL5, as well as L9 and M116 of TRMT112 (**Fig. 4e** and **Extended Data Fig. 7b**). This series of interactions aligns with the steep SAR displayed by various C6 analogues (**Fig. 3b, c**). In contrast, the C4 appendage extends away from the binding pocket into a solvent-exposed region (**Extended Data Fig. 7b**), which is consistent with the limited impact of C4 modifications on bicyclopyrrolidine acrylamide engagement of the TRMT112:METTL5 complex.

A comparison of the structure of the FWG-33B-liganded TRMT112:METTL5 complex with previously reported apo structures of TRMT112:METTL5 (**Fig. 4f)**, TRMT112:N6AMT1 (**Fig. 4g**), and TRMT112:BUD23 (**Fig. 4h**) helped to explain the basis for exclusive reactivity of bicyclopyrrolidine acrylamides with TRMT112 when bound to METTL5, as neither the TRMT112:N6AMT1 or TRMT112:BUD23 complex displayed evidence of a solvent-accessible pocket in proximity to C100 of TRMT112. These differences appear to be at least partly explained by the location of a dynamic loop in close proximity to C100 (a.a. L9-F21 of TRMT112), which is open in apo TRMT112:METTL5 structures (**Extended Data Fig. 7d, e**), but closed in other TRMT112:MT structures (**Extended Data Fig. 7f-h**, though not fully resolved in all structures (**Extended Data Fig. 7f)**).

Site mapping performed in Schrödinger Maestro supported major differences in the apparent solvent accessibility near TRMT112_C100, with only the TRMT112:METTL5 complex having a well-defined predicted pocket (164 Å^3^, SiteScore 0.985; **Fig. 4i**). We speculate that these structural data not only rationalize the complexoform-restricted liganding of TRMT112 when bound to METTL5, but may also explain the discordance between our cysteine– and protein-directed ABPP data for C100 of TRMT112 (**Fig. 2a**). Indeed, it is possible that the IA-DTB probe used in cysteine-directed ABPP experiments can react with C100 in multiple TRMT112 complexoforms, while, in contrast, the alkynylated bicyclopyrrolidine acrylamides used in protein-directed ABPP experiments would be expected to exclusively engage TRMT112 when bound to METTL5. The resulting profiles would lead to maximal and sub-maximal engagement estimates for bicyclopyrrolidine acrylamide competitors in protein– and cysteine-directed ABPP experiments, respectively, with the latter matching the fraction of total IA-DTB-reactive TRMT112_C100 residing in TRMT112:METTL5 complexes (**Fig. 4j**).

### TRMT112 ligands allosterically potentiate METTL5 activity

A more detailed comparison of the structures of the apo and FWG-33B-liganded TRMT112:METTL5 complexes revealed a marked rearrangement in a flexible loop of METTL5 (L111-S117) near the SAM binding site (**Fig. 5a**). In the unliganded structures, S113-S117 forms a single turn helix that unfolds upon FWG-33B binding to place the ε carbon of METTL5_M116 and the β and γ carbons of METTL5_R115 against the chlorophenyl moiety. The extensive interactions between L111-S117 and FWG-33B, along with the hydrogen bond-mediated coordination of the adenine base of the SAM cofactor by the proximal residues D108 and V109, suggested that FWG-33B reactivity with TRMT112_C100 might allosterically modulate the catalytic activity of METTL5.

**Fig. 5.**
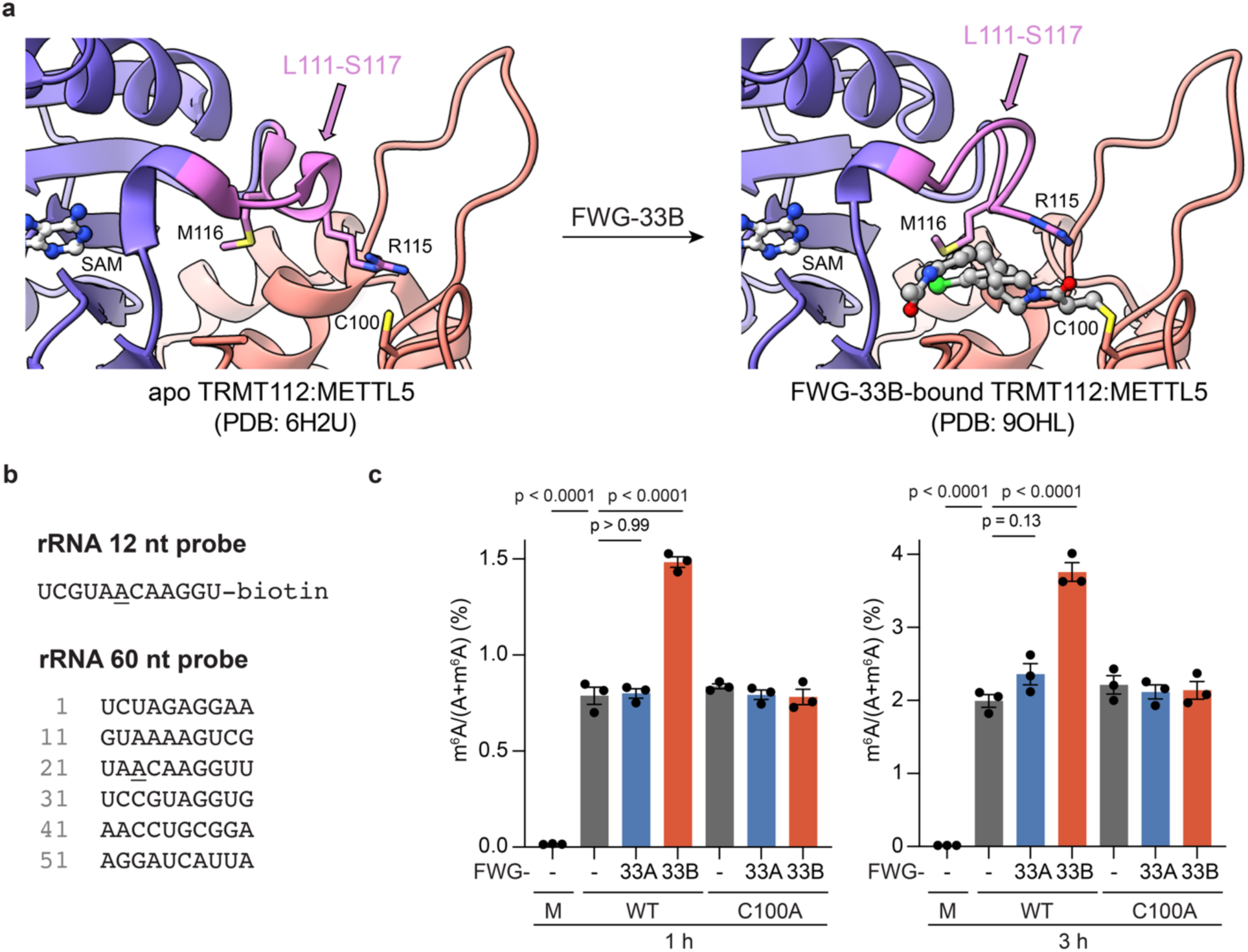
FWG-33B enhances the methyltransferase activity of METTL5. **a**, FWG-33B induces allosteric changes in the TRMT112:METTL5 structure, including movement of a flexible loop (colored in magenta, a.a. METTL5_L111-S117) and unwinding of single turn helix in proximity to the SAM binding site. **b**, Model RNA substrates (12 or 60 nucleotide (nt) containing the A_1832_ motif of the 18S rRNA (A_1832_ underlined). **c**, Effects of FWG-33B on the catalytic activity of WT-TRMT112:His-METTL5 and C100A-TRMT112: His-METTL5 complexes using a 12 nt substrate. WT-TRMT112:His-METTL5 and C100A-TRMT112:His-METTL5 complexes (20 μM) were treated with FWG-33A or FWG-33B (40 μM, 30 min) or DMSO in presence of SAM (1 mM). The samples were then diluted 20X into the METTL5 activity assay buffer (10 μM RNA, 1 mM SAM, 5 mM MgCl_2_, 50 mM NaCl, 50 mM TRIS, 1 mM DTT) for 1 or 3 h (37 °C), followed by digestion for the RNA substrates into single nucleosides and targeted (QQQ) LC-MS analysis. Data are from a single experiment containing three technical replicates and representative of two independent experiments. Data are average values ± s.e.m. normalized to DMSO for three independent biological experiments. Statistical significance was assessed using one-way analysis of variance (ANOVA).

The TRMT112:METTL5 complex catalyzes the *N*^6^-methyladenosine (m^6^A) modification at A_1832_ of the 18S small ribosomal subunit^44–46^. This modification occurs near the ribosomal decoding center, where it influences translation efficiency and regulates the translation of specific transcripts^44–46^. We tested the impact of FWG-33B on METTL5 activity using nucleotide single-stranded RNA substrates containing the UA<u>A</u>CA motif derived from the 18S rRNA where the underlined A corresponds to A_1832_ (**Fig 5b**)^46–48^. The purified WT-TRMT112 or C100A-TRMT112:METTL5 complexes (20 μΜ protein) were treated FWG-33A or FWG-33B (40 μΜ; 2:1 compound-to-protein ratio) or DMSO control for 30 min, after which the proteins were diluted 20-fold into the substrate assay containing 10 µM of RNA substrate and 1 mM SAM and, after 1 or 3 h, the methylated RNA product quantified by measuring the ratio of adenosine (A) and m^6^A by LC-MS as described previously^46,47^. The DMSO control experiments confirmed that both WT-TRMT112:METTL5 and C100A-TRMT112:METTL5 complexes exhibited similar time-dependent methylation activities with the short (12-nucleotide) or long (60-nucleotide) substrate (**Fig. 5c** and **Extended Data Fig. 8a-c**). These experiments revealed a ∼two-fold increase in methylation activity for the FWG-33B-treated WT-TRMT112:METTL5 complex with either RNA substrate that was both stereoselective (not observed with FWG-33A) and site-specific (not observed with the C100A-TRMT112:METTL5 complex) (**Fig. 5c** and **Extended Data Fig. 8b, c**).

METTL5 has been reported to display a *K*_m_ value for SAM of ∼1 µM^48^, which is well below the concentration used in our enzymatic assays (1 mM). The FWG-33B-induced enhancement of METTL5 activity therefore seems unlikely to be related to changes in SAM binding affinity and might instead reflect other impacts on the METTL5 mechanism (e.g., FWG-33B-induced changes in product release). Regardless, our combined structural data and enzymatic assays support that covalent binding of FWG-33B to an interface pocket of the TRMT112:METTL5 complex leads to allosteric enhancement of METTL5 catalytic activity, and thus provides, to our knowledge, the first chemical tool for studying the potentiation of METTL5 activity in biological systems.

## Discussion

The massive structural and functional diversity conferred by proteoforms presents an exciting opportunity for chemical biology and drug discovery^1,7,49–51^. Small molecules that target specific proteoforms have the potential to impact cell and human biology with a level of precision that exceeds the complete genetic or pharmacological disruption of proteins (and consequently all of their respective proteoforms). While in select instances a specific proteoform of interest may be amenable to production (e.g., semisynthesis^52^), purification, and screening, the general pursuit of proteoform-restricted ligands would benefit from methodologies that can operate in native biological systems where a more complete range of post-translational protein states can be assayed. Here, we have shown that chemical proteomics offers a compelling strategy for identifying covalent ligands that target a single complexation state (or complexoform^5,6^) of the general MT adaptor TRMT112.

The discordant profiles of TRMT112 in cysteine– and protein-directed ABPP datasets – incomplete and complete liganding, respectively – provided the first clue that bicyclopyrrolidine acrylamide stereoprobes engage this protein in an atypical manner. The extent to which such discrepancies can serve as a general hallmark of proteoform-restricted liganding events will depend on the depth of coverage in cysteine-directed ABPP datasets, as alternative explanations, such as the failure to detect the relevant cysteine for a liganding event mapped exclusively by protein-directed ABPP, remain possible. Nonetheless, we believe that some of the additional experimental approaches deployed in this study may prove generally useful for mapping complexoform-restricted small molecule interactions. For instance, we envision that integrating ABPP with abundance-based proteomics of SEC fractions from cells treated with electrophilic stereoprobes may provide a robust way to identify liganding events that exclusively occur with rare complexoforms of proteins. Combining ABPP with top-down proteomics^5,53^ or other global measurements of protein modifications (e.g., phosphoproteomics^7,54^) may also provide complementary approaches for relating covalent liganding events to individual proteoforms in cells.

The role of METTL5-mediated 18S rRNA m^6^A_1832_ modification in regulating protein translation and downstream cellular processes is under active investigation in fields ranging from neuroscience^55,56^ to metabolism^57^ to cancer^57,58^. Deleterious mutations in METTL5 lead to intellectual disability and microcephaly in humans^59^, pointing to an important role for this enzyme in brain development. Chemical probes, however, are lacking for METTL5, and we therefore believe the allosteric agonists reported herein should offer valuable tools for studying the functions of 18S rRNA m^6^A_1832_ modification in diverse cellular processes. We should note, however, that the basal level of 18S rRNA m^6^A_1832_ modification is quite high (> 98%) in immortalized cells grown under standard cell culture conditions ^45,57,60^, which indicates that alternative cell models may be needed to study the impact of METTL5 agonism on ribosome function and protein translation. Some studies suggest that METTL5 may also regulate the methylation of mRNAs^61^ and evaluating the effects of FWG-33B on the global m^6^A profiles of cells^62–64^ may help to establish the full substrate scope of METTL5. Considering that allosteric sites on other proteins have been found to support both small molecule-mediated agonism and antagonism^65,66^, we also wonder if the TRMT112:METTL5 composite pocket may serve as a future source for METTL5 inhibitors.

Small molecules that induce protein-protein interactions through the formation of ternary complexes, or molecular glues, have become a major source of chemical probes and drugs^67^. Such induced proximity mechanisms can strengthen *dynamic* protein-protein interactions or promote novel ones. Our findings highlight an alternative and complementary mechanism for small molecule regulation of *stable* protein complexes. By binding at a composite pocket unique to the interface of TRMT112 and METTL5, FWG-33B not only achieves high selectivity for a single MT partner of TRMT112, but also causes allosteric effects that increase the methylation activity of METTL5. This type of “proximity-induced” liganding event was recently observed for allosteric inhibitors of the CCNE1:CDK2 complex, which bind to a composite pocket distal from the CDK2 active site and similarly display complexoform-restricted activity (in this case over other cyclin:CDK2 complexes, such as CCNE2:CDK2)^68^. We do not yet understand the frequency of allosteric composite pockets across the human proteome, but the cases reported to date underscore the importance of considering each complexoform as an independent source.

Looking forward, we believe that additional general insights can be drawn from the overall chemical proteomic profiles of the bicyclopyrrolidine acrylamides. These compounds, despite exhibiting lower intrinsic reactivity compared to previously described azetidine and tryptoline acrylamide stereoprobes, nonetheless engaged several unique proteins in human cells (**Table 1**). Our results, alongside studies of additional classes of lower reactivity stereoprobes^21^, thus emphasize the value of scaffold diversity in the design of electrophilic compound libraries. Toward this end, the catalytic stereoselective synthetic strategy described for the bicyclopyrrolidine acrylamides may offer a useful approach for scaffold diversification compared to past routes for stereoprobe construction, which mostly leveraged enantiopure starting materials^18,20,23^. We also acknowledge that our initial chemical proteomic datasets were generated with one cell line (Ramos cells) in a basal state and may therefore overlook liganding events that occur with dynamic or regulated proteoforms. Future studies of other cell types and cell states may uncover additional context-dependent proteoform liganding events and, through doing so, further expand the toolbox of chemistries that can perturb biological systems with high precision.

## Data Availability

Proteomic data are available via ProteomeXchange with identifier PXD063358^69^. The atomic coordinates and structure factors have been deposited in the Protein Data Bank, www.pdb.org (PDB ID code 6HOL).

## Supporting information

Supplementary Biology Information

Supplementary Chemistry Information

PDB Validation Report

## Acknowledgements.

This work was supported by the NCI (R35 CA231991 for B. F. C). F. W. G. and B. F. C thank the George E. Hewitt Foundation for Medical Research for a Postdoctoral Fellowship. We are grateful to Dr. G. J. Kroon and Dr. L. Pasternack (Scripps Research; NMR Facility) for NMR spectroscopic assistance; Quynh Nguyen Wong and Jason Lee (Scripps Research; Automated Synthesis Facility) for HRMS measurements and SFC separations; and WuXi AppTec for the synthesis of (±)-1. X-ray diffraction data were collected at beamline 5.0.2 at the Advanced Light Source. The Berkeley Center for Structural Biology is supported by the Howard Hughes Medical Institute, Participating Research Team members, and the National Institutes of Health, National Institute of General Medical Sciences, ALS-ENABLE grant P30 GM124169. The Advanced Light Source is a Department of Energy Office of Science User Facility under Contract No. DE-AC02-05CH11231.

---

**Extended Data Fig. 1.**
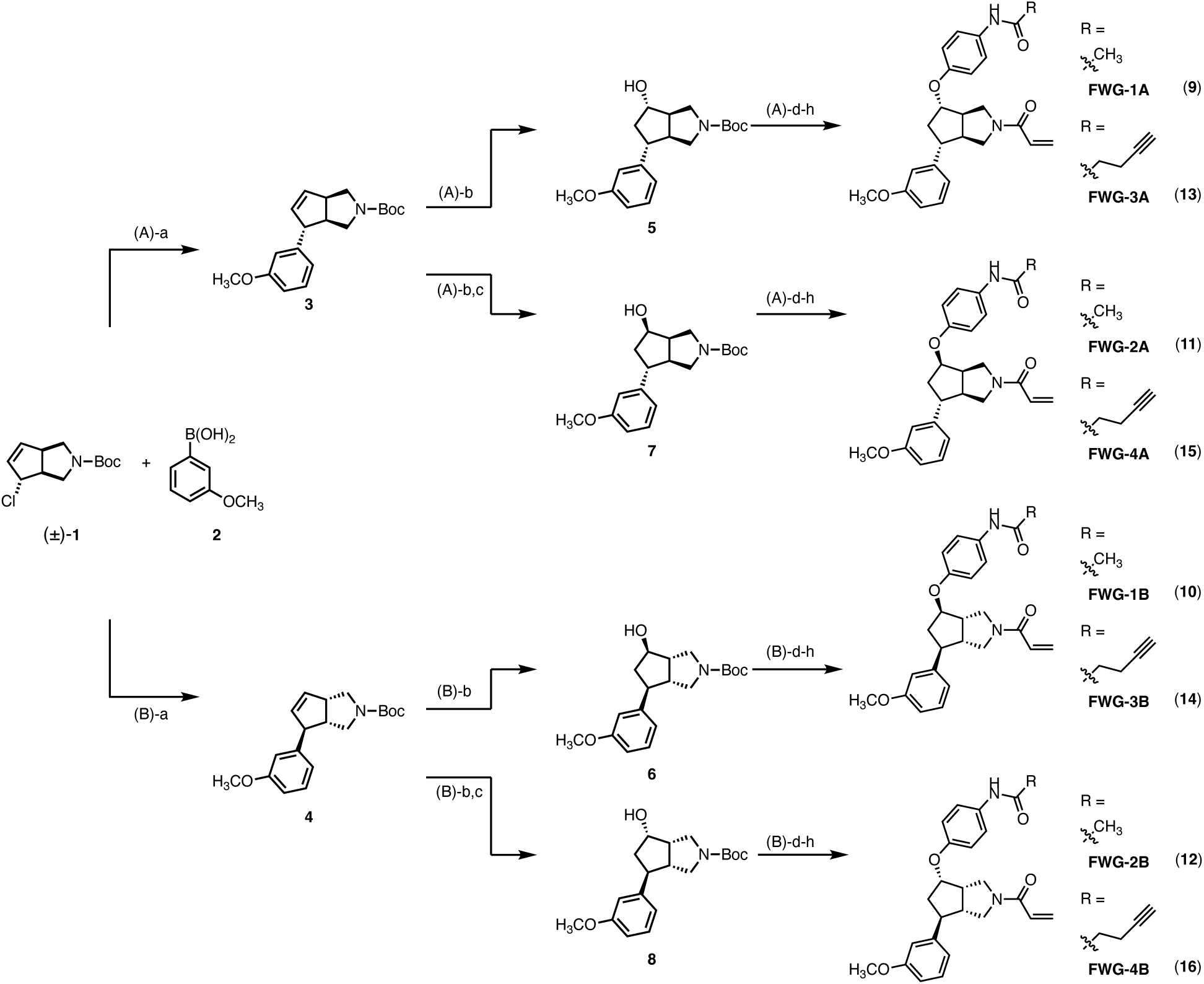
Scheme for the catalytic stereodivergent synthesis of bicyclopyrrolidine stereoprobes. *General conditions* (representative for **FWG-3A** and **FWG-4A**; also see **Supporting Synthetic Chemistry Information**): a. 3-H_3_CO-Ph-B(OH)_2_, aq. CsOH, (*S*)-SEGPHOS (6 mol%), [Rh(cod)OH]_2_ (2.5 mol%), THF/H_2_O, 65 °C, overnight, 75%, 99% ee; b. HBpin, Cs_2_CO_3_, [Rh(PPh_3_)_3_Cl] (10 mol%), THF, r.t., overnight, then H_2_O_2_, aq. NaOH, 0 °C, 1 h, 51%; c. (1) 4-NO_2_-Ph-CO_2_H, PPh_3_, DIAD, THF, 0 °C to r.t., overnight, (2) K_2_CO_3_, MeOH/H_2_O, r.t., overnight, 68%; for FWG-3A: d. 4-NO_2_-Ph-F, KHMDS, PhCH_3_/DMF, 0 °C to r.t., overnight, 80%; e. H_2_, Pd/C, MeOH, 3 h, r.t., 88%; f. HC≡C(CH)_2_CO_2_H, HATU, DIPEA, CH_2_Cl_2_, r.t., overnight (no purification); g. TFA, CH_2_Cl_2_, r.t., 1 h (no purification); h. acryloyl chloride, DIPEA, CH_2_Cl_2_, r.t., 2 h, 39% (over 3 steps); for FWG-4A: d. 4-NO_2_-Ph-F, KHMDS, PhCH_3_/DMF, 0 °C to r.t., overnight, 33%; e. H_2_, Pd/C, MeOH, 3 h, r.t., 88%; f. HC≡C(CH)_2_CO_2_H, HATU, DIPEA, CH_2_Cl_2_, r.t., overnight (no purification); g. TFA, CH_2_Cl_2_, r.t., 1 h (no purification); h. acryloyl chloride, DIPEA, EtOAc, r.t., 2 h, 64% (over 3 steps). *Abbreviations:* cod = 1,5-cyclooctadiene; HATU = hexafluorophosphate azabenzotriazole tetramethyl uranium; DIAD = diisopropyl azodicarboxylate; DIPEA = *N,N*-diisopropylethylamine; DMF = *N,N*-dimethylformamide; KHMDS = potassium bis(trimethylsilyl)amide; pin = pinacolato; TFA = trifluoroacetic acid; THF = tetrahydrofuran.

**Extended Data Fig. 2.**
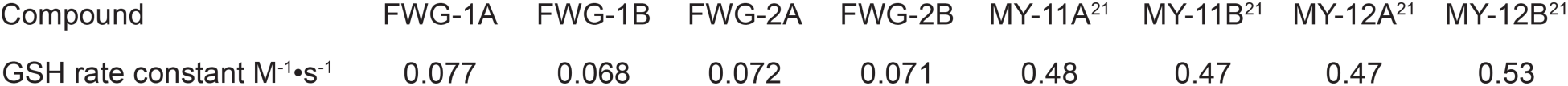
Glutathione (GSH) reactivity measurements for FWG-1A/B and FWG-2A/B compared to previously described azetidine acrylamides MY-11A/B and MY-12A/B^21,30^.

**Extended Data Fig. 3.**
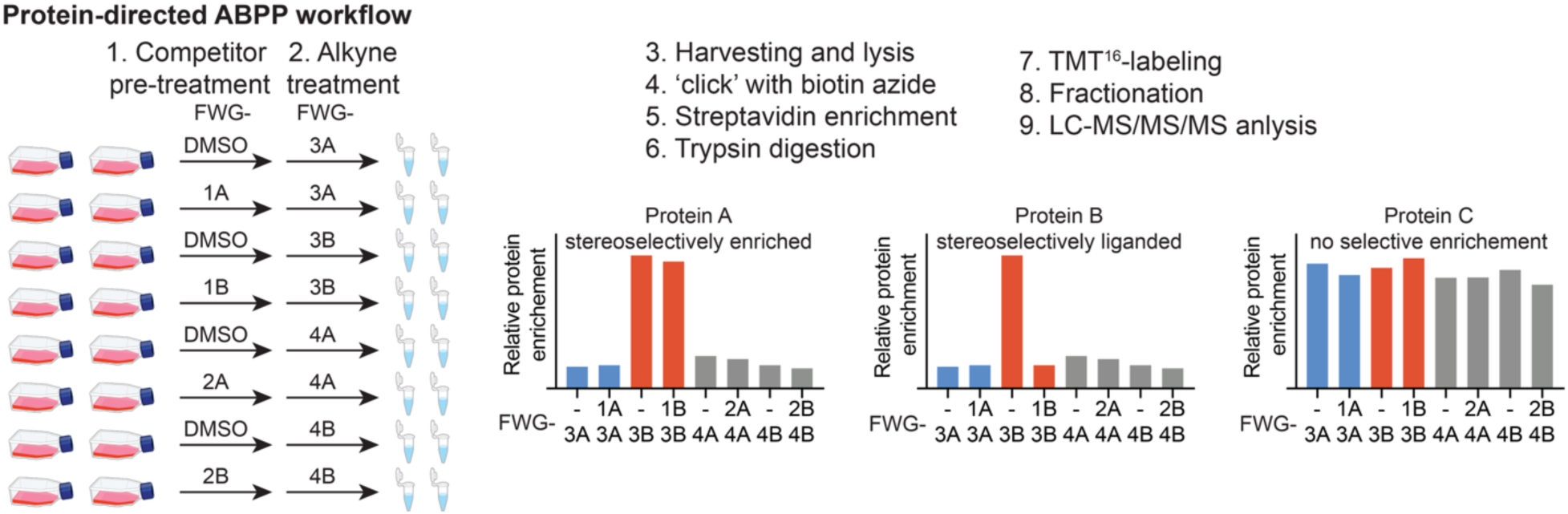
General workflow for quantifying stereoselectively liganded proteins in protein-directed ABPP performed with bicyclopyrrolidine acrylamide stereoprobes. Multiplexed format (TMT^16plex^) for protein-directed ABPP experiments (performed as described previously^23^) for quantifying stereoselective enrichment of proteins by alkyne stereoprobes FWG-3A, 3B, 4A, and 4B, and the competitive blockade of this enrichment by non-alkynes FWG-1A, 1B, 2A and 2B. Two replicates of each group are generated per TMT^16plex^ experiment. Examples of proteins showing stereoselectively enrichment, but not competition (protein A), stereoselective enrichment and competition (protein B), and neither stereoselective enrichment or competition (protein C).

**Extended Data Fig. 4.**
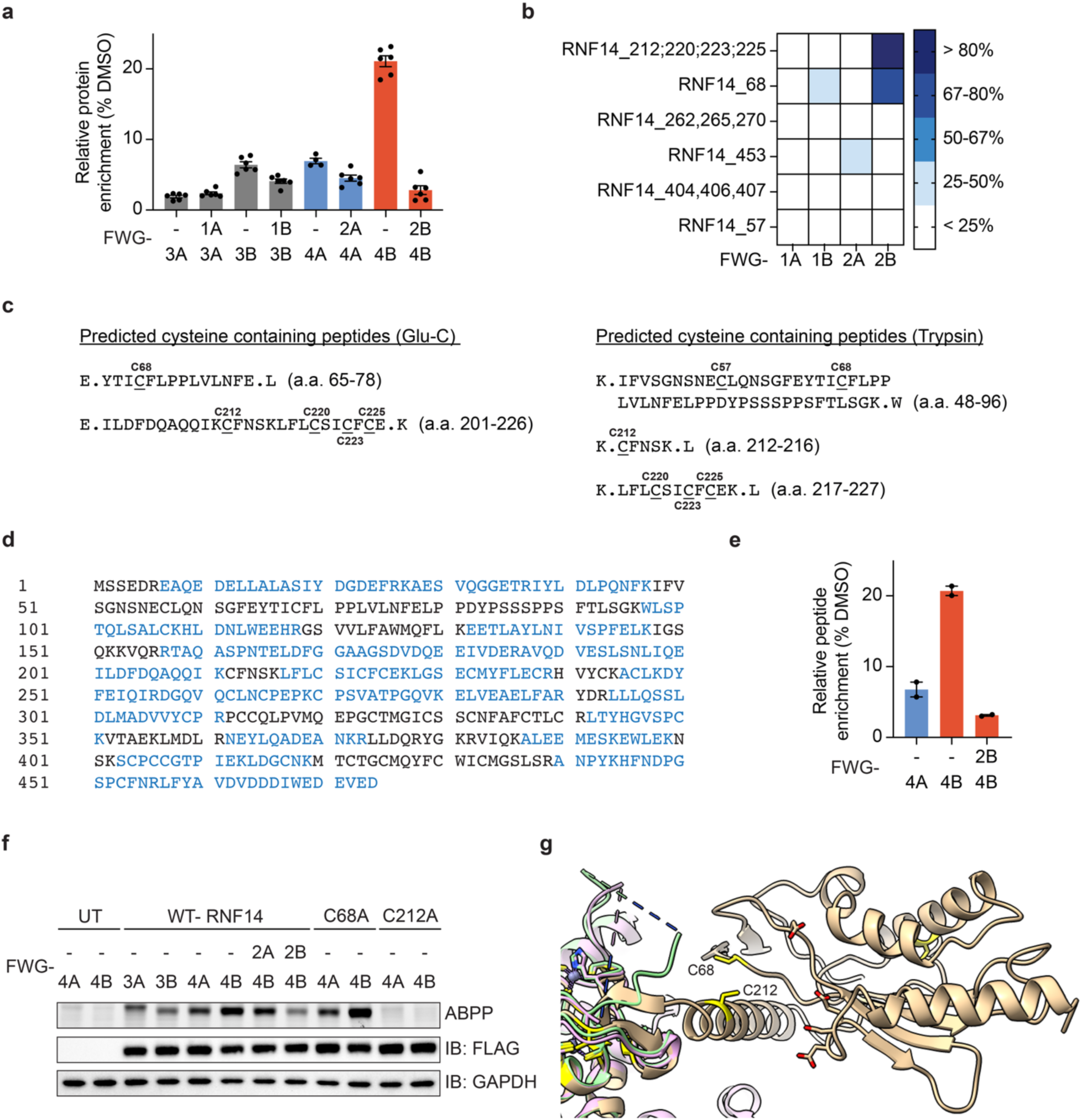
Mapping cysteine-212 (C212) as the site of bicyclopyrrolidine acrylamide engagement in RNF14. **a**, Bar graph showing protein-directed ABPP data for RNF14 (from experiments summarized in **Fig. 1c**). Data are average values ± s.e.m. for four-six independent biological experiments. **b**, Heatmap showing RNF14 peptides from cysteine-directed ABPP experiments performed on HEK293T cells recombinantly expressing RNF14 and using a Glu-C protease digestion protocol. The RNF14_C212,220,223,225 and RNF14_C68 peptides showed stereoselective blockade of IA-DTB enrichment by FWG-2B. HEK293T cells were transfected with WT-RNF14, treated with FWG-1A, 1B, 2A, and 2B (50 μM, 3 h), followed by lysis and treatment with IA-DTB (100 μM, 1 h) and cysteine-directed ABPP with Glu-C digestion. Data are average values ± s.e.m. for two independent biological experiments. **c**, Predicted RNF14 peptides containing C212,220,223,225 and C68 from Glu-C and trypsin digestion. **d**. Amino acid sequence of RNF14 highlighting in blue the tryptic peptide sequences that were quantified by protein-directed ABPP experiments performed on HEK293T cells recombinantly expressing WT-RNF14 and treated with FWG-1A, 1B, 2A, and 2B (50 μM, 3 h), followed by stereomatched alkynes FWG-3A, 3B, 4A, and 4B (10 μM,1 h). Data are from two independent biological experiments. **e**, Stereoselective enrichment of the RNF14_220,223,225 tryptic peptide in FWG-4B-treated cells from protein-directed ABPP experiments performed on HEK293T cells recombinantly expressing WT-RNF14 (for treatment conditions see **Extended Data Fig. 4d**). Data are average values ± s.e.m. for two independent biological experiments. **f**, Gel-ABPP data demonstrating stereoselective and site-specific engagement of recombinant WT-RNF14-FLAG and C68A-RNF14-FLAG, but not C212A-RNF14-FLAG by FWG-4B. HEK293T cells expressing the indicated recombinant RNF14 proteins were pre-treated with non-alkyne competitors FWG-2A and 2B (50 μM, 3 h), followed by FWG-3A, 3B, 4A, and 4B (10 μM, 1 h), and analyzed by gel-ABPP (as described in **Fig. 1b**). Data are from a single experiment representative of two independent experiments. UT = untransfected. **g**, AlphaFold model of RNF14 predicting proximity between C68 and C212 (8 Å). RNF14_C212 is on the N-terminal side of the RNF14 RBR/TRIAD supradomain. The alignment of the TRIAD supradomain of RNF14 with determined TRIAD1 structures in green and pink (PDB: 7ONI and PDB: 7OD1).

**Extended Data Fig. 5.**
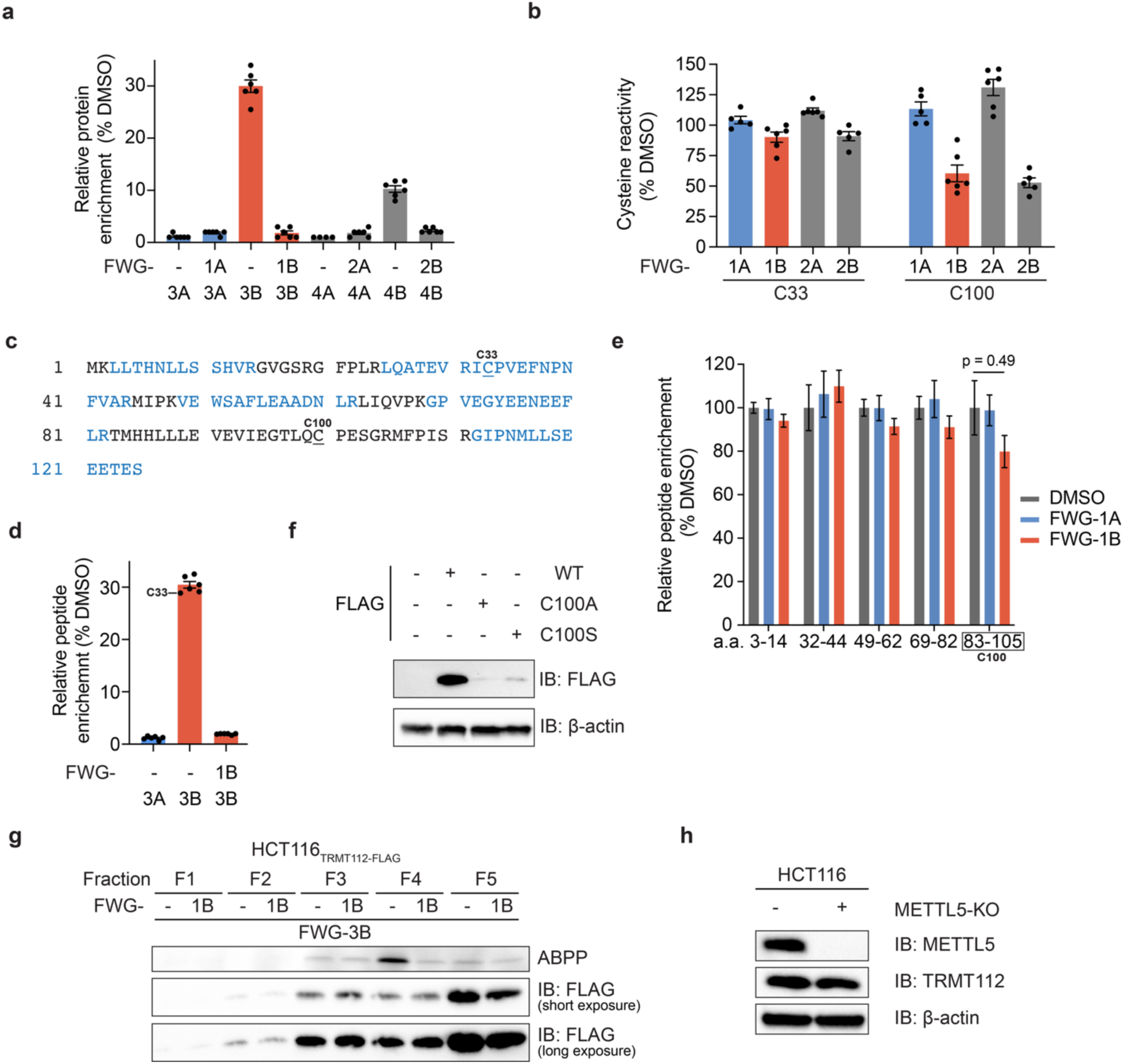
Biyclopyrrolidine acrylamides engage a TRMT112:METTL5 complexoform in cells. **a**, Bar graph showing protein-directed ABPP data for TRMT112. Data are from Ramos cells treated with non-alkyne competitors FWG-1A, 1B, 2A, and 2B (50 μM, 3 h), followed by stereomatched alkynes FWG-3A, 3B, 4A, and 4B (10 μM,1 h) and protein-directed ABPP analysis. Data are average values ± s.e.m. for four-six independent biological experiments. **b**, Bar graph showing cysteine-directed ABPP for TRMT112_C33 and C100. Data are from Ramos cells treated with FWG-1A, 1B, 2A, and FWG-2B (50 μM, 3 h), followed by lysis and treatment with IA-DTB (100 μM,1 h) and cysteine-directed ABPP analysis. Data are average values ± s.e.m. for five-six independent biological experiments. **c**, Amino acid sequence of TRMT112. Amino acids in blue were quantified by protein-directed ABPP. **d**, Quantification of TRMT112 tryptic peptides, including the C33 containing peptide by protein-directed ABPP. **e**, Quantification of the indicated tryptic peptides for TRMT112 from anti-FLAG IP-MS experiments performed with HCT116_TRMT112-FLAG_ cells treated with DMSO or FWG-1A or FWG-1B (50 µM, 4 h). Minimal loss of the unmodified tryptic peptide containing C100 was observed in FWG-1B-treated cells, consistent with only a small fraction of total recombinant TRMT112 reacting with FWG-1B. Data are average values ± s.e.m. normalized to DMSO for eight independent biological experiments. Statistical significance between the DMSO and the FWG-1B treated peptides was assessed using unpaired-multiple *t*-tests (Holm-Šídák approach to multiple comparisons; reported p values are adjusted p values). **f**, Western blot showing negligible expression of C100A– and C100S-TRMT112 mutants relative to WT-TRMT112 in HCT116 cells. β-actin shown as a loading control. Data are from a single experiment representative of two independent experiments. **g**, Gel-ABPP demonstrating blockade of the FWG-3B-TRMT112 interaction in SEC fraction 4 by pre-treatment with FWG-1B. HCT116_TRMT112-FLAG_ cells were pre-treated with DMSO and FWG-1B (50 μM, 3 h) and then FWG-3B (10 μM, 1 h), followed by the workflow presented in **Fig. 3a**. Data are from a single experiment representative of two independent experiments. **h**, Western blot showing similar expression of TRMT112 in parental vs METTL5-KO HCT116 cells. β-actin shown as a loading control. Data are from a single experiment representative of two independent experiments.

**Extended Data Fig. 6.**
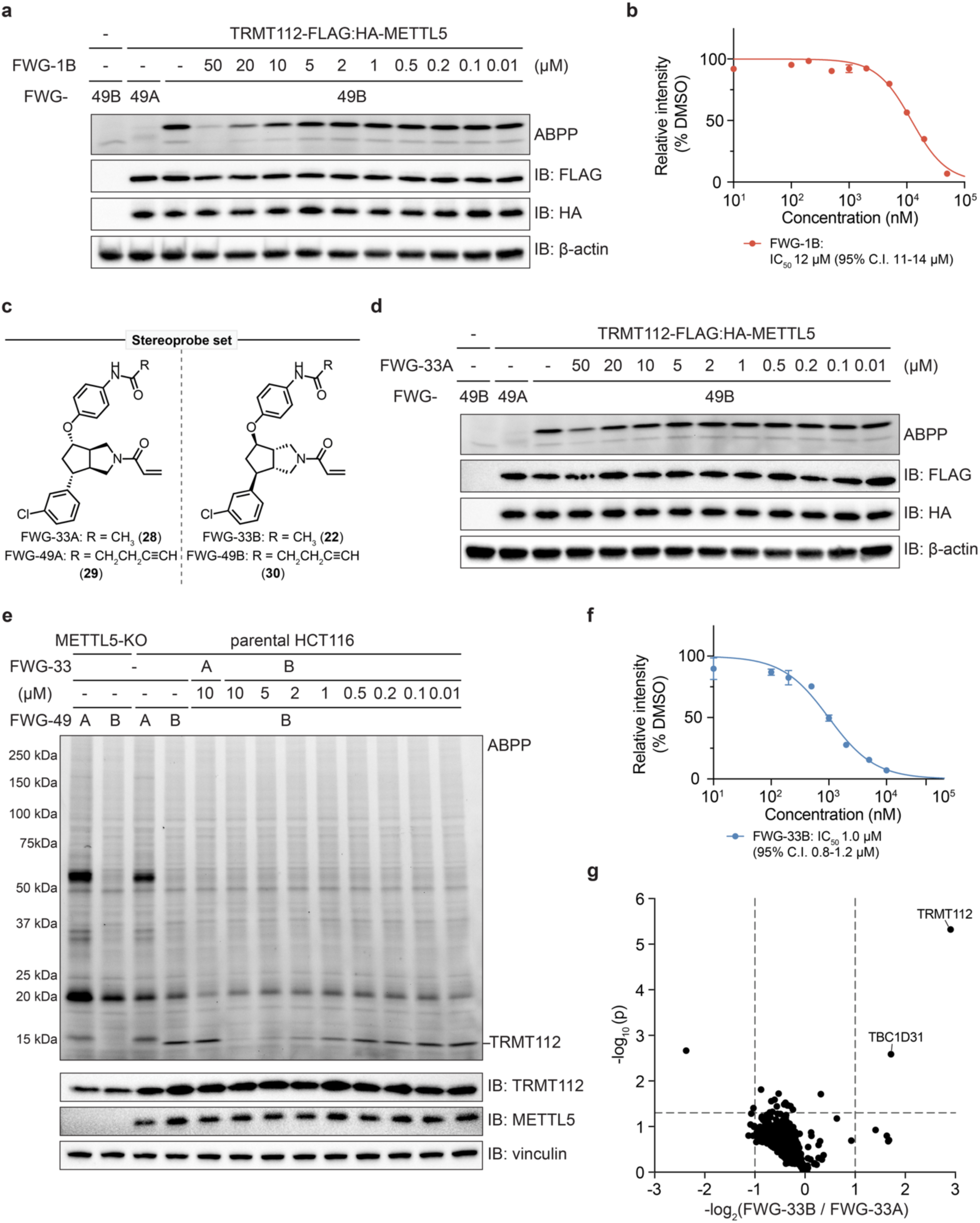
Optimization of bicyclopyrrolidine acrylamide stereoprobes targeting the TRMT112:METTL5 complex. **a**, Gel-ABPP data showing concentration-dependent blockade of the FWG-49B-TRMT112 interaction by FWG-1B. HCT116_TRMT112-FLAG:HA-METTL5_ cells were pre-treated with the indicated concentrations of FWG-1B for 3 h, followed by FWG-49A and 49B (2 μM, 1 h), and gel-ABPP analysis. Data are from a single experiment representative of two independent experiments. **b**, Quantification of gel-ABPP data from panel **a** for engagement of TRMT112 by FWG-1B in HCT116_TRMT112-FLAG:HA-METTL5_ cells. Data are average values ± s.e.m. for two independent biological experiments. **c**, Structures of non-alkyne (FWG-33A (**28**), FWG-33B (**22**)) and alkyne (FWG-49A (**29**), FWG-33B (**30**)) bicyclopyrrolidine acrylamide stereoprobes. **d**, Gel-ABPP data showing concentration-dependent blockade of the FWG-49B-TRMT112 interaction by FWG-33A. HCT116_TRMT112-FLAG:HA-METTL5_ cells were pre-treated with the indicated concentrations of FWG-33A for 3 h, followed by FWG-49A and 49B (2 μM, 1 h), and gel-ABPP analysis. Data are from a single experiment representative of two independent experiments. **e**, Gel-ABPP data showing METTL5-dependent stereoselective liganding of TRMT112. Parental HCT116 cells or METTL5-KO cells were treated with FWG-33A (10 μM, 3 h) or 33B (10 to 0.01 μM, 3 h), followed by FWG-49A and 49B (2 μM, 1 h), and analyzed by gel-ABPP (as described in **Fig. 1b**). Data are from a single experiment representative of two independent experiments. **f**, Quantification of gel-ABPP data from panel **e** for engagement of endogenous TRMT112 by FWG-33B in parental HCT116 cells. Data are average values ± s.e.m. for two independent biological experiments. **g**, Volcano plot showing protein-directed ABPP for blockade of FWG-49B enrichment of proteins by FWG-33B versus FWG-33A (same protein-directed ABPP experiment as shown in **Fig. 3f**). Data are average values for three-four independent biological experiments. Statistical significance was assessed using Welch two sample *t*-tests.

**Extended Data Fig. 7.**
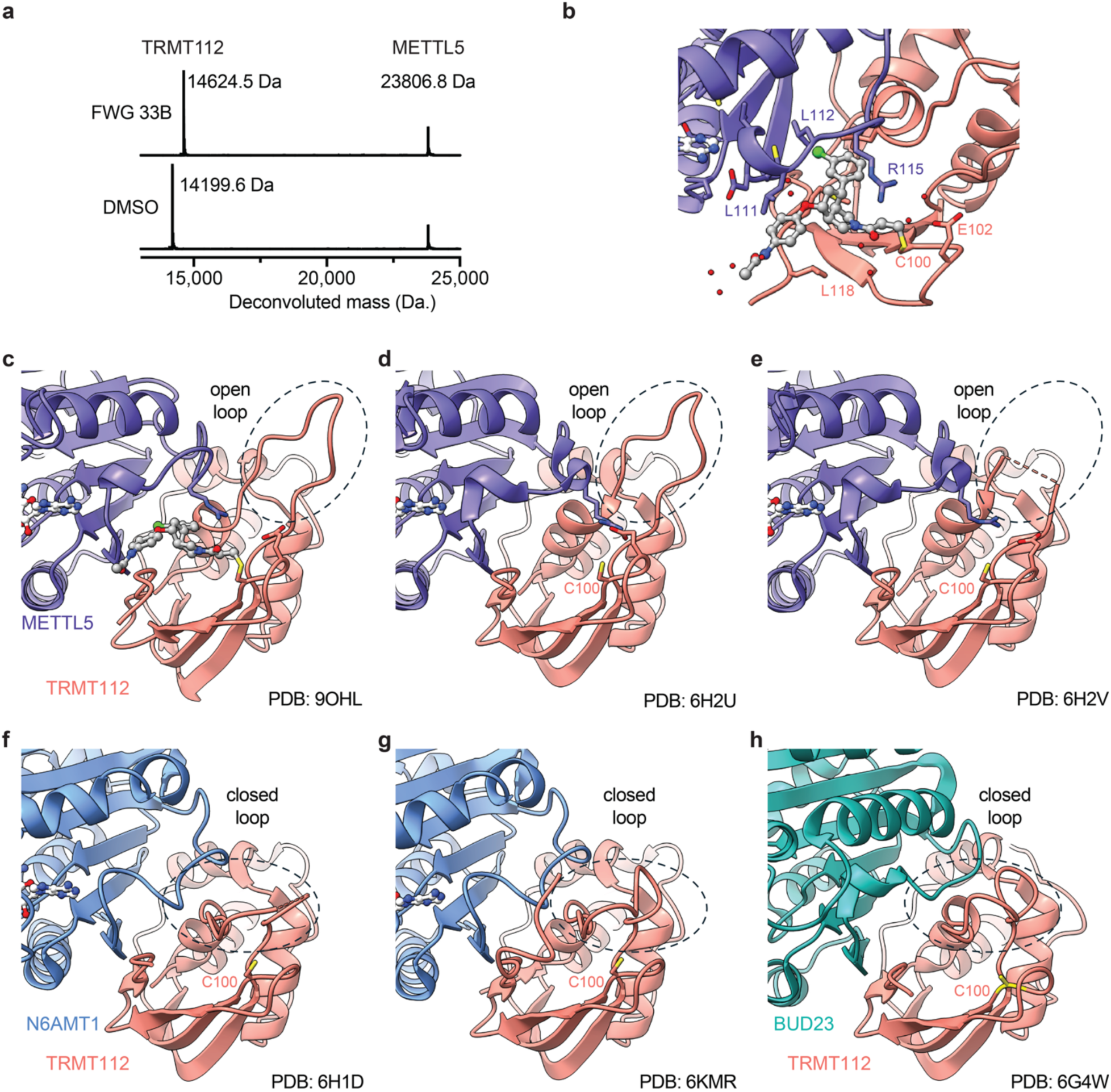
Crystal structure of an FWG-33B-modified TRMT112:METTL5 complex. **a**, Intact MS showing complete modification of TRMT112 by FWG-33B. Purified WT-TRMT112:WT-METTL5 complex (20 μM) treated with FWG-33B (100 μM, 30 min) in presence of SAM (1 mM), and analyzed by Q-TOF MS. Data are from a single experiment. **b**, Key residues in the FWG-33B binding site. Blue, METTL5; salmon, TRMT112. **c-h**, TRMT112 structures in TRMT112:METTL5 (**c**-**e**), TRMT112:N6AMT1 (**f**-**g**) and TRMT112:BUD23 (**h**) showing mostly good alignment. A dynamic loop (black circle, TRMT112: a.a. L9-F21) in proximity to C100 is highlighted to show its open conformation in apo TRMT112:METTL5 structures (PDB 6H2U (**d**) and in PDB 6H2V (**e**)) and closed conformation in other TRMT112:MT structures (TRMT112:N6AMT1: PDB 6H1D (**f**) and PDB 6KMR (**g**); TRMT112:BUD23: PDB 6G4W (**h**)).

**Extended Data Fig. 8.**
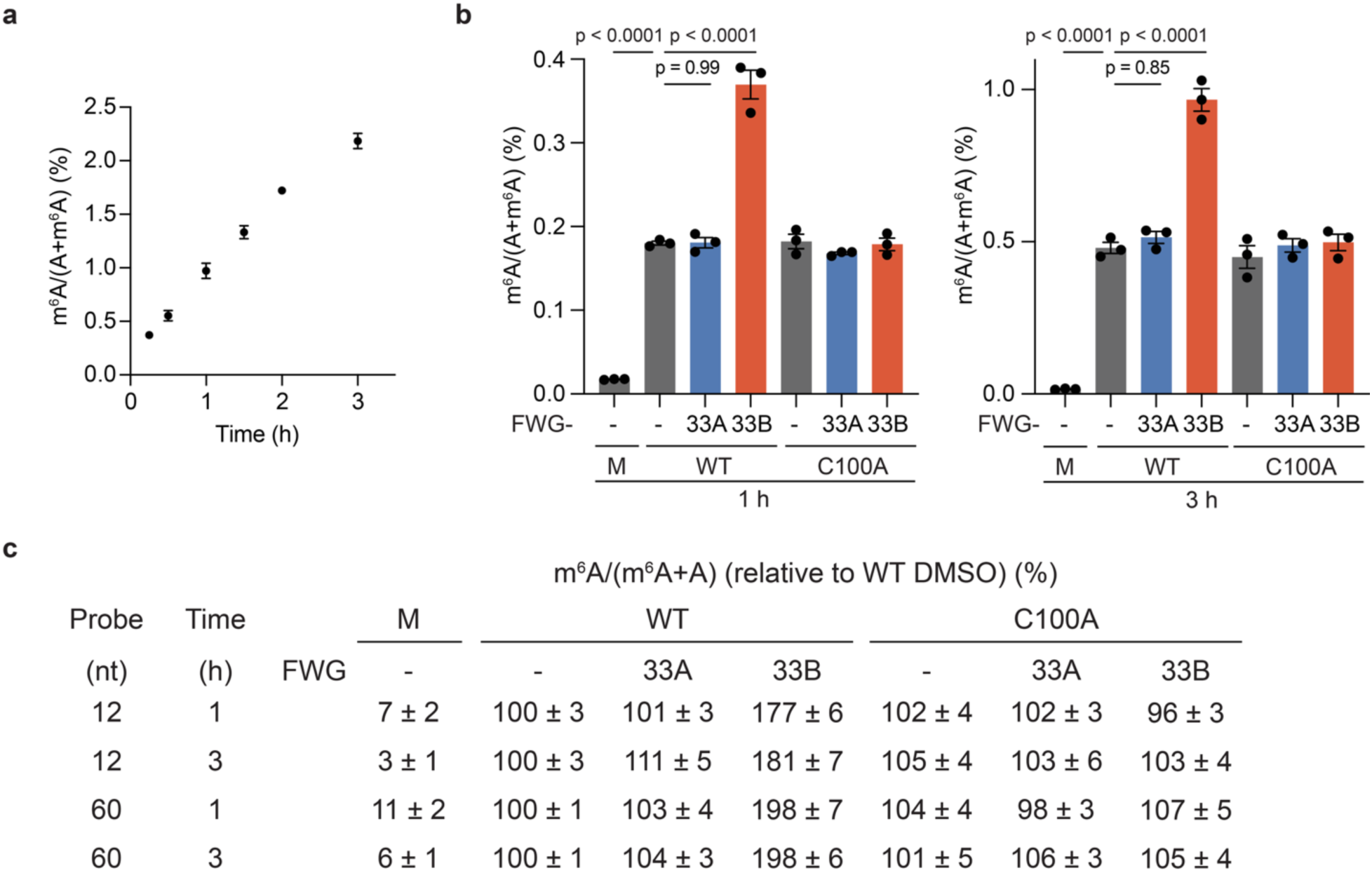
FWG-33B enhances the methyltransferase activity of METTL5. **a**, Time-dependence of the methyltransferase activity of the WT-TRMT112:His-METTL5 complex using a 12 nt RNA substrate. Samples were processed and analyzed as described in **Fig. 5c**. Data are average values ± s.e.m. for three independent biological experiments. **b**, Effects of FWG-33B on the catalytic activity of WT-TRMT112:His-METTL5 and C100A-TRMT112:His-METTL5 complexes using a 60 nt substrate. WT-TRMT112:His-METTL5 and C100A-TRMT112:His-METTL5 complexes (20 μM) were treated with FWG-33A or FWG-33B (40 μM, 30 min) or DMSO in presence of SAM (1 mM). The samples were then diluted 20X into the METTL5 activity assay buffer (10 μM RNA, 1 mM SAM, 5 mM MgCl_2_, 50 mM NaCl, 50 mM TRIS, 1 mM DTT) for 1 or 3 h (37 °C), followed by digestion for the RNA substrates into single nucleosides and targeted (QQQ) LC-MS analysis. Data are from a single experiment containing three technical replicates and representative of two independent experiments. Data are average values ± s.e.m. normalized to DMSO for three independent biological experiments. Statistical significance was assessed using one-way analysis of variance (ANOVA). **c**, Table summarizing METT5 activity data normalized to the DMSO-treated WT-TRMT112:METTL5 condition. Data are average values ± s.e.m. for two experiments with each three technical replicates.

## References

1. Aebersold, R. et al. How many human proteoforms are there? Nat. Chem. Biol. 14, 206–214 (2018).

2. Stumpf, M. P. H. et al. Estimating the size of the human interactome. Proc. Natl. Acad. Sci. U.S.A. 105, 6959–6964 (2008).

3. Lu, H. et al. Recent advances in the development of protein–protein interactions modulators: mechanisms and clinical trials. Signal Transduct. Target. Ther. 5, 213 (2020).

4. Smith, L. M. et al. The Human Proteoform Project: Defining the human proteome. Sci. Adv. 7, eabk0734 (2021).

5. Gomes, F. P. et al. Native top-down proteomics enables discovery in endocrine-resistant breast cancer. Nat. Chem. Biol. (2025) doi:10.1038/s41589-025-01866-8.

6. Jensen, M. H., Morris, E. J., Tran, H., Nash, M. A. & Tan, C. Stochastic ordering of complexoform protein assembly by genetic circuits. PLoS Comput. Biol. 16, e1007997 (2020).

7. Kemper, E. K., Zhang, Y., Dix, M. M. & Cravatt, B. F. Global profiling of phosphorylation-dependent changes in cysteine reactivity. Nat. Methods 19, 341– 352 (2022).

8. Castellón, J. O. et al. Chemoproteomics Identifies State-Dependent and Proteoform-Selective Caspase-2 Inhibitors. J. Am. Chem. Soc. 146, 14972–14988 (2024).

9. Backus, K. M. et al. Proteome-wide covalent ligand discovery in native biological systems. Nature 534, 570–574 (2016).

10. Shaw, T. I. et al. Multi-omics approach to identifying isoform variants as therapeutic targets in cancer patients. Front. Oncol. 12, 1051487 (2022).

11. Nuti, E. et al. Bivalent Inhibitor with Selectivity for Trimeric MMP-9 Amplifies Neutrophil Chemotaxis and Enables Functional Studies on MMP-9 Proteoforms. Cells 9, 1634 (2020).

12. Gil, G. et al. Proteoform-Specific Protein Binding of Small Molecules in Complex Matrices. ACS Chem. Biol. 12, 389–397 (2017).

13. Niphakis, M. J. & Cravatt, B. F. Ligand discovery by activity-based protein profiling. Cell Chem. Biol. 31, 1636–1651 (2024).

14. Niphakis, M. J. & Cravatt, B. F. Enzyme inhibitor discovery by activity-based protein profiling. Annu. Rev. Biochem. 83, 341–77 (2014).

15. Spradlin, J. N., Zhang, E. & Nomura, D. K. Reimagining Druggability Using Chemoproteomic Platforms. Acc. Chem. Res. 54, 1801–1813 (2021).

16. Maurais, A. J. & Weerapana, E. Reactive-cysteine profiling for drug discovery. Curr. Opin. Chem. Biol. 50, 29–36 (2019).

17. Wang, Y. et al. Expedited mapping of the ligandable proteome using fully functionalized enantiomeric probe pairs. Nat. Chem. 11, 1113–1123 (2019).

18. Vinogradova, E. V. et al. An Activity-Guided Map of Electrophile-Cysteine Interactions in Primary Human T Cells. Cell 182, 1009–1026.e29 (2020).

19. Won, S. J. et al. Redirecting the pioneering function of FOXA1 with covalent small molecules. Mol. Cell 84, 4125–4141.e10 (2024).

20. Lazear, M. R. et al. Proteomic discovery of chemical probes that perturb protein complexes in human cells. Mol. Cell 83, 1725–1742.e12 (2023).

21. Liu, Z. et al. Proteomic Ligandability Maps of Spirocycle Acrylamide Stereoprobes Identify Covalent ERCC3 Degraders. J. Am. Chem. Soc. 146, 10393–10406 (2024).

22. Chen, Y. et al. Direct mapping of ligandable tyrosines and lysines in cells with chiral sulfonyl fluoride probes. Nat. Chem. 15, 1616–1625 (2023).

23. Njomen, E. et al. Multi-tiered chemical proteomic maps of tryptoline acrylamide– protein interactions in cancer cells. Nat. Chem. 16, 1592–1604 (2024).

24. Tao, Y. et al. Targeted Protein Degradation by Electrophilic PROTACs that Stereoselectively and Site-Specifically Engage DCAF1. J. Am. Chem. Soc. 144, 18688–18699 (2022).

25. Grygorenko, O. O., Volochnyuk, D. M. & Vashchenko, B. V. Emerging Building Blocks for Medicinal Chemistry: Recent Synthetic Advances. Eur. J. Org. Chem. 2021, 6478–6510 (2021).

26. Blakemore, D. C. et al. Organic synthesis provides opportunities to transform drug discovery. Nat. Chem. 10, 383–394 (2018).

27. Goetzke, F. W., Mortimore, M. & Fletcher, S. P. Enantio– and Diastereoselective Suzuki–Miyaura Coupling with Racemic Bicycles. Angew. Chem. Int. Ed. 58, 12128–12132 (2019).

28. Mishra, S., Modicom, F. C. T., Dean, C. L. & Fletcher, S. P. Catalytic asymmetric synthesis of carbocyclic C-nucleosides. Commun. Chem. 5, 154 (2022).

29. Henderson, A. S., Medina, S., Bower, J. F. & Galan, M. C. Nucleophilic Aromatic Substitution (SNAr) as an Approach to Challenging Carbohydrate-Aryl Ethers. Org. Lett. 17, 4846–4849 (2015).

30. Tao, Y. et al. Chemical Proteomic Discovery of Isotype-Selective Covalent Inhibitors of the RNA Methyltransferase NSUN2. Angew. Chem. Int. Ed. 62, e202311924 (2023).

31. Rostovtsev, V. V., Green, L. G., Fokin, V. V. & Sharpless, K. B. A Stepwise Huisgen Cycloaddition Process: Copper(I)-Catalyzed Regioselective “Ligation” of Azides and Terminal Alkynes. Angew. Chem. Int. Ed. 41, 2596–2599 (2002).

32. Tornøe, C. W., Christensen, C. & Meldal, M. Peptidotriazoles on Solid Phase: [1,2,3]-Triazoles by Regiospecific Copper(I)-Catalyzed 1,3-Dipolar Cycloadditions of Terminal Alkynes to Azides. J. Org. Chem. 67, 3057–3064 (2002).

33. Jumper, J. et al. Highly accurate protein structure prediction with AlphaFold. Nature 596, 583–589 (2021).

34. Varadi, M. et al. AlphaFold Protein Structure Database in 2024: providing structure coverage for over 214 million protein sequences. Nucleic Acids Res. 52, D368–D375 (2024).

35. Brūmele, B. et al. Human TRMT112-Methyltransferase Network Consists of Seven Partners Interacting with a Common Co-Factor. Int. J. Mol. Sci. 22, 13593 (2021).

36. Van Tran, N. et al. The human 18S rRNA m6A methyltransferase METTL5 is stabilized by TRMT112. Nucleic Acids Res. 47, 7719–7733 (2019).

37. Figaro, S. et al. Trm112 Is Required for Bud23-Mediated Methylation of the 18S rRNA at Position G1575. Mol. Cell. Biol. 32, 2254–2267 (2012).

38. Figaro, S., Scrima, N., Buckingham, R. H. & Heurgué-Hamard, V. HemK2 protein, encoded on human chromosome 21, methylates translation termination factor eRF1. FEBS Lett. 582, 2352–2356 (2008).

39. Yang, W.-Q. et al. THUMPD3–TRMT112 is a m2G methyltransferase working on a broad range of tRNA substrates. Nucleic Acids Res. 49, 11900–11919 (2021).

40. Songe-Møller, L. et al. Mammalian ALKBH8 Possesses tRNA Methyltransferase Activity Required for the Biogenesis of Multiple Wobble Uridine Modifications Implicated in Translational Decoding. Mol. Cell. Biol. 30, 1814–1827 (2010).

41. Fu, D. et al. Human AlkB Homolog ABH8 Is a tRNA Methyltransferase Required for Wobble Uridine Modification and DNA Damage Survival. Mol. Cell. Biol. 30, 2449–2459 (2010).

42. Wang, C. et al. N2-methylguanosine modifications on human tRNAs and snRNA U6 are important for cell proliferation, protein translation and pre-mRNA splicing. Nucleic Acids Res. 51, 7496–7519 (2023).

43. Senatore, E. et al. The TBC1D31/praja2 complex controls primary ciliogenesis through PKA-directed OFD1 ubiquitylation. EMBO J. 40, (2021).

44. van Tran, N. et al. The human 18S rRNA m6A methyltransferase METTL5 is stabilized by TRMT112. Nucleic Acids Res. 47, 7719–7733 (2019).

45. Turkalj, E. M. & Vissers, C. The emerging importance of METTL5-mediated ribosomal RNA methylation. Exp. Mol. Med. 54, 1617–1625 (2022).

46. Sepich-Poore, C. et al. The METTL5-TRMT112 N6-methyladenosine methyltransferase complex regulates mRNA translation via 18S rRNA methylation. J. Biol. Chem. 298, 101590 (2022).

47. Rong, B. et al. Ribosome 18S m6A Methyltransferase METTL5 Promotes Translation Initiation and Breast Cancer Cell Growth. Cell Rep. 33, 108544 (2020).

48. Yu, D., Kaur, G., Blumenthal, R. M., Zhang, X. & Cheng, X. Enzymatic characterization of three human RNA adenosine methyltransferases reveals diverse substrate affinities and reaction optima. J. Biol. Chem. 296, 100270 (2021).

49. Edwards, A. N. & Hsu, K.-L. Emerging opportunities for intact and native protein analysis using chemical proteomics. Anal. Chim. Acta 1338, 343551 (2025).

50. Ai, H., Pan, M. & Liu, L. Chemical Synthesis of Human Proteoforms and Application in Biomedicine. ACS Cent. Sci. 10, 1442–1459 (2024).

51. Green, J. J., Grimm, C., Fristo, A., Byrum, J. & Kelleher, N. L. Parsing 20 Years of Public Data by AI Maps Trends in Proteomics and Forecasts Technology. J. Proteome Res. 23, 523–531 (2024).

52. Thompson, R. E. & Muir, T. W. Chemoenzymatic Semisynthesis of Proteins. Chem. Rev. 120, 3051–3126 (2020).

53. Toby, T. K., Fornelli, L. & Kelleher, N. L. Progress in Top-Down Proteomics and the Analysis of Proteoforms. Annu. Rev. Anal. Chem. 9, 499–519 (2016).

54. Gassaway, B. M. et al. A multi-purpose, regenerable, proteome-scale, human phosphoserine resource for phosphoproteomics. Nat. Methods 19, 1371–1375 (2022).

55. Wang, L. et al. Mettl5 mediated 18S rRNA N6-methyladenosine (m6A) modification controls stem cell fate determination and neural function. Genes Dis. 9, 268–274 (2022).

56. Li, Z. et al. METTL5-mediated 18S rRNA m6A modification promotes corticospinal tract sprouting after unilateral traumatic brain injury. Exp. Neurol. 383, 115000 (2025).

57. Peng, H. et al. N6-methyladenosine (m6A) in 18S rRNA promotes fatty acid metabolism and oncogenic transformation. Nat. Metab. 4, 1041–1054 (2022).

58. Chen, B. et al. N6-methyladenosine modification in 18S rRNA promotes tumorigenesis and chemoresistance via HSF4b/HSP90B1/mutant p53 axis. Cell Chem. Biol. 30, 144–158.e10 (2023).

59. Richard, E. M. et al. Bi-allelic Variants in METTL5 Cause Autosomal-Recessive Intellectual Disability and Microcephaly. Am. J. Hum. Genet. 105, 869–878 (2019).

60. Liu, N. et al. Probing N6-methyladenosine RNA modification status at single nucleotide resolution in mRNA and long noncoding RNA. RNA 19, 1848–1856 (2013).

61. Luo, M. et al. METTL5 enhances the mRNA stability of TPRKB through m6A modification to facilitate the aggressive phenotypes of hepatocellular carcinoma cell. Exp. Cell Res. 442, 114219 (2024).

62. Wang, P., Ye, C., Zhao, M., Jiang, B. & He, C. Small-molecule-catalysed deamination enables transcriptome-wide profiling of N6-methyladenosine in RNA. Nat. Chem. (2025) doi:10.1038/s41557-025-01801-3.

63. Xiao, Y.-L. et al. Transcriptome-wide profiling and quantification of N6-methyladenosine by enzyme-assisted adenosine deamination. Nat. Biotechnol. 41, 993–1003 (2023).

64. Liu, C. et al. Absolute quantification of single-base m6A methylation in the mammalian transcriptome using GLORI. Nat. Biotechnol. 41, 355–366 (2023).

65. Roy, N. et al. Suppression of NRF2-dependent cancer growth by a covalent allosteric molecular glue. Preprint at 10.1101/2024.10.04.616592 (2024).

66. Jahnke, W. et al. Binding or Bending: Distinction of Allosteric Abl Kinase Agonists from Antagonists by an NMR-Based Conformational Assay. J. Am. Chem. Soc. 132, 7043–7048 (2010).

67. Schreiber, S. L. Molecular glues and bifunctional compounds: Therapeutic modalities based on induced proximity. Cell Chem. Biol. 31, 1050–1063 (2024).

68. Zhang, Y. et al. An allosteric cyclin E-CDK2 site mapped by paralog hopping with covalent probes. Nat. Chem. Biol. 21, 420–431 (2025).

69. Perez-Riverol, Y. et al. The PRIDE database at 20 years: 2025 update. Nucleic Acids Res. 53, D543–D553 (2025).

