## Supplementary Biology Information for "Complexoform-restricted covalent TRMT112 ligands that allosterically agonize METTL5"

### MATERIALS AND METHODS

#### Cell Lines and Cell Culture

Ramos (ATCC, CRL-1596) cells were grown in Roswell Park Memorial Institute medium (RPMI) supplemented with 10% fetal bovine serum (FBS), 2 mM L-glutamine, penicillin (100 U $mL^{-1}$), and streptomycin (100 µg $mL^{-1}$). HEK293T (ATCC, CRL-3216), Lenti-X (Takara, #632180) and HCT116 (ATCC, CCL-247) were grown in Dulbecco's modified Eagle medium (DMEM) supplemented with 10% fetal bovine serum (FBS), 2 mM L-glutamine, penicillin (100 U $mL^{-1}$), and streptomycin (100 µg $mL^{-1}$). Cells were cultured in a humidified, 37 °C/5% $CO_2$ tissue culture incubator. All cell lines were routinely inspected for mycoplasma contamination.

#### Cloning and mutagenesis

The expression plasmid pRK5_RNF14-FLAG was previously prepared by our lab[1]. Mutagenesis was performed using the Q5 site-directed mutagenesis kit (NEB, E0554S) with the following primers.

RNF14_C68A_FWD: ATACACCATTgccTTTCTGCCTC

RNF14_C68A_REV: TCAAAGCCACTATTCTGG

RNF14_C212A_FWD: GCAGATAAAAgccTTTAATAGTAAATTGTTCC

RNF14_C212A_FWD: TGAGCTTGATCAAAGTCC

ORFs of HA-N6AMT1 and HA-BUD23 were synthesized from IDT. The ORF of METTL5 was synthesized from IDT and was PCR amplified (two consecutive PCRs) with primers containing a N-terminal HA tag, using the primers described below.

HA-METTL5_FWD_1:

TACCCATACGATGTTCCAGATTACGCTGGTACCATGAAGAAAGTAAGGCTTAA

HA-METTL5_REV_1: CCTAATTCGGTTTTCCTTT

HA-METTL5_FWD_2:

ggggacaagtttgtacaaaaaagcaggctgccaccATGTACCCATACGATGTTCCAG

HA-METTL5_REV_2: accactttgtacaagaaagctgggttcaAAAGGAAAACCGAATTAGGTCC

HA-N6AMT1, HA-METTL5 and HA-BUD23 were cloned into a pRK5 expression vector using Gateway technology (Thermo, #11791019 and #11789013) (for ORF see Supplementary Table 2).

**Generation of HCT116 cells stably expressing WT-TRMT112-FLAG, C100A-TRMT112-FLAG or C100S-TRMT112-FLAG**

Cloning and mutagenesis

Codon optimized WT-TRMT112-FLAG and C100A-TRMT112-FLAG were synthesized from IDT and cloned into a pLV416 expression vector using Gateway technology (Thermo, #11791019 and #11789013; for ORF see Supplementary Table 2). The plasmid pLV416_ C100S-TRMT112_FLAG was cloned by mutagenesis using a Q5 site-directed mutagenesis kit using the primers described below.

TRMT112_C100S_FWD: AACACTGCAGagcCCTGAGAGCG

TRMT112_C100S_REV: CCCTCGATCACCTCAAC

Virus production

HEK293T cells ($6.0 \times 10^5$ in 2 mL DMEM/well) were seeded into 6-well dishes and allowed to grow for one day. pLV416_WT-TRMT112-FLAG or mutants (1 µg), pCMV-dR8.91 (1 µg) and pCMV-VSV-G (0.1 µg) were mixed in 100 µL Opti-MEM™. Then, 10 µL of 1 mg $mL^{-1}$ PEI (Polysciences) was added and incubated for 15 min. The mixture was gently added to the cells. After 8 h, the media was removed and exchanged with DMEM supplemented with 30% fetal FBS (L-glutamine, penicillin, streptomycin). The virus was collected after 2 and 3 days, combined and filtered with 0.45 µm syringe filters (Millipore).

Transduction

HCT116 ($4.0 \times 10^5$ in 1 mL DMEM/well) cells were mixed in a 12-well dish with 200 µL virus supernatant and 8 µg $mL^{-1}$ polybrene (Millipore). Spin infection was performed (900 x *g*, 1 h at 30 °C). On day 1, the virus containing media was removed, and fresh DMEM was added. On day 2, 600 µg $mL^{-1}$ G418 was added to start the selection (2 weeks).

**Generation of HCT116 cells stably expressing WT-TRMT112-FLAG and HA-METTL5**

The open ORF of HA-METTL5 was cloned into the lentiviral expression vector pLEX307 using Gateway technology (Thermo, #11791019 and #11789013). Virus production and transduction of $HCT116_{FLAG-TRMT112}$ cells was performed as described above. On day 2 after the transduction, 2 µg $mL^{-1}$ puromycin was added to start the selection (3 days).

**Generation of HCT116 cells with METTL5-KO**

Stable knockout cell lines were generated by transduction of HCT116 cells with lentiCRISPR v2 vector (sgRNAs cloned into the vector, Addgene #52961) via BsmBI (Neb, #R0739S) restriction and annealed oligo ligation cloning (see primers below). Virus production transduction of HCT116 cells was performed as described above except with Lenti-X cells rather than HEK293T. On day 2 after the transduction, 2 µg $mL^{-1}$ puromycin was added to start the selection (3 days) and the cells were subsequently expanded for 2 weeks. Successful knock-out was validated by immunoblotting and gDNA sequencing.

sgMETTL5_FWD: caccgCCAGGCCGCACATTGCAGGT

sgMETTL5_REV: aaacACCTGCAATGTGCGGCCTGGc

**Western blotting**

Proteins were blotted onto PVDF membrane (60 V, 2 h), blocked with 5% milk in TBST for 30 min at room temperature and incubated with primary antibodies (see Supplementary Information Table 3 for dilution) at 4 °C overnight. Membranes were washed with TBST (3 x) before treatment with secondary antibodies for 2 h at room temperature or directly imaged if a primary HRP-conjugate was used. Membranes were developed with SuperSignal™ West Pico PLUS Chemiluminescent Substrate and visualized by chemiluminescence scan on a ChemiDoc MP Imaging System (Bio-Rad).

**Gel-ABPP for proteome-wide reactivity**

Ramos cells (5 mL of 3 x $10^6$ cells $mL^{-1}$) were treated with alkyne probes for 1 h. Cells were collected by centrifugation at 500 x *g* for 5 min at 4 °C and washed twice with cold DPBS. Cell pellets were resuspended in 250 µL cold DPBS and lysed by pulse sonication (3 x 8 pulses, 10% power output) and normalized to 100 µL of 1 mg $mL^{-1}$ whole cells lysates (Pierce BCA protein assay). Samples were treated with 11 µL of click Master-mix

[6 μL of 1.7 mM TBTA in 4:1 *t*-BuOH:DMSO, 2 μL of 50 mM $CuSO_4$ in $H_2O$, 2 μL of freshly prepared 50 mM Tris(2-carboxyethyl)phosphine in $H_2O$, 0.8 μL of 1.25 mM tetramethylrhodamine (TAMRA)-PEG3-azide)] for 1 h, followed by the addition 4X SDS gel loading buffer (36 μL). Proteins were resolved by SDS-PAGE (275 V, 4 h, 10% Tris-glycine, made in-house) and visualized by in-gel fluorescence on a ChemiDoc MP Imaging System (Bio-Rad). The images were processed using Image Lab software (version 6.1.0). Gels were stained with Coomassie InstantBlue® Protein Stain and visualized by Coomassie gel scan on a ChemiDoc MP Imaging System (Bio-Rad).

**Gel-ABPP with recombinant RNF14**

HEK293T cells (3.5 x $10^5$ in 2 mL DMEM/well) were seeded into 6-well dishes the day before transfection with pRK5_RNF14-FLAG (1 μg) (PEI:DNA = 3:1) for 2 days. Untransfected (UT) cells were treated only with PEI. After 2 days, the media was exchanged (1 mL DMEM/well). Cells were treated with DMSO or competitor for 3 h, followed by alkyne for 1 h. Cells were scraped in cold DPBS and collected by centrifugation at 500 x *g* for 5 min at 4 °C and washed with cold DPBS. The cell pellets were resuspended in 200 μL cold DPBS and lysed by pulse sonication (3 x 8 pulses, 10% power output). After centrifugation at 21,000 x *g* for 5 min at 4 °C, the supernatants were normalized to 100 μL of 1 mg $mL^{-1}$ (DC protein assay) and were treated with 11 μL of click Master-mix, as described above. Proteins were resolved by SDS-PAGE gel (160 V, 1 h, 4–20%, Tris-glycine: Invitrogen, XP04205BOX) and visualized as described above, followed by Western blotting.

**Gel-ABPP with stable HCT116 cell lines**

Stable HCT116 cells (7.5 x $10^5$ in 2 mL DMEM/well) were seeded into 6-well dishes the day before treatment. The media was exchanged (1 mL DMEM/well) and cells were treated with DMSO or competitor for 3 h, followed by treatment with alkyne for 1 h. Cells were scraped in cold DPBS and collected by centrifugation at 500 x *g* for 5 min at 4 °C and washed with cold DPBS. The cell pellets were resuspended in 200 μL cold DPBS and lysed by pulse sonication (3 x 8 pulses, 10% power output). After centrifugation at 21,000 x *g* for 5 min, the supernatants were normalized to 100 μL of 1 mg $mL^{-1}$ (DC protein assay) and were treated with 11 µL of click Master-mix, as described above. Proteins were resolved by SDS-PAGE gel (160 V, 1.25 h, 4–20% Tris-glycine) and visualized as described above, followed by Western blotting.

Modification for transient co-expression with MTs

HCT116$_{TRMT112\text{-}FLAG}$ cells (6.25 x $10^5$ in 2 mL DMEM/well) were seeded into 6-well dishes the day before transfection with pRK5_HA-MT (2.5 µg) (10 µL of Lipofectamine LTX (Invitrogen, #15338100) in 500 µL Opti-MEM™ (complexed for 30 min) for 1 day. Untransfected (UT) cells were treated only with Lipofectamine LTX. Samples were further processed as described as above.

Quantification

Band intensities were quantified using the Image Lab (6.1.0) software (Bio-Rad). $IC_{50}$ curves were generated using GraphPad Prism version 10.4.1, applying a four-parameter variable slope nonlinear regression with the top and bottom constraints set to 100% and 0%, respectively.

**IP-Gel-ABPP for TRMT112**

HCT116 (parental control) or HCT116$_{TRMT112\text{-}WT\text{-}FLAG}$ cells (5.0 x $10^6$ in 10 mL DMEM/dish) were seeded into 10 cm plates the day before. On the day of the treatment, the media was exchanged (10 mL DMEM/dish). Cells were treated with DMSO or competitor for 3 h, followed by treatment with alkyne for 1 h. and harvested. Cells were scraped in cold DPBS, collected by centrifugation at 500 x *g* for 5 min at 4 °C and washed with cold DPBS. The cell pellets were resuspended in 500 µL cold DPBS containing 1% NP-40 and cOmplete Protease Inhibitor Cocktail (Roche) and lysed by pulse sonication (8 pulses, 10% power output). The lysate was centrifugated at 21,000 x *g* for 5 min.

For the input: 50 µL of 1 mg $mL^{-1}$ of normalized supernatants (DC protein assay) were treated with 5.5 µL of click Master-mix and processed as described above.

For the IP: 500 µL of 2 mg $mL^{-1}$ of supernatants were incubated with washed Pierce™ Anti-DYKDDDDK Magnetic Agarose (Thermo, #A36797; 40 µL of 25% slurry/sample) for 2 h at 4 °C with rotation. After incubation, the beads were isolated with a magnetic stand

and washed with 0.2% NP-40 in DPBS (3 x 1 mL), followed by DPBS (1 x 1 mL). The beads were resuspended in 50 µL DPBS and treated with 5.5 µL of click Master-mix and further processed as described above, followed by Western blotting.

**GSH reactivity assay**

As described previously[2], glutathione (GSH) was diluted to a final concentration of 50 µM in 0.1 M Tris-HCl pH 8.8, 30% acetonitrile. In triplicate, 100 µL of the GSH solution was added to a clear 384-well plate (Greiner, #781101). Stereoprobes (5 µL of 10 mM) were then added to the GSH solution to achieve a final probe concentration of 500 µM, and the reaction was incubated for 2 h at r.t.. Ellman's reagent (5 µL of 100 mM) was then added to the plate and the absorbance read at 440 nm. The concentration of GSH remaining was derived from a standard curve and the observed rate ($k_{obs}$/[I]) was calculated assuming pseudo-first-order reaction kinetics from the following equations:

(eq. 1) $d[GSH]/dt = -k \cdot [GSH]$

(eq. 2) $[GSH] = [GSH]_{t0} \cdot e^{-kt}$.

**Protein-directed ABPP**

*In situ* treatment and sample processing

As described previously[2], Ramos cells (10 mL of 3 x $10^6$ cells $mL^{-1}$) were treated with DMSO or competitor for 3 h, followed by treatment with a stereo-matched alkyne for 1 h. The cells were harvested on ice, washed twice with cold DPBS and stored at −80 °C. The cell pellets were resuspended in 500 µL cold DPBS and lysed by pulse sonication (3 x 8 pulses, 10% power output). 500 µL of 1 mg $mL^{-1}$ of normalized whole cells lysates (Pierce BCA protein assay) were treated with 55 µL of click MS-Master-mix [30 µL of 1.7 mM TBTA in 4:1 *t*-BuOH:DMSO, 10 µL of 50 mM $CuSO_4$ in $H_2O$, 10 µL of freshly prepared 50 mM Tris(2-carboxyethyl)phosphine in $H_2O$, 10 µL of 10 mM Biotin-PEG4-azide]. Proteins were precipitated with cold methanol (600 µL), chloroform (200 µL) and water (100 µL), vortexed, and then centrifuged at 16,000 x *g* for 10 min. The top and bottom layers were aspirated, and the protein disk was sonicated in 500 µL of methanol and pelleted at 16,000 x *g* for 10 min. After the methanol was completely aspirated, protein pellets were immediately processed or stored at −80 °C. Pellets were resuspended in 500 μL freshly prepared 8 M urea in DPBS, followed by the addition of 10 μL of 10 wt% SDS. Samples were then pulse-sonicated until clear. The samples were reduced with 25 μL of 200 mM dithiothreitol (DTT) at 65 °C for 15 min, followed by alkylation with 25 μL of 400 mM iodoacetamide at 37 °C for 30 min. Then, 65 μL of 20 wt% SDS was added, and the samples were transferred to a 15-mL tube in a total volume of 6 mL with DPBS (0.2% final SDS). Washed streptavidin beads (Thermo, #20353; 100 μL of 50% slurry/sample) were then added and proteins were enriched for 1.5 h at r.t. with rotation. After incubation, the beads were pelleted (2 min at 2,000 x *g*) and washed with 0.2% wt% SDS in DPBS (2 x 10 mL), DPBS (1 x 5 mL), HPLC-grade water (2 x 1 mL) and 200 mM 4-(2-hydroxyethyl)-1-piperazinepropanesulfonic acid (EPPS; 1 mL, pH 8.0). Enriched proteins were digested on-bead overnight with 200 μL of trypsin mix (2 M urea, 1 mM $CaCl_2$, 10 μg $mL^{-1}$ trypsin (Promega, #V5111), 200 mM EPPS, pH 8.0). The beads were pelleted at 2,000 x *g*, the supernatant was collected and then diluted with 100 μL acetonitrile (30% final). Samples were then labeled with 6 μL 20 mg $mL^{-1}$ (in dry acetonitrile) $TMTpro^{16}plex$ tag (Thermo, #A44520) or $TMT^{10}plex$ (Thermo, #90406) for 1.5 h at r.t (vortex every 30 min). TMT labelling was quenched by the addition of hydroxylamine (6 μL 5% solution in $H_2O$) and incubated for 15 min at r.t.. Samples were then acidified with 20 μL formic acid, combined and dried using a SpeedVac at 46 °C. Samples were desalted with a Sep-Pak column Vac 1 cc (50 mg) (Waters, #WAT054955) and then high pH fractionated into ten fractions (for 16-plex) or five fractions (10-plex) using Pierce peptide desalting spin columns (Thermo, #89852) and acetonitrile/$NH_4HCO_3$ (10 mM) gradient (as described in Supplementary Table 4 and 5) and analyzed by mass spectrometry (see TMT LC-MS Analysis).

<u>Modifications for adherent cells</u>

<u>Protein-directed ABPP for RNF14:</u> HEK293T cells (3.5 x $10^6$ in 10 mL DMEM) were seeded into four 10 cm plates the day before transfection with pRK5_RNF14-FLAG (5 μg/dish) (PEI:DNA = 3:1). After 1 day, the cells were pooled and distributed over eighteen 10 cm plates with 10 mL DMEM each. The next day, the media was exchanged (7 mL DMEM/dish) and cells were treated with DMSO or competitor for 3 h, followed by

treatment with alkyne for 1 h. Cells were scraped in media on ice, washed twice with cold DPBS and stored at −80 °C. The samples were further processed as described as above.

Comparative protein-directed ABPP: HCT116 or $HCT116_{METTL5\text{-}KO}$ cells (4.5 x $10^6$ in 10 mL DMEM/dish) were seeded into 10 cm plates the day before. On the day of the treatment, the media was exchanged (7 mL DMEM/dish). Cells were treated with alkyne for 1 h. Cells were scraped in cold DPBS, washed twice with cold DPBS and stored at −80 °C. The samples were further processed as described as above.

TMT LC-MS Analysis

Fractions were resuspended in buffer A (5% acetonitrile, 0.1% formic acid in water) and analyzed by liquid chromatography tandem mass-spectrometry using an Orbitrap Fusion Tribrid Mass Spectrometer (Thermo Scientific) coupled to an UltiMate 3000 Series Rapid Separation LC system and autosampler (Thermo Scientific Dionex). The peptides were eluted onto a capillary column (75-µm-inner-diameter fused silica, packed with C18 (Waters, Acquity BEH C18, 1.7 µm, 25 cm) or an EASY-Spray HPLC column (Thermo, #ES902, #ES903) using an Acclaim PepMap 100 (Thermo, #164535) loading column, and separated at a flow rate of 0.25 µL $min^{-1}$. Peptides were separated across a 10 min gradient of 5%, 150 min gradient of 5-20%, 20 min 20-45%, and then 5 min 45-95% acetonitrile (0.1% formic acid) in $H_2O$ (0.1% formic acid) followed by column equilibration. Data were acquired using an MS3-based TMT method on Orbitrap Fusion or Eclipse Tribrid mass spectrometers.

Fusion instruments: The scan sequence began with an MS1 master scan (Orbitrap analysis, resolution 120,000, 400-1,700 m/z, RF lens 60%, maximum injection time 50 ms) with dynamic exclusion enabled (repeat count 1, duration 15 s). The top precursors were then selected for MS2/MS3 analysis. MS2 analysis consisted of quadrupole isolation (isolation window 0.7) of precursor ion followed by collision-induced dissociation in the ion trap (collision energy 35%, maximum injection time 120 ms). Following the acquisition of each MS2 spectrum, synchronous precursor selection enabled the selection of up to 10 MS2 fragment ions for MS3 analysis. MS3 precursors were fragmented by higher-energy collisional dissociation (HCD) and analyzed using the Orbitrap (collision energy

55, maximum injection time 120 ms, resolution 50,000). For MS3 analysis, we used charge state-dependent isolation windows. For charge state z = 2, the MS isolation window was set at 1.2; for z = 3–6 the MS isolation window was set at 0.7.

Eclipse Tribrid instrument: The scan sequence began with an MS1 master scan (Orbitrap analysis, resolution 120,000, 400-1,700 m/z, RF lens 30%, maximum injection time 50 ms) with dynamic exclusion enabled (repeat count 1, duration 30 s). The top precursors were then selected for MS2/MS3 analysis. MS2 analysis consisted of quadrupole isolation (isolation window 0.7) of precursor ion followed by higher-energy collisional dissociation (HCD) in the ion trap (collision energy 36%, maximum injection time 120 ms). Following the acquisition of each MS2 spectrum, synchronous precursor selection enabled the selection of up to 10 MS2 fragment ions for MS3 analysis. MS3 precursors were fragmented by HCD and analyzed using the Orbitrap (collision energy 55%, maximum injection time 120 ms, resolution 30,000). For MS3 analysis, we used charge state-dependent isolation windows. For charge state z = 2, the MS isolation window was set at 1.2; for z = 3 the MS isolation window was set at 0.7; for z = 4–6, the MS isolation window was set at 0.4.

Data processing

Raw files were uploaded to the Integrated Proteomics Pipeline (IP2, version 6.7.1) available at http://ip2.scripps.edu/ip2/mainMenu.html, and MS2 and MS3 files were extracted from the raw files using RAW Converter (version 1.1.0.22, available at http://fields.scripps.edu/rawconv/) and searched using the ProLuCID algorithm using a reverse concatenated, non-redundant variant of the Human UniProt database (release 2016-07). Cysteine residues were searched with a static modification for carboxyamidomethylation (+57.02146 Da). N termini and lysine residues were also searched with a static modification corresponding to the TMT tag (+229.1629 Da for 10-plex and +304.2071 Da for 16-plex). Peptides were required to be at least six amino acids long. ProLuCID data were filtered through DTASelect (version 2.0) to achieve a spectrum false-positive rate below 1%. We included a keratin filter. The MS3-based peptide quantification was performed with reporter ion mass tolerance set to 20 ppm with the Integrated Proteomics Pipeline (IP2).

Data analysis

Enrichment ratios (probe versus probe) were calculated for each peptide-spectrum match (PSM) by dividing the TMT reporter ion intensity by the total intensity across all channels. PSMs were grouped by protein ID, excluding peptides with summed reporter ion intensities <10,000, signal-to-noise ratio (S/N) <1.0, or isolation purity <0.5. Proteins supported by fewer than two peptides were also excluded. For two experiments, one TMT channel showed substantially reduced reporter ion intensity and was excluded from analysis. To maintain 16-channel balance during normalization, the corresponding replicate channel was temporarily duplicated, and protein signals were normalized by total intensity across all channels (sum = 100), including the duplicated channel. The temporarily duplicated channel was finally removed prior to any downstream statistical analysis.

Replicate channels were grouped across each experiment, and mean values were calculated for each protein. To assess measurement variability, the coefficient of variation (CV), defined as the ratio of the standard deviation to the mean, was calculated across replicate channels. Proteins with a CV ≥ 0.2 in the most enriched (DMSO + alkyne) channel were excluded from further analysis.

Criteria for liganding in protein-directed ABPP

Proteins were initially defined as liganded if they exhibited >2-fold enantioselective enrichment and >50% competition. All proteins passing the initial filters for liganding were manually reviewed to remove proteins showing additional evidence of high variability.

**Cysteine-directed ABPP**

*In situ* treatment and sample processing

As described previously[2], Ramos cells (10 mL of 3 x $10^6$ cells $mL^{-1}$) were seeded 30 min prior to the experiment. Cells were treated with DMSO or competitor for 3 h. The cells were harvested on ice, washed twice with cold DPBS and stored at −80 °C. The cell pellets were resuspended in 500 µL cold DPBS and lysed by pulse sonication (3 x 8 pulses, 10% power output). The total protein content of whole-cell lysates was measured using a Pierce BCA protein assay kit and the samples were normalized to 2 mg mL$^{-1}$ and 500 µL. Samples were treated with 5 µL of 10 mM IA-DTB (in DMSO) for 1 h at r.t. with vortexing every 20 min. Proteins were precipitated by the addition of cold methanol (600 µL), chloroform (200 µL) and HPLC-grade water (100 µL), followed by vortexing and centrifugation at 16,000 x *g* for 10 min. Without disrupting the protein disk, both the top and bottom layers were aspirated, and the protein disk was sonicated again in 500 µL of methanol and centrifuged at 16,000 x *g* for 10 min. After the methanol was completely aspirated, protein pellets were immediately processed or frozen at −80 °C. Pellets were resuspended in 90 µL of denaturing/reducing buffer (9 M urea, 10 mM DTT, 50 mM triethylammonium bicarbonate (TEAB) pH 8.5). The samples were reduced by heating at 65 °C for 20 min, followed by alkylation with 10 µL of 500 mM iodoacetamide at 37 °C for 30 min. The samples were then centrifuged at 16,000 x *g* for 2 min to pellet any insoluble precipitate and probe-sonicated once more to ensure complete resuspension, and then diluted with 300 µL of 50 mM TEAB pH 8.5 to reach a final urea concentration of 2 M. Trypsin (4 µL of 0.25 µg µL$^{-1}$ in trypsin resuspension buffer with 25 mM $CaCl_2$) was added to each sample and digested at 37 °C overnight. Digested samples were then diluted with 300 µL of enrichment buffer (50 mM TEAB pH 8.5, 150 mM NaCl, 0.2% NP-40) containing streptavidin-agarose beads (50 µL of 50% slurry/sample) and were rotated at r.t. for 2 h. The samples were centrifuged (2,000 x *g*, 2 min) and the entire content transferred to BioSpin columns and washed (3 x 1 mL wash buffer, 3 x 1 mL DPBS, 3 x 1 mL water). Enriched peptides were eluted from the beads with 300 µL of 50% acetonitrile with 0.1% formic acid and dried using a SpeedVac at 46 °C Enriched peptides were resuspended in 100 µL EPPS buffer (200 mM, pH 8.0) with 30% acetonitrile, vortexed and water bath-sonicated. The samples were TMT-labelled by the addition of 3 µL of 20 mg mL$^{-1}$ (in dry acetonitrile) of corresponding TMT$^{10}$plex tag for 1.5 h at r.t with vortexing every 30 min. TMT labelling was quenched by the addition of hydroxylamine (3 µL 5% solution in $H_2O$) and incubated for 15 min at r.t.. Samples were then acidified with 5 µL formic acid, combined and dried using a SpeedVac. Samples were desalted with a Sep-Pak column and then high-pH-fractionated by HPLC (described in the following) into a 96-well plate and recombined into 12 fractions (total).

As previously described[2], the cysteine-directed ABPP samples were resuspended in 500 µL of buffer A and fractionated with an Agilent HPLC system into a 96-deep-well plate containing 20 µL of 20% formic acid to acidify the eluting peptides. The peptides were eluted onto a capillary column (ZORBAX 300Extend-C18, 3.5 µm) and separated at a flow rate of 0.5 mL $min^{-1}$ using the following gradient: 100% buffer A from 0 min to 2 min, 0-13% buffer B from 2 min to 3 min, 13–42% buffer B from 3 min to 60 min, 42-100% buffer B from 60 min to 61 min, 100% buffer B from 61 min to 65 min, 100–0% buffer B from 65 min to 66 min, 100% buffer A from 66 min to 75 min, 0-13% buffer B from 75 min to 78 min, 13–80% buffer B from 78 min to 80 min, 80% buffer B from 80 min to 85 min, 100% buffer A from 86 min to 91 min, 0-13% buffer B from 91 min to 94 min, 13–80% buffer B from 94 min to 96 min, 80% buffer B from 96 min to 101 min, and 80–0% buffer B from 101 min to 102 min (buffer A, 10 mM aqueous $NH_4HCO_3$; buffer B, acetonitrile). The plates were evaporated to dryness using a SpeedVac and peptides resuspended in 80% acetonitrile, with 0.1% formic acid, and combined to a total of 12 fractions (for example, fraction1 = well 1A + 1B … 1H, fraction 2 = well 2A + 2B …. 2H) (3 x 300 µL per column). Samples were dried with a SpeedVac and analyzed by mass spectrometry (see TMT LC-MS Analysis above).

Data processing

Raw files processed as described above with the following modification: A dynamic modification for IA-DTB labelling (+398.25292 Da) was included with a maximum number of two differential modifications per peptide.

Data analysis

Cysteine engagement ratios (DMSO versus compound) were calculated for each peptide-spectra match (PSM) by dividing each TMT reporter ion intensity by the average intensity for the DMSO channels. Peptide–spectra matches were then grouped based on protein ID and residue number, excluding peptides with summed reporter ion intensities for the two DMSO channels of <10,000, and a coefficient of variation (CV) for DMSO channels of >0.5 in each experiment. Replicate channels were grouped across each experiment, and average values were computed for each cysteine site and peptides with MAD/mean > 0.2 were excluded.

Criteria for liganding in cysteine-directed ABPP

Sites were initially defined as liganded if they exhibited >50% competition and >2-fold enantioselectivity, and if at least one of the following additional criteria was met to distinguish liganding from changes driven by protein abundance:

If the protein passed the 'liganded' criteria in protein-directed ABPP, then detection of liganding at a single site was considered sufficient.

If the protein did not pass the 'liganded' criteria in protein-directed ABPP, we required evidence of at least one additional peptide that did not show liganding by cysteine-directed ABPP in order to classify the site as liganded.

All proteins passing the initial filters for liganding were manually reviewed to remove proteins showing additional evidence of high variability.

Modifications for cysteine-directed ABPP for RNF14 with Glu-C digestion

HEK293T cells (3.5 x $10^6$ in 10 mL DMEM) were seeded into five 10 cm plates the day before transfection with pRK5_RNF14-FLAG (5 µg/dish) (PEI:DNA = 3:1). After 1 day, the cells were pooled and distributed over eleven 15 cm plates with 25 mL media each. The following day, the media was exchanged (13 mL DMEM/dish) and cells were treated with DMSO or competitor for 3 h. Cells were scraped in media on ice, washed twice with cold DPBS and stored at −80 °C. The cell pellets were resuspended in 500 µL cold DPBS and lysed by pulse sonication (3 x 8 pulses, 10% power output). The lysate was cleared by ultracentrifugation 100,000 x *g* for 30 min. Normalized clarified lysates (2 mg $mL^{-1}$ and 500 µL) were further processed as described as above with the following modifications: After methanol precipitation, the samples were resuspended in 90 µL of denaturing/reducing buffer (8 M urea, 10 mM DTT, 50 mM ammonium bicarbonate ($NH_4HCO_3$). After reduction and alkylation, 800 µL of 50 mM $NH_4HCO_3$ were added (final urea 0.8 M). Glu-C (4 µg) (Promega, #V1651) was added to each sample and digested at 37 °C overnight. Digested samples were then diluted with 300 µL of enrichment buffer (50 mM TEAB pH 8.5, 150 mM NaCl, 0.3% NP-40) containing streptavidin-agarose beads (50 µL of 50% slurry/sample) and further processed as described above. In the data processing step, we searched with C-terminal aspartic or glutamic acid cleavage instead of C-terminal lysine or arginine cleavage.

**Immunoprecipitation (IP)-MS experiments for TRMT112**

*In situ* treatment and sample processing

HCT116 (parental control) or stable HCT116 cells (5.0 x $10^6$ in 10 mL DMEM/dish) were seeded into 10 cm plates the day before. On the day of the treatment, the media was exchanged (10 mL DMEM/dish). Cells were treated with DMSO or competitor for 4 h. Cells were scraped in cold DPBS, washed twice with cold DPBS and stored at −80 °C. The cell pellets were resuspended in 500 µL lysis buffer (50 mM EPPS, pH 7.5, 150 mM NaCl) containing 1% NP-40 and cOmplete Protease Inhibitor Cocktail (Roche) and lysed by rotation at 4 °C for 1 h. After centrifugation at 21,000 x *g* for 5 min, 500 µL of 2 mg $mL^{-1}$ of normalized supernatant (BCA protein assay kit) were incubated with washed Pierce™ Anti-DYKDDDDK Magnetic Agarose (Thermo, #A36797; 40 µL of 25% slurry/sample) or Pierce™ Anti-HA Magnetic Beads (Thermo, #A36797; 40 µL of slurry/sample) for 2 h at 4 °C with rotation. After incubation, the beads were isolated with a magnetic stand and washed with wash buffer (25 mM EPPS, pH 7.5, 150 mM NaCl) containing 0.2% NP-40 (3 x 1 mL), followed by 50 mM EPPS (pH 8.0, 1 x 1 mL). Proteins were eluted from beads in 40 µL 8 M urea in DPBS at 65 °C for 10 min. The samples were reduced with 2 µL of 200 mM dithiothreitol (DTT) at 65 °C for 15 min, followed by alkylation with 2 µL of 400 mM iodoacetamide at 37 °C for 30 min. The total volume was brought to 160 µL with 50 mM EPPS (pH 8) (2M final urea). Enriched proteins were digested overnight 37 °C with trypsin (1 µL of 100 mM $CaCl_2$, 1 µg trypsin per sample). Then, 70 µL acetonitrile (30% final) was added, followed by 6 µL of 20 mg $mL^{-1}$ (in dry acetonitrile) of the corresponding $TMTpro^{16}plex$ tag for 1.5 h at r.t with vortexing every 30 min. TMT labelling was quenched by the addition of hydroxylamine (6 µL 5% solution in $H_2O$) and incubated for 15 min at r.t.. Samples were then acidified with 5 µL formic acid, combined and dried using a

SpeedVac at 46 °C. Samples were desalted with a Sep-Pak column and then high pH fractionated into three fractions using peptide desalting spin columns and acetonitrile/$NH_4HCO_3$ (10 mM) gradient into three fractions (as described in Supplementary Table 6) and analyzed by mass spectrometry.

Data processing

Raw files were processed as described for the protein-directed samples above.

**SEC ABPP experiment**

HCT116$_{TRMT112\text{-}WT\text{-}FLAG}$ cells (10 x $10^6$ in 25 mL DMEM/dish) were seeded into 15 cm plates the day before. On the day of the treatment, the media was exchanged (13 mL DMEM/dish). Cells were treated with alkyne for 1 h (comparative SEC ABPP) or pre-treated with DMSO or competitor for 3 h (competitive SEC ABPP), followed by alkyne for 1 h. Cells were scraped in cold DPBS and washed twice with cold DPBS. The cell pellets were resuspended in cold DPBS (600 µL) containing cOmplete Protease Inhibitor Cocktail (Roche) and PhosSTOP Phosphatase Inhibitor Cocktail (Roche) and lysed by pulse sonication (1 x 8 pulses, 10% power output). The lysate was cleared by ultracentrifugation 100,000 x *g* for 30 min. Normalized clarified lysates (2 mg $mL^{-1}$, DC protein assay) were filtered using a centrifugal filter (Milipore, #UFC30GV; 20 µM PVDF) and 500 µL of clarified lysate was injected into a Superdex 200 Increase 10/300 GL column attached to an ÄKTA pure FPLC system (Cytiva). Proteins were fractionated using an isocratic gradient (DPBS) running at 0.5 mL $min^{-1}$ into 5 x 2 mL fractions, beginning at 8 mL and ending at 18 mL. Eluate was collected into 15 mL tubes containing 0.5 mL of PBS with 1% NP-40 and containing 1% NP-40 and cOmplete Protease Inhibitor Cocktail (Roche). SEC fractions were incubated with washed ANTI-FLAG® M2 Affinity Gel (Sigma-Aldrich, #A2220; 60 µL of 50% slurry/sample) overnight at 4 °C with rotation. After incubation, the beads were pelleted (2 min at 2,000 x *g*) and washed with 0.2% NP-40 in DPBS (3 x 1 mL), followed by DPBS (1 x 1 mL). The beads were resuspended in 50 µL DPBS and treated with 5.5 µL of click Master-mix for 1 h, followed by the addition 4X SDS gel loading buffer (18 µL). Proteins were resolved by SDS-PAGE gel (160 V, 1.25 h, 4–20% Tris-glycine) and visualized as described above, followed by Western blotting.

**Abundance-based SEC analysis**

$HCT116_{TRMT112\text{-}FLAG}$ cell pellets were seeded the day prior to the experiment, scraped in cold DPBS and washed twice with cold DPBS, and processed as described above (SEC ABPP experiment). The eluate from the FPLC was collected into tubes containing each 12 mL of acetone at 4 °C. Proteins were precipitated overnight at -20 °C, then centrifuged at 4,500 x *g* for 20 min, and the acetone/DPBS mixture was decanted off the pellets. The samples were dried at room temperature and resuspended in 1X SDS gel loading buffer (100 µL), followed by water-bath sonication and heating to 98 °C for 5 min. Proteins were resolved by SDS-PAGE gel (160 V, 1.25 h, 4–20% Tris-glycine) and visualized as described above, followed by Western blotting.

**Protein-purification**

Cloning

The full-length human TRMT112 and METTL5 genes were codon-optimized for *E. coli* cell expression were synthesized from IDT. The ORF of TRMT112 was cloned into a pACYCDuet-1 vector via NdeI (NEB, #R0111S) and BglII (NEB, #R0144S) restriction and Gibson Assembly (NEB, # E2621S). Mutagenesis was carried out using a Q5 site-directed mutagenesis kit (NEB, E0554S), using the following primers.

TRMT112_C100A_FWD: AACTTTGCAGgcgCCAGAATCTGGG

TRMT112_C100A_REV: CCCAGATTCTGGcgcCTGCAAAGTT

The ORF of METTL5 was synthesized from IDT and was PCR amplified with primer containing a N-terminal His tag followed by a TEV protease cleavage site (for ORF see Supplementary Table 2), using the following primers. The PCR product was cloned into a pET21a vector via NdeI (NEB, #R0111S) and NotI (NEB, #R0189S) restriction and Gibson Assembly (NEB, # E2621S).

His-METTL5_Gibson_FWD:

taactttaagaaggagatatacATATGCACCACCACCACCACCACGAAAATTTGTATTTCCAG
TCAATGAAGAAGGTACGTCTG

His-METTL5_Gibson_REV:
GGTGGTGCTCGAGTGCGGCCGCTCAAAAACTGAAGCGGATCA

Protein-expression of TRMT112:His-METTL5

With modifications to a previously described procedure[3], the TRMT112 and His-METTL5 complex was co-expressed in *E. coli* BL21 (DE3). *E. coli* were grown in 1 L of culture in auto-inducible Terrific Broth (Formedium, #AIMTB0260) containing carbenicillin (50 µg $mL^{-1}$) and chloramphenicol (34 µg $mL^{-1}$). The culture was initially incubated at 37°C for 3.5 h and then at 18°C overnight. The cells were harvested by centrifugation at 4,000 x *g* for 30 min and flash frozen in liquid nitrogen. Cell pellets were resuspended in 30 mL cold lysis buffer (50 mM Tris-HCl pH 7.5, 200 mM NaCl, 1 mM TCEP with cOmplete Protease Inhibitor Cocktail (Roche) and Lysozyme). The cells were lysed by pulse-sonication on ice and the lysate was cleared by centrifugation at 20,000 x *g* for 45 min. The supernatant was incubated with pre-washed HisPur™ Ni-NTA Resin (Thermo, #88221; 1 mL of 50% slurry) and 10 mM imidazole at 4°C for 1 h with rotation. The beads were washed in a gravity-flow column with 10 mL of cold wash buffer 1 (50 mM Tris-HCl pH 7.5, 1 M NaCl, 1 mM TCEP, 10 mM imidazole), 10 mL wash buffer 2 (50 mM Tris-HCl pH 7.5, 200 M NaCl, 1 mM TCEP, 10 mM imidazole) and 10 mL wash buffer 3 (50 mM Tris-HCl pH 7.5, 200 M NaCl, 1 mM TCEP, 20 mM imidazole). Proteins were eluted from beads with 5 mL elution buffer (50 mM Tris-HCl pH 7.5, 200 M NaCl, 1 mM TCEP, 400 mM imidazole). Proteins were dialyzed using a Slide-A-Lyzer™ Dialysis Cassettes, 10K MWCO (Thermo, #66830) into anion-exchange buffer A (50 mM Tris-HCl pH 7.5, 100 mM NaCl, 0.5 mM TCEP) overnight at 4°C.

Proteins were injected into a HiTrap Q FF anion exchange chromatography column (Cytiva, #17515601) attached to an ÄKTA pure FPLC system (Cytiva). Proteins were fractionated using a linear gradient from anion-exchange buffer A to anion-exchange buffer B (50 mM Tris-HCl pH 7.5, 1 M NaCl, 0.5 mM TCEP) over 8 column volumes at 5 mL $min^{-1}$. Eluate was collected in 500 µL fractions. TRMT112:METTL5 containing fractions were combined, concentrated with an Amicon® Ultra Centrifugal Filter, 3 kDa MWCO (Milipore, #UFC900308) and then injected into a Superdex 200 Increase 10/300

GL column attached to an ÄKTA pure FPLC system (Cytiva). Proteins were fractionated using an isocratic gradient (50 mM Tris-HCl pH 7.5, 200 mM NaCl, 1 mM TCEP) running at 0.5 mL $min^{-1}$ and collected in 500 µL fractions. TRMT112:METTL5 containing fractions were combined quantified by NanoDrop and flash frozen in liquid nitrogen. Typical yields were 5 to 10 mg of protein-complex per 1 L culture.

Protein-expression of tag-free TRMT112:METTL5

Tag-free TRMT112:METTL5 was expressed from the same constructs as described above. After the dialysis step, 0.5 mg His-tagged TEV protease (gift from Vividion Therapeutics) was added, and the protein was incubated at 4 °C for 1 day. The protein was then incubated with pre-washed HisPur™ Ni-NTA Resin (Thermo, #88221; 1 mL of 50% slurry) at 4°C for 1 h with rotation. The supernatant was filtered using a gravity-flow column and further processed as described above (Protein-expression of TRMT112:His-METTL5).

**Intact protein mass spectrometry and rate determination**

Purified 0.5 µM TRMT112:His-METTL5 complex was incubated with 5 µM FWG-33B in 50 mM Tris-HCl pH 7.5, 200 mM NaCl, 1 mM DTT, 5% DMSO. Reactions were stopped after 5 and 10 min by addition of formic acid.

The observed rate ($k_{obs}$/[I]) was calculated assuming pseudo-first-order reaction kinetics:

(eq. 3) $d[TRMT112:METTL5]/dt = -k \cdot [TRMT112:METTL5]$

(eq. 4) $[TRMT112:METTL5]_t = [TRMT112:METTL5]_{t0} \cdot e^{-kt}$.

Samples were analyzed on an Agilent LC1290 Infinity II instrument coupled to a 6545 QTOF liquid chromatography–mass spectrometer (Agilent Technologies). A sample volume of 10 µL was injected. The protein was desalted and separated on an AERIS 3.6-µM-wide-bore XB-C8 liquid chromatography column (50 x 2.1 $mm^2$, Phenomenex) at 60 °C at a flow rate of 0.5 mL $min^{-1}$. Mass spectra were acquired from 700 to 1,700 Da at a resolution of 25,000. A Dual Agilent Jet Stream Electrospray Ionization Source was used for ionization. The resulting data files were deconvoluted to protein masses using Agilent MassHunter BioConfirm Software, v.11.0.

**Protein Crystallization and Structure Determination**

Tag-free 20 µM TRMT112:METTL5 complex was incubated with 100 µM FWG-33B in 50 mM Tris-HCl pH 7.5, 200 mM NaCl, 1 mM TCEP, 2% DMSO, and 1 mM S-(5′-Adenosyl)-L-methionine chloride dihydrochloride (SAM) (Sigma-Aldrich, #A7007) and the reaction was monitored by intact protein MS. Upon completion (<30 min), the protein sample was buffer exchanged into 50 mM Tris-HCl pH 7.5, 200 mM NaCl, 1 mM TCEP using a PD-10 desalting column (Cytvia, #17085101) and concentrated to 24 mg $mL^{-1}$. Initial crystals were identified in numerous conditions but diffracted poorly. These were used for matrix micro-seeding into commercial screens. Crystals used in diffraction experiments were grown in drops consisting of 1:1, 1:2 and 2:1 ratios of protein to reservoir solution equilibrated against 21% PEG3350, 0.36 M ammonium sulfate, 0.1 M Bis-Tris pH 5.5 at 4 °C. Crystals were cryoprotected by rapid transfer into reservoir solution supplemented with 30% glycerol before flash freezing in liquid nitrogen. Diffraction data were collected on Advanced Light Source Beamline 5.0.2 and processed with XDS. The structure was determined by molecular replacement in Phaser using TRMT112 and METTL5 from PDB ID 6H2U as independent search models. The structure was refined using iterative rounds of refinement in REFMAC5 with manual inspection and model building in COOT. Ligand restraints for FWG-33B and the covalent bond with Cys100 were generated in JLigand. Waters were automatically added in COOT and REFMAC5 and manually inspected. Data collection and refinement statistics can be found in Supplementary Table 1.

**Binding-site mapping of TRMT112 complexes**

Binding-site detection was performed on crystallographic structures, using SiteMap on Schrödinger Maestro (version 13.2.128, MMshare Version 5.8.128, Release 2022-2, Platform Darwin-x86_64).

**Gel-ABPP with purified TRMT112:METTL5**

49 µL of purified 1 µM TRMT112:His-METTL5 complex in DPBS containing 1 mM TCEP and 1 mM SAM were treated with DMSO or competitor for 1 h (2% final DMSO), followed by treatment with alkyne for 1 h. 5.5 µL of click Master-mix were then added, as described

above. Proteins were resolved by SDS-PAGE gel (160 V, 1.25 h, 4–20%) and visualized as described above.

***In vitro* methyltransferase assay**

rRNA probes (12 nt and 60 nt; **Fig. 5b**) were synthesized from IDT.

Purified 20 μM WT-TRMT112:His-METTL5 or C100A-TRMT112:His-METTL5 complex was incubated with 40 μM FWG-33B in reaction buffer (50 mM Tris-HCl pH 7.5, 200 mM NaCl, 1 mM DTT, 5% DMSO, and 1 mM S-(5′-Adenosyl)-L-methionine chloride dihydrochloride (SAM) (Sigma-Aldrich, #A7007)) for 30 min at r.t.. 1 μL (20X, final 1 μM) of this protein mixture was then diluted into 19 μL of the substrate assay (final concentrations: Tris-HCl, pH 7.5, 50 mM, NaCl 50 mM, $MgCl_2$ 5 mM, 1 mM DTT, 1 U SUPERase·In™ RNase Inhibitor (Invitrogen, #AM2694), 10 μM 12 nt RNA probe, 1 mM SAM) for 1 or 3 h at 37 °C. The samples were then diluted with 350 μL cold IP buffer (50 mM Tris-HCl pH 7.5, 150 mM NaCl, 0.1% NP-40) and incubated with pre-washed Dynabeads™ MyOne™ Streptavidin T1 (Invitrogen, #65601; 20 μL of slurry/sample). After incubation, the beads were isolated with a magnetic stand and washed with IP buffer (2 x 1 mL), followed by HPLC-grade water (1 x 1 mL). The eluted RNA was then digested by nucleoside digestion mix (NEB, #M0649S). The digested nucleosides were subjected to LC-MS analysis.

The experiment with the 60 nt probe was performed as described above. After the time points, the liquid was then subjected to Monarch® Spin RNA Cleanup Kit (10 μg) (NEB, #T2030L) and eluted with 17 μL water, and digested overnight as described above

The nucleosides were analyzed by LC/MS–based multiple reaction monitoring (MRM) (Agilent Technologies 6460 or 6470 Triple Quad). MS analysis was performed using positive mode ESI with the following parameters: drying gas temperature, 350°C; drying gas flow, 9 $L^{-1}$; nebulizer pressure, 45 psi; sheath gas temperature, 375°C; sheath gas flow, 10 = L $min^{-1}$; fragmentor voltage, 135 V; and capillary voltage, 4.5 kV. The separation of the analyte was achieved using a 50 mm × 4.6 mm 5 μm Gemini C18 column (Phenomenex) coupled to a guard column (Gemini: C18: 4 × 3 mm). The LC solvents were as follows: solvent A, 5 mM $NH_4OAc$; and solvent B, acetonitrile. The LC gradient following injection increase from 0% to 5% B at 0.5 mL $min^{-1}$ over 5 min; then increase to 90% B at 0.5 mL $min^{-1}$ over 10 min; decrease to 0% B at 0.5 mL $min^{-1}$ over 5 min. *N*6-Methyladenosine (282.1 *m/z* → 150.1 *m/z*), adenosine (268.1 *m/z* → 136.0 *m/z*). Adenosine and *N*6-methyladenosine concentrations were calculated from external calibration curves.

**Statistics and reproducibility**

Statistical analyses and data visualization in this paper were performed using GraphPad Prism 9 (v.10.4.1). To compare group means, we performed multiple unpaired *t*-tests between group pairs, using the Holm–Šídák method to correct for multiple comparisons; reported p values are adjusted accordingly. For experiments comparing multiple treatments to a single treatment group, we used one-way ANOVA followed by Dunnett's post hoc test.

### References


1. Ogasawara, D. et al. Chemical tools to expand the ligandable proteome: Diversity-oriented synthesis-based photoreactive stereoprobes. *Cell. Chem. Biol.* **31**, 2138-2155.e32 (2024).
2. Njomen, E. et al. Multi-tiered chemical proteomic maps of tryptoline acrylamide–protein interactions in cancer cells. *Nat. Chem.* **16**, 1592-1604 (2024).
3. Van Tran, N. et al. The human 18S rRNA m6A methyltransferase METTL5 is stabilized by TRMT112. *Nucleic Acids Res.* **47**, 7719–7733 (2019).
