## Supplementary Chemistry Information for "Complexoform-restricted covalent TRMT112 ligands that allosterically agonize METTL5"

### Synthetic procedure and analytical data

**Table of Content**


| | |
|---|---|
| General Methods | p. 3 – 4 |
| Representative Procedures | p. 5 – 8 |
| Analytical Data for intermediates and FWG-1A/B, FWG-2A/B, FWG-3A/B and FWG-4A/B | p. 9 – 23 |
| Analytical Data for FWG-9A/B and FWG-11A/B | p. 24 – 27 |
| Analytical Data for Intermediates and FWG-33A/B and FWG-49A/B | p. 28 – 34 |
| Analytical Data for Intermediates and FWG-19B, FWG-21B and FWG-23B | p. 35 – 42 |
| Analytical Data for Intermediates and FWG-35B, FWG-41B, FWG-43B, FWG-45B and FWG-47B | p. 43 – 59 |
| $^{1}H$, $^{13}C$ and $^{19}F$ NMR Spectra | p. 60 – 112 |
| SFC Traces | p. 113 – 136 |
| References | p. 137 |

#### General methods

The following commercially available reagents were purchased from Sigma Aldrich: $[Rh(cod)OH]_2$, aq. CsOH (50 wt%), $[Rh(PPh_3)_3Cl]$, $Cs_2CO_3$, HBpin, KHMDS solution in toluene (0.5 M), THF (≥99.9%, anhydrous, inhibitor-free), DMF (≥99.8%, anhydrous), Pd/C, pyridine (anhydrous, 99.8%), trifluoroacetic acid, dichloromethane (anhydrous, ≥99.8%, contains 40-150 ppm amylene as stabilizer), ethyl acetate (anhydrous, 99.8%), *N,N*-diisopropylethylamine (≥99%), acryloyl chloride (≥97%, contains ~400 ppm phenothiazine as stabilizer). (*S*)-Segphos and (*R*)-Segphos were bought from TCI. Boronic acids and esters were bought from Combi-blocks. All listed commercially available reagents were used without further purification. Allyl chloride (±)-**1** was synthesized according to literature protocols.[1]

Preparative thin layer chromatography (prep. TLC) was performed on TLC silica gel 60G $F_{254}$ 25 glass plates (60F-254). Normal phase automated medium-pressure chromatography (MPLC) was performed on Teledyne ISCO CombiFlash NextGen 300+ using RediSep Gold® or Silver® cartridges. Reverse-phase automated medium-pressure chromatography (RP-MPLC) was performed on Teledyne ISCO CombiFlash NextGen 300+ using RediSep Gold® Reverse-phase C18 cartridges. Reverse-phase automated high-pressure chromatography (RP-HPLC) was performed on Teledyne ISCO ACCQPrep HP150 system using a RediSep® Prep C18 Column (20 mm x 150 mm).

Nuclear magnetic resonance (NMR) spectroscopy measurements were carried out at room temperature. NMR spectra were recorded on the following instruments: Bruker AVIII HD 600 MHz NMR (equipped with 5 mm CPQCI and 1.7 mm CPTCI CryoProbes), Bruker AVIII HD 600 MHz NMR (equipped with 5 mm CPDCH CryoProbe), Bruker AV NEO 500 MHz NMR (equipped with 5 mm BBFO Probe) or Bruker AV NEO 399 MHz NMR (equipped with 5 mm BBFO Probe). Chemical shifts (δ) are reported in ppm relative to the residual solvent peak ($CHCl_3$, 7.26 ppm for $^1H$ NMR and 77.16 ppm for $^{13}C$ NMR) with corresponding coupling constants (*J*) in Hertz (Hz) and apparent multiplicities (i.e., s: singlet, d: doublet, t: triplet, q: quartet, m: multiplet, app, or a combination of these multiplicities). The $^{19}F$ chemical shifts are not referenced.

Mass measurements using high-resolution mass spectrometry (HRMS) were performed on a Waters Xevo G2-XS TOF instrument calibrated against sodium formate clusters and using a LeuEnk lockmass. Expected monoisotopic masses were calculated using MassLynx 4.1 and the m/z values for calibrant and lockmass were MassLynx-default values.

Chiral SFC (supercritical fluid chromatography) separations were conducted on a Waters Acquity UPC2 system using Daicel columns as stationary phase and Waters Empower software. Racemic samples of (±)-**3**, (±)-**41**, (±)-**52**, (±)-**55**, (±)-**58** and (±)-**61** were obtained by following **Procedure A** using *rac*-BINAP instead of Segphos. Racemic samples of **FWG-1A/B**, **FWG-2A/B**, **FWG-3A/B**, **FWG-4A/B**, **FWG-9A/B**, **FWG-11A/B**, **FWG-33A/B** and **FWG-49A/B** were obtained by mixing both enantiomers in a 1:1 ratio.

The absolute and relative stereochemistry was initially assigned by analogy to previous literature reports.[1,2] Our protein co-crystal structure confirms this stereochemistry for **FWG-33B**.

All final compounds were >95% pure by LC-MS (UV) and showed >95% ee, either as intermediates, as final compounds or both.

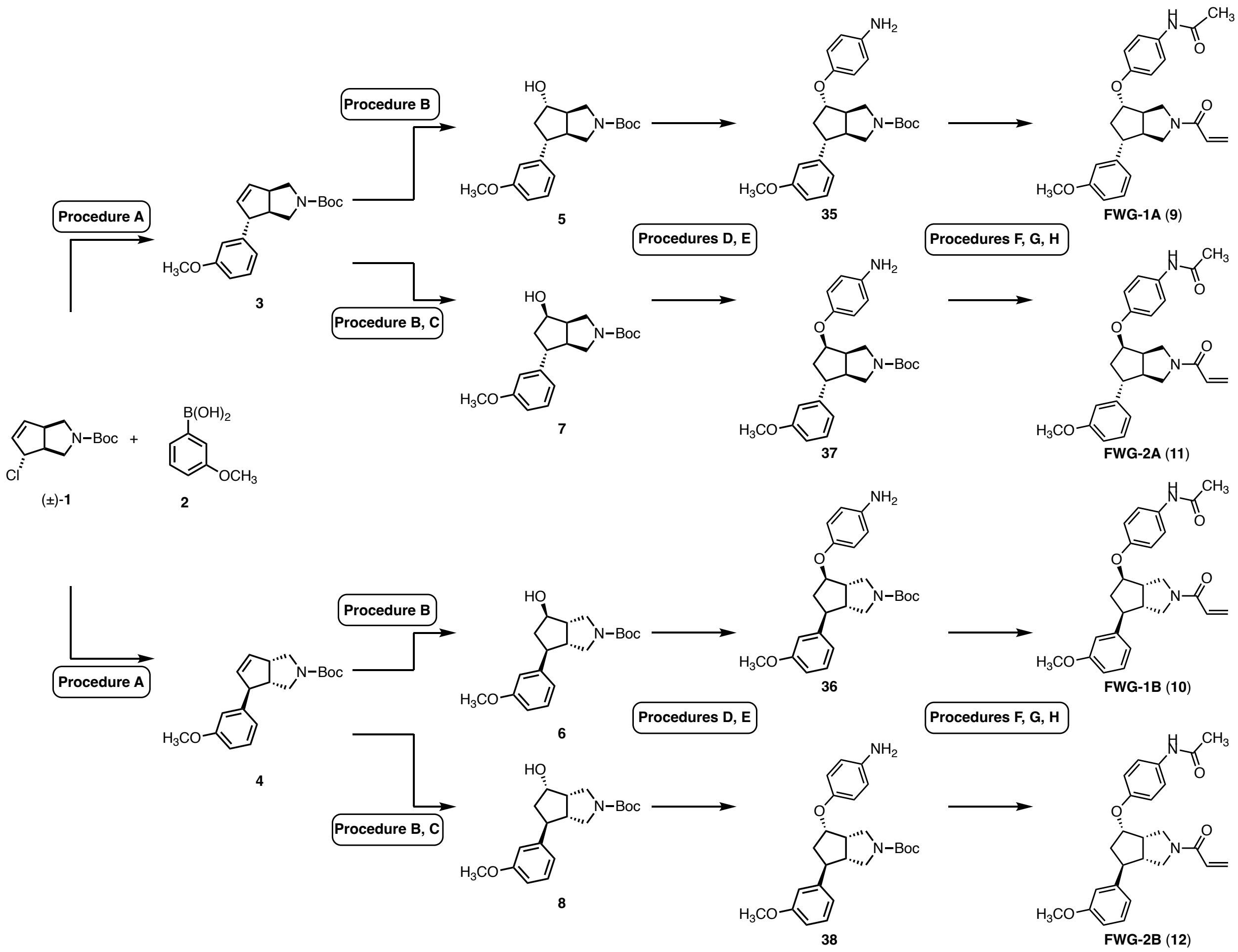


**Figure S1:** Representative synthesis scheme for bicyclopyrrolidine acrylamides.

Representative synthetic procedures corresponding to the transformations shown in Fig. S1 are found below. Analytical data is provided for specific compounds.

##### Procedure A

This is a representative procedure on a 4 mmol scale:

$[Rh(cod)OH]_2$ (45.6 mg, 0.1 mmol, 2.5 mol%) and (*S*)-Segphos (146 mg, 0.24 mmol, 6.0 mol%) were added to a 50 mL round bottom flask, sealed with a rubber septum under a $N_2$ atmosphere, dissolved in THF (7.2 mL) and stirred at room temperature. CsOH (50 wt% aq. solution, 700 µL, 4.0 mmol, 1.0 eq) was added and the mixture was heated to 60 °C. After 30 min, a solution (or suspension) of boronic acid (or pinacol ester) (12.0 mmol, 3.0 eq) and allylic chloride (±)-**1** (4.0 mmol, 1.0 eq) in THF (8.8 mL) and $H_2O$ (2.0 mL) was added via syringe and the flask was rinsed with THF (2.0 mL). The resulting mixture was then stirred at 60 °C overnight. The mixture was then cooled to room temperature before $SiO_2$ (2 g) was added and the solvents were removed under reduced pressure. Purification by automated medium pressure liquid chromatography (hexane:EtOAc 100:0 to 60:40) afforded the desired product **3**.

**Procedure B**

This is a representative procedure on a 2.95 mmol scale:

A mixture of **3** (2.95 mmol, 1.0 eq), $Cs_2CO_3$ (961 mg, 2.95 mmol, 1.0 eq) and $[Rh(PPh_3)_3Cl]$ (273 mg, 0.295 mmol, 10 mol%) were added to a 25 mL round bottom flask, sealed with a rubber septum under a $N_2$ atmosphere, suspended in THF (5.9 mL) and stirred at room temperature. HBpin (860 µL, 5.9 mmol, 2.0 eq) were added dropwise and the mixture was stirred at r.t. overnight. The mixture was then cooled to 0 °C. An aq. solution of $H_2O_2$ (3.0 mL), followed by a 2 *N* solution of NaOH (7.3 mL) were added and the resulting mixture was stirred for 1 h. The reaction mixture was extracted with EtOAc (3 x 25 mL). The combined organic layers were washed with brine (25 mL), dried over $MgSO_4$ and concentrated under reduced pressure. Purification by automated medium pressure liquid chromatography (hexane:EtOAc 100:0 to 50:50) afforded the desired product **5**.

**Procedure C**

This is a representative procedure on a 750 µmol scale:

A mixture of **5** (750 µmol, 1.0 eq), $PPh_3$ (392 mg, 1.5 mmol, 2.0 eq) and 4-nitrobenzoic acid (250 mg, 1.5 mmol, 2.0 eq) were added to a 25 mL round bottom flask, sealed with a rubber septum under a $N_2$ atmosphere, and dissolved in dry THF (6.0 mL). The solution was cooled to 0 °C and diisopropyl azodicarboxylate (300 µL, 1.5 mmol, 2.0 eq) were added dropwise. The mixture was allowed to reach r.t. overnight. An aq. solution of $NaHCO_3$ (20 mL) was added. The reaction mixture was extracted with EtOAc (2 x 50 mL). The combined organic layers were washed with brine (20 mL), dried over $MgSO_4$ and concentrated under reduced pressure. Purification by automated medium pressure liquid chromatography (hexane:EtOAc 100:0 to 50:50) afforded a mixture containing the 4-nitrobenzoic ester (LC-MS). This crude mixture was suspended in MeOH (7.0 mL) and $H_2O$ (0.5 mL) in a 25 mL round bottom flask. $K_2CO_3$ (207 mg, 2.0 eq) was added and the resulting mixture was stirred under an open atmosphere at r.t. overnight. The mixture was concentrated under reduced pressure, resuspended in EtOAc (10 mL), filtered, and concentrated under reduced pressure. Purification by automated medium pressure liquid chromatography (hexane:EtOAc 100:0 to 50:50) afforded the desired product **7**.

**Procedure D**

This is a representative procedure on a 750 µmol scale:

*Caution:* This reaction is sensitive to the quality of DMF and exotherms during addition (cooling is required).

A solution of alcohol **6** (750 µmol, 1.0 eq) was stirred in dry DMF (3.8 mL) in a 2-dram vial equipped with a septum under a $N_2$ atmosphere. The mixture was cooled to 0 °C and solution of KHMDS in toluene (0.5 M, 2.3 mL, 1.13 mmol, 1.5 eq) was added dropwise. After 10 min, 4-nitrofluorobenzene (95 µL, 900 µmol, 1.2 eq) was added dropwise. The reaction mixture was allowed to reach r.t. overnight. Then, a sat. aq. solution of $NaHCO_3$ (2 mL), followed by $H_2O$ (3 mL) were added. The mixture was extracted with EtOAc (3 x 10 mL). The combined organic layers were washed with brine (10 mL), dried over $MgSO_4$ and concentrated under reduced pressure. Purification by automated medium pressure liquid chromatography (hexane:EtOAc 100:0 to 50:50) afforded the desired product **32**.

**Procedure E**

This is a representative procedure on a 100 µmol scale:

*Caution:* We noticed that extended reaction times (overnight) give lower yields of the desired anilines.

A solution of nitrobenzene **31** (262 µmol, 1.0 eq) in MeOH (5.2 mL) was added to Pd/C (12 mg, 10wt%) in a 10 mL flask, sealed with a rubber septum under a $N_2$ atmosphere. A balloon filled with $H_2$ was then attached and bubbled through the reaction mixture for 15 min. The reaction mixture was then sealed and allowed to continue incubating under the $H_2$ atmosphere for the indicated amount of time. The reaction mixture was then filtered over Celite and concentrated under reduced pressure. The product was either used crude without any additional purification or purified by automated medium pressure liquid chromatography (hexane:EtOAc 100:0 to 30:70) to afford aniline **35**.

**Procedure F1**

This is a representative procedure on a 50 µmol scale:

Pyridine (140 µL, 1.7 mmol, 35 eq.) and acetic anhydride (95 µL, 1.0 mmol, 20 eq.) were added to aniline **35** (50 µmol, 1.0 eq) in 1-dram vial and the mixture was stirred at r.t. overnight. The reaction mixture was concentrated by blowing an airflow over the solution, diluted with a sat. aq. solution of $NaHCO_3$ (2 mL) and $H_2O$ (3 mL). The mixture was extracted with EtOAc (3 x 10 mL). The combined organic layers were washed with brine (10 mL), dried over $MgSO_4$ and concentrated under reduced pressure. The product was used crude without any additional purification step.

**Procedure F2**

This is a representative procedure on a 50 µmol scale:

A mixture of hexafluorophosphate azabenzotriazole tetramethyl uronium (60 µmol, 1.2 eq) and carboxylic acid (55 µmol, 1.1 eq) and *N*,*N*-diisopropylethylamine (16 µL, 125 µmoL, 2.5 eq) in dichloromethane (330 µL) was stirred for 10 min at r.t. under an open atmosphere in a 1-dram vial. Then, a solution of aniline **35** (50 µmol, 1.0 eq) in dichloromethane (170 µL) was added. The vial was flushed with $N_2$ and stirred overnight at r.t.. A sat. aq. solution of $NaHCO_3$ (2 mL) and then $H_2O$ (3 mL) were added. The mixture was extracted with EtOAc (3 x 10 mL). The combined organic layers were washed with brine (10 mL), dried over $MgSO_4$ and concentrated under reduced pressure. The product was used crude without any additional purification step.

**Procedure G**

This is a representative procedure on a 50 µmol scale:

Trifluoroacetic acid (170 µL, 2.2 mmol, 22 eq.) was added to a solution of the crude product from **Procedure F1** or **F2** (100 µmol, 1.0 eq) in dichloromethane (330 µL) in a 1-dram vial. The mixture was stirred for 1 h at r.t.. The reaction mixture was then concentrated under blowing an airflow over the solution, followed by reduced pressure (<5 mbar). The product was used crude without any additional purification step.

**Procedure H**

This is a representative procedure on a 50 µmol scale (the eluent used in preparative chromatography is provided with the analytical data):

Dichloromethane was added to the crude product from **Procedure G** (50 µmol, 1.0 eq) in a 1-dram vial. *N*,*N*-diisopropylethylamine (34 µL, 200 µmoL, 4 eq) was added, followed by dropwise addition of acroyl chloride (8 µL, 100 µmoL, 2 eq). The vial was flushed with a $N_2$ atmosphere and stirred for 2 h at r.t.. The reaction mixture was then directly loaded and purified by preparative thin-layer chromatography. Subsequent reverse-phase purification afforded the desired acrylamides.

Synthesis of **3** and **4**

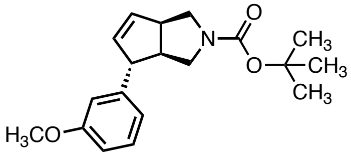

**3**

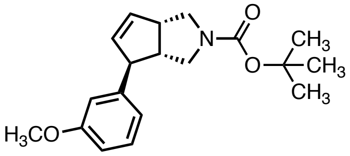

**4**

Compound **3** was synthesized according to **Procedure A** (4.0 mmol scale) using 3-methoxyphenylboronic acid and was obtained as a pale-yellow oil (943 mg, 75%) with 99% ee.

The enantiomer **4** was synthesized according to **Procedure A** (4.0 mmol scale; using (*R*)-Segphos) using 3-methoxyphenylboronic acid and was obtained as a pale-yellow oil (916 mg, 73%) with 97% ee.

**$^1$H NMR** (600 MHz, $CDCl_3$, **3**) δ 7.22 (app. t, *J* = 7.9 Hz, 1H), 6.78 – 6.74 (app. dt, *J* = 8.3, 2.3 Hz, 2H), 6.70 (app. t, *J* = 2.1 Hz, 1H), 5.82 (s, 1H), 5.79 – 5.72 (m, 1H), 3.79 (s, 3H), 3.68 (dd, *J* = 11.2, 8.7 Hz, 2H), 3.53 – 3.19 (m, 4H), 2.77 (s, 1H), 1.46 (s, 9H) ppm.

**$^1$H NMR** (600 MHz, $CDCl_3$, **4**) δ 7.22 (app. t, *J* = 7.9 Hz, 1H), 6.79 – 6.74 (m, 2H), 6.70 (app. t, *J* = 2.1 Hz, 1H), 5.82 (dt, *J* = 5.7, 1.8 Hz, 1H), 5.76 (dd, *J* = 5.7, 2.2 Hz, 1H), 3.79 (s, 3H), 3.72 – 3.65 (m, 2H), 3.56 – 3.39 (m, 3H), 3.34 – 3.24 (m, 1H), 2.77 (dtd, *J* = 8.7, 5.9, 2.9 Hz, 1H), 1.46 (s, 9H) ppm.

**$^{13}$C NMR** (151 MHz, $CDCl_3$, **3**) 160.0, 154.6, 146.0, 134.6*, 134.2*, 133.7*, 129.7, 119.7, 113.1, 111.7, 79.4, 57.9, 55.3 (3C), 52.5$^†$, 52.1$^†$, 51.0$^†$, 50.2$^†$, 49.9$^†$, 49.7$^†$, 48.8$^†$, 28.7 (3C) ppm (in total 4 additional rotameric peaks detected).

* rotamers; corresponding to 2C

$^†$ rotamers; corresponding to 4C

**HRMS** (**3**) m/z calc. for $C_{14}H_{18}NO^+$ $[M\text{-}Boc+2H]^+$ 216.1388 found 216.1392.

**HRMS** (**4**) m/z calc. for $C_{19}H_{25}NNaO_3^+$ $[M+Na]^+$ 338.1732 found 338.1743.

**SFC** (Waters UPC2 SFC with a Daicel IA column (3 µm, 4.6x250 mm) under isocratic conditions (3.3 mL/min, 15% MeOH / $CO_2$, 1600 psi backpressure) at 30 °C) ($t_R$ (**3**) = 1.70 min; $t_R$ (**4**) = 2.48 min).

Synthesis of **5** and **6**

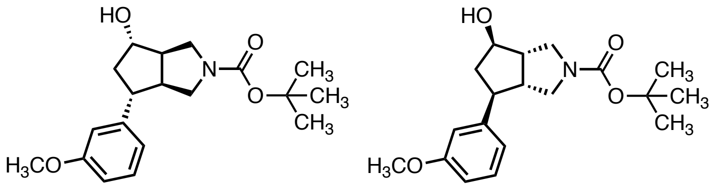

**5** **6**

Compound **5** was synthesized according to **Procedure B** (2.95 mmol scale) and was obtained as a highly viscous pale-yellow oil (501 mg, 51%).

The enantiomer **6** was synthesized according to **Procedure B** (2.75 mmol scale) and was obtained as a colorless oil (440 mg, 48%).

**$^1$H NMR** (600 MHz, $CDCl_3$, **5**) δ 7.23 (app. t, *J* = 8.0 Hz, 1H), 6.86 (d, *J* = 7.6 Hz, 1H), 6.82 (s, 1H), 6.77 (dd, *J* = 8.2, 2.5 Hz, 1H), 4.18 (s, 1H), 3.80 (s, 3H), 3.61 – 3.23 (m, 4H), 2.81 (s, 2H), 2.67 (dt, *J* = 8.6, 3.8 Hz, 1H), 2.52 (app. h, *J* = 4.2 Hz, 1H), 1.89 (ddd, *J* = 12.9, 11.0, 7.6 Hz, 1H), 1.46 (s, 9H) ppm (OH proton not detected).

**$^1$H NMR** (600 MHz, $CDCl_3$, **6**) 7.23 (app. t, *J* = 7.9 Hz, 1H), 6.89 – 6.84 (m, 1H), 6.82 (app. t, *J* = 2.1 Hz, 1H), 6.78 – 6.72 (m, 1H), 4.17 (dt, *J* = 7.9, 5.5 Hz, 1H), 3.80 (s, 3H), 3.52 (dd, J = 11.5, 8.7 Hz, 1H), 3.44 – 3.36 (m, 2H), 3.32 (dd, *J* = 11.5, 6.5 Hz, 1H), 2.88 – 2.74 (m, 2H), 2.67 (tt, *J* = 8.5, 4.9 Hz, 1H), 2.51 (dt, *J* = 12.6, 6.2 Hz, 1H), 2.35 – 2.05 (br. m, 1H), 1.89 (ddd, *J* = 13.1, 10.8, 7.5 Hz, 1H), 1.46 (s, 9H) ppm.

**$^{13}$C NMR** (151 MHz, $CDCl_3$, **5**) 159.9, 155.0, 145.6, 129.7, 119.8, 113.5, 111.6, 79.6, 77.9, 55.3, 52.5$^†$, 51.6$^†$, 51.0$^†$, 50.5$^†$, 50.0$^†$, 49.7$^†$, 49.3, 44.7, 28.7 (3C) ppm (in total 2 additional rotameric peaks detected).

$^†$ rotamers; corresponding to 4C

**HRMS** (**5**) m/z calc. for $C_{14}H_{20}NO_2^+$ $[M\text{-}Boc+2H]^+$ 234.1494 found 234.1500.

**HRMS** (**6**) m/z calc. for $C_{14}H_{20}NO_2^+$ $[M\text{-}Boc+2H]^+$ 234.1494 found 234.1502.

Synthesis of **7** and **8**

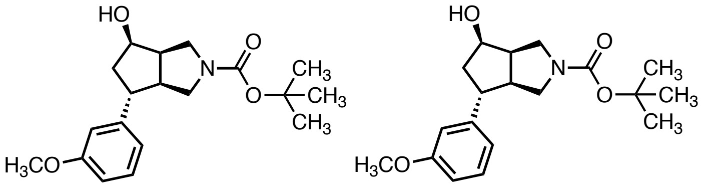

**7** **8**

Compound **7** was synthesized according to **Procedure C** (750 µmol scale) and was obtained as a highly colorless oil (170 mg, 68%).

The enantiomer **8** was synthesized according to **Procedure C** (900 µmol scale) and was obtained as a colorless oil (203 mg, 68%).

**$^{1}$H NMR** (600 MHz, $CDCl_3$, **7**) δ 7.20 (app. t, *J* = 7.9 Hz, 1H), 6.79 (d, *J* = 7.7 Hz, 1H), 6.76 – 6.66 (m, 2H), 4.44 (d, *J* = 6.9 Hz, 1H), 3.77 (s, 3H), 3.74 – 3.70 (m, 1H), 3.47 (dd, *J* = 11.2, 7.9 Hz, 1H), 3.41 – 3.03 (m, 4H), 2.87 (dt, *J* = 9.3, 4.2 Hz, 1H), 2.67 (d, *J* = 9.9 Hz, 1H), 2.25 (ddd, *J* = 13.5, 7.5, 3.0 Hz, 1H), 2.05 – 1.95 (m, 1H), 1.44 (s, 9H) ppm.

**$^{1}$H NMR** (600 MHz, $CDCl_3$, **8**) δ 7.23 (app. td, *J* = 7.6, 1.0 Hz, 1H), 6.81 (d, *J* = 7.6 Hz, 1H), 6.78 – 6.71 (m, 2H), 4.47 (qd, *J* = 5.2, 2.9 Hz, 1H), 3.80 (s, 3H), 3.76 – 3.69 (m, 1H), 3.48 (dd, *J* = 11.2, 8.0 Hz, 1H), 3.40 – 3.31 (m, 2H), 3.15 (dt, *J* = 10.7, 7.5 Hz, 1H), 2.90 (tt, *J* = 9.5, 5.1 Hz, 1H), 2.77 – 2.66 (m, 1H), 2.25 (ddd, *J* = 13.5, 7.4, 2.8 Hz, 1H), 2.11 – 1.95 (br. m, 2H), 1.45 (s, 9H) ppm.

**$^{13}$C NMR** (151 MHz, $CDCl_3$, **7**) δ 159.9, 154.9, 146.4, 129.7, 119.7, 113.4, 111.4, 79.5, 73.9, 55.3 (3C), 52.2, 50.8, 49.1, 48.6, 45.4, 44.9, 28.7 (3C) ppm.

**HRMS** (**7**) m/z calc. for $C_{19}H_{27}NNaO_4^+$ $[M+Na]^+$ 356.1838 found 356.1851.

**HRMS** (**8**) m/z calc. for $C_{19}H_{27}NNaO_4^+$ $[M+Na]^+$ 356.1838 found 356.1838.

Synthesis of **31** and **32**

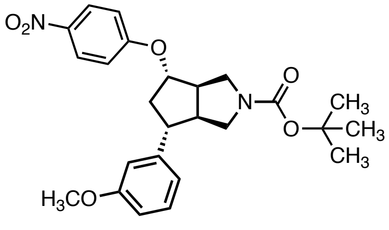

**31**

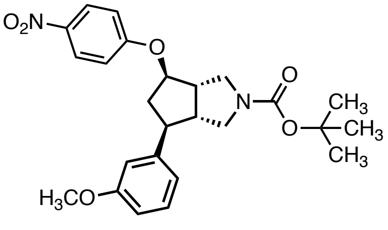

**32**

Compound **31** was synthesized according to **Procedure D** (300 μmol scale) using 1-fluoro-4-nitrobenzene (1.1 eq) and KHMDS (1.1 eq) and was obtained as a yellow foam (109 mg, 80%).

Compound **32** was synthesized according to **Procedure D** (750 μmol scale) using 1-fluoro-4-nitrobenzene and was obtained as a yellow foam (294 mg, 86%).

**$^1$H NMR** (600 MHz, $CDCl_3$, **31**) δ 8.21 (d, *J* = 9.1 Hz, 2H), 7.25 (app. t, *J* = 8.5 Hz, 1H), 6.94 (d, *J* = 9.3 Hz, 2H), 6.86 (d, *J* = 7.9 Hz, 1H), 6.82 (app. t, *J* = 2.1 Hz, 1H), 6.78 (dd, *J* = 8.4, 2.5 Hz, 1H), 4.75 (t, *J* = 5.9 Hz, 1H), 3.81 (s, 3H), 3.81 – 3.68 (m, 1H), 3.56 – 3.24 (m, 3H), 3.01 – 2.90 (m, 3H), 2.85 (dt, *J* = 13.9, 6.8 Hz, 1H), 2.10 (ddd, *J* = 13.8, 10.5, 5.7 Hz, 1H), 1.48 (s, 9H) ppm.

**$^1$H NMR** (500 MHz, $CDCl_3$, **32**) 8.21 (d, *J* = 9.2 Hz, 2H), 7.25 (app. t, *J* = 7.9 Hz, 1H), 6.94 (d, *J* = 9.3 Hz, 2H), 6.86 (dd, *J* = 7.8, 1.2 Hz, 1H), 6.81 (t, *J* = 2.1 Hz, 1H), 6.78 (ddd, *J* = 8.2, 2.6, 1.0 Hz, 1H), 4.74 (td, *J* = 6.0, 2.5 Hz, 1H), 3.80 (s, 3H), 3.75 (dd, *J* = 11.4, 9.2 Hz, 1H), 3.45 (d, *J* = 11.1 Hz, 1H), 3.37 (ddd, *J* = 11.6, 6.6, 5.1 Hz, 2H), 3.05 – 2.88 (m, 3H), 2.85 (dt, *J* = 13.9, 6.9 Hz, 1H), 2.10 (ddd, *J* = 13.9, 10.0, 5.8 Hz, 1H), 1.48 (s, 9H) ppm.

**$^{13}$C NMR** (151 MHz, $CDCl_3$, **31**) δ 162.9, 160.1, 154.9, 144.4, 141.7, 129.9, 126.2 (2C), 119.9, 115.4 (2C), 113.6, 111.9, 82.9, 79.9, 55.4 (3C), 50.7, 50.2, 49.3, 49.2, 41.9, 28.7 (3C) ppm (one aliphatic C not detected).

**HRMS** (**31**) m/z calc. for $C_{25}H_{30}N_2NaO_6^+$ $[M+Na]^+$ 477.2002 found 477.1991.

**HRMS** (**32**) m/z calc. for $C_{20}H_{23}N_2O_4^+$ $[M\text{-Boc}+2H]^+$ 355.1658 found 355.1671.

Synthesis of **33** and **34**

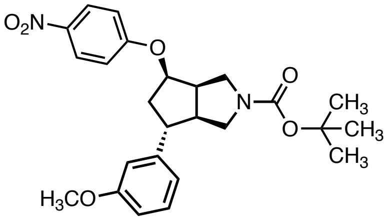

**33**

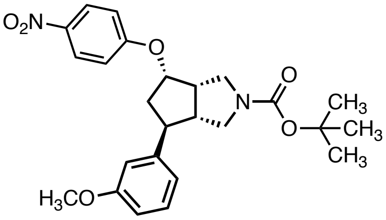

**34**

Compound **33** was synthesized according to **Procedure D** (1.2 mmol scale) using 1-fluoro-4-nitrobenzene and was obtained as a yellow foam (179 mg, 33%) along with unreacted *epi*-**4** (176 mg, 44%).

Compound **34** was synthesized according to **Procedure D** (500 μmol scale) using 1-fluoro-4-nitrobenzene and was obtained as a yellow foam (139 mg, 60%).

**$^{1}$H NMR** (600 MHz, $CDCl_3$, **33**) δ 8.19 (d, *J* = 9.2 Hz, 2H), 7.25 (app., *J* = 7.8 Hz, 1H), 6.96 (d, *J* = 9.3 Hz, 2H), 6.85 – 6.81 (m, 1H), 6.79 – 6.72 (m, 2H), 5.03 (td, *J* = 6.3, 2.7 Hz, 1H), 3.81 (s, 3H), 3.74 (dd, *J* = 11.7, 5.9 Hz, 1H), 3.55 – 3.35 (m, 3H), 3.22 (tt, *J* = 9.0, 6.1 Hz, 1H), 3.15 (dt, *J* = 10.7, 8.1 Hz, 1H), 2.92 – 2.78 (m, 1H), 2.47 (ddd, *J* = 14.1, 7.6, 2.7 Hz, 1H), 2.25 (ddd, *J* = 14.1, 10.8, 6.2 Hz, 1H), 1.46 (s, 9H) ppm.

**$^{1}$H NMR** (600 MHz, $CDCl_3$, **34**) δ 8.20 (d, *J* = 9.3 Hz, 2H), 7.26 (app. t, *J* = 7.8 Hz, 1H), 6.98 (d, *J* = 9.3 Hz, 2H), 6.84 (d, *J* = 7.5 Hz, 1H), 6.79 (d, *J* = 7.7 Hz, 2H), 5.05 (td, *J* = 6.3, 2.6 Hz, 1H), 3.82 (s, 3H), 3.75 (dt, *J* = 11.8, 5.6 Hz, 1H), 3.58 – 3.38 (m, 3H), 3.23 (ddd, *J* = 15.3, 9.0, 6.2 Hz, 1H), 3.16 (dt, *J* = 10.7, 8.1 Hz, 1H), 2.84 (qd, *J* = 8.5, 3.2 Hz, 1H), 2.48 (ddd, *J* = 14.2, 7.6, 2.6 Hz, 1H), 2.27 (ddd, *J* = 14.1, 10.7, 6.2 Hz, 1H), 1.48 (s, 9H) ppm.

**$^{13}$C NMR** (151 MHz, $CDCl_3$, **33**) δ 163.1, 160.0, 154.6, 144.8, 141.7, 129.9, 126.1 (2C), 119.6, 115.3 (2C), 113.5, 111.7, 79.7, 79.6, 55.4 (3C), 51.1, 50.9, 48.4, 47.4, 45.3, 41.8, 28.7 (3C) ppm.

**HRMS** (**33**) m/z calc. for $C_{20}H_{23}N_2O_4^+$ $[M\text{-}Boc+2H]^+$ 355.1658 found 355.1660.

**HRMS** (**34**) m/z calc. for $C_{20}H_{23}N_2O_4^+$ $[M\text{-}Boc+2H]^+$ 355.1658 found 355.1666.

Synthesis of **35** and **36**

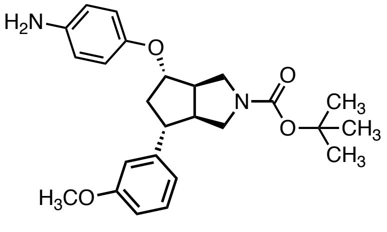

**35**

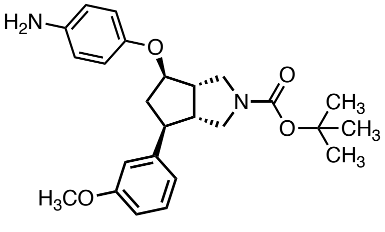

**36**

Compound **35** was synthesized according to **Procedure E** (262 µmol scale; 3 h) and was obtained as a pale-brown foam (98 mg, 88%).

Compound **36** was synthesized according to **Procedure E** (225 µmol scale; 3 h) and was obtained as a pale-brown foam (71 mg, 77%).

**$^1$H NMR** (600 MHz, $CDCl_3$, **35**) δ 7.23 (s, 1H), 6.87 (d, *J* = 7.6 Hz, 1H), 6.84 (s, 1H), 6.77 (d, *J* = 8.1 Hz, 1H), 6.73 (d, *J* = 8.1 Hz, 2H), 6.64 (d, *J* = 8.2 Hz, 2H), 4.53 (s, 1H), 3.80 (s, 3H), 3.77 – 3.20 (m, 6H), 3.02 – 2.82 (m, 3H), 2.71 (dt, *J* = 13.4, 6.5 Hz, 1H), 2.12 – 2.06 (m, 1H), 1.47 (s, 9H) ppm.

**$^1$H NMR** (600 MHz, $CDCl_3$, **36**) δ 7.23 (s, 1H), 6.87 (d, *J* = 7.6 Hz, 1H), 6.84 (s, 1H), 6.77 (d, *J* = 8.2 Hz, 1H), 6.73 (d, *J* = 8.3 Hz, 2H), 6.65 (d, *J* = 8.2 Hz, 2H), 4.53 (s, 1H), 3.80 (s, 3H), 3.76 – 3.21 (m, 6H), 2.98 – 2.82 (m, 3H), 2.72 (dt, *J* = 13.4, 6.6 Hz, 1H), 2.07 (ddt, *J* = 17.1, 9.5, 4.7 Hz, 1H), 1.47 (s, 9H) ppm.

**$^{13}$C NMR** (151 MHz, $CDCl_3$, **35**) δ 159.9, 154.9, 150.9, 145.1, 140.3, 129.7, 120.0, 117.1 (2C), 116.6 (2C), 113.5*, 113.2*, 112.0*, 111.8*, 83.0, 79.6, 55.3 (3C), 50.7, 50.3, 50.02, 49.97, 49.7, 49.1, 49.0, 42.3, 41.9, 28.6 (3C) ppm (in total for 2 additional rotameric peaks detected).

* rotamers; corresponding to 2C

**HRMS** (**35**) m/z calc. for $C_{25}H_{32}N_2NaO_4^+$ $[M+Na]^+$ 447.2260 found 447.2274.

**HRMS** (**36**) m/z calc. for $C_{25}H_{32}N_2NaO_4^+$ $[M+Na]^+$ 447.2260 found 447.2272.

Synthesis of **37** and **38**

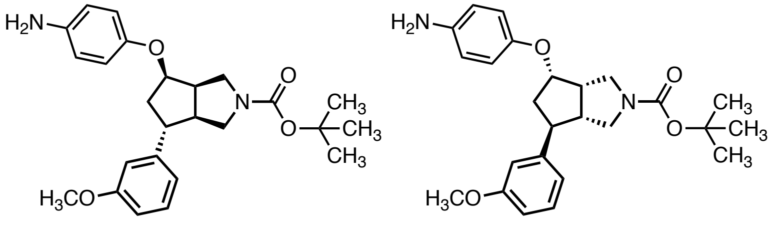

**37** **38**

Compound **37** was synthesized according to **General Procedure E** (178 μmol scale; 3 h) and was obtained as a pale-brown foam (72 mg, 88%).

Compound **38** was synthesized according to **General Procedure E** (280 μmol scale; 3 h) and was obtained as a pale-brown foam (90 mg, 75%).

**$^1$H NMR** (600 MHz, $CDCl_3$, **37**) δ 7.25 – 7.18 (m, 1H), 6.82 (d, *J* = 7.6 Hz, 1H), 6.80 – 6.71 (m, 4H), 6.71 – 6.58 (m, 2H), 4.81 (td, *J* = 6.3, 3.0 Hz, 1H), 3.80 (s, 3H), 3.79 – 3.70 (m, 3H), 3.56 – 3.29 (m, 3H), 3.22 – 2.99 (m, 2H), 2.76 (s, 1H), 2.45 (s, 1H), 2.12 (ddd, *J* = 13.9, 10.3, 6.2 Hz, 1H), 1.46 (s, 9H) ppm.

**$^1$H NMR** (600 MHz, $CDCl_3$, **38**) δ 7.25 – 7.16 (m, 1H), 6.82 (d, *J* = 7.7 Hz, 1H), 6.79 – 6.69 (m, 4H), 6.67 – 6.59 (m, 2H), 4.80 (td, *J* = 6.2, 3.0 Hz, 1H), 3.80 (s, 3H), 3.78 – 3.74 (m, 1H), 3.57 – 3.23 (m, 5H), 3.13 (ddt, *J* = 15.7, 8.6, 5.4 Hz, 2H), 2.76 (s, 1H), 2.45 (s, 1H), 2.12 (ddd, *J* = 13.9, 10.3, 6.2 Hz, 1H), 1.46 (s, 9H) ppm.

**$^{13}$C NMR** (151 MHz, $CDCl_3$, **37**) δ 159.8, 154.7*, 154.6*, 151.3, 145.9*, 145.7*, 140.0*, 139.7*, 129.7, 119.6, 116.9*, 116.8*, 116.7*, 113.5*, 113.3*, 111.5*, 111.3*, 79.5$^†$, 79.2, 79.1$^†$, 55.3 (3C), 51.3$^†$, 51.2$^†$, 51.1$^†$, 50.5$^†$, 48.3$^†$, 48.2$^†$, 48.0$^†$, 47.2$^†$, 45.5$^†$, 45.1$^†$, 42.2$^†$, 41.5$^†$, 28.6 (3C) ppm (in total 13 additional rotameric peaks detected).

* rotamers; corresponding to 6C

$^†$ rotamers; corresponding to 7C

**HRMS** (**37**) m/z calc. for $C_{20}H_{25}N_2O_2^+$ $[M\text{-}Boc+2H]^+$ 325.1916 found 325.1927.

**HRMS** (**38**) m/z calc. for $C_{20}H_{25}N_2O_2^+$ $[M\text{-}Boc+2H]^+$ 325.1916 found 325.1919.

Synthesis of **FWG-1A** (**9**) and **FWG-1B** (**10**)

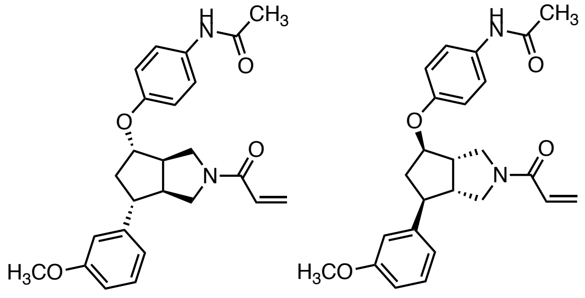


Compound **FWG-1A** (**9**) was synthesized from **35** according to **General Procedure F1** (50 μmol scale), followed by **General Procedure G** and **General Procedure H** in $CH_2Cl_2$ and was obtained as a colorless solid (9.1 mg, 43% over 3 steps; prep. TLC hexane/EtOAc 25:75 and RP-HPLC acetonitrile/$H_2O$) with 99% ee.

Compound **FWG-1B** (**10**) was synthesized from **36** according to **General Procedure F1** (150 μmol scale), followed by **General Procedure G** and **General Procedure H** in $CH_2Cl_2$ and was obtained as a colorless solid (33 mg, 52% over 3 steps; prep. TLC EtOAc/MeOH 98:2 and RP-HPLC acetonitrile/$H_2O$) with 99% ee.

**$^1$H NMR** (600 MHz, $CDCl_3$, **FWG-1A**) δ 7.46 – 7.35 (m, 2H), 7.27 (s, 1H), 7.25 – 7.20 (m, 1H), 6.87 – 6.74 (m, 5H), 6.53 – 6.32 (m, 2H), 5.81 – 5.60 (m, 1H), 4.71 – 4.57 (m, 1H), 4.02 – 3.85 (m, 1H), 3.80 (s, 3H), 3.78 – 3.45 (m, 3H), 3.12 – 2.73 (m, 4H), 2.15 (s, 3H), 2.14 – 2.06 (m, 1H) ppm.

**$^1$H NMR** (600 MHz, $CDCl_3$, **FWG-1B**) δ 7.47 – 7.37 (m, 3H), 7.24 (s, 1H), 6.91 – 6.74 (m, 5H), 6.50 – 6.37 (m, 2H), 5.70 (s, 1H), 4.64 (s, 1H), 4.02 – 3.87 (m, 1H), 3.80 (s, 3H), 3.73 – 3.48 (m, 3H), 3.11 – 2.72 (m, 4H), 2.16 (s, 3H), 2.12 (s, 1H) ppm.

**$^{13}$C NMR** (151 MHz, $CDCl_3$, **FWG-1B**) δ 168.5, 165.1, 160.0, 154.5, 144.5*, 144.3*, 131.6, 129.9, 128.4, 128.3, 122.1, 119.9, 115.9, 113.5*, 113.3*, 112.1, 82.1$^\dagger$, 81.9$^\dagger$, 55.4 (3C), 51.2$^\dagger$, 50.5$^\dagger$, 50.1$^\dagger$, 49.5$^\dagger$, 49.1$^\dagger$, 48.3$^\dagger$, 42.3$^\dagger$, 41.8$^\dagger$, 24.5 (3C) ppm (in total 5 additional rotameric peaks detected).

* rotamers; corresponding to 7xC

$^\dagger$ rotamers; corresponding to 2xC

**HRMS** (**FWG-1A**) m/z calc. for $C_{25}H_{29}N_2O_4^+$ $[M+H]^+$ 421.2127 and 421.2121.

**HRMS** (**FWG-1B**) m/z calc. for $C_{25}H_{29}N_2O_4^+$ $[M+H]^+$ 421.2127 and 421.2128.

**SFC** (Waters UPC2 SFC with a Daicel IA column (3 µm, 4.6x250 mm) under isocratic conditions (3.3 mL/min, 45% MeOH / $CO_2$, 1600 psi backpressure) at 30 °C) ($t_R$ (**FWG-1A**) = 3.02 min; $t_R$ (**FWG-1B**) = 3.64 min).

Synthesis of **FWG-2A** (**11**) and **FWG-2B** (**12**)

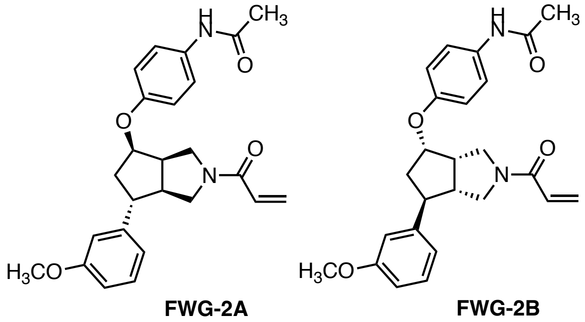

Compound **FWG-2A** (**11**) was synthesized from **37** according to **General Procedure F1** (50 µmol scale), followed by **General Procedure G** and **General Procedure H** in EtOAc and was obtained as a colorless solid (11 mg, 54% over 3 steps; prep. TLC hexane/EtOAc 25:75 and RP-HPLC acetonitrile/$H_2O$) with >99% ee.

Compound **FWG-2B** (**12**) was synthesized from **38** according to **General Procedure F1** (100 µmol scale), followed by **General Procedure G** and **General Procedure H** in EtOAc and was obtained as a colorless solid (18 mg, 56% over 3 steps; prep. TLC EtOAc and RP-HPLC acetonitrile/$H_2O$) with >99% ee.

**$^1$H NMR** (600 MHz, $CDCl_3$, **FWG-2A**) δ 7.52 – 7.36 (m, 3H), 7.25 – 7.21 (m, 1H), 6.89 – 6.72 (m, 5H), 6.51 – 6.41 (m, 1H), 6.40 – 6.33 (m, 1H), 5.67 (dt, *J* = 10.0, 2.3 Hz, 1H), 4.92 (dq, *J* = 6.2, 2.9 Hz, 1H), 4.04 – 3.96 (m, 1H), 3.85 – 3.73 (m, 4H), 3.71 – 3.54 (m, 2H), 3.30 – 3.16 (m, 1H), 3.11 (td, *J* = 11.3, 5.6 Hz, 1H), 2.93 – 2.79 (m, 1H), 2.51 – 2.41 (m, 1H), 2.23 – 2.10 (m, 4H) ppm.

**$^1$H NMR** (600 MHz, $CDCl_3$, **FWG-2B**) δ 8.06 – 7.83 (m, 1H), 7.41 (dd, *J* = 8.8, 7.0 Hz, 2H), 7.25 – 7.19 (m, 1H), 6.83 – 6.73 (m, 5H), 6.53 – 6.41 (m, 1H), 6.39 – 6.32 (m, 1H), 5.67 (dd, *J* = 10.2, 2.1 Hz, 1H), 4.97 – 4.87 (m, 1H), 4.00 (td, *J* = 12.6, 5.7 Hz, 1H), 3.87 – 3.71 (m, 4H), 3.69 – 3.51 (m, 2H), 3.28 – 3.15 (m, 1H), 3.14 – 3.06 (m, 1H), 2.93 – 2.77 (m, 1H), 2.53 – 2.38 (m, 1H), 2.21 – 2.10 (m, 4H) ppm.

**$^{13}$C NMR** (151 MHz, $CDCl_3$, **FWG-2B**) δ 168.6, 164.7*, 164.7*, 160.0*, 159.9*, 154.4*, 154.2*, 144.93*, 144.89*, 131.9, 129.9*, 129.8*, 128.8*, 128.7*, 127.8*, 127.7*, 122.0*, 121.9*, 119.61*, 119.59*, 115.8*, 115.7*, 113.5*, 113.3*, 111.8*, 111.7*, 78.80†, 78.77†, 55.35†, 55.33†, 51.7†, 51.6†, 50.9†, 49.9†, 48.7†, 48.3†, 48.2†, 46.5†, 46.1†, 45.4†, 42.2†, 41.6†, 24.4 ppm (in total 20 additional rotameric peaks detected).

* rotamers; corresponding to 12xC

† rotamers; corresponding to 8xC

**HRMS** (**FWG-2A**) m/z calc. for $C_{25}H_{29}N_2O_4^+$ $[M+H]^+$ 421.2127 and 421.2121.

**HRMS** (**FWG-2B**) m/z calc. for $C_{25}H_{29}N_2O_4^+$ $[M+H]^+$ 421.2127 and 421.2133.

**SFC** (Waters UPC2 SFC with a Daicel IA column (3 µm, 4.6x250 mm) under isocratic conditions (3.3 mL/min, 35% MeOH / $CO_2$, 1600 psi backpressure) at 30 °C) ($t_R$ (**FWG-2A**) = 7.05 min; $t_R$ (**FWG-2B**) = 8.41 min).

Synthesis of **FWG-3A** (**13**) and **FWG-3B** (**14**)

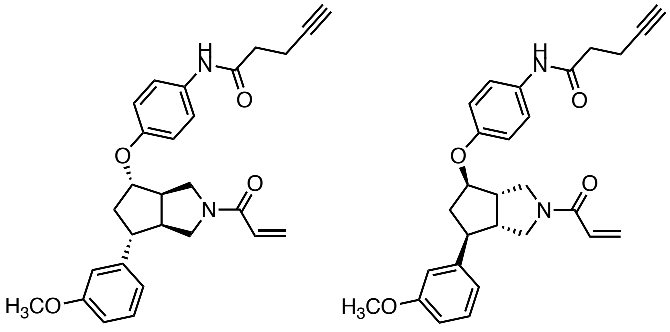


Compound **FWG-3A** (**13**) was synthesized from **35** according to **General Procedure F2** (50 μmol scale), followed by **General Procedure G** and **General Procedure H** in $CH_2Cl$ and was obtained as a colorless solid (9.0 mg, 39% over 3 steps; prep. TLC hexane/EtOAc 25:75 and RP-HPLC acetonitrile/$H_2O$) with >99% ee.

Compound **FWG-3B** (**14**) was synthesized from **36** according to **General Procedure F2** (100 μmol scale), followed by **General Procedure G** and **General Procedure H** in EtOAc and was obtained as a colorless solid (26 mg, 56% over 3 steps; prep. TLC EtOAc and RP-HPLC acetonitrile/$H_2O$) with 99% ee.

**$^1$H NMR** (600 MHz, $CDCl_3$, **FWG-3A**) δ 7.50 – 7.38 (m, 3H), 7.25 – 7.19 (m, 1H), 6.90 – 6.74 (m, 5H), 6.51 – 6.32 (m, 2H), 5.74 – 5.67 (m, 1H), 4.69 – 4.57 (m, 1H), 4.01 – 3.86 (m, 1H), 3.83 – 3.78 (m, 3H), 3.78 – 3.44 (m, 3H), 3.13 – 2.98 (m, 1H), 2.96 – 2.72 (m, 3H), 2.66 – 2.53 (m, 4H), 2.18 – 2.08 (m, 1H), 2.05 (t, *J* = 2.5 Hz, 1H) ppm.

**$^1$H NMR** (600 MHz, $CDCl_3$, **FWG-3B**) δ 7.88 – 7.80 (m, 1H), 7.44 (app. t, *J* = 8.5 Hz, 2H), 7.25 – 7.18 (m, 1H), 6.89 – 6.73 (m, 5H), 6.51 – 6.35 (m, 2H), 5.75 – 5.66 (m, 1H), 4.68 – 4.56 (m, 1H), 4.02 – 3.85 (m, 1H), 3.81 – 3.77 (m, 3H), 3.76 – 3.45 (m, 3H), 3.10 – 2.96 (m, 1H), 2.95 – 2.72 (m, 3H), 2.62 – 2.53 (m, 4H), 2.16 – 2.06 (m, 1H), 2.03 (t, *J* = 2.5 Hz, 1H) ppm.

**$^{13}$C NMR** (151 MHz, $CDCl_3$, **FWG-3B**) δ 169.3, 165.1*, 165.02*, 159.96*, 160.0*, 154.43*, 154.35*, 144.5*, 144.2*, 131.6*, 131.5*, 129.9*, 129.8*, 128.5*, 128.34*, 128.27*, 128.25*, 122.07*, 122.01*, 119.86*, 119.84*, 116.0*, 115.8*, 113.5*, 113.2*, 112.09*, 112.01*, 83.1$^†$, 83.0$^†$, 82.2$^†$, 81.9$^†$, 69.67$^†$, 69.65$^†$, 55.34$^†$, 55.32$^†$, 51.2$^†$, 50.5$^†$, 50.4$^†$, 50.3$^†$, 50.1$^†$, 49.4$^†$, 49.2$^†$, 49.1$^†$, 48.3$^†$, 42.2$^†$, 41.7$^†$, 36. 1$^†$, 15.0 ppm (in total 21 additional rotameric peaks detected).

* rotamers; corresponding to 13C

† rotamers; corresponding to 11C

**HRMS** (**FWG-3A**) m/z calc. for $C_{28}H_{31}N_2O_4^+$ [M+H]$^+$ 459.2284 found 459.2296.

**HRMS** (**FWG-3B**) m/z calc. for $C_{28}H_{31}N_2O_4^+$ [M+H]$^+$ 459.2284 found 459.2283.

**SFC** (Waters UPC2 SFC with a Daicel IA column (3 µm, 4.6x250 mm) under isocratic conditions (3.3 mL/min, 50% MeOH / $CO_2$, 1600 psi backpressure) at 30 °C) ($t_R$ (**FWG-3A**) = 3.18 min; $t_R$ (**FWG-3B**) = 4.30 min).

Synthesis of **FWG-4A** (**15**) and **FWG-4B** (**16**)

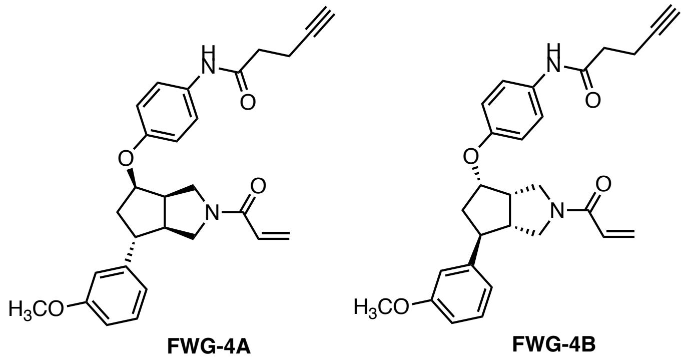

Compound **FWG-4A** (**15**) was synthesized from **37** according to **General Procedure F2** (50 µmol scale), followed by **General Procedure G** and **General Procedure H** in EtOAc and was obtained as a colorless solid (15 mg, 64% over 3 steps; prep. TLC hexane/EtOAc 25:75 and RP-HPLC acetonitrile/$H_2O$) with >99% ee.

Compound **FWG-4B** (**16**) was synthesized from **34** according to **General Procedure E** (100 µmol scale; 17 h), followed by **General Procedure F2** (100 µmol scale), **General Procedure G** and **General Procedure H** in EtOAc and was obtained as a colorless solid (17 mg, 30% over 4 steps; prep. TLC hexane/EtOAc 25:75 and RP-HPLC acetonitrile/$H_2O$) with >99% ee.

**$^1$H NMR** (600 MHz, $CDCl_3$, **FWG-4A**) δ 8.03 – 7.83 (m, 1H), 7.49 – 7.36 (m, 2H), 7.25 – 7.14 (m, 1H), 6.90 – 6.72 (m, 5H), 6.51 – 6.40 (m, 1H), 6.40 – 6.33 (m, 1H), 5.67 (dt, *J* = 10.4, 1.6 Hz, 1H), 4.91 (dq, *J* = 6.2, 3.0 Hz, 1H), 4.07 – 3.94 (m, 1H), 3.85 – 3.73 (m, 4H), 3.70 – 3.53 (m, 2H), 3.32 – 3.16 (m, 1H), 3.15 – 3.06 (m, 1H), 2.95 – 2.77 (m, 1H), 2.65 – 2.54 (m, 4H), 2.51 – 2.40 (m, 1H), 2.25 – 2.09 (m, 1H), 2.02 (dd, *J* = 3.9, 2.4 Hz, 1H) ppm.

**$^1$H NMR** (600 MHz, $CDCl_3$, **FWG-4B**) δ 8.07 – 7.80 (m, 1H), 7.61 – 7.36 (m, 2H), 7.25 – 7.17 (m, 1H), 6.92 – 6.68 (m, 5H), 6.59 – 6.41 (m, 1H), 6.40 – 6.29 (m, 1H), 5.67 (dt, *J* = 10.6, 1.4 Hz, 1H), 5.10 – 4.67 (m, 1H), 4.01 (td, *J* = 11.8, 5.7 Hz, 1H), 3.88 – 3.73 (m, 4H), 3.71 – 3.53 (m, 2H), 3.30 – 3.16 (m, 1H), 3.11 (ddt, *J* = 15.3, 10.6, 7.9 Hz, 1H), 2.94 – 2.77 (m, 1H), 2.65 – 2.54 (m, 4H), 2.50 – 2.40 (m, 1H), 2.22 – 2.10 (m, 1H), 2.03 – 2.00 (m, 1H) ppm.

**$^{13}$C NMR** (151 MHz, $CDCl_3$, **FWG-4B**) δ 169.35*, 169.33*, 164.72*, 164.69*, 160.0*, 159.9*, 154.5*, 154.4*, 144.93*, 144.91*, 131.6*, 131.6*, 130.0*, 129.8*, 128.8*, 128.7*, 127.83*, 127.76*, 122.0*, 121.9*, 119.62*, 119.61*, 115.8*, 115.7*, 113.5*, 113.3*, 111.8*, 111.7*, 83.13$^†$, 83.08$^†$, 78.82$^†$, 78.80$^†$, 69.64$^†$, 69.57$^†$, 55.37$^†$, 55.35$^†$, 51.7$^†$, 51.6$^†$, 50.9$^†$, 49.9$^†$, 48.7$^†$, 48.3$^†$,

48.2$^{\dagger}$, 46.5$^{\dagger}$, 46.1$^{\dagger}$, 45.4$^{\dagger}$, 42.2$^{\dagger}$, 41.6$^{\dagger}$, 36.2, 15.0 ppm (in total 24 additional rotameric peaks detected).

* rotamers; corresponding to 14C

$^{\dagger}$ rotamers; corresponding to 10C

**HRMS** (**FWG-4A**) m/z calc. for $C_{28}H_{31}N_2O_4^+$ $[M+H]^+$ 459.2284 found 459.2277.

**HRMS** (**FWG-4B**) m/z calc. for $C_{28}H_{31}N_2O_4^+$ $[M+H]^+$ 459.2284 found 459.2290.

**SFC** (Waters UPC2 SFC with a Daicel IA column (3 μm, 4.6x250 mm) under isocratic conditions (3.3 mL/min, 35% IPA / $CO_2$, 1600 psi backpressure) at 30 °C) ($t_R$ (**FWG-4A**) = 7.78 min; $t_R$ (**FWG-4B**) = 8.56 min).

Synthesis of **FWG-9A** (**39**) and **FWG-9B** (**17**)

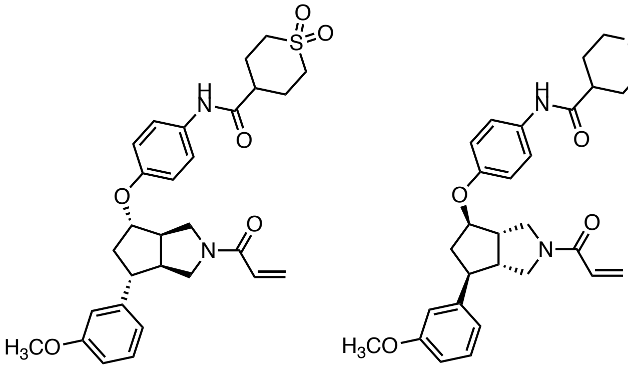


Compound **FWG-9A** (**39**) was synthesized from **35** according to **General Procedure F2** (50 µmol scale), followed by **General Procedure G** and **General Procedure H** in $CH_2Cl_2$ and was obtained as a colorless solid (19 mg, 69% over 3 steps; prep. TLC EtOAc/MeOH 95:5 and RP-HPLC acetonitrile/$H_2O$) with >99% ee.

Compound **FWG-9B** (**17**) was synthesized from **36** according to **General Procedure F2** (50 µmol scale), followed by **General Procedure G** and **General Procedure H** in EtOAc and was obtained as a colorless solid (17 mg, 63% over 3 steps; prep. TLC EtOAc/MeOH 95:5 and RP-HPLC acetonitrile/$H_2O$) with >99% ee

**$^1$H NMR** (600 MHz, $CDCl_3$, **FWG-9A**) δ 7.94 – 7.77 (m, 1H), 7.42 (dd, *J* = 8.8, 5.6 Hz, 2H), 7.25 – 7.18 (m, 1H), 6.94 – 6.71 (m, 5H), 6.54 – 6.33 (m, 2H), 5.79 – 5.66 (m, 1H), 4.62 (dt, *J* = 7.0, 3.5 Hz, 1H), 4.01 – 3.87 (m, 1H), 3.86 – 3.78 (m, 3H), 3.77 – 3.47 (m, 3H), 3.43 – 3.33 (m, 2H), 3.10 – 2.73 (m, 6H), 2.60 (td, *J* = 8.1, 4.1 Hz, 1H), 2.49 – 2.41 (m, 2H), 2.37 (ddt, *J* = 11.2, 8.1, 3.8 Hz, 2H), 2.17 – 2.05 (m, 1H) ppm.

**$^1$H NMR** (600 MHz, $CDCl_3$, **FWG-9B**) δ 8.06 – 7.86 (m, 1H), 7.46 – 7.37 (m, 2H), 7.25 – 7.19 (m, 1H), 6.90 – 6.73 (m, 5H), 6.52 – 6.32 (m, 2H), 5.80 – 5.67 (m, 1H), 4.61 (td, *J* = 6.1, 2.1 Hz, 1H), 3.98 – 3.85 (m, 1H), 3.83 – 3.78 (m, 3H), 3.77 – 3.46 (m, 3H), 3.40 – 3.32 (m, 2H), 3.11 – 2.89 (m, 4H), 2.88 – 2.80 (m, 1H), 2.79 – 2.69 (m, 1H), 2.60 (qt, *J* = 7.4, 3.8 Hz, 1H), 2.51 – 2.41 (m, 2H), 2.40 – 2.30 (m, 2H), 2.20 – 2.06 (m, 1H) ppm.

**$^{13}$C NMR** (151 MHz, $CDCl_3$, **FWG-9B**) δ 171.21*, 171.16*, 165.1*, 165.0*, 160.1*, 160.0*, 154.72*, 154.68*, 144.4*, 144.1*, 131.2*, 130.0*, 129.9*, 128.5*, 128.4*, 128.34*, 128.31*, 122.27*, 122.2*, 119.84*, 119.82*, 116.03*, 115.97*, 113.6*, 113.4*, 112.02*, 111.98*, 82.14†,

82.10$^{\dagger}$, 55.37$^{\dagger}$, 55.36$^{\dagger}$, 51.1$^{\dagger}$, 50.5$^{\dagger}$, 50.4$^{\dagger}$, 50.3$^{\dagger}$, 50.1$^{\dagger}$, 49.83$^{\dagger}$, 49.80$^{\dagger}$, 49.6$^{\dagger}$, 49.2$^{\dagger}$, 49.03$^{\dagger}$, 49.00$^{\dagger}$, 48.2$^{\dagger}$, 42.3$^{\dagger}$, 41.7$^{\dagger}$, 40.9$^{\dagger}$, 27.4 (2C) ppm (in total 22 additional rotameric peaks detected).

* rotamers; corresponding to 14C

$^{\dagger}$ rotamers; corresponding to 11C

**HRMS** (**FWG-9A**) m/z calc. for $C_{29}H_{35}N_2O_6S^+$ $[M+H]^+$ 539.2216 found 539.2228.

**HRMS** (**FWG-9B**) m/z calc. for $C_{29}H_{35}N_2O_6S^+$ $[M+H]^+$ 539.2216 found 539.2225.

**SFC** (Waters UPC2 SFC with a Daicel IK column (3 μm, 4.6x250 mm) under isocratic conditions (3.3 mL/min, 50% MeOH / $CO_2$, 1600 psi backpressure) at 30 °C) ($t_R$ (**FWG-9B**) = 14.92 min; $t_R$ (**FWG-9A**) = 16.52 min).

Synthesis of **FWG-11A** (**40**) and **FWG-11B** (**18**)

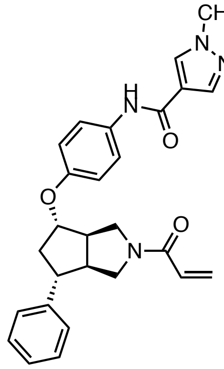

**FWG-11A**

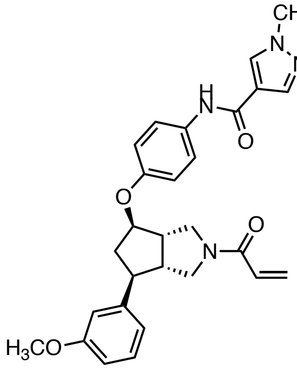

**FWG-11B**

Compound **FWG-11A** (**40**) was synthesized from **35** according to **Procedure F2** (50 µmol scale), followed by **Procedure G** and **Procedure H** in EtOAc and was obtained as a colorless solid (6.0 mg, 25% over 3 steps; prep. TLC EtOAc/MeOH 95:5 and RP-HPLC acetonitrile/$H_2O$) with 99% ee.

Compound **FWG-11B** (**18**) was synthesized from **32** according to **Procedure E** (100 µmol scale, 2h), followed by **Procedure F2**, **Procedure G** and **Procedure H** in EtOAc and was obtained as a colorless solid (24 mg, 49% over 4 steps; prep. TLC EtOAc/MeOH EtOAc/MeOH 95:5 and RP-MPLC acetonitrile/$H_2O$) with >99% ee.

**$^1$H NMR** (600 MHz, $CDCl_3$, **FWG-11A**) δ 7.92 (s, 1H), 7.83 (s, 1H), 7.65 – 7.55 (m, 1H), 7.50 (t, *J* = 8.3 Hz, 2H), 7.25 – 7.19 (m, 1H), 6.92 – 6.74 (m, 5H), 6.53 – 6.34 (m, 2H), 5.71 (ddd, *J* = 15.3, 8.8, 3.5 Hz, 1H), 4.68 – 4.60 (m, 1H), 4.06 – 3.88 (m, 4H), 3.87 – 3.78 (m, 3H), 3.78 – 3.43 (m, 3H), 3.13 – 2.98 (m, 1H), 2.96 – 2.73 (m, 3H), 2.13 (tdd, *J* = 13.8, 10.1, 5.6 Hz, 1H) ppm.

**$^1$H NMR** (600 MHz, $CDCl_3$, **FWG-11B**) δ 8.01 – 7.81 (m, 3H), 7.50 (t, *J* = 9.7 Hz, 2H), 7.25 – 7.17 (m, 1H), 6.91 – 6.72 (m, 5H), 6.52 – 6.33 (m, 2H), 5.76 – 5.67 (m, 1H), 4.66 – 4.58 (m, 1H), 4.11 – 3.87 (m, 4H), 3.84 – 3.79 (m, 3H), 3.77 – 3.40 (m, 3H), 3.12 – 2.95 (m, 1H), 2.95 – 2.70 (m, 3H), 2.11 (qd, *J* = 13.2, 5.7 Hz, 1H) ppm.

**$^{13}$C NMR** (151 MHz, $CDCl_3$, **FWG-11B**) δ 165.1*, 165.0*, 160.8, 160.04*, 159.97*, 154.5*, 154.4*, 144.5*, 144.2*, 138.3*, 138.2*, 132.3, 131.6*, 131.5*, 129.9*, 129.8*, 128.5*, 128.4*, 128.3 (2C), 122.5*, 122.4*, 119.88*, 119.86*, 119.2, 116.0*, 115.9*, 113.5*, 113.2*, 112.1*, 112.0*, 82.2$^{\dagger}$, 81.9$^{\dagger}$, 55.4, 51.2, 50.5$^{\dagger}$, 50.4$^{\dagger}$, 50.3$^{\dagger}$, 50.1$^{\dagger}$, 49.5$^{\dagger}$, 49.2$^{\dagger}$, 49.0$^{\dagger}$, 48.3$^{\dagger}$, 42.3$^{\dagger}$, 41.7$^{\dagger}$, 39.5 ppm
(in total 19 additional rotameric peaks detected).

* rotamers; corresponding to 13C

† rotamers; corresponding to 6C

**HRMS** (**FWG-11A**) m/z calc. for $C_{28}H_{31}N_4O_4^+$ $[M+H]^+$ 487.2345 found 487.2343.

**HRMS** (**FWG-11B**) m/z calc. for $C_{28}H_{31}N_4O_4^+$ $[M+H]^+$ 487.2345 found 487.2361.

**SFC** (Waters UPC2 SFC with a Daicel IK column (3 µm, 4.6x250 mm) under isocratic conditions (3.3 mL/min, 50% MeOH / $CO_2$, 1600 psi backpressure) at 30 °C) ($t_R$ (**FWG-11A**) = 3.35 min; $t_R$ (**FWG-11B**) = 4.53 min).

Synthesis of **41** and **42**

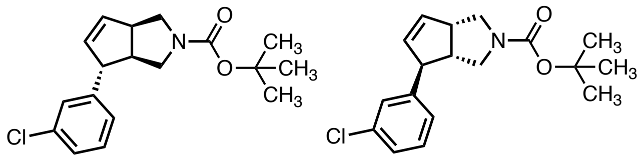

**41** **42**

Compound **41** was synthesized according to **Procedure A** (2.0 mmol scale) using 3-chlorophenylboronic acid) and was obtained as a colorless oil (506 mg, 79%) with 96% ee.

The enantiomer **42** was synthesized according to **Procedure A** (2.0 mmol scale; using (*R*)-Segphos) and 3-chlorophenylboronic acid) and was obtained as a colorless oil (543 mg, 85%) with 98% ee.

**$^{1}$H NMR** (600 MHz, $CDCl_3$, **41**) δ 7.22 (app. t, *J* = 7.7 Hz, 1H), 7.20 – 7.17 (m, 1H), 7.13 (app. t, *J* = 1.9 Hz, 1H), 7.04 (d, *J* = 7.5 Hz, 1H), 5.86 (s, 1H), 5.79 – 5.62 (m, 1H), 3.68 (dd, *J* = 11.3, 8.7 Hz, 2H), 3.63 – 3.19 (m, 4H), 2.74 (s, 1H), 1.46 (s, 9H) ppm.

**$^{1}$H NMR** (600 MHz, $CDCl_3$, **42**) δ 7.22 (app. t, *J* = 7.6 Hz, 1H), 7.18 (app. dt, *J* = 8.0, 1.6 Hz, 1H), 7.13 (app. t, *J* = 1.9 Hz, 1H), 7.04 (app. dt, *J* = 7.3, 1.6 Hz, 1H), 5.85 (s, 1H), 5.74 (dd, *J* = 6.0, 2.4 Hz, 1H), 3.68 (dd, *J* = 11.4, 8.6 Hz, 2H), 3.56 – 3.20 (m, 4H), 2.74 (s, 1H), 1.46 (s, 9H) ppm.

**$^{13}$C NMR** (151 MHz, $CDCl_3$, **42**) δ 154.6, 146.4, 135.2*, 134.8*, 134.6, 133.5*, 133.2*, 130.0, 127.4, 126.8, 125.5, 79.4, 57.5$^{†}$, 52.4$^{†}$, 52.0$^{†}$, 50.9$^{†}$, 50.1$^{†}$, 49.7$^{†}$, 48.7$^{†}$, 28.6 (3C) ppm (in total 4 additional rotameric peaks detected).

* rotamers; corresponding to 2C

$^{†}$ rotamers; corresponding to 5C

**HRMS** (**41**) m/z calc. for $C_{13}H_{15}ClN^+$ [M-Boc+2H]$^+$ 220.0893 found 220.0896.

**HRMS** (**42**) m/z calc. for $C_{13}H_{15}ClN^+$ [M-Boc+2H]$^+$ 220.0893 found 220.0895.

**SFC** (Waters UPC2 SFC with a Daicel IG column (3 µm, 4.6x250 mm) under isocratic conditions (3.3 mL/min, 25% MeOH / $CO_2$, 1600 psi backpressure) at 30 °C) ($t_R$ (**41**) = 1.88 min; $t_R$ (**42**) = 2.60 min).

Synthesis of **43** and **44**

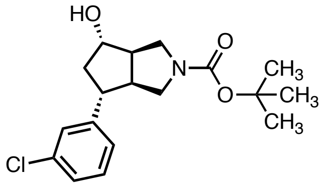

**43**

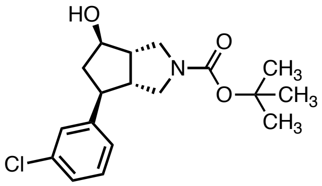

**44**

Compound **43** was synthesized according to **Procedure B** (1.55 mmol scale) and was obtained as a highly viscous colorless oil (164 mg, 31%).

The enantiomer **44** was synthesized according to **Procedure B** (1.65 mmol scale) and was obtained as a colorless oil (284 mg, 51%).

**$^1$H NMR** (600 MHz, $CDCl_3$, **43**) δ 7.29– 7.12 (m, 4H; overlapping with $CHCl_3$ peak), 4.19 (s, 1H), 3.63 – 3.16 (m, 4H), 2.88 – 2.75 (m, 2H), 2.68 (s, 1H), 2.53 (dt, *J* = 13.0, 5.9 Hz, 1H), 1.93 – 1.80 (m, 1H), 1.47 (s, 9H) ppm (OH not detected).

**$^1$H NMR** (500 MHz, $CDCl_3$, **44**) δ 7.30 – 7.10 (m, 4H; overlapping with $CHCl_3$ peak), 4.16 (s, 1H), 3.65 – 3.13 (m, 4H), 2.87 – 2.74 (m, 2H), 2.68 (t, *J* = 8.2 Hz, 1H), 2.51 (dt, *J* = 12.8, 6.1 Hz, 1H), 1.86 (td, *J* = 11.8, 7.2 Hz, 1H), 1.46 (s, 9H) ppm (OH not detected).

**$^{13}$C NMR** (126 MHz, $CDCl_3$, **44**) 155.0, 146.1, 134.5, 130.0, 127.6, 126.7, 125.7, 79.7, 77.6, 52.3$^\dagger$, 51.6$^\dagger$, 51.0$^\dagger$, 50.8$^\dagger$, 50.5$^\dagger$, 49.7$^\dagger$, 48.9, 44.5, 28.6 (3C) ppm (in total 2 additional rotameric peaks detected).

$^\dagger$ rotamers; corresponding to 4C

**HRMS** (**43**) m/z calc. for $C_{13}H_{17}ClNO^+$ [M-Boc+2H]$^+$ 238.0999 found 238.1005.

**HRMS** (**44**) m/z calc. for $C_{13}H_{17}ClNO^+$ [M-Boc+2H]$^+$ 238.0999 found 238.0996.

Synthesis of **45** and **46**

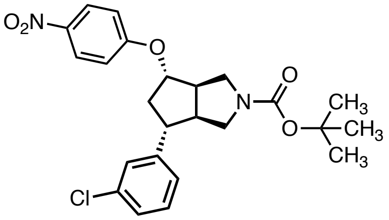

**45**

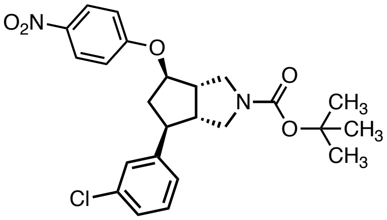

**46**

Compound **45** was synthesized according to **Procedure D** (200 μmol scale) using 1-fluoro-4-nitrobenzene and was obtained as a yellow foam (81 mg, 88%).

Compound **46** was synthesized according to **Procedure D** (440 μmol scale) using 1-fluoro-4-nitrobenzene and was obtained as a yellow foam (152 mg, 75%).

**$^1$H NMR** (600 MHz, $CDCl_3$, **45**) δ 8.20 (d, *J* = 9.1 Hz, 2H), 7.31 – 7.18 (m, 3H), 7.14 (d, *J* = 7.5 Hz, 1H), 6.94 (d, *J* = 8.7 Hz, 2H), 4.76 (s, 1H), 3.76 (s, 1H), 3.50 – 3.28 (m, 3H), 3.11 – 2.75 (m, 4H), 2.08 (ddd, *J* = 15.0, 10.2, 5.3 Hz, 1H), 1.48 (s, 9H) ppm.

**$^1$H NMR** (500 MHz, $CDCl_3$, **46**) δ 8.20 (d, *J* = 9.2 Hz, 2H), 7.30 – 7.18 (m, 3H), 7.14 (dt, *J* = 7.5, 1.5 Hz, 1H), 6.94 (d, *J* = 9.2 Hz, 2H), 4.76 (td, *J* = 5.8, 2.4 Hz, 1H), 3.76 (t, *J* = 10.4 Hz, 1H), 3.49 – 3.30 (m, 3H), 3.06 – 2.76 (m, 4H), 2.08 (ddd, *J* = 13.8, 10.2, 5.4 Hz, 1H), 1.48 (s, 9H) ppm.

**$^{13}$C NMR** (126 MHz, $CDCl_3$, **46**) δ 162.7, 154.8, 144.9, 141.7, 134.7, 130.2, 127.6, 127.2, 126.2 (2C), 125.8, 115.3 (2C), 82.7, 80.0, 50.4, 50.1, 49.6, 49.2, 48.9, 41.7, 28.6 (3C) ppm.

**HRMS** (**46**) m/z calc. for $C_{19}H_{20}ClN_2O_3^+$ $[M\text{-}Boc+2H]^+$ 359.1162 found 359.1160.

Synthesis of **FWG-33A** (**28**) and **FWG-33B** (**22**)

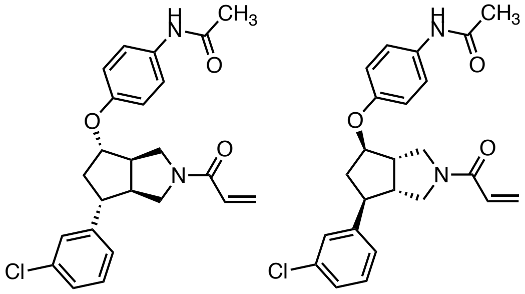

**FWG-33A** **FWG-33B**

Compound **FWG-33A** (**28**) was synthesized from **45** according to **Procedure E** (100 µmol scale; 30 min with 5 wt% Pd/C), followed by **Procedure F1**, **Procedure G** and **Procedure H** in EtOAc and was obtained as a colorless solid (13 mg, 31% over 4 steps; prep. TLC EtOAc and RP-MPLC acetonitrile/$H_2O$) with 97% ee.

Compound **FWG-33B** (**22**) was synthesized from **46** according to **Procedure E** (150 µmol scale; 90 min with 5 wt% Pd/C), followed by **Procedure F1**, **Procedure G** and **Procedure H** in EtOAc and was obtained as a colorless solid (26 mg, 40% over 4 steps; prep. TLC EtOAc and RP-HPLC acetonitrile/$H_2O$) with >99% ee.

**$^1$H NMR** (600 MHz, $CDCl_3$, **FWG-33A**) δ 7.52 – 7.36 (m, 3H), 7.27 (app. t, *J* = 1.9 Hz, 1H), 7.26 – 7.19 (m, 2H), 7.17 – 7.09 (m, 1H), 6.90 – 6.77 (m, 2H), 6.56 – 6.30 (m, 2H), 5.76 – 5.68 (m, 1H), 4.75 – 4.53 (m, 1H), 4.09 – 3.87 (m, 1H), 3.81 – 3.41 (m, 3H), 3.18 – 2.74 (m, 4H), 2.16 (s, 3H), 2.13 – 2.06 (m, 1H) ppm.

**$^1$H NMR** (600 MHz, $CDCl_3$, **FWG-33B**) δ 7.84 – 7.72 (m, 1H), 7.50 – 7.34 (m, 2H), 7.33 – 7.16 (m, 3H; overlapping with $CHCl_3$ peak), 7.15 – 7.07 (m, 1H), 6.90 – 6.73 (m, 2H), 6.56 – 6.27 (m, 2H), 5.78 – 5.62 (m, 1H), 4.73 – 4.54 (m, 1H), 4.11 – 3.82 (m, 1H), 3.79 – 3.38 (m, 3H), 3.20 – 2.59 (m, 4H), 2.14 (s, 3H), 2.12 – 2.04 (m, 1H) ppm.

**$^{13}$C NMR** (151 MHz, $CDCl_3$, **FWG-33B**) δ 168.6, 165.1*, 165.0*, 154.2*, 154.1*, 145.0*, 144.9*, 134.7*, 134.6*, 131.9*, 131.8*, 130.2*, 130.1*, 128.4*, 128.3*, 127.7*, 127.6*, 127.2*, 127.1*, 125.84*, 125.81*, 122.2* (2C), 122.1* (2C), 115.9* (2C), 115.8* (2C), 82.0$^†$, 81.7$^†$, 51.2$^†$, 50.5$^†$, 50.4$^†$, 50.1$^†$, 50.0$^†$, 49.3$^†$, 49.1$^†$, 48.87$^†$, 48.85$^†$, 48.3$^†$, 42.1$^†$, 41.6$^†$, 24.4 ppm (in total 18 additional rotameric peaks detected).

* rotamers; corresponding to 13C

† rotamers; corresponding to 7C

**HRMS** (**FWG-33A**) m/z calc. for $C_{24}H_{26}ClN_2O_3^+$ $[M+H]^+$ 425.1632 found 425.1645.

**HRMS** (**FWG-33B**) m/z calc. for $C_{24}H_{26}ClN_2O_3^+$ $[M+H]^+$ 425.1632 found 425.1635.

**SFC** (Waters UPC2 SFC with a Daicel IA column (3 µm, 4.6x250 mm) under isocratic conditions (3.3 mL/min, 45% MeOH / $CO_2$, 1600 psi backpressure) at 30 °C) ($t_R$ (**FWG-33A**) = 3.15 min; $t_R$ (**FWG-33B**) = 3.90 min).

Synthesis of **FWG-49A** (**29**) and **FWG-49B** (**30**)

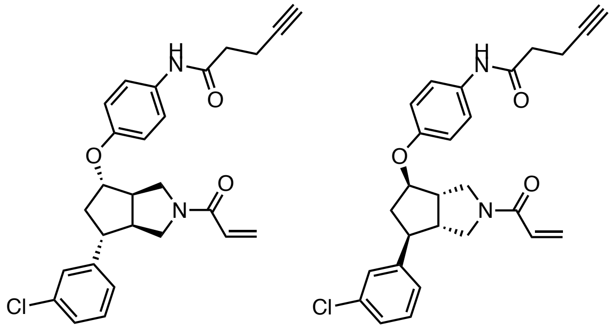

**FWG-49A** **FWG-49B**

Compound **FWG-49A** (**29**) was synthesized from **45** according to **Procedure E** (75 µmol scale; 30 min with 5 wt% Pd/C), followed by **Procedure F2**, **Procedure G** and **Procedure H** in EtOAc and was obtained as a colorless solid (14 mg, 40% over 4 steps; prep. TLC EtOAc and RP-MPLC acetonitrile/$H_2O$) with 98% ee.

Compound **FWG-49B** (**30**) was synthesized from **46** according to **Procedure E** (100 µmol scale; 30 min with 5 wt% Pd/C), followed by **Procedure F2**, **Procedure G** and **Procedure H** in EtOAc and was obtained as a colorless solid (20 mg, 44% over 4 steps; prep. TLC EtOAc and RP-MPLC acetonitrile/$H_2O$) with 98% ee.

**$^1$H NMR** (600 MHz, $CDCl_3$, **FWG-49A**) δ 7.71 – 7.61 (m, 1H), 7.51 – 7.38 (m, 2H), 7.32 – 7.17 (m, 3H; overlapping with $CHCl_3$ peak), 7.17 – 7.06 (m, 1H), 6.87 – 6.79 (m, 2H), 6.51 – 6.33 (m, 2H), 5.81 – 5.62 (m, 1H), 4.70 – 4.59 (m, 1H), 4.09 – 3.87 (m, 1H), 3.82 – 3.42 (m, 3H), 3.16 – 2.73 (m, 4H), 2.66 – 2.51 (m, 4H), 2.13 – 2.06 (m, 1H), 2.04 (s, 1H) ppm.

**$^1$H NMR** (600 MHz, $CDCl_3$, **FWG-49B**) δ 7.63 (s, 1H), 7.54 – 7.39 (m, 2H), 7.27 (app. t, *J* = 1.9 Hz, 1H), 7.25 – 7.18 (m, 2H), 7.16 – 7.06 (m, 1H), 6.95 – 6.70 (m, 2H), 6.53 – 6.36 (m, 2H), 5.83 – 5.63 (m, 1H), 4.71 – 4.56 (m, 1H), 4.03 – 3.86 (m, 1H), 3.80 – 3.40 (m, 3H), 3.18 – 2.75 (m, 4H), 2.70 – 2.50 (m, 4H), 2.14 – 2.06 (m, 1H), 2.04 (t, *J* = 2.5 Hz, 1H) ppm.

**$^{13}$C NMR** (151 MHz, $CDCl_3$, **FWG-49B**) δ 169.3, 165.1*, 165.0*, 154.3, 145.02*, 144.95*, 134.7, 131.6*, 131.5*, 130.2*, 130.1*, 128.4*, 128.3*, 127.7*, 127.6*, 127.2*, 127.1*, 125.9, 122.1, 116.0*, 115.9*, 83.0, 82.1$^\dagger$, 81.8$^\dagger$, 69.8, 51.3$^\dagger$, 50.5$^\dagger$, 50.4$^\dagger$, 50.2$^\dagger$, 50.0$^\dagger$, 49.3$^\dagger$, 49.1$^\dagger$, 48.9, 48.4$^\dagger$, 42.1$^\dagger$, 41.6$^\dagger$, 36.2, 15.0 ppm (in total 14 additional rotameric peaks detected).

* rotamers; corresponding to 8C

$^\dagger$ rotamers; corresponding to 6C

**HRMS** (**FWG-49A**) m/z calc. for $C_{27}H_{28}ClN_2O_3^+$ $[M+H]^+$ 463.1788 found 463.1800.

**HRMS** (**FWG-49B**) m/z calc. for $C_{27}H_{28}ClN_2O_3^+$ $[M+H]^+$ 463.1788 found 463.1787.

**SFC** (Waters UPC2 SFC with a Daicel IBN column (3 μm, 4.6x100 mm) under isocratic conditions (3.3 mL/min, 60% MeOH / $CO_2$, 1600 psi backpressure) at 30 °C) ($t_R$ (**FWG-49B**) = 1.24 min; $t_R$ (**FWG-49A**) = 1.60 min).

Synthesis of **47**

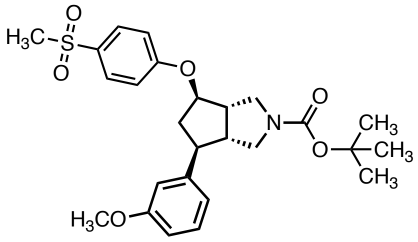

**47**

Compound **47** was synthesized according to **Procedure D** (150 µmol scale; *modification:* aryl fluoride and alcohol were pre-mixed) using 1-fluoro-4-(methylsulfonyl)benzene (44 mg, 60%; purified by MPLC hexane/EtOAc 100:0 to 30:70 and prep. TLC hexane/EtOAc 50:50).

**$^{1}$H NMR** (500 MHz, $CDCl_3$, **47**) δ 7.86 (d, *J* = 8.9 Hz, 2H), 7.24 (app. t, *J* = 8.0 Hz, 1H), 7.00 (d, *J* = 8.9 Hz, 2H), 6.86 (app. dt, *J* = 7.7, 1.2 Hz, 1H), 6.81 (app. t, *J* = 2.1 Hz, 1H), 6.79 – 6.74 (m, 1H), 4.73 (td, *J* = 6.0, 2.4 Hz, 1H), 3.80 (s, 3H), 3.74 (app. t, *J* = 10.3 Hz, 1H), 3.44 (d, *J* = 11.6 Hz, 1H), 3.36 (app. p, *J* = 5.6 Hz, 2H), 3.03 (s, 3H), 3.00 – 2.87 (m, 3H), 2.83 (dt, *J* = 13.9, 7.0 Hz, 1H), 2.08 (ddd, *J* = 13.7, 10.1, 5.7 Hz, 1H), 1.47 (s, 9H) ppm.

**HRMS** (**47**) m/z calc. for $C_{26}H_{33}NNaO_6S^+$ $[M+Na]^+$ 510.1926 found 510.1937.

Synthesis of **FWG-19B** (**17**)

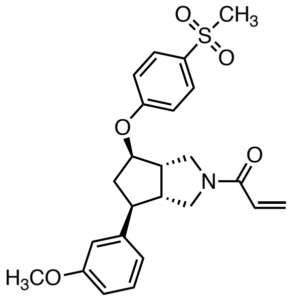

**FWG-19B**

Compound **FWG-19B** (**17**) was synthesized from **47** according to **Procedure G** (85 µmol scale) and **Procedure H** in EtOAc and was obtained as a colorless solid (32 mg, 84% over 2 steps; prep. TLC hexane/EtOAc 33:67).

**$^1$H NMR** (600 MHz, $CDCl_3$, **FWG-19B**) δ 7.87 (d, *J* = 8.9 Hz, 2H), 7.24 (app. t, *J* = 8.1 Hz, 1H), 7.00 (d, *J* = 8.9 Hz, 2H), 6.87 – 6.82 (m, 1H), 6.81 – 6.74 (m, 2H), 6.48 – 6.37 (m, 2H), 5.71 (t, *J* = 6.1 Hz, 1H), 4.77 (s, 1H), 4.09 – 3.95 (m, 1H), 3.79 (s, 3H), 3.71 – 3.46 (m, 3H), 3.22 – 2.79 (m, 7H), 2.30 – 2.08 (m, 1H) ppm.

**HRMS** (**FWG-19B**) m/z calc. for $C_{24}H_{28}NO_5S^+$ $[M+H]^+$ 442.1688 found 442.1684.

Synthesis of **48**

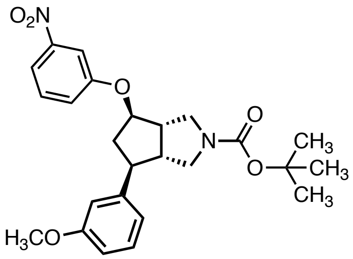

**48**

A mixture of **8** (150 μmol, 1.0 eq), $PPh_3$ (59 mg, 225 μmol, 1.5 eq) and 3-nitrophenol (31 mg, 225 μmol, 1.5 eq) were added 2-dram vial equipped with septum under a $N_2$ atmosphere, and dissolved in dry THF (1.5 mL). The solution was cooled to 0 °C and diisopropyl azodicarboxylate (45 μL, 225 mmol, 1.5 eq) was added dropwise. The mixture was allowed to reach r.t. overnight. A sat. aq. solution of $NaHCO_3$ (2 mL) and $H_2O$ (2 mL) were added. The reaction mixture was extracted with EtOAc (3 x 5 mL). The combined organic layers were washed with brine (5 mL), dried over $MgSO_4$ and concentrated under reduced pressure. Purification by automated medium pressure liquid chromatography (hexane:EtOAc 100:0 to 20:80) afforded **48** as colorless oil (37 mg, 54%).

**$^1$H NMR** (600 MHz, $CDCl_3$, **48**) δ 7.83 (ddd, *J* = 8.2, 2.1, 0.9 Hz, 1H), 7.71 (app. t, *J* = 2.3 Hz, 1H), 7.44 (app. t, *J* = 8.2 Hz, 1H), 7.24 (d, *J* = 7.9 Hz, 1H), 7.22 – 7.20 (m, 1H), 6.87 (d, *J* = 7.7 Hz, 1H), 6.82 (app. t, *J* = 2.1 Hz, 1H), 6.81 – 6.75 (m, 1H), 4.72 (td, *J* = 6.0, 2.7 Hz, 1H), 3.81 (s, 3H), 3.75 (dd, *J* = 11.4, 9.3 Hz, 1H), 3.46 (d, *J* = 11.5 Hz, 1H), 3.38 (td, *J* = 11.0, 8.2 Hz, 2H), 3.05 – 2.89 (m, 3H), 2.85 (ddd, *J* = 13.9, 7.6, 6.2 Hz, 1H), 2.09 (ddd, *J* = 13.8, 10.3, 5.8 Hz, 1H), 1.48 (s, 9H) ppm.

**HRMS** (**48**) m/z calc. for $C_{25}H_{30}N_2NaO_6^+$ $[M+Na]^+$ 477.2002 found 477.2012.

Synthesis of **49**

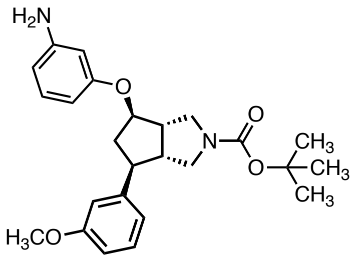

**49**

Compound **49** was synthesized according to **Procedure E** (82 μmol scale; 16 h) and was obtained as a pale-brown foam (23 mg, 65%).

**$^{1}$H NMR** (500 MHz, $CDCl_3$, **49**) δ 7.23 (d, *J* = 7.9 Hz, 1H), 7.06 (app. t, *J* = 8.0 Hz, 1H), 6.88 (d, *J* = 7.7 Hz, 1H), 6.84 (app. t, *J* = 2.1 Hz, 1H), 6.78 (d, *J* = 8.1 Hz, 1H), 6.38 – 6.20 (m, 3H), 4.62 (s, 1H), 4.10 (br. s, 2H), 3.81 (s, 3H), 3.57 – 3.22 (m, 4H), 2.87 (s, 3H), 2.76 (dt, *J* = 13.5, 6.6 Hz, 1H), 2.10 (s, 1H). 1.48 (s, 9H) ppm.

**HRMS** (**49**) m/z calc. for $C_{25}H_{33}N_2O_4^+$ $[M+H]^+$ 425.2440 found 425.2445.

Synthesis of **FWG-21B** (**18**)

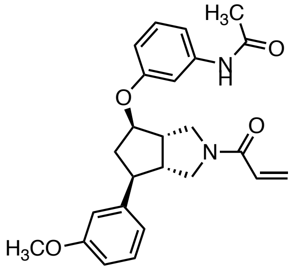

**FWG-21B**

Compound **FWG-21B** (**18**) was synthesized from **49** according to **Procedure F1** (50 µmol scale), followed by **Procedure G** and **Procedure H** in EtOAc and was obtained as a colorless solid (14 mg, 66% over 3 steps; prep. TLC hexane/EtOAc 33:67).

**$^{1}$H NMR** (500 MHz, $CDCl_3$, **FWG-21B**) δ 7.74 – 7.42 (m, 1.4H), 7.26 – 7.22 (m, 1H), 7.19 (app. *t*, J = 8.1 Hz, 1H), 7.02 (s, 0.6H), 6.85 (app. dt, *J* = 7.7, 1.2 Hz, 1H), 6.81 (app. t, *J* = 2.4 Hz, 1H), 6.78 (d, *J* = 8.3 Hz, 1H), 6.62 (d, *J* = 8.2 Hz, 1H), 6.56 – 6.33 (m, 2H), 5.71 (t, *J* = 6.2 Hz, 1H), 4.65 (td, *J* = 6.1, 2.1 Hz, 1H), 4.01 (d, *J* = 10.6 Hz, 1H), 3.80 (s, 3H), 3.56 (s, 3H), 3.15 – 2.62 (m, 5H), 2.18 (s, 3H), 2.13 (ddd, *J* = 12.9, 6.7, 4.1 Hz, 1H) ppm.

**HRMS** (**FWG-21B**) m/z calc. for $C_{25}H_{29}N_2O_4^+$ $[M+H]^+$ 421.2127 found 421.2121.

Synthesis of **50**

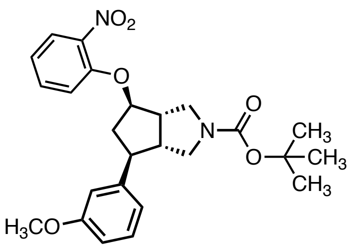

**50**

Compound **50** was synthesized according to **Procedure D** (150 μmol scale) using 1-fluoro-2-nitrobenzene (28 mg, 41%; purified by MPLC hexane/EtOAc 100:0 to 40:60).

**$^{1}$H NMR** (600 MHz, $CDCl_3$, **50**) δ 7.83 (dd, *J* = 8.0, 1.7 Hz, 1H), 7.50 (ddd, *J* = 8.2, 7.5, 1.7 Hz, 1H), 7.24 (app. t, *J* = 7.9 Hz, 1H), 7.10 – 6.99 (m, 2H), 6.88 (app. dt, *J* = 7.7, 1.2 Hz, 1H), 6.83 (app. *t*, J = 2.1 Hz, 1H), 6.78 (ddd, *J* = 8.2, 2.6, 0.9 Hz, 1H), 4.79 (td, *J* = 5.8, 2.5 Hz, 1H), 3.81 (s, 3H), 3.73 (dd, *J* = 11.4, 9.4 Hz, 1H), 3.44 (d, *J* = 11.4 Hz, 1H), 3.34 (ddd, *J* = 11.3, 6.7, 4.4 Hz, 2H), 3.12 – 3.00 (m, 1H), 2.98 – 2.89 (m, 2H), 2.84 (ddd, *J* = 13.7, 7.6, 6.2 Hz, 1H), 2.18 (ddd, *J* = 13.8, 9.9, 5.5 Hz, 1H), 1.47 (s, 9H) ppm.

**HRMS** (**50**) m/z calc. for $C_{25}H_{31}N_2O_6^+$ $[M+H]^+$ 455.2182 found 455.2180.

Synthesis of **51**

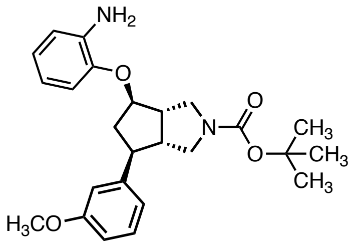

**51**

Compound **51** was synthesized according to **Procedure E** (62 μmol scale; 16 h) and was obtained as a colorless foam (22 mg, 84%).

**$^{1}$H NMR** (600 MHz, $CDCl_3$, **51**) δ 7.23 (app. t, *J* = 7.8 Hz, 1H), 6.88 (d, *J* = 7.7 Hz, 1H), 6.83 (t, *J* = 2.0 Hz, 1H), 6.81 – 6.70 (m, 5H), 4.66 (s, 1H), 4.20 (br. s, 1H), 3.80 (s, 3H), 3.73 – 3.25 (m, 4H), 3.04 – 2.87 (m, 3H), 2.80 (dt, *J* = 13.4, 6.6 Hz, 1H), 2.15 (ddd, *J* = 13.8, 10.7, 6.0 Hz, 1H), 1.48 (s, 9H) (one NH not detected) ppm.

**HRMS** (**51**) m/z calc. for $C_{25}H_{32}N_2NaO_4^+$ $[M+Na]^+$ 447.2260 found 447.2258.

Synthesis of **FWG-23B** (**19**)

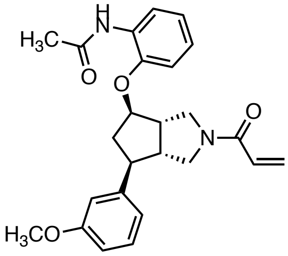

**FWG-23B**

Compound **FWG-23B** (**19**) was synthesized from **51** according to **Procedure F1** (50 µmol scale), followed by **Procedure G** and **Procedure H** in EtOAc and was obtained as a colorless solid (11 mg, 54% over 3 steps; prep. TLC hexane/EtOAc 33:67).

**$^1$H NMR** (500 MHz, $CDCl_3$, **FWG-23B**) δ 8.36 (d, *J* = 8.5 Hz, 1H), 7.65 (s, 1H), 7.29 – 7.22 (m, 1H), 7.02 (td, *J* = 7.8, 1.8 Hz, 1H), 6.97 (td, *J* = 7.8, 1.5 Hz, 1H), 6.86 (dt, *J* = 7.6, 1.2 Hz, 1H), 6.84 – 6.75 (m, 3H), 6.41 (d, *J* = 6.7 Hz, 2H), 5.71 (d, *J* = 8.3 Hz, 1H), 4.74 (s, 1H), 4.09 – 3.89 (m, 1H), 3.80 (s, 3H), 3.64 – 3.50 (m, 3H), 3.14 – 2.92 (m, 3H), 2.85 (dt, *J* = 13.7, 6.8 Hz, 1H), 2.19 (s, 4H) ppm.

**HRMS** (**FWG-23B**) m/z calc. for $C_{25}H_{29}N_2O_4^+$ $[M+H]^+$ 421.2127 found 421.2124.

Synthesis of **52**

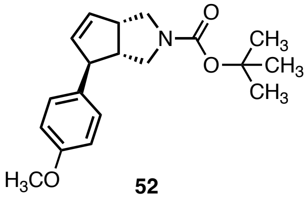

**52**

Compound **52** was synthesized according to **Procedure A** (1.0 mmol scale; using (*R*)-Segphos and 4-methoxyphenylboronic acid) and was obtained as a colorless oil (248 mg, 79%) with 99% ee.

**$^1$H NMR** (600 MHz, $CDCl_3$, **52**) δ 7.07 (d, *J* = 8.6 Hz, 2H), 6.84 (d, *J* = 8.7 Hz, 2H), 5.80 (d, *J* = 5.6 Hz, 1H), 5.77 – 5.73 (m, 1H), 3.79 (s, 3H), 3.73 – 3.60 (m, 2H), 3.55 – 3.18 (m, 4H), 2.72 (td, *J* = 8.5, 4.1 Hz, 1H), 1.46 (s, 9H) ppm.

**HRMS** (**52**) m/z calc. for $C_{14}H_{18}NO^+$ $[M\text{-}Boc+2H]^+$ 216.1388 found 216.1390.

**SFC** (Waters UPC2 SFC with a Daicel IG column (3 µm, 4.6x250 mm) under isocratic conditions (3.3 mL/min, 25% MeOH / $CO_2$, 1600 psi backpressure) at 30 °C) ($t_R$ (**52**) = 2.76 min; $t_R$ (*ent*-**52**) = 3.11 min).

Synthesis of **53**

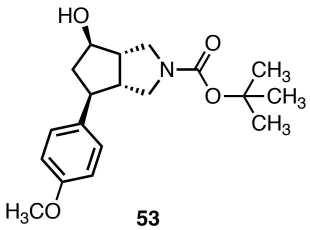

**53**

Compound **53** was synthesized according to **Procedure B** (750 µmol scale; *modification:* the reaction was performed with 5 mol% [$Rh(PPh_3)_3Cl$] at 50 °C) and was obtained as a viscous colorless oil (64 mg, 30%).

**$^1$H NMR** (600 MHz, $CDCl_3$, **53**) δ 7.18 (d, *J* = 8.3 Hz, 2H), 6.95 – 6.77 (m, 2H), 4.16 (s, 1H), 3.79 (s, 3H), 3.64 – 3.23 (m, 4H), 2.91 – 2.57 (m, 3H), 2.49 (dt, *J* = 12.9, 5.9 Hz, 1H), 2.27 (br. s, 1H), 1.94 – 1.75 (m, 1H), 1.46 (s, 9H) ppm.

**HRMS** (**53**) m/z calc. for $C_{14}H_{20}NO_2^+$ $[M\text{-}Boc+2H]^+$ 234.1494 found 234.1496.

Synthesis of **54**

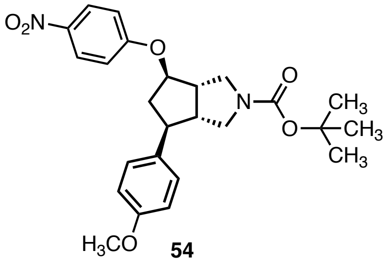


**54**

Compound **54** was synthesized according to **Procedure D** (180 μmol scale) using 1-fluoro-4-nitrobenzene (1.2 eq) and KHMDS (1.5 eq) and was obtained as a yellow foam (60 mg, 73%).

**$^1$H NMR** (500 MHz, $CDCl_3$, **54**) δ 8.20 (d, *J* = 9.0 Hz, 2H), 7.18 (d, *J* = 8.7 Hz, 2H), 6.93 (d, *J* = 9.2 Hz, 2H), 6.86 (d, *J* = 8.1 Hz, 2H), 4.73 (td, *J* = 6.1, 2.5 Hz, 1H), 3.79 (s, 4H), 3.50 – 3.28 (m, 3H), 3.05 – 2.68 (m, 4H), 2.12 – 2.02 (m, 1H), 1.48 (s, 9H) ppm.

**HRMS** (**54**) m/z calc. for $C_{20}H_{23}N_2O_4^+$ $[M\text{-}Boc+2H]^+$ 355.1658 found 355.1666.

Synthesis of **FWG-35B** (**23**)

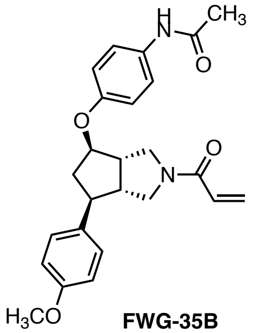

**FWG-35B**

Compound **FWG-35B** (**23**) was synthesized from **54** according to **Procedure E** (125 µmol scale; 20 h), followed by **Procedure F1, Procedure G** and **Procedure H** in EtOAc and was obtained as a colorless solid (20 mg, 37% over 4 steps; prep. TLC EtOAc and RP-MPLC acetonitrile/$H_2O$).

**$^1$H NMR** (500 MHz, $CDCl_3$, **FWG-35B**) δ 7.65 – 7.56 (m, 1H), 7.41 (dd, *J* = 9.0, 6.9 Hz, 2H), 7.23 – 7.14 (m, 2H), 6.91 – 6.78 (m, 4H), 6.53 – 6.34 (m, 2H), 5.70 (ddd, *J* = 14.2, 9.0, 3.2 Hz, 1H), 4.72 – 4.52 (m, 1H), 4.05 – 3.86 (m, 1H), 3.81 – 3.78 (m, 3H), 3.77 – 3.41 (m, 3H), 3.15 – 2.66 (m, 4H), 2.14 (s, 3H), 2.11 – 2.04 (m, 1H) ppm.

**HRMS** (**FWG-35B**) m/z calc. for $C_{25}H_{29}N_2O_4^+$ $[M+H]^+$ 421.2127 found 421.2128.

Synthesis of **FWG-41B** (**24**)

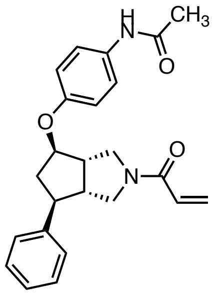

**FWG-41B**

Compound **FWG-41B** (**24**) was synthesized from **46** according to **Procedure E** (100 µmol scale; 18 h with 20 wt% Pd/C), followed by **Procedure F1, Procedure G** and **Procedure H** in EtOAc and was obtained as a colorless solid (8 mg, 21% over 4 steps; prep. TLC EtOAc and RP-MPLC acetonitrile/$H_2O$).

**$^1$H NMR** (500 MHz, $CDCl_3$, **FWG-41B**) δ 7.44 – 7.38 (m, 2H), 7.33 (app. t, *J* = 7.5 Hz, 2H), 7.29 – 7.15 (m, 4H; overlapping with $CHCl_3$), 6.88 – 6.80 (m, 2H), 6.48 – 6.35 (m, 2H), 5.77 – 5.65 (m, 1H), 4.65 (s, 1H), 4.07 – 3.43 (m, 4H), 3.12 – 2.69 (m, 4H), 2.19 – 2.08 (m, 4H) ppm.

**HRMS** (**FWG-41B**) m/z calc. for $C_{24}H_{27}N_2O_3^+$ [M+H]$^+$ 391.2022 found 391.2020.

Synthesis of **55**

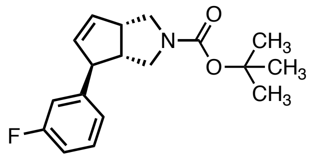


**55**

Compound **55** was synthesized according to **Procedure A** (1.0 mmol scale; using (*R*)-Segphos and 3-fluorophenylboronic acid pinacol ester) and was obtained as a colorless oil (262 mg, 86%) with 98% ee.

**$^{1}$H NMR** (400 MHz, $CDCl_3$, **55**) δ 7.31 – 7.19 (m, 1H), 7.01 – 6.78 (m, 3H), 5.85 (d, *J* = 5.6 Hz, 1H), 5.79 – 5.72 (m, 1H), 3.76 – 3.62 (m, 2H), 3.62 – 3.07 (m, 4H), 2.75 (s, 1H), 1.46 (s, 9H) ppm.

**$^{19}$F NMR** {1H} (376 MHz, $CDCl_3$, **55**) δ -115.82 (s, 1F) ppm.

**HRMS** (**55**) m/z calc. for $C_{13}H_{15}FN^{+}$ [M-Boc+2H]$^{+}$ 204.1189 found 204.1189.

**SFC** (Waters UPC2 SFC with a Daicel IG column (3 µm, 4.6x100 mm) under isocratic conditions (3.3 mL/min, 25% MeOH / $CO_2$, 1600 psi backpressure) at 30 °C) ($t_R$ (*ent*-**55**) = 0.74 min; $t_R$ (**55**) = 0.90 min).

Synthesis of **56**

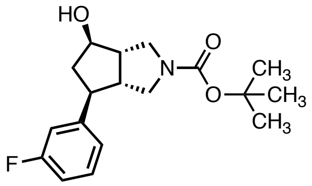

**56**

Compound **56** was synthesized according to **Procedure B** (800 μmol scale) and was obtained as a viscous colorless oil (114 mg, 44%).

**$^{1}$H NMR** (400 MHz, $CDCl_3$, **56**) δ 7.22 (app. t, *J* = 7.2 Hz, 1H), 7.01 (dd, *J* = 7.7, 1.2 Hz, 1H), 6.97 (app. dt, *J* = 10.3, 2.1 Hz, 1H), 6.89 (app. td, *J* = 8.5, 2.6 Hz, 1H), 4.15 (q, *J* = 6.1 Hz, 1H), 3.62 – 3.27 (m, 4H), 3.13 (br. s, 0.6H, OH), 2.85 – 2.63 (m, 3H), 2.50 (dt, *J* = 12.9, 6.3 Hz, 1H), 2.15 (br. s, 0.4H, OH), 1.86 (ddd, *J* = 13.0, 10.6, 7.4 Hz, 1H), 1.45 (s, 9H) ppm.

**$^{19}$F NMR** {1H} (376 MHz, $CDCl_3$, **56**) δ -115.69 (s, 1F) ppm.

**HRMS** (**56**) m/z calc. for $C_{13}H_{17}FNO^+$ [M-Boc+2H]$^+$ 222.1294 found 222.1295.

Synthesis of **57**

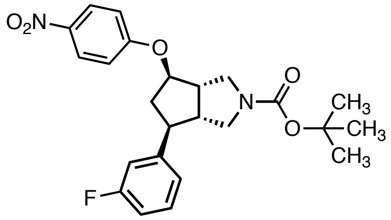

**57**

Compound **57** was synthesized according to **Procedure D** (300 μmol scale) using 1-fluoro-4-nitrobenzene (1.2 eq) and KHMDS (1.5 eq) and was obtained as a pale-yellow foam (96 mg, 71%).

**$^{1}$H NMR** (400 MHz, $CDCl_3$, **57**) δ 8.23 – 8.16 (m, 2H), 7.32 – 7.23 (m, 1H), 7.06 – 7.01 (m, 1H), 7.00 – 6.88 (m, 4H), 4.76 (td, *J* = 5.8, 2.4 Hz, 1H), 3.76 (dd, *J* = 11.4, 9.3 Hz, 1H), 3.50 – 3.29 (m, 3H), 3.07 – 2.77 (m, 4H), 2.07 (ddt, *J* = 18.1, 9.7, 4.8 Hz, 1H), 1.47 (s, 9H).
ppm.

**$^{19}$F NMR** {1H} (376 MHz, $CDCl_3$, **57**) δ -115.30 (s, 1F) ppm.

**HRMS** (**57**) m/z calc. for $C_{19}H_{20}FN_2O_3^+$ $[M\text{-}Boc+2H]^+$ 343.1458 found 343.1460.

Synthesis of **FWG-43B** (**25**)

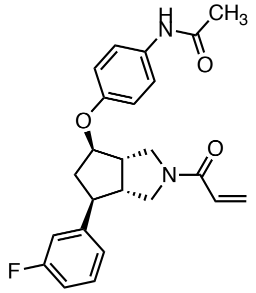

**FWG-43B**

Compound **FWG-43B** (**25**) was synthesized from **57** according to **Procedure E** (100 µmol scale; 1 h), followed by **Procedure F1, Procedure G** and **Procedure H** in EtOAc and was obtained as a colorless solid (24 mg, 58% over 4 steps; prep. TLC EtOAc and RP-MPLC acetonitrile/$H_2O$).

**$^1$H NMR** (400 MHz, $CDCl_3$, **FWG-43B**) δ 7.55 – 7.38 (m, 3H), 7.31 – 7.20 (m, 1H; overlapping with $CHCl_3$), 7.15 – 6.82 (m, 5H), 6.52 – 6.29 (m, 2H), 5.79 – 5.60 (m, 1H), 4.76 – 4.55 (m, 1H), 4.03 – 3.86 (m, 1H), 3.81 – 3.43 (m, 3H), 3.14 – 2.65 (m, 4H), 2.15 (s, 3H), 2.12 – 2.03 (m, 1H) ppm.

**$^{19}$F NMR** {1H} (376 MHz, $CDCl_3$, **FWG-43B**) δ -115.20 (s, 1F, 0.6F), -115.40 (s, 0.4F) ppm.

**HRMS** (**FWG-43B**) m/z calc. for $C_{24}H_{26}FN_2O_3^+$ $[M+H]^+$ 409.1927 found 409.1930

Synthesis of **58**

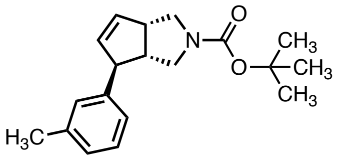

**58**

Compound **58** was synthesized according to **Procedure A** (1.0 mmol scale; using (*R*)-Segphos and 3-methylphenylboronic acid pinacol ester) and was obtained as a colorless oil (253 mg, 85%) with 97% ee.

**$^1$H NMR** (500 MHz, $CDCl_3$, **58**) δ 7.19 (t, *J* = 7.5 Hz, 1H), 7.06 – 7.01 (m, 1H), 6.96 (d, *J* = 8.7 Hz, 2H), 5.88 – 5.81 (m, 1H), 5.79 – 5.74 (m, 1H), 3.68 (dd, *J* = 11.2, 8.7 Hz, 2H), 3.58 – 3.19 (m, 4H), 2.77 (s, 1H), 2.33 (s, 3H), 1.47 (s, 9H) ppm.

**HRMS** (**58**) m/z calc. for $C_{14}H_{18}N^+$ $[M\text{-}Boc+2H]^+$ 200.1439 found 200.1441.

**SFC** (Waters UPC2 SFC with a Daicel IA column (3 µm, 4.6x100 mm) under isocratic conditions (3.3 mL/min, 3% MeOH / $CO_2$, 1600 psi backpressure) at 30 °C) ($t_R$ (*ent*-**58**) = 1.41 min; $t_R$ (**58**) = 1.63 min).

Synthesis of **59**

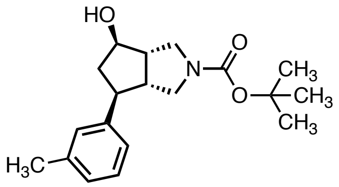

**59**

Compound **59** was synthesized according to **Procedure B** (800 μmol scale) and was obtained as a viscous colorless oil (126 mg, 50%).

**$^1$H NMR** (500 MHz, $CDCl_3$, **59**) δ 7.20 (app. t, *J* = 7.5 Hz, 1H), 7.13 – 6.99 (m, 3H), 4.17 (s, 1H), 3.66 – 3.22 (m, 4H), 2.90 – 2.43 (m, 5H), 2.34 (s, 3H), 1.89 (td, *J* = 11.8, 7.5 Hz, 1H), 1.47 (s, 9H) ppm.

**HRMS** (**59**) m/z calc. for $C_{14}H_{20}NO^+$ $[M\text{-}Boc+2H]^+$ 218.1545 found 218.1544

Synthesis of **60**

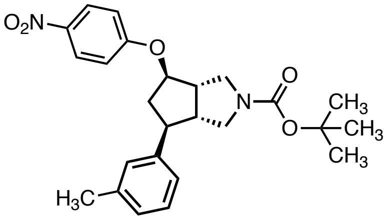

**60**

Compound **60** was synthesized according to **Procedure D** (350 µmol scale) using 1-fluoro-4-nitrobenzene (1.2 eq) and KHMDS (1.5 eq) and was obtained as a pale-yellow foam (103 mg, 67%).

**$^{1}$H NMR** (500 MHz, $CDCl_3$, **60**) δ 8.20 (d, *J* = 9.2 Hz, 2H), 7.22 (app. t, *J* = 7.5 Hz, 1H), 7.12 – 7.02 (m, 3H), 6.94 (d, *J* = 9.2 Hz, 2H), 4.75 (td, *J* = 6.0, 2.5 Hz, 1H), 3.76 (dd, *J* = 11.4, 9.3 Hz, 1H), 3.45 (d, *J* = 11.5 Hz, 1H), 3.38 (ddd, *J* = 15.3, 11.4, 6.5 Hz, 2H), 3.02 – 2.78 (m, 4H), 2.35 (s, 3H), 2.11 (ddd, *J* = 13.6, 10.2, 5.7 Hz, 1H), 1.49 (s, 9H) ppm.

**HRMS** (**60**) m/z calc. for $C_{20}H_{23}N_2O_3^+$ $[M\text{-}Boc+2H]^+$ 339.1709 found 339.1710.

Synthesis of **FWG-45B**

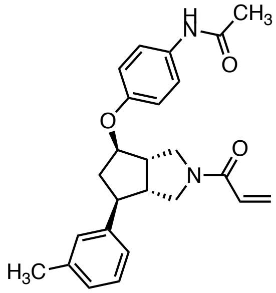

**FWG-45B**

Compound **FWG-45B** (**26**) was synthesized from **60** according to **Procedure E** (100 µmol scale; 2 h), followed by **Procedure F1, Procedure G** and **Procedure H** in EtOAc and was obtained as a colorless solid (11 mg, 28% over 4 steps; prep. TLC EtOAc and RP-MPLC acetonitrile/$H_2O$).

**$^1$H NMR** (500 MHz, $CDCl_3$, **FWG-45B**) δ 7.47 – 7.36 (m, 2H), 7.28 – 7.17 (m, 2H), 7.11 – 7.02 (m, 3H), 6.90 – 6.82 (m, 2H), 6.49 – 6.37 (m, 2H), 5.75 – 5.65 (m, 1H), 4.64 (s, 1H), 3.96 (s, 1H), 3.85 – 3.42 (m, 3H), 3.10 – 2.72 (m, 4H), 2.34 (s, 3H), 2.16 (s, 3H), 2.14 – 2.08 (m, 1H) ppm.

**HRMS** (**FWG-45B**) m/z calc. for $C_{25}H_{29}N_2O_3^+$ $[M+H]^+$ 405.2178 found 405.2178.

Synthesis of **61**

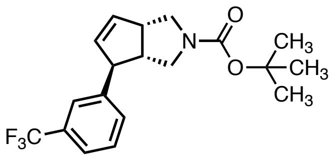


**61**

Compound **61** was synthesized according to **Procedure A** (1.0 mmol scale; using (*R*)-Segphos and 3-trifluoromethylphenylboronic acid pinacol ester) and was obtained as a colorless oil (304 mg, 86%) with 98% ee.

**$^1$H NMR** (400 MHz, $CDCl_3$, **61**) δ 7.49 – 7.46 (m, 1H), 7.44 – 7.38 (m, 2H), 7.37 – 7.32 (m, 1H), 5.89 (d, *J* = 5.5 Hz, 1H), 5.82 – 5.66 (m, 1H), 3.77 (s, 1H), 3.70 (dd, *J* = 11.3, 8.7 Hz, 1H), 3.61 – 3.18 (m, 4H), 2.75 (s, 1H), 1.46 (s, 9H) ppm.

**$^{19}$F NMR** {1H} (376 MHz, $CDCl_3$, **61**) δ -65.20 (s, 3F) ppm.

**HRMS** (**61**) m/z calc. for $C_{14}H_{15}F_3N^+$ [M-Boc+2H]$^+$ 254.1157 found 254.1156.

**SFC** (Waters UPC2 SFC with a Daicel IG column (3 μm, 4.6x100 mm) under isocratic conditions (3.3 mL/min, 5% IPA / $CO_2$, 1600 psi backpressure) at 30 °C) ($t_R$ (*ent*-**61**) = 1.17 min; $t_R$ (**61**) = 1.42 min).

Synthesis of **62**

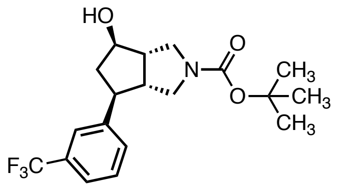

**62**

Compound **62** was synthesized according to **Procedure B** (800 μmol scale) and was obtained as a viscous colorless oil (114 mg, 49%).

**$^{1}$H NMR** (400 MHz, $CDCl_3$, **62**) δ 7.51 (s, 1H), 7.48 – 7.35 (m, 3H), 4.17 (q, *J* = 6.0 Hz, 1H), 3.65 – 3.01 (m, 5H), 2.92 – 2.64 (m, 3H), 2.53 (dt, J = 13.2, 6.5 Hz, 1H), 1.88 (ddd, *J* = 13.1, 10.6, 7.4 Hz, 1H), 1.45 (s, 9H) ppm.

**$^{19}$F NMR** {1H} (376 MHz, $CDCl_3$, **62**) δ - 65.18 (s, 3F) ppm.

**HRMS** (**62**) m/z calc. for $C_{14}H_{17}F_3NO^+$ $[M\text{-}Boc+2H]^+$ 272.1262 found 272.1264.

Synthesis of **63**

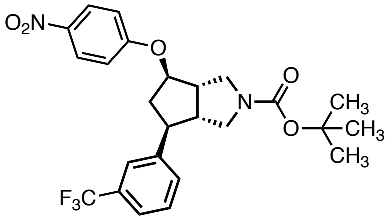

**63**

Compound **63** was synthesized according to **Procedure D** (250 μmol scale) using 1-fluoro-4-nitrobenzene (1.2 eq) and KHMDS (1.5 eq) and was obtained as a pale-yellow foam (101 mg, 82%).

**$^{1}$H NMR** (400 MHz, $CDCl_3$, **63**) δ 8.20 (d, *J* = 8.7 Hz, 2H), 7.56 – 7.42 (m, 4H), 6.94 (d, *J* = 8.9 Hz, 2H), 4.79 (t, *J* = 5.8 Hz, 1H), 3.77 (s, 1H), 3.61 – 3.26 (m, 3H), 3.12 – 2.81 (m, 4H), 2.11 (ddd, *J* = 14.6, 9.9, 5.1 Hz, 1H), 1.48 (s, 9H) ppm.

**$^{19}$F NMR** {1H} (376 MHz, $CDCl_3$, **63**) δ -62.53 (s, 3F) ppm.

**HRMS** (**63**) m/z calc. for $C_{20}H_{20}F_3N_2O_3^+$ $[M\text{-}Boc+2H]^+$ 393.1426 found 393.1429.

Synthesis of **FWG-47B** (**27**)

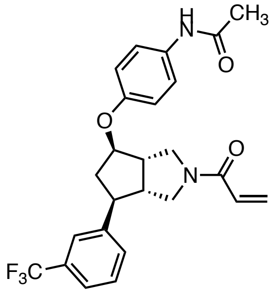

**FWG-47B**

Compound **FWG-47B** (**27**) was synthesized from **63** according to **Procedure E** (100 μmol scale; 2 h), followed by **Procedure F1, Procedure G** and **Procedure H** in EtOAc and was obtained as a colorless solid (31 mg, 69% over 4 steps; prep. TLC EtOAc and RP-MPLC acetonitrile/$H_2O$).

**$^1$H NMR** (400 MHz, $CDCl_3$, **FWG-47B**) δ 7.65 – 7.38 (m, 7H), 6.91 – 6.77 (m, 2H), 6.57 – 6.33 (m, 2H), 5.71 (q, *J* = 7.0 Hz, 1H), 4.73 – 4.60 (m, 1H), 4.12 – 3.88 (m, 1H), 3.83 – 3.46 (m, 3H), 3.16 – 2.88 (m, 3H), 2.87 – 2.72 (m, 1H), 2.14 (s, 3H), 2.13 – 2.04 (m, 1H) ppm.

**$^{19}$F NMR** {1H} (376 MHz, $CDCl_3$, **FWG-47B**) δ -65.16 (s, 1.1F), -65.19 (s, 1.9F) ppm.

**HRMS** (**FWG-47B**) m/z calc. for $C_{25}H_{26}F_3N_2O_3{}^+$ $[M+H]^+$ 405.2178 found 405.2178.

#### NMR Spectra

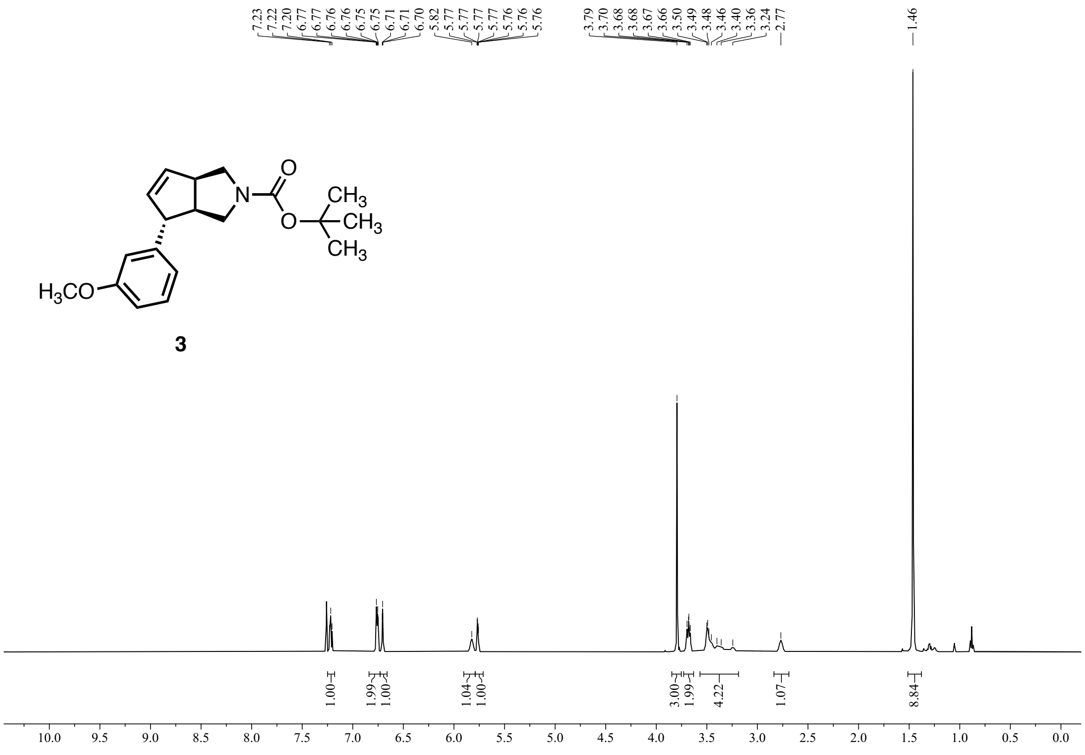


**Figure S2:** $^{1}$H NMR (600 MHz, $CDCl_3$) of **3**.

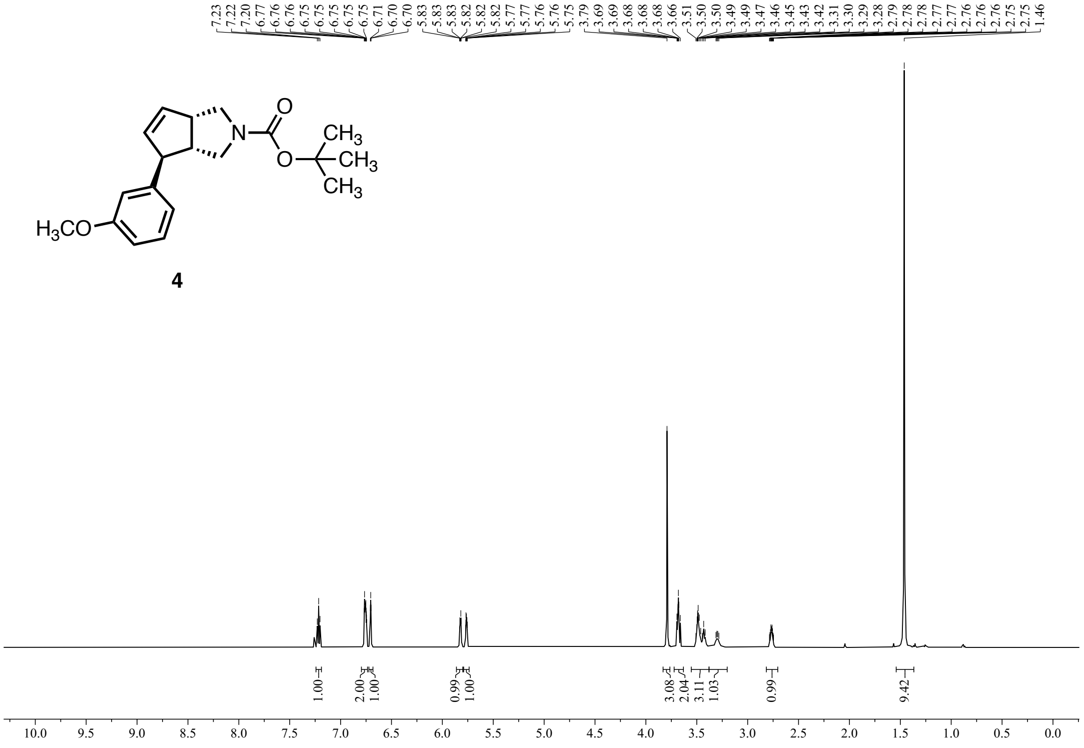


**Figure S3:** $^{1}$H NMR (600 MHz, $CDCl_3$) of **4**.

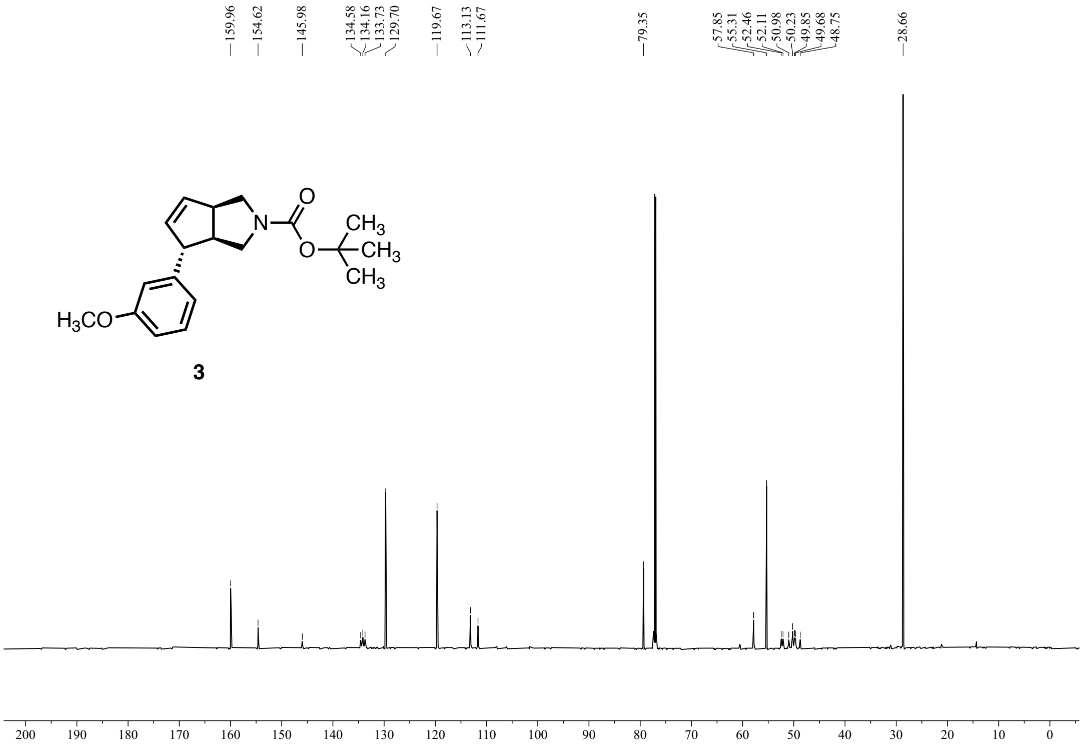


**Figure S4:** $^{13}C$ NMR (151 MHz, $CDCl_3$) of **3**.

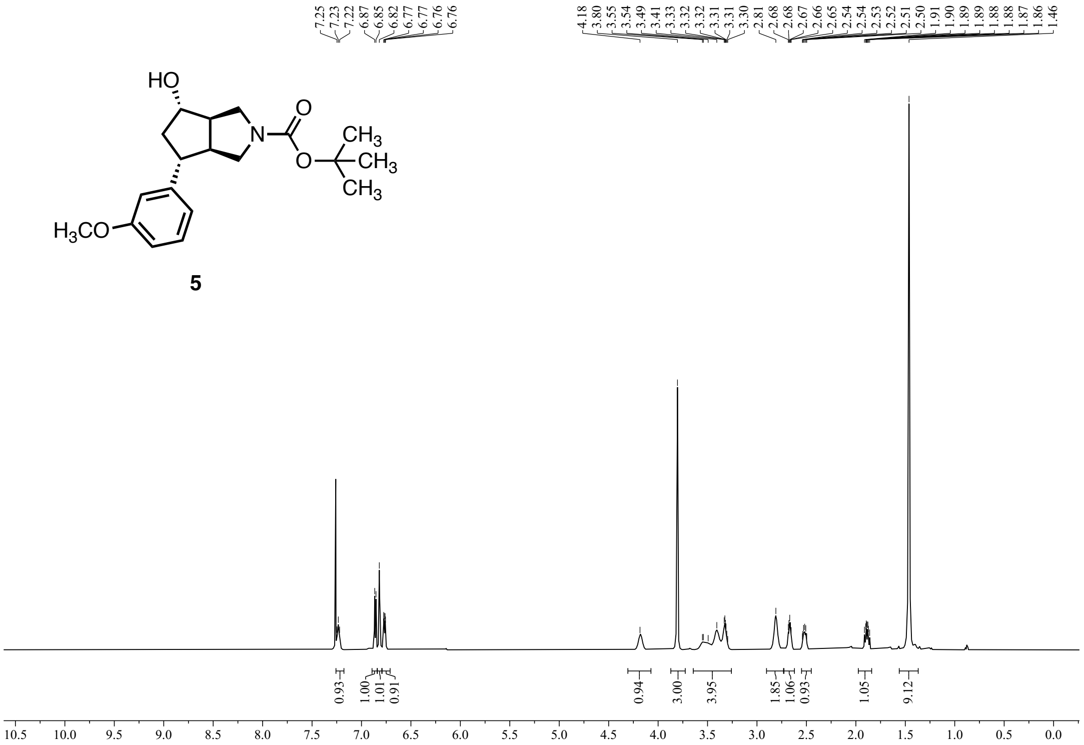


**Figure S5:** $^{1}$H NMR (600 MHz, $CDCl_3$) of **5**.

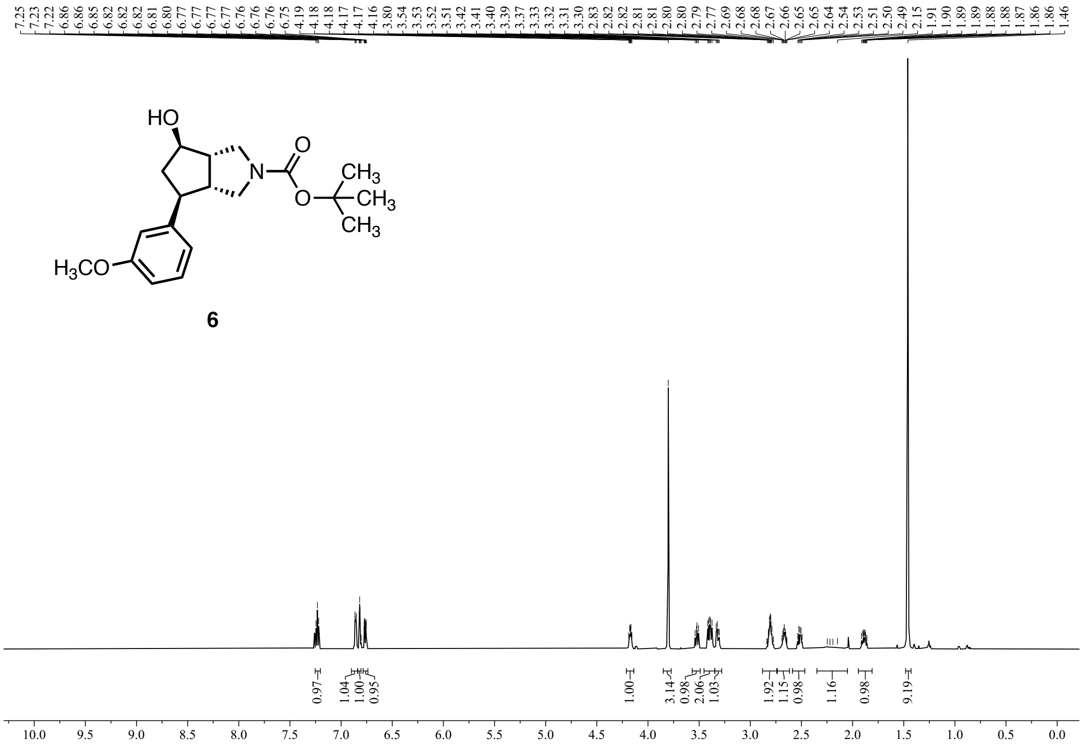


**Figure S6:** $^{1}$H NMR (600 MHz, $CDCl_3$) of **6**.

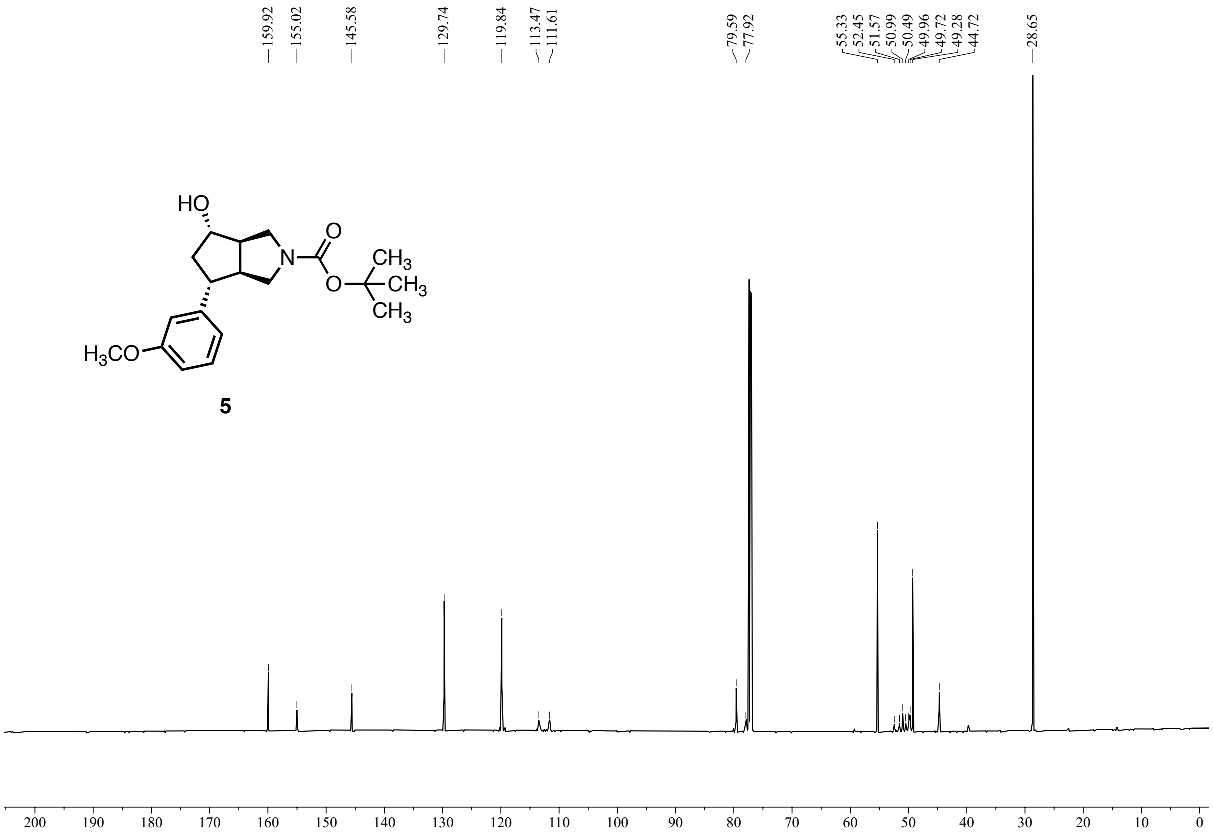


**Figure S7:** $^{13}$C NMR (151 MHz, CDCl$_3$) of **5**.

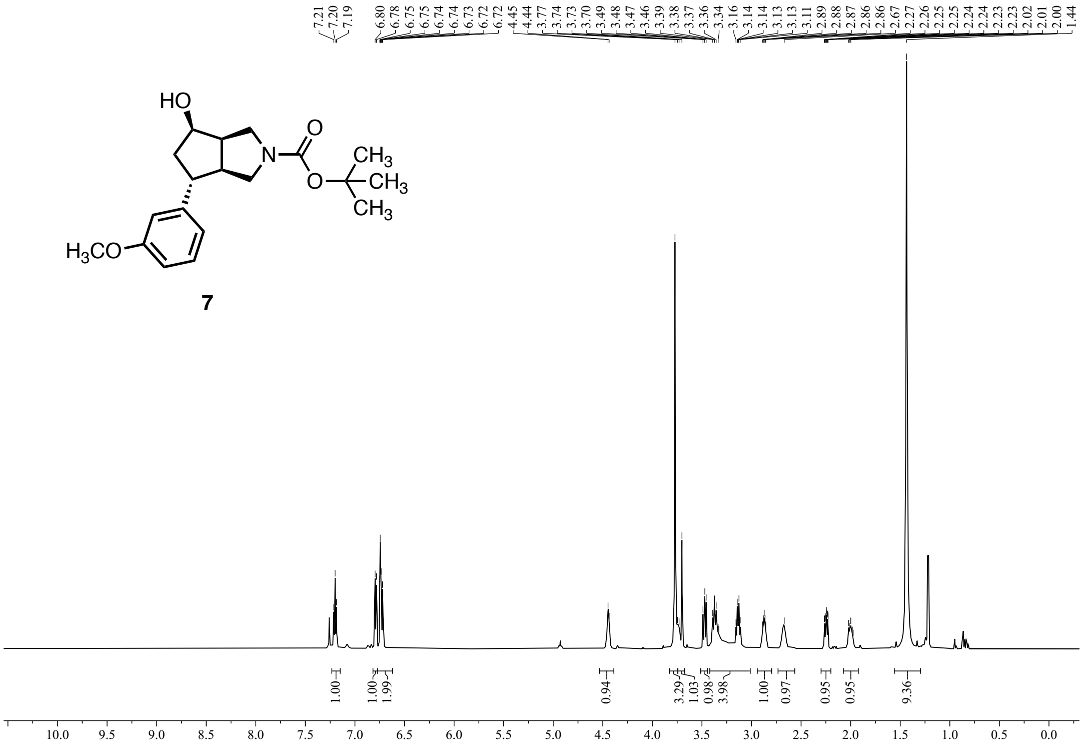


**Figure S8:** $^{1}$H NMR (600 MHz, $CDCl_3$) of **7**.

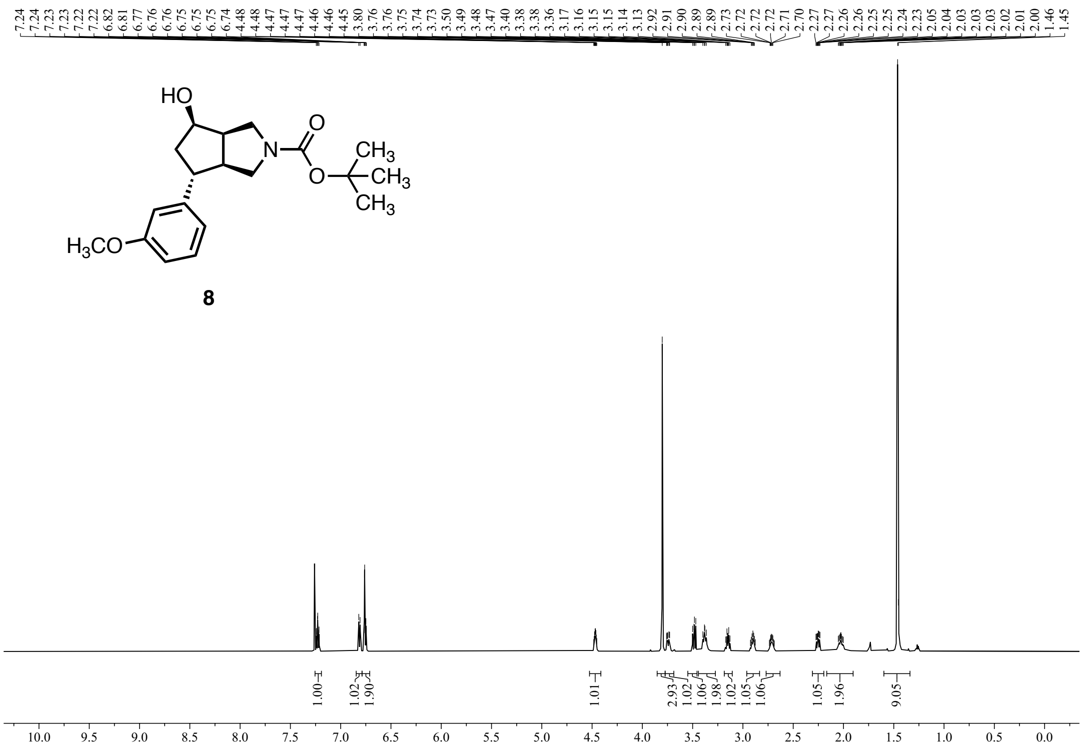


**Figure S9:** $^{1}$H NMR (600 MHz, $CDCl_3$) of **8**.

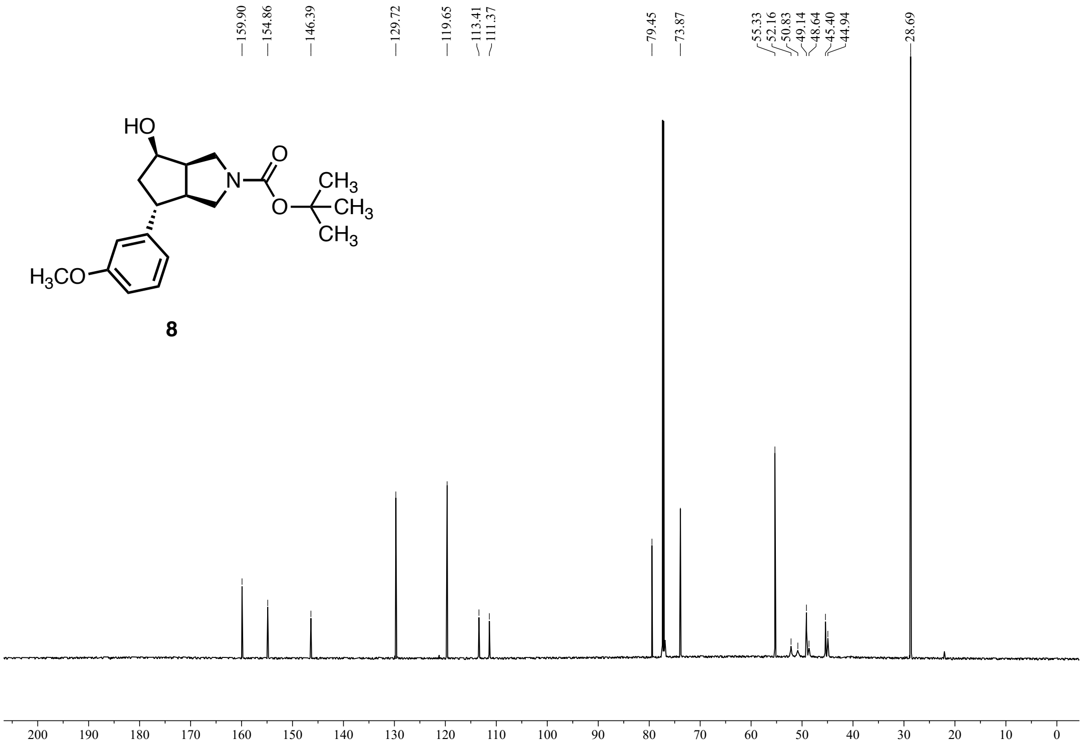


**Figure S10:** $^{13}$C NMR (151 MHz, $CDCl_3$) of **8**.

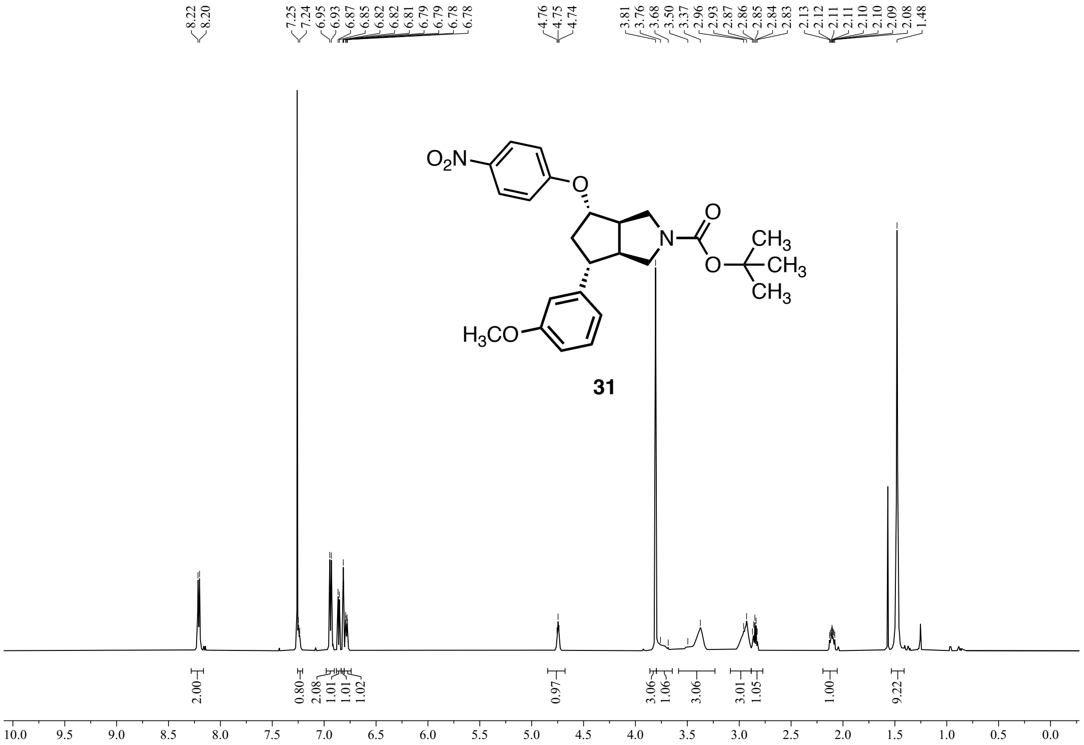


**Figure S11:** $^{1}$H NMR (600 MHz, $CDCl_3$) of **31**.

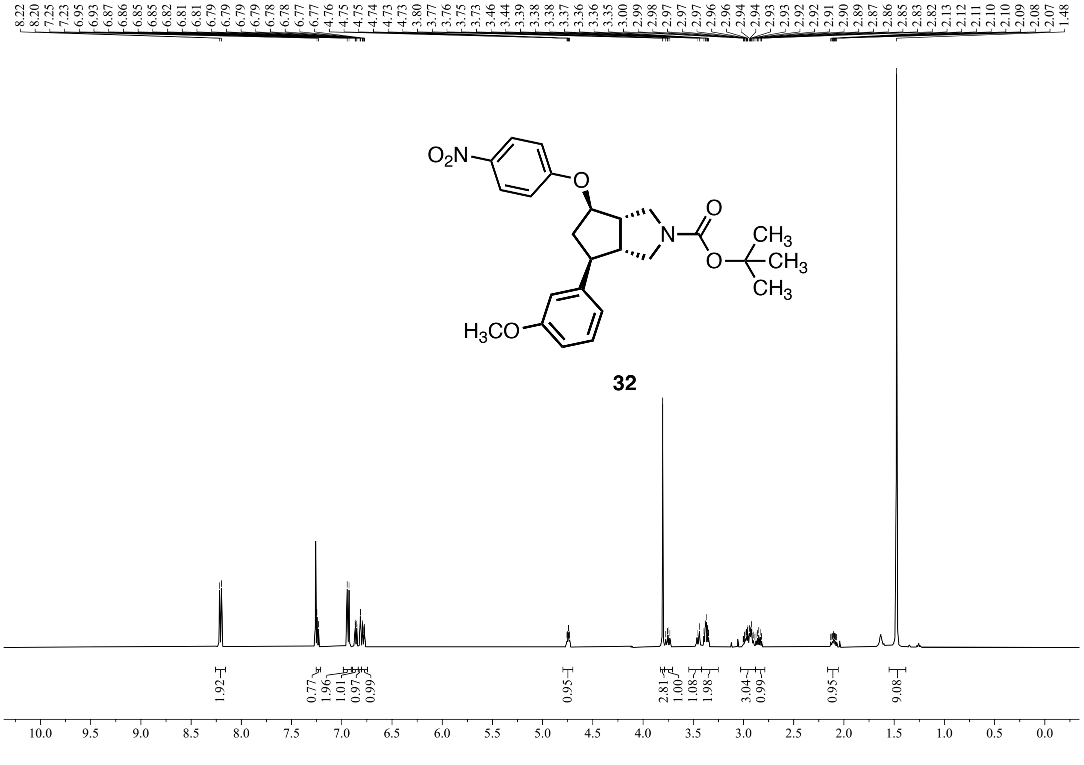


**Figure S12:** $^{1}$H NMR (500 MHz, $CDCl_3$) of **32**.

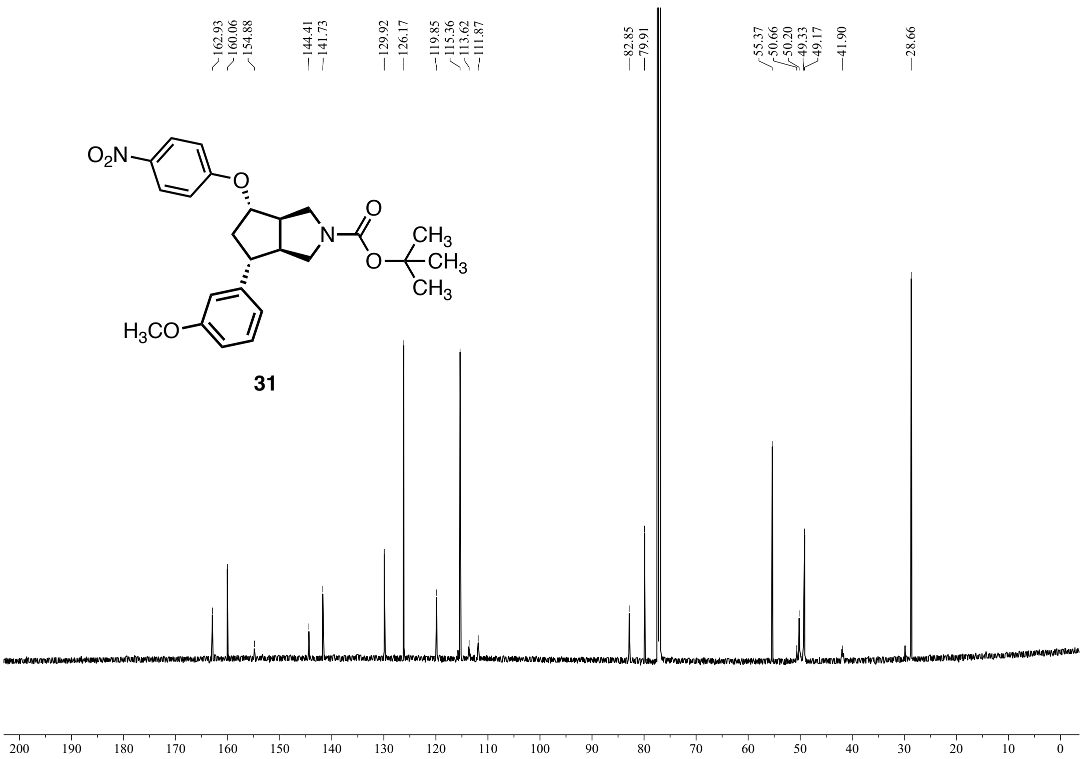


**Figure S13:** $^{13}C$ NMR (151 MHz, $CDCl_3$) of **31**.

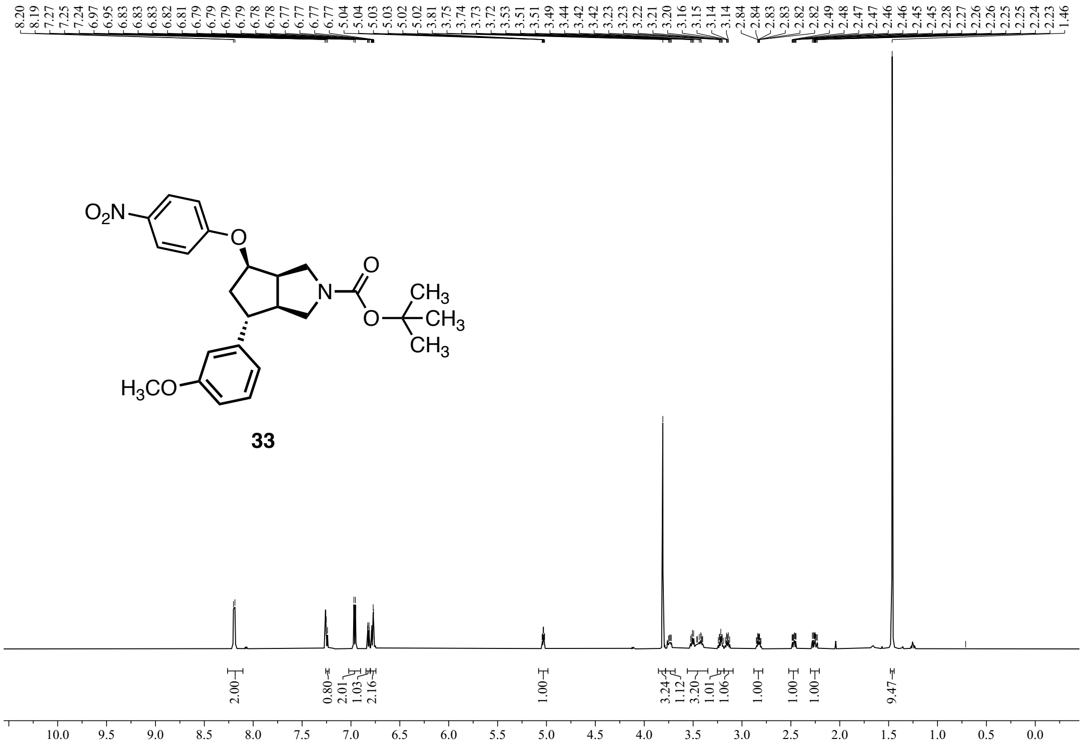


**Figure S14:** $^{1}$H NMR (600 MHz, $CDCl_3$) of **33**.

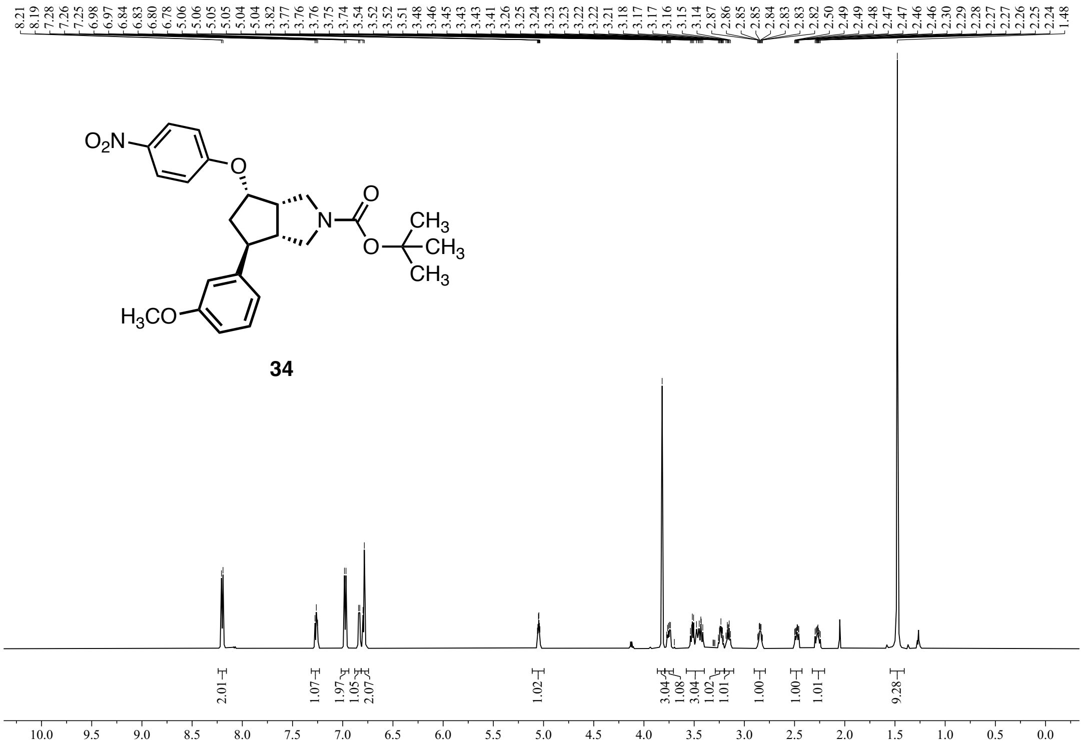


**Figure S15:** $^{1}$H NMR (600 MHz, $CDCl_3$) of **34**.

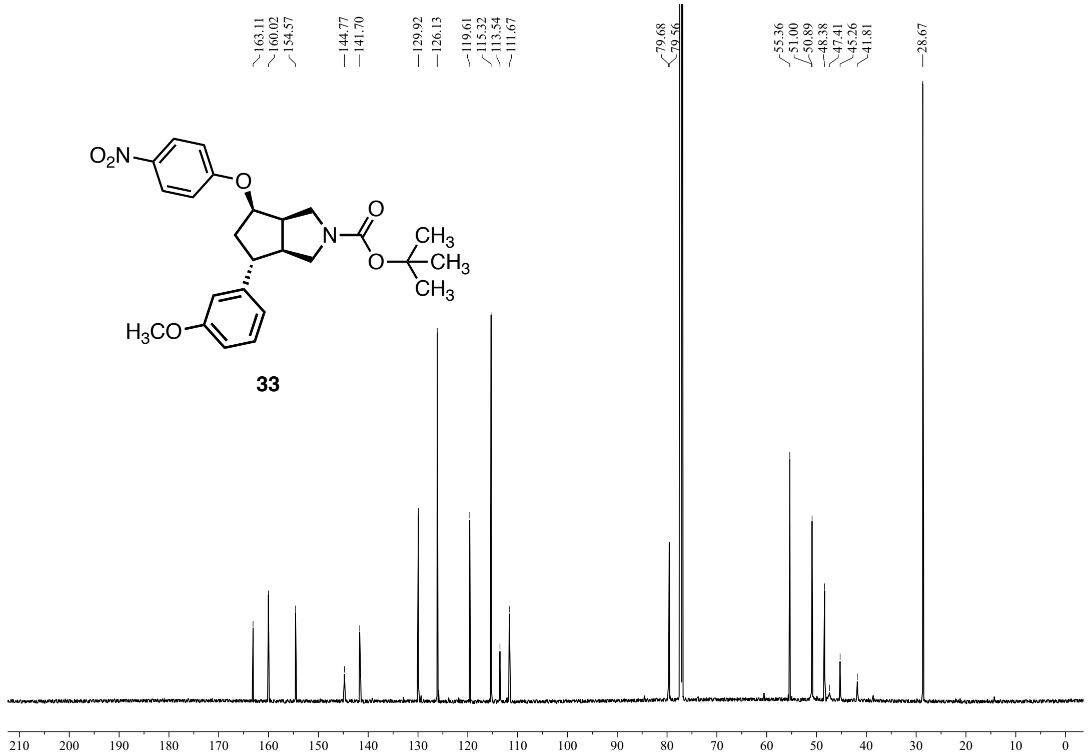


**Figure S16:** $^{13}C$ NMR (151 MHz, $CDCl_3$) of **33**.

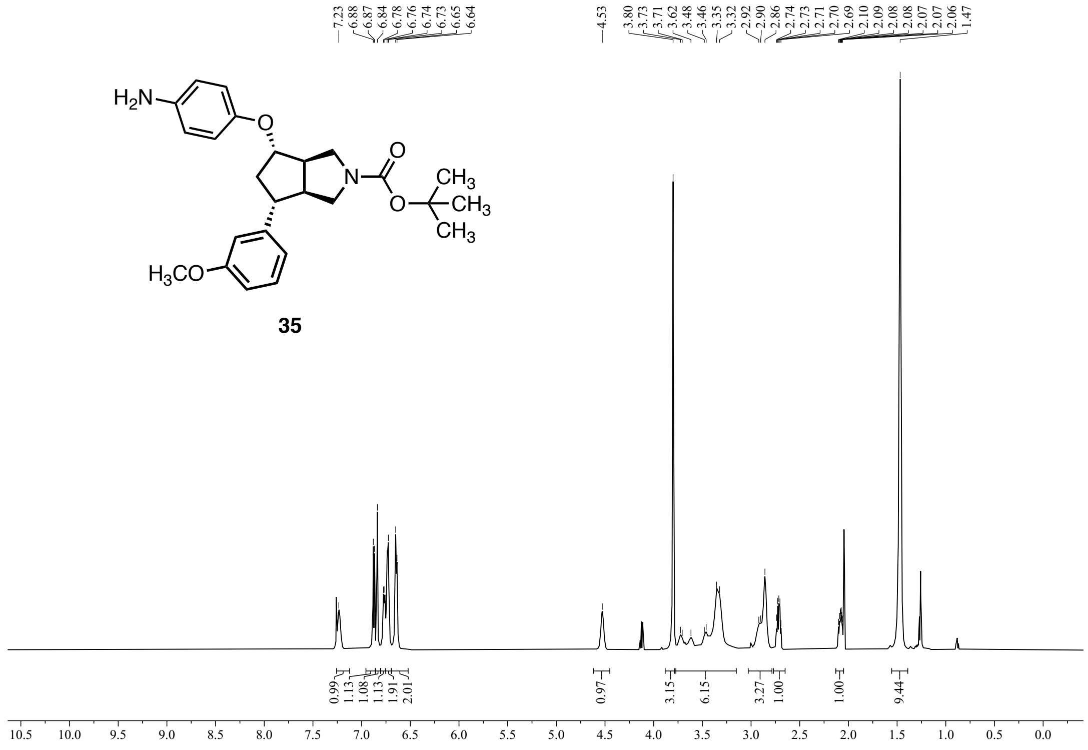


**Figure S17:** $^{1}$H NMR (600 MHz, $CDCl_3$) of **35**.

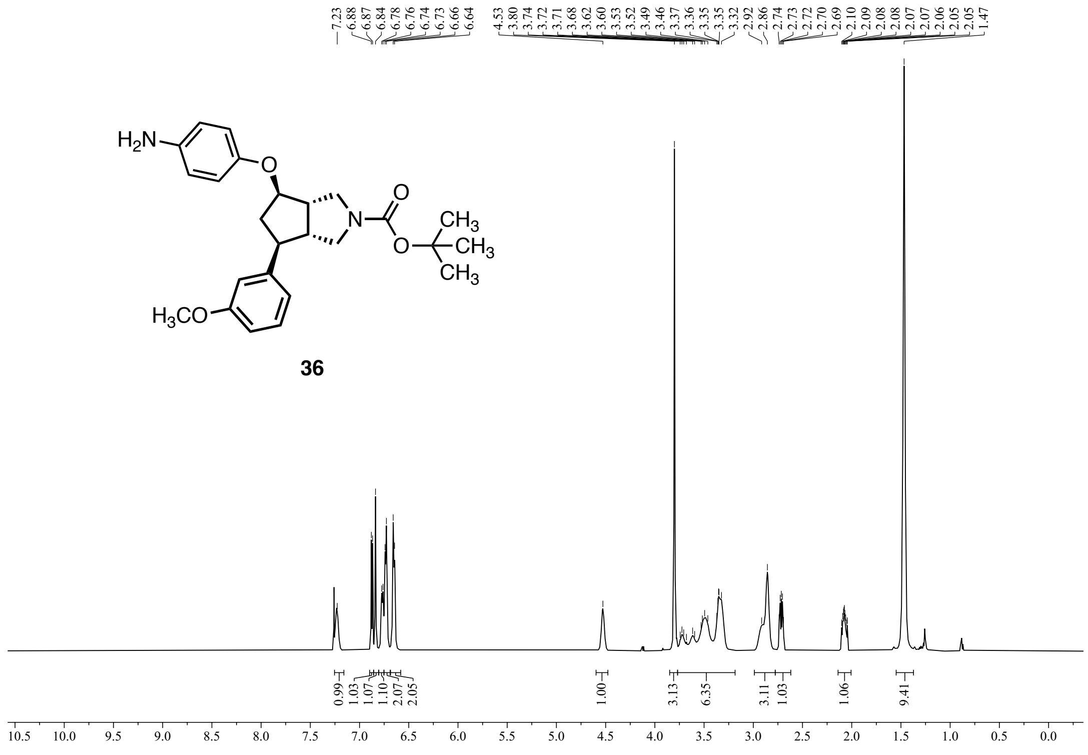


**Figure S18:** $^{1}$H NMR (600 MHz, $CDCl_3$) of **36**.

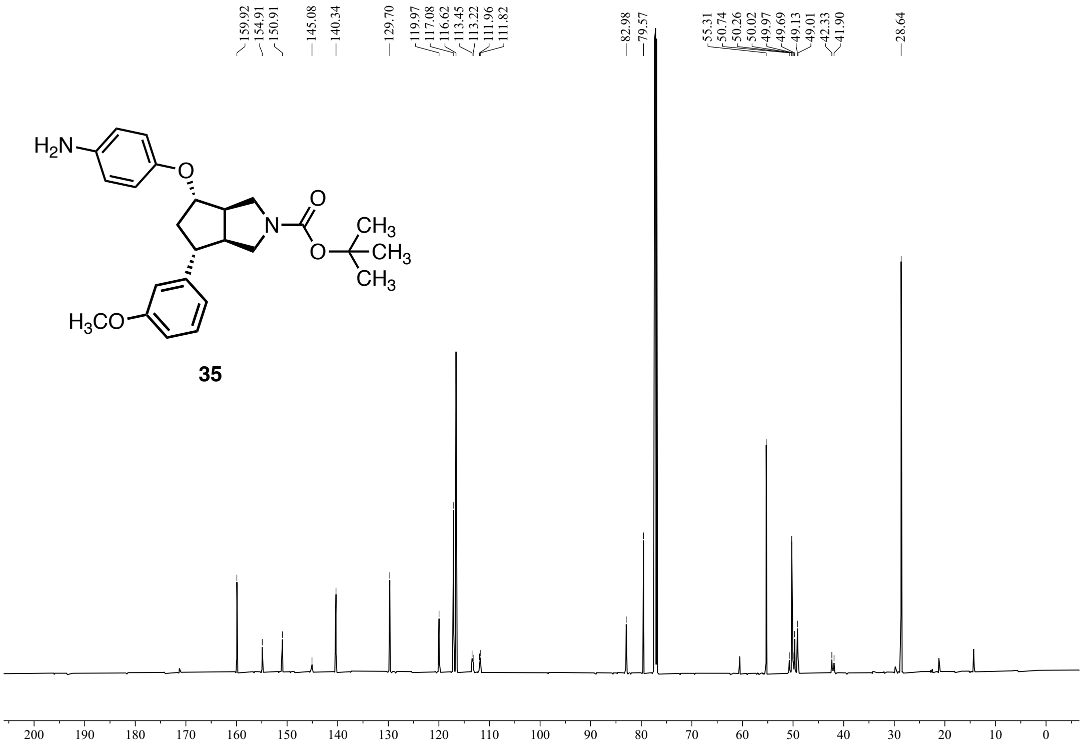


**Figure S19:** $^{13}C$ NMR (151 MHz, $CDCl_3$) of **35**.

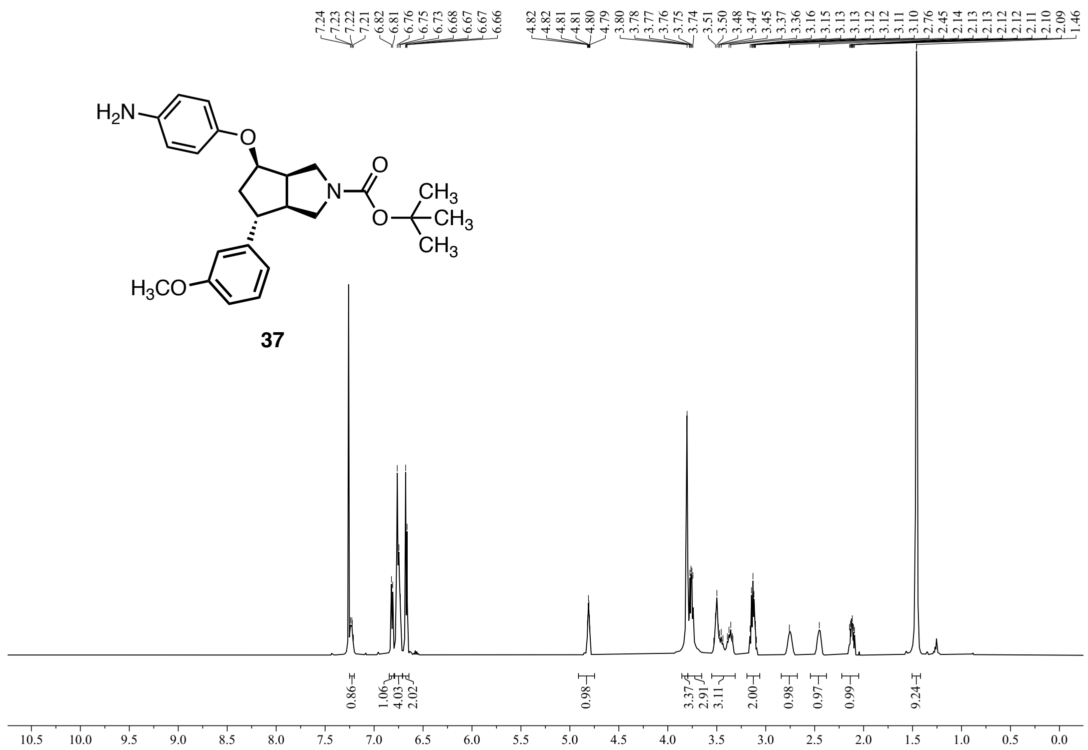

**Figure S20:** $^{1}$H NMR (600 MHz) of **37**.

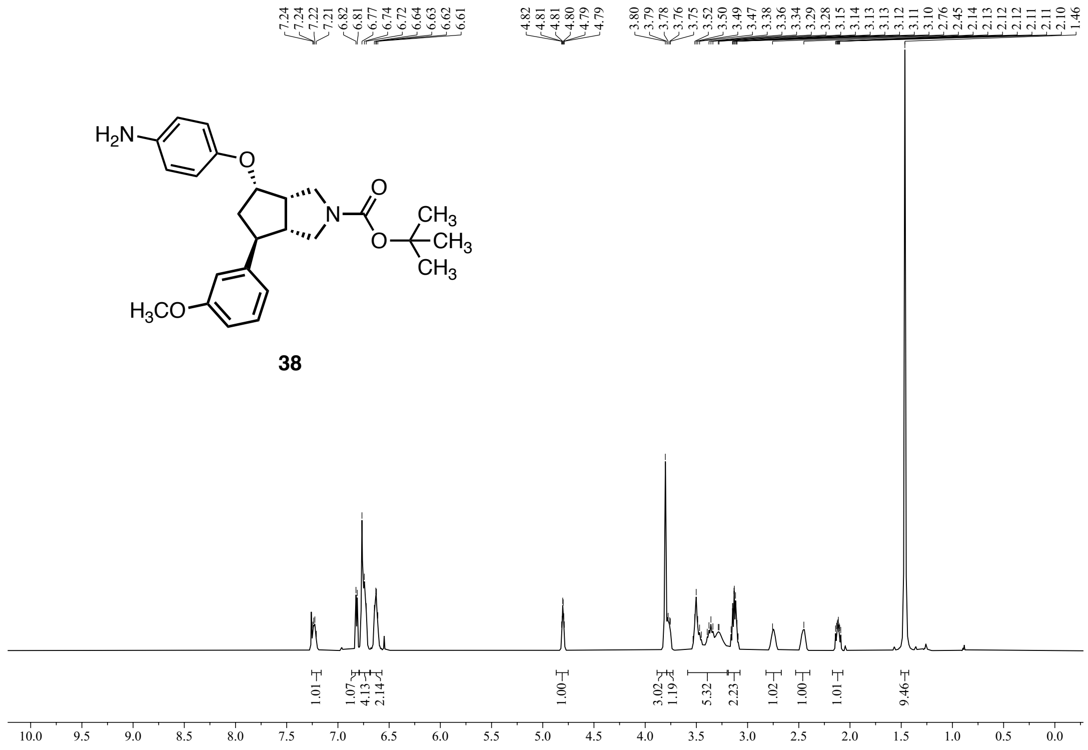

**Figure S21:** $^{1}$H NMR (600 MHz, $CDCl_3$) of **38**.

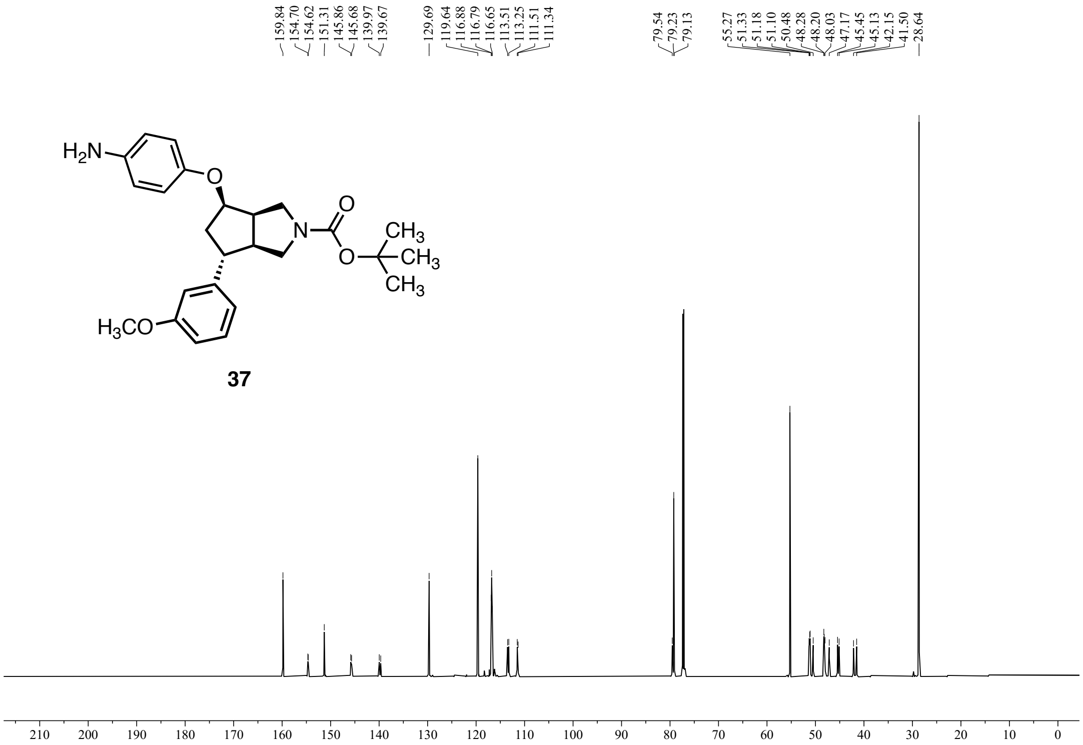


37

**Figure S22:** $^{13}C$ NMR (151 MHz, $CDCl_3$) of **37**.

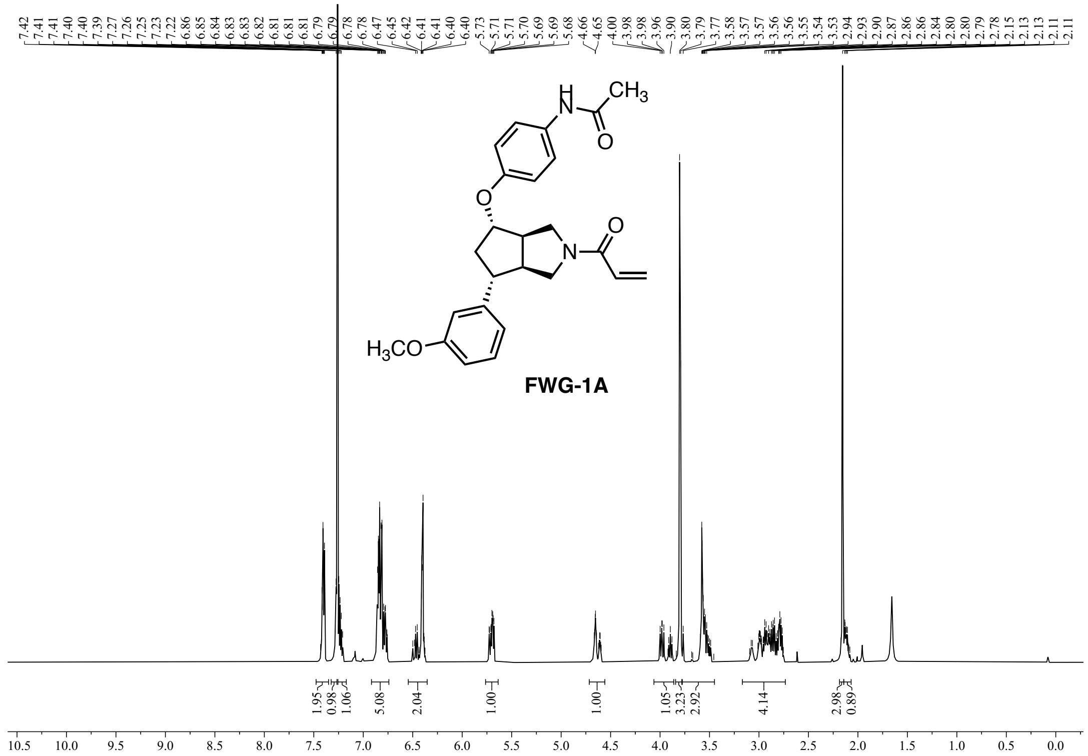

**Figure S23:** $^{1}H$ NMR (600 MHz, $CDCl_3$) of **FWG-1A**.

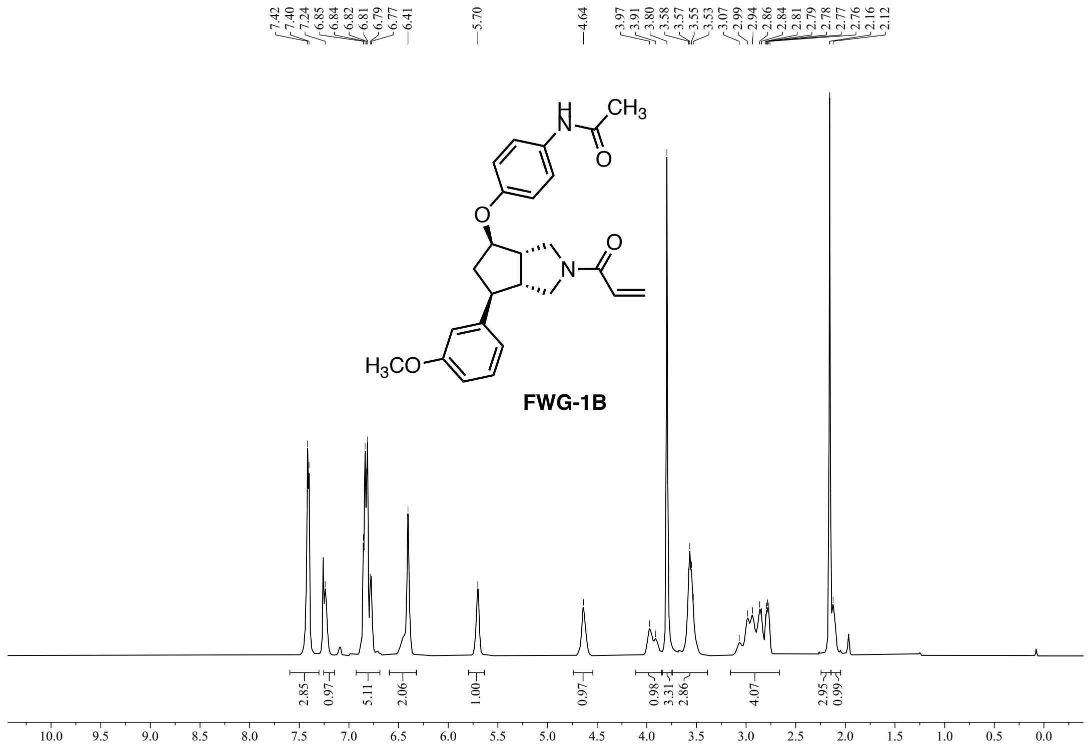

**Figure S24:** $^{1}H$ NMR (600 MHz, $CDCl_3$) of **FWG-1B**.

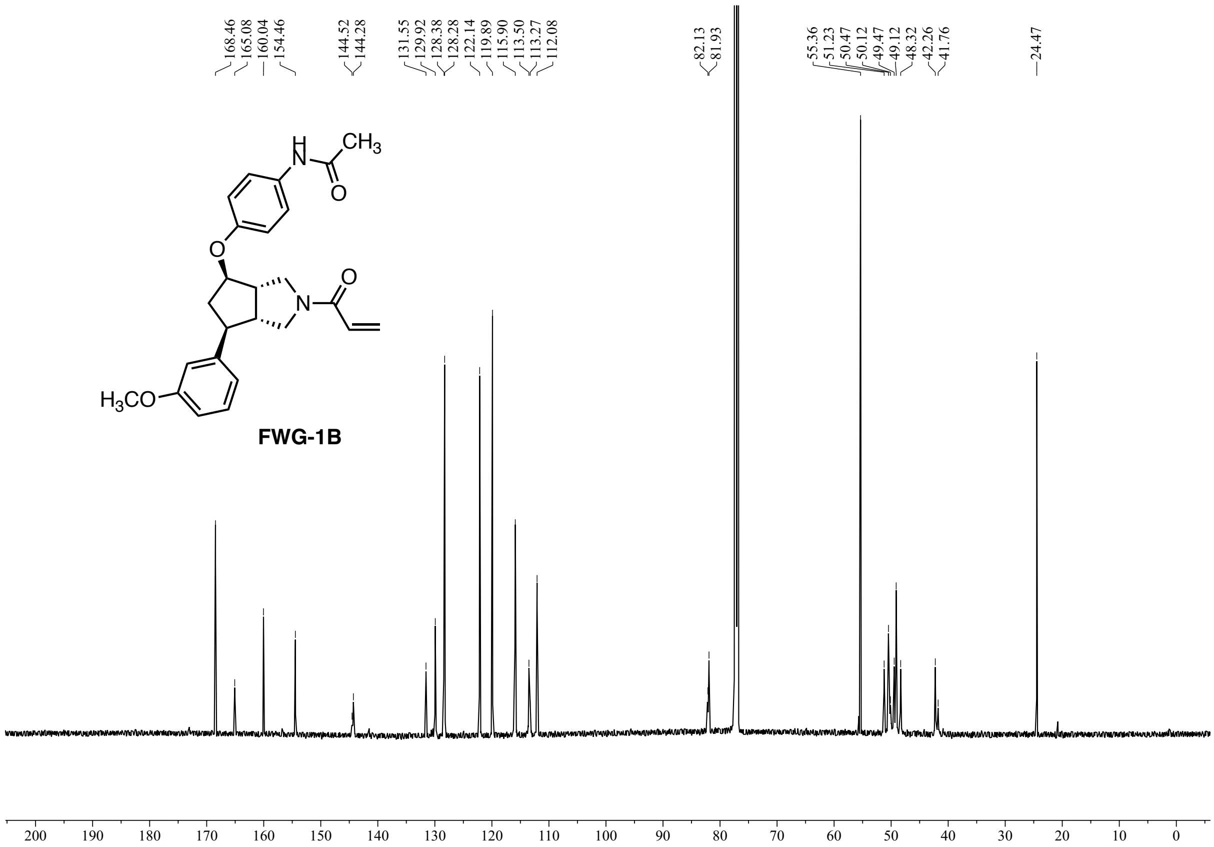


**Figure S25:** $^{13}$C NMR (151 MHz, $CDCl_3$) of **FWG-1B**.

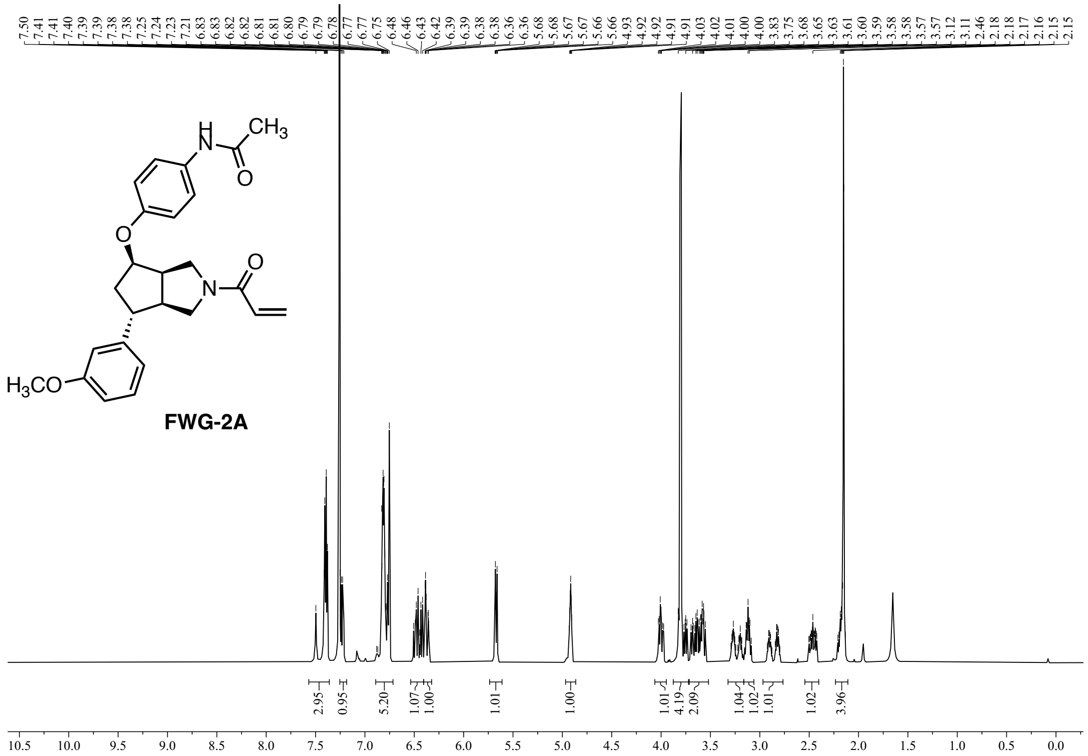

Figure S26: $^{1}$H NMR (600 MHz, $CDCl_3$) of **FWG-2A**.

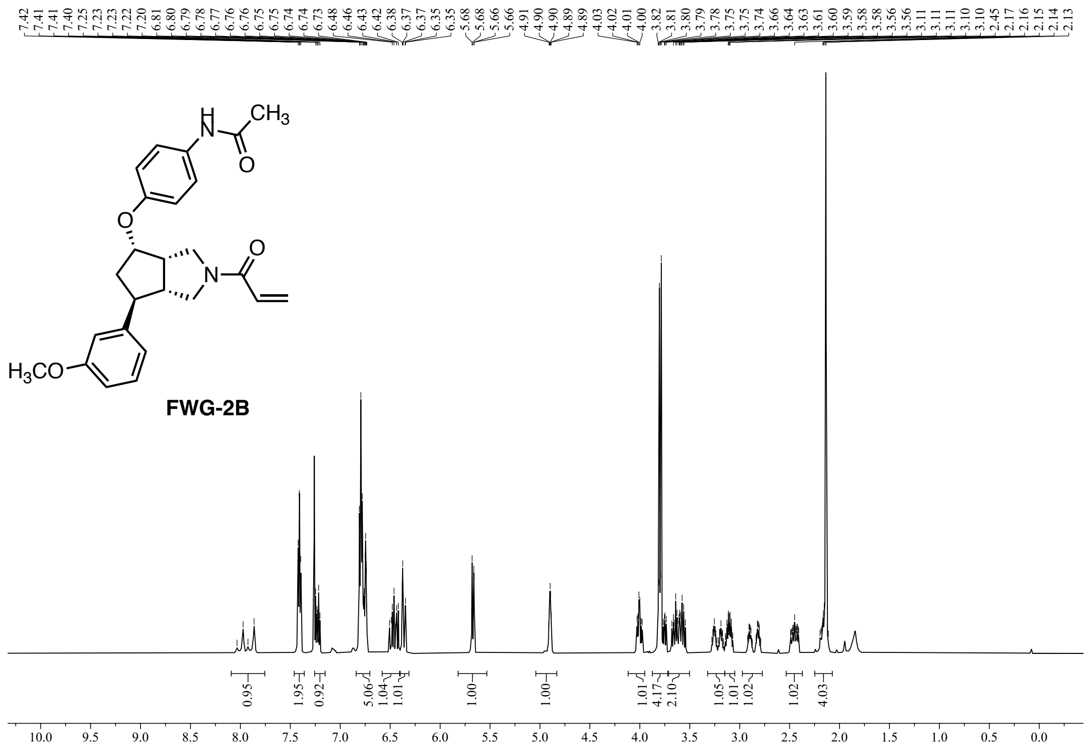

Figure S27: $^{1}$H NMR (600 MHz, $CDCl_3$) of **FWG-2B**.

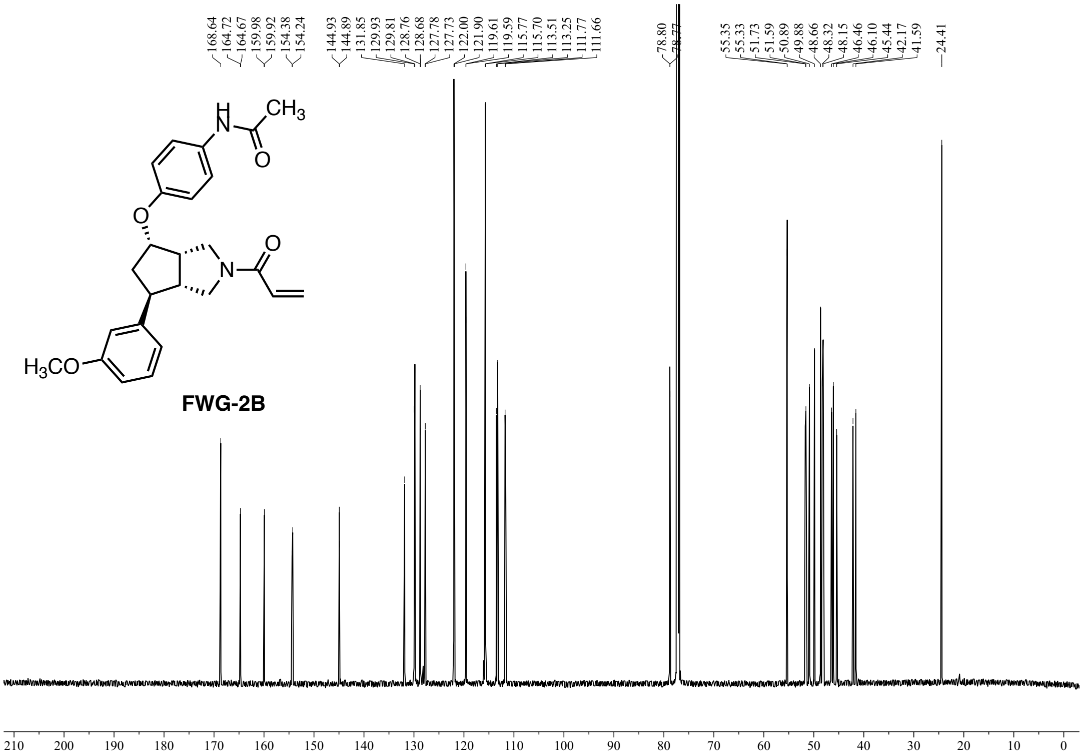

**Figure S28:** $^{13}$C NMR (151 MHz, $CDCl_3$) of **FWG-2B**.

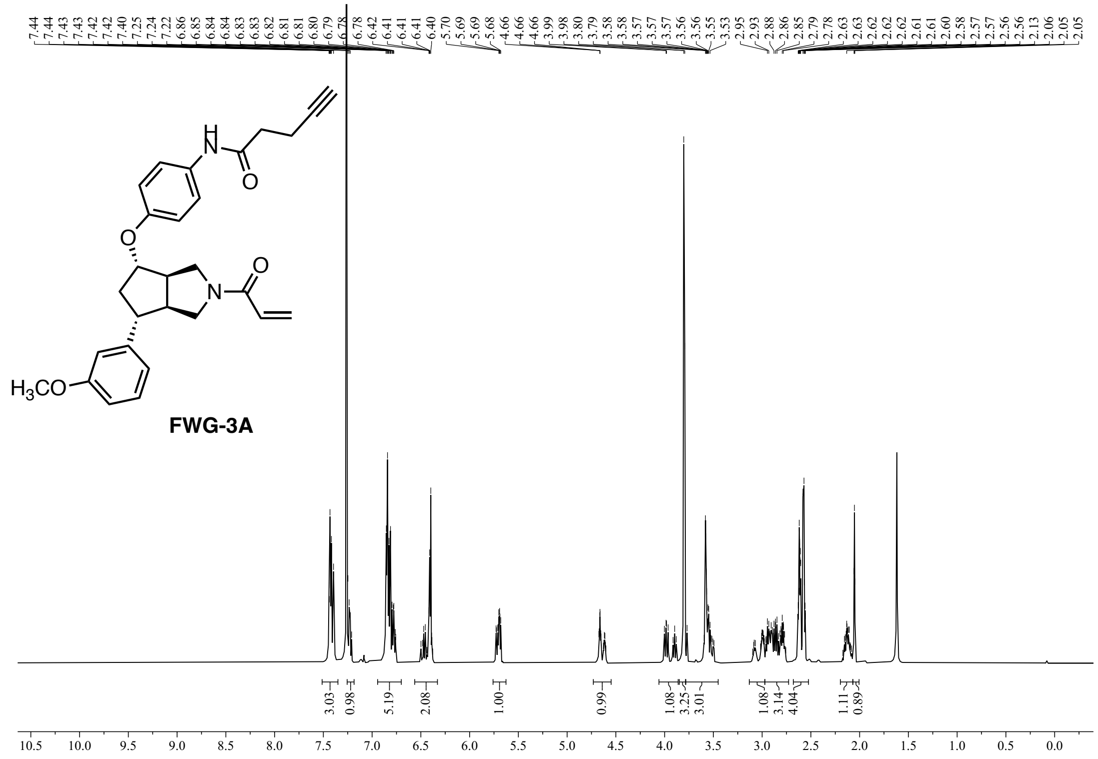


**Figure S29:** $^{1}$H NMR (600 MHz, $CDCl_3$) of **FWG-3A**.

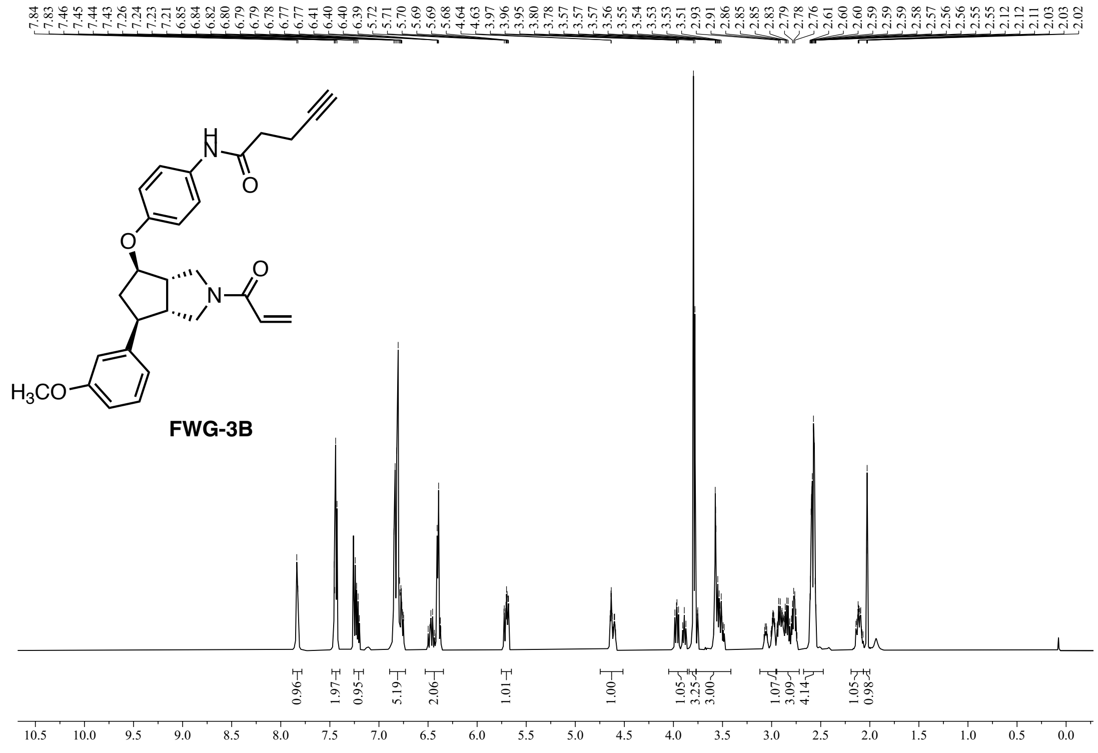


**Figure S30:** $^{1}$H NMR (600 MHz, $CDCl_3$) of **FWG-3B**.

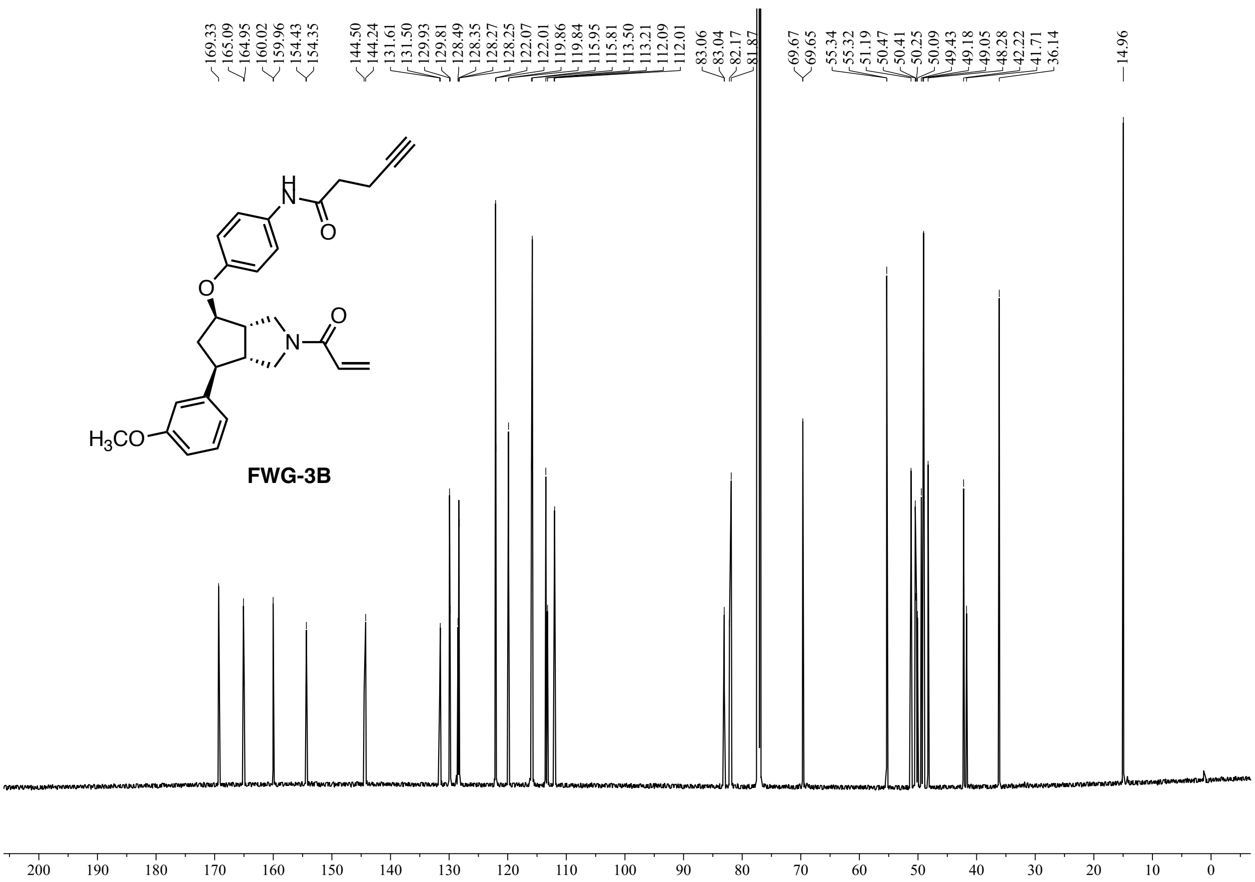


**Figure S31:** $^{13}C$ NMR (151 MHz, $CDCl_3$) of **FWG-3B**.

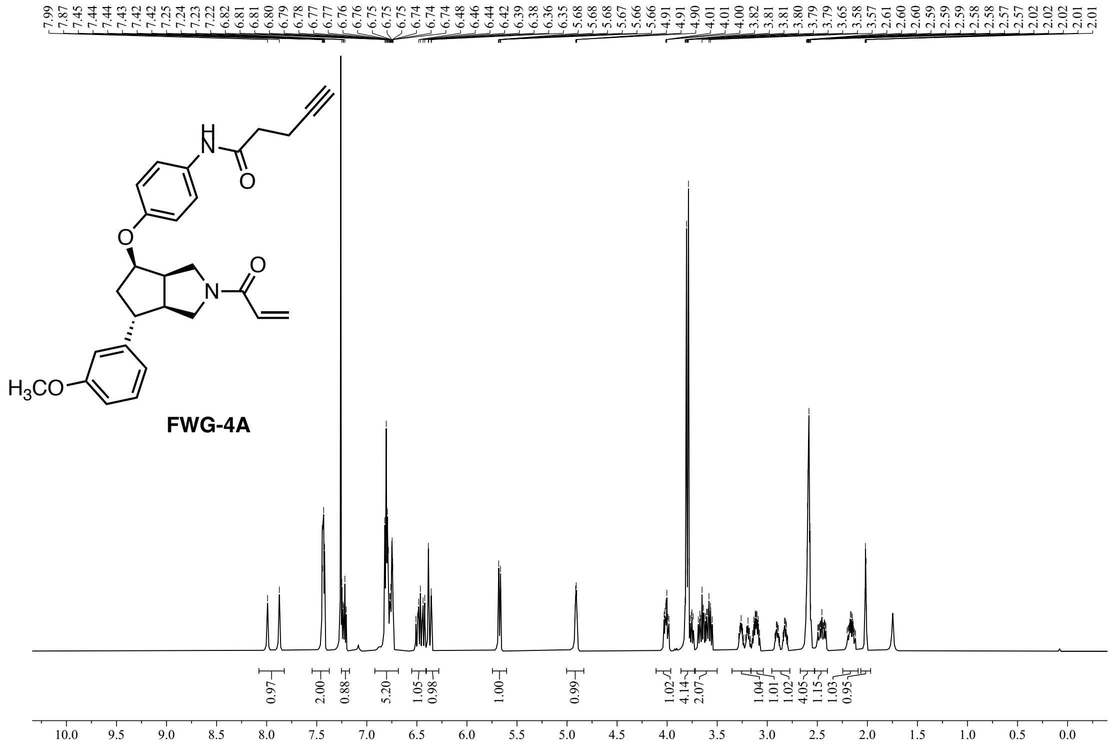

**Figure S32:** $^1$H NMR (600 MHz, $CDCl_3$) of **FWG-4A**.

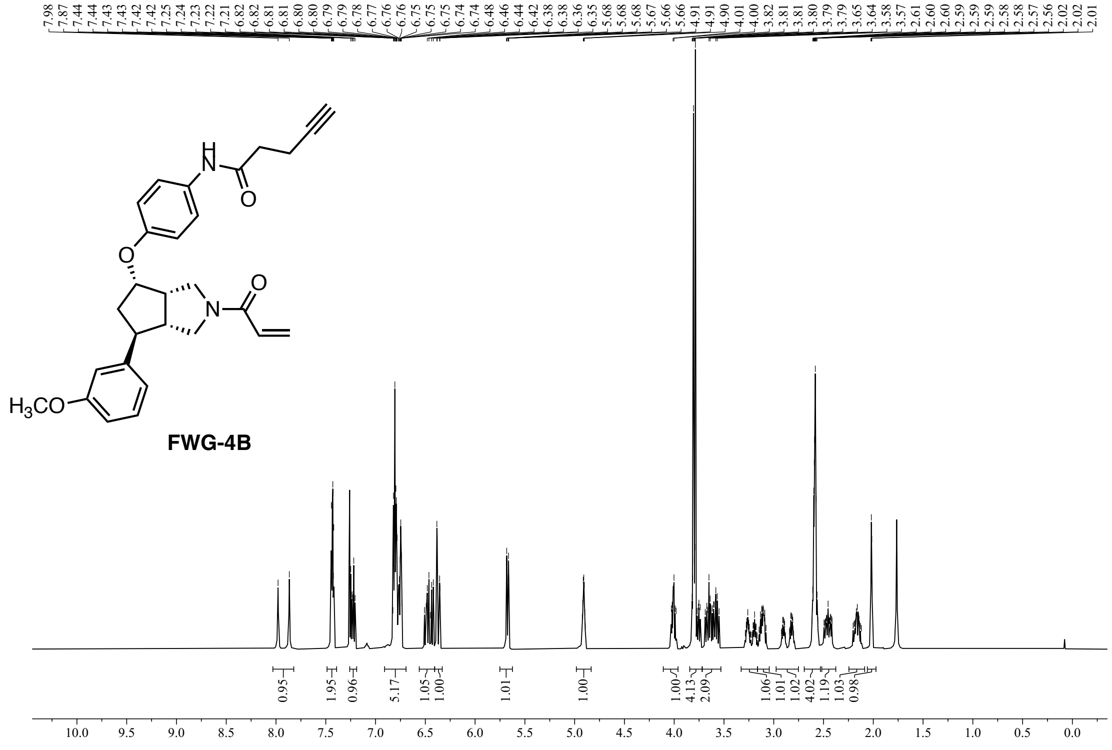

**Figure S33:** $^1$H NMR (600 MHz, $CDCl_3$) of **FWG-4B**.

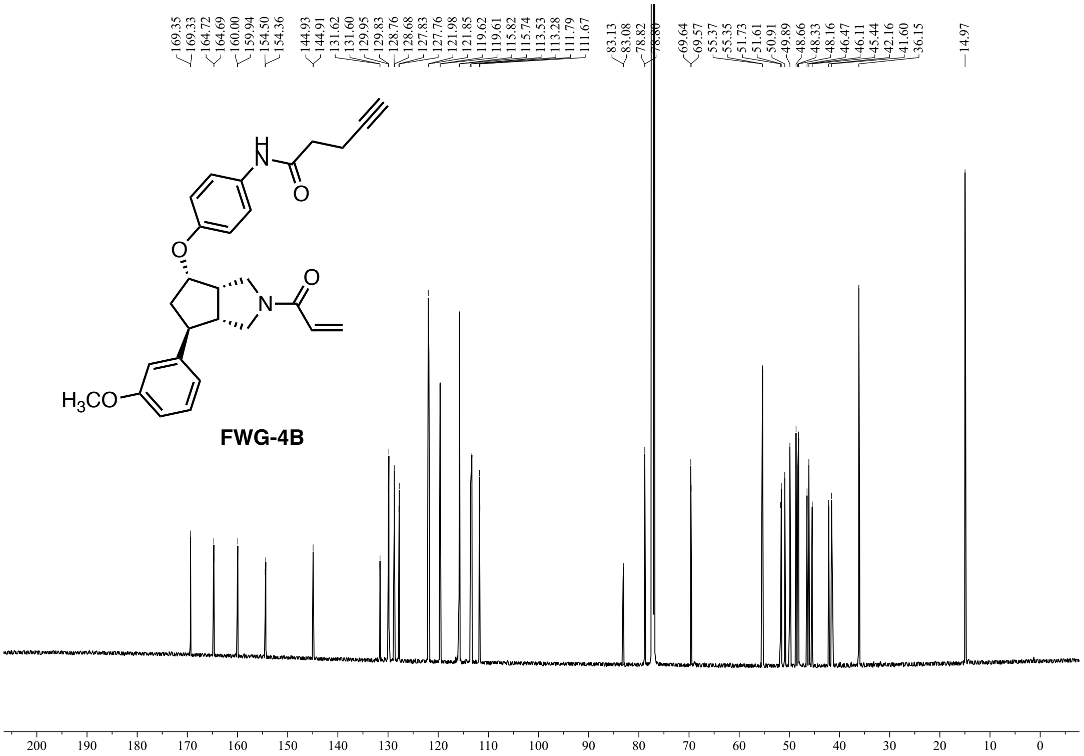


**Figure S34:** $^{13}C$ NMR (151 MHz, $CDCl_3$) of **FWG-4B**.

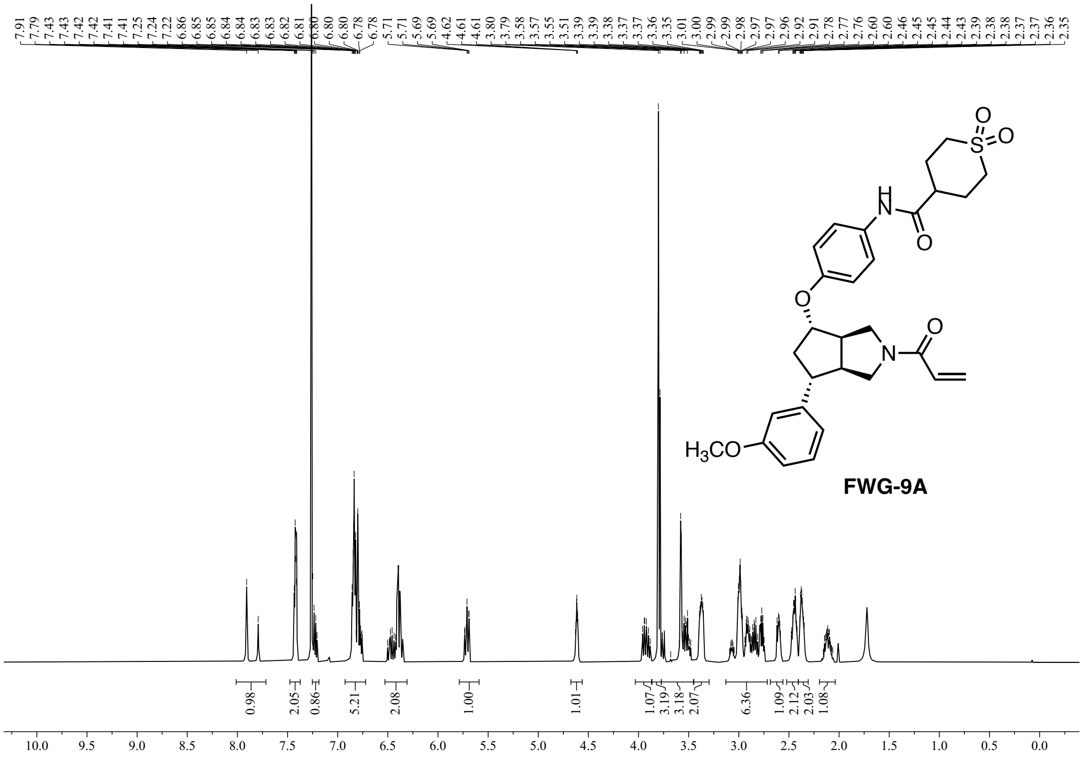

**Figure S35:** $^{1}H$ NMR (600 MHz, $CDCl_3$) of **FWG-9A**.

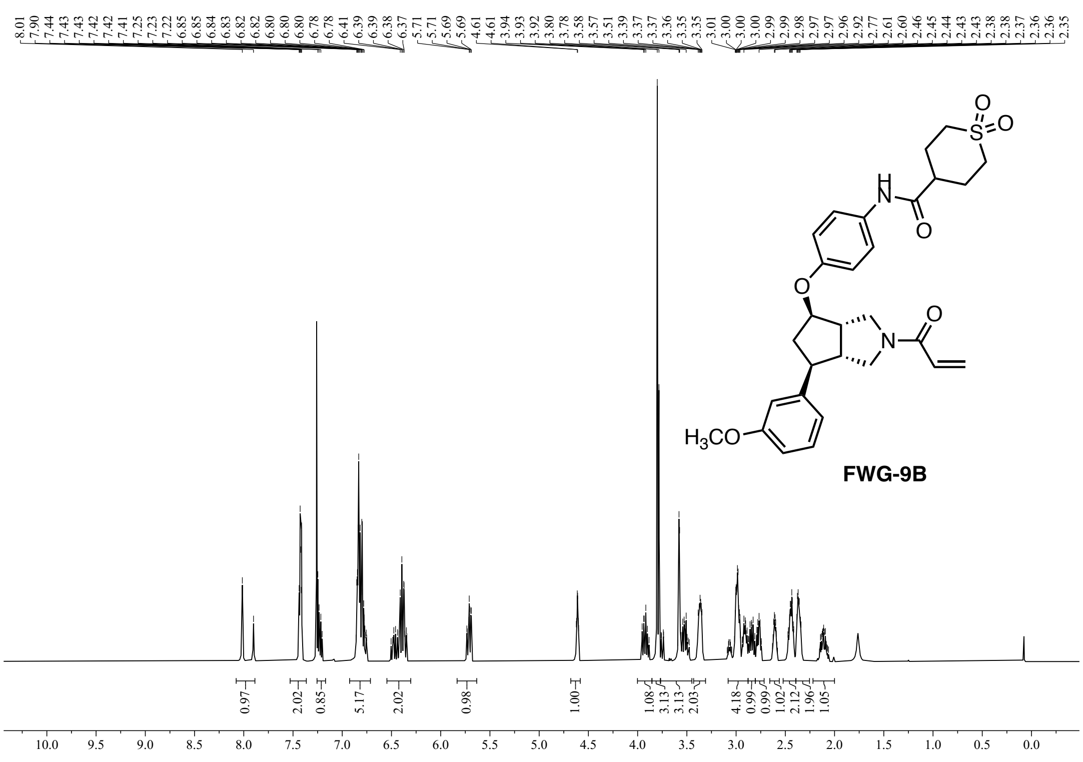

**Figure S36:** $^{1}H$ NMR (600 MHz, $CDCl_3$) of **FWG-9B**.

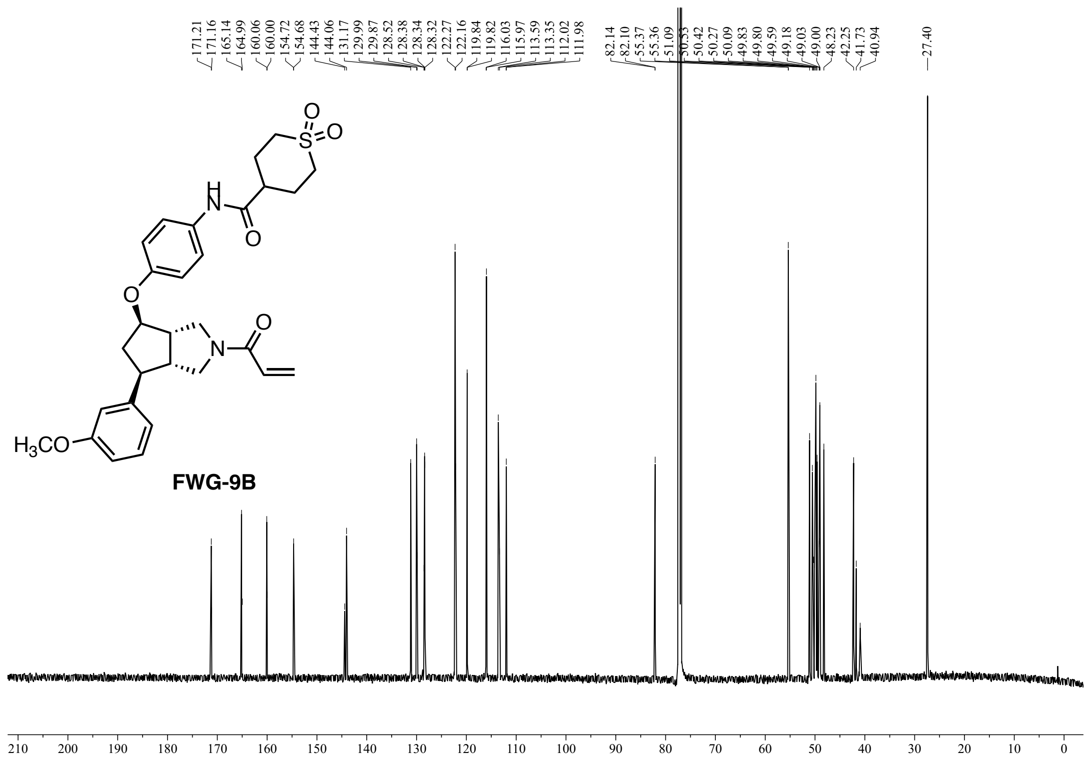

**Figure S37:** $^{13}$C NMR (151 MHz, $CDCl_3$) of **FWG-9B**.

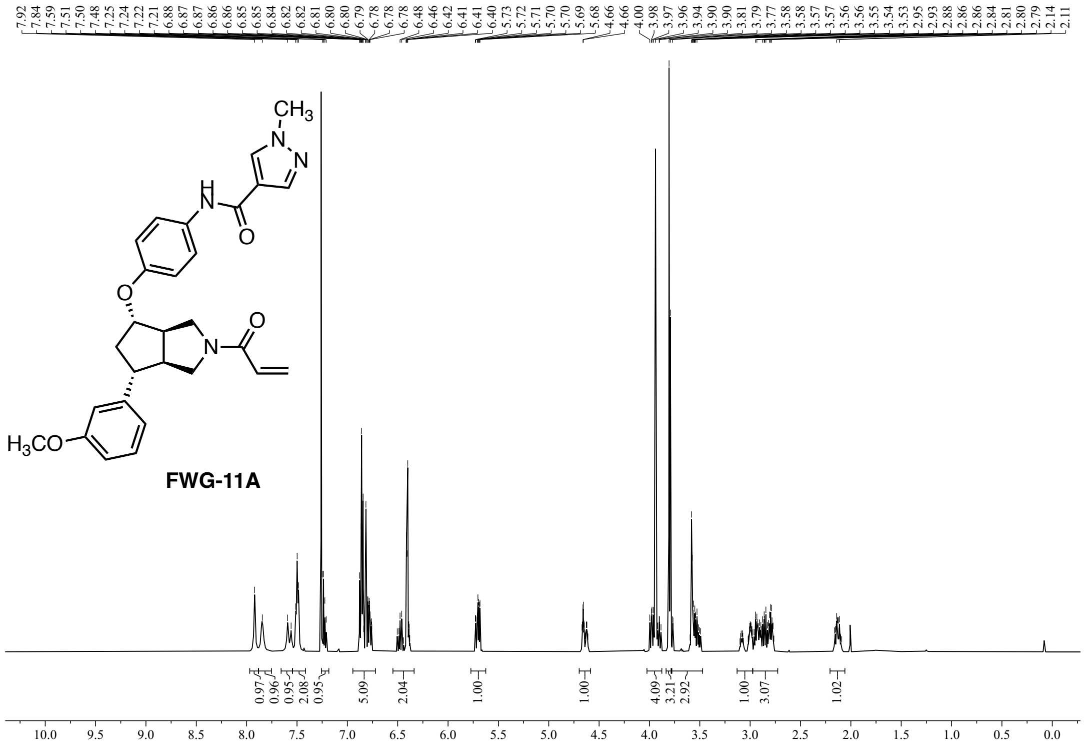


**Figure S38:** $^{1}$H NMR (600 MHz, $CDCl_3$) of **FWG-11A**.

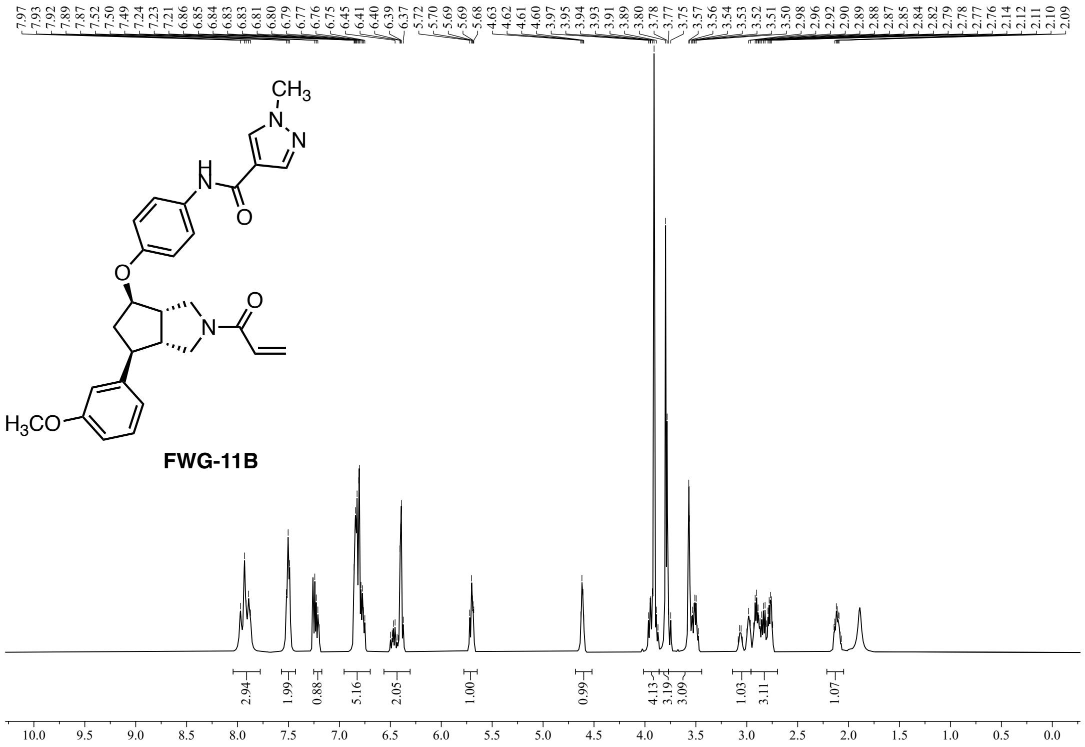


**Figure S39:** $^{1}$H NMR (600 MHz, $CDCl_3$) of **FWG-11B**.

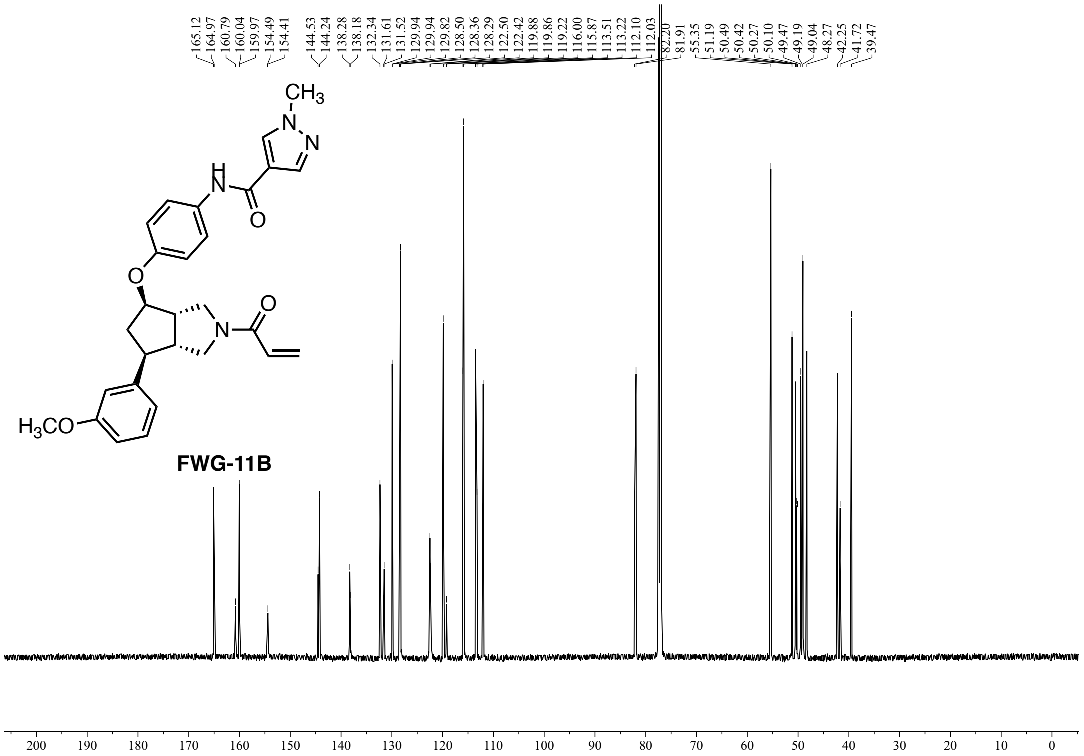

**Figure S40:** $^{13}$C NMR (151 MHz, $CDCl_3$) of **FWG-11B**.

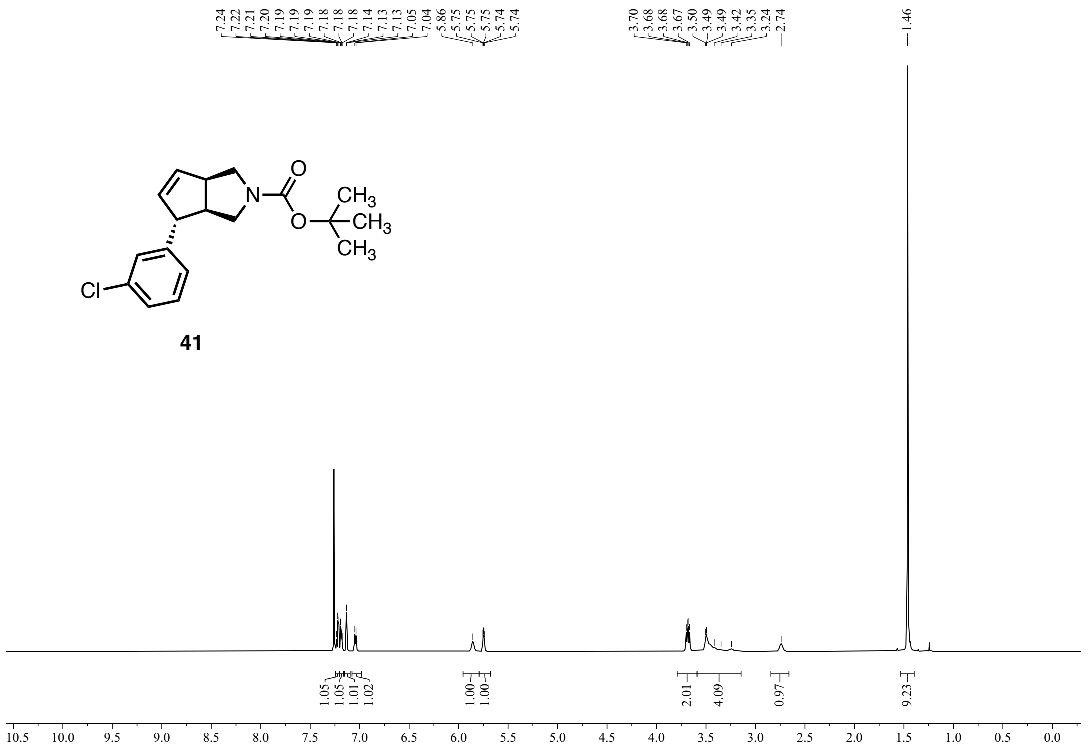


**Figure S41:** $^{1}$H NMR (600 MHz, $CDCl_3$) of **41**.

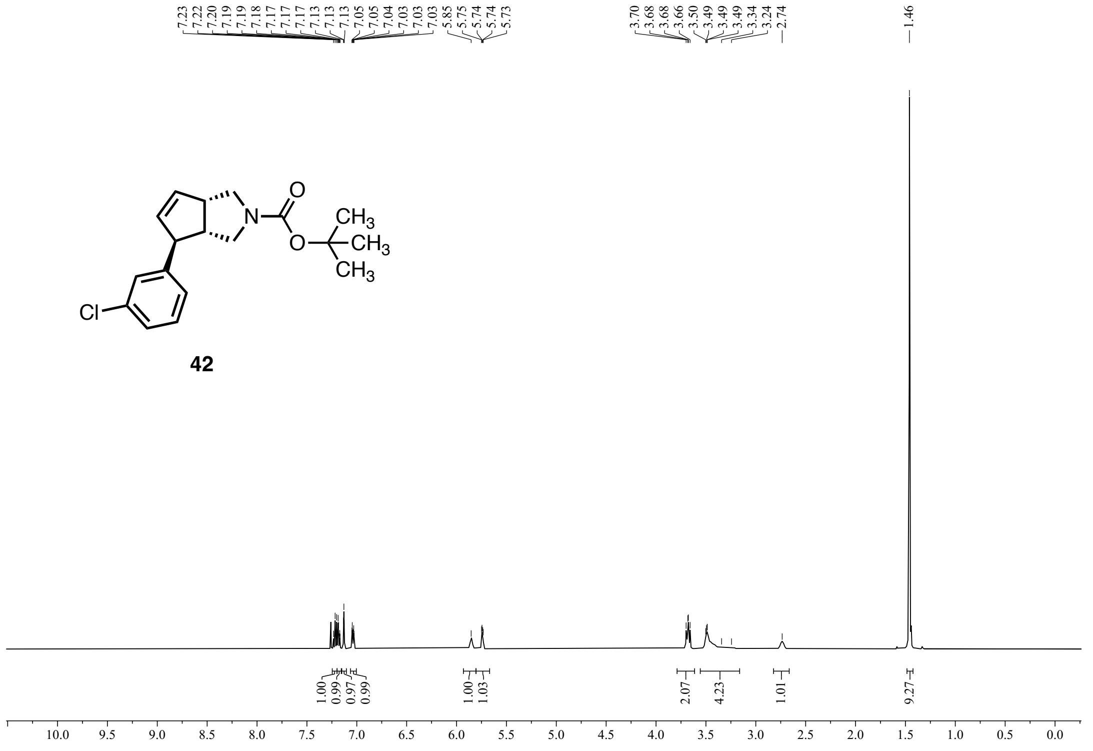


**Figure S42:** $^{1}$H NMR (600 MHz, $CDCl_3$) of **42**.

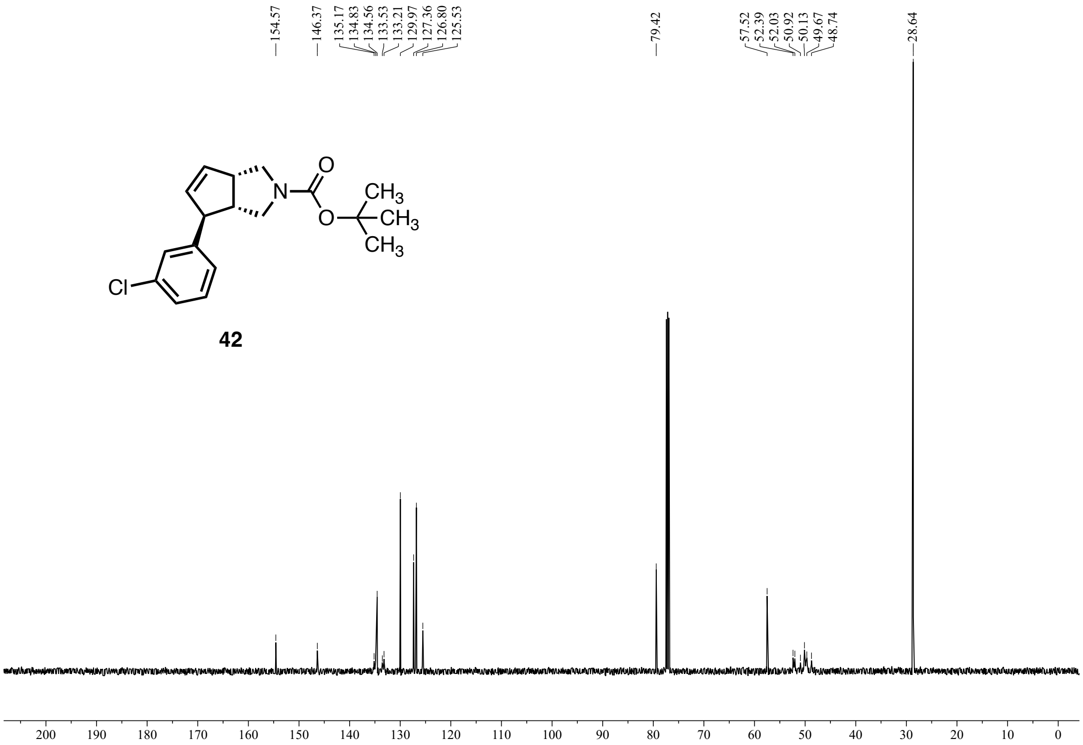


**Figure S43:** $^{13}C$ NMR (151 MHz, $CDCl_3$) of **42**.

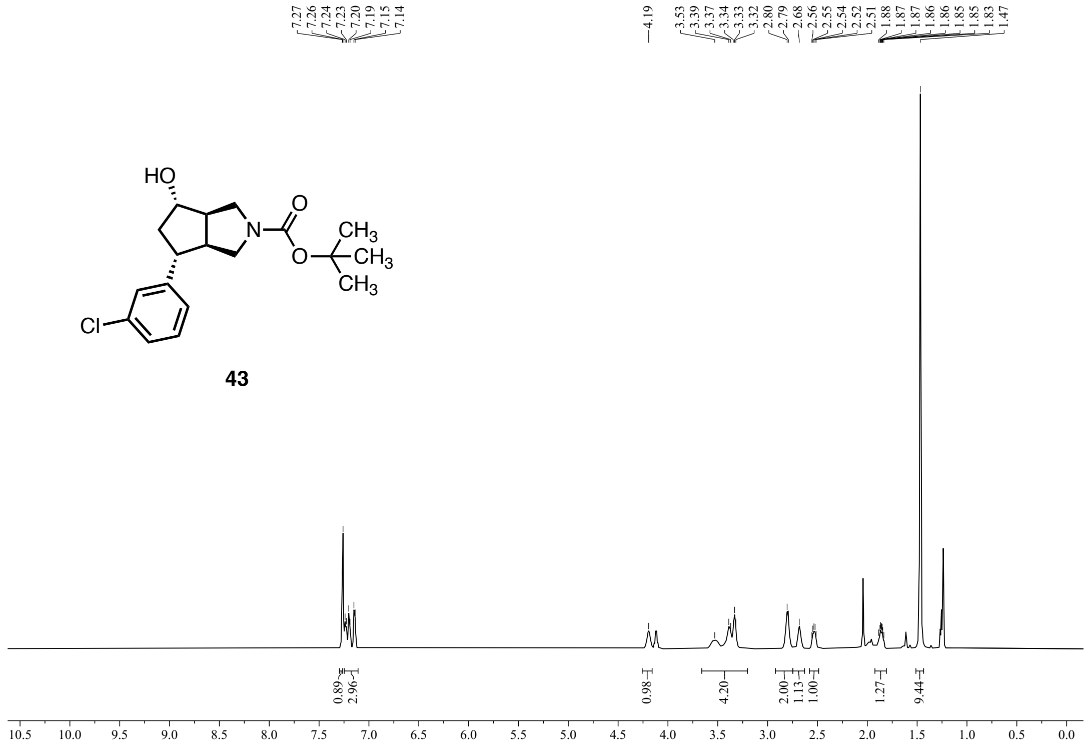


**43**

**Figure S44:** $^1$H NMR (600 MHz, $CDCl_3$) of **43** (spectra contains a small amount of EtOAc).

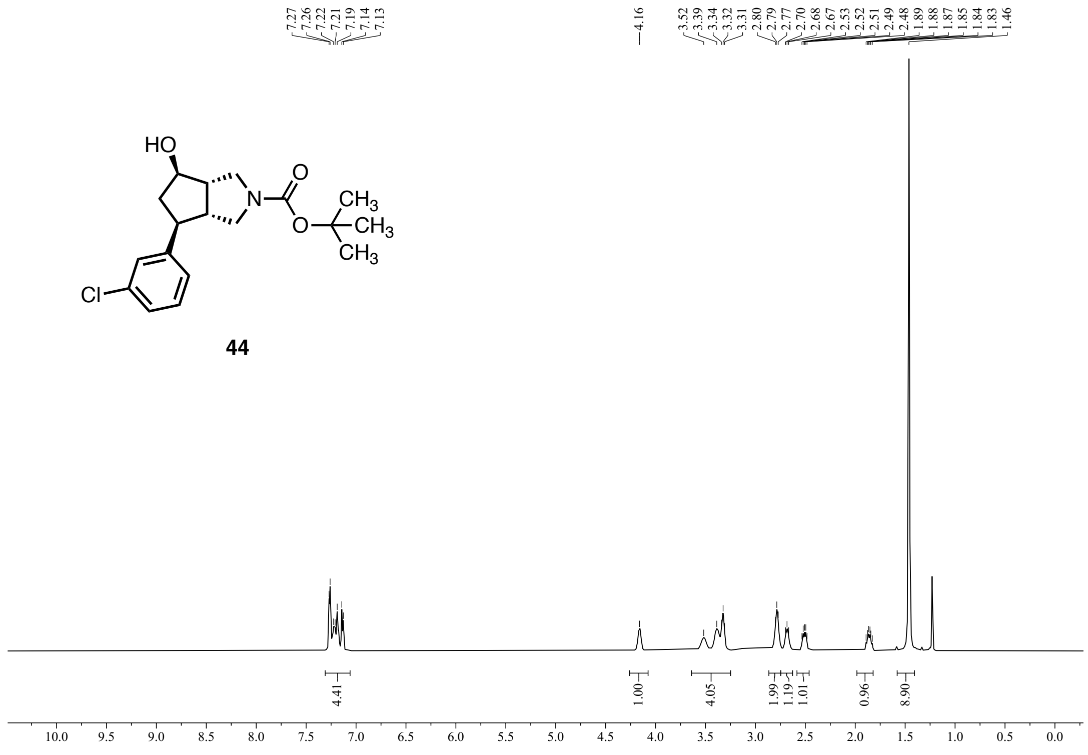


**44**

**Figure S45:** $^1$H NMR (500 MHz, $CDCl_3$) of **44**.

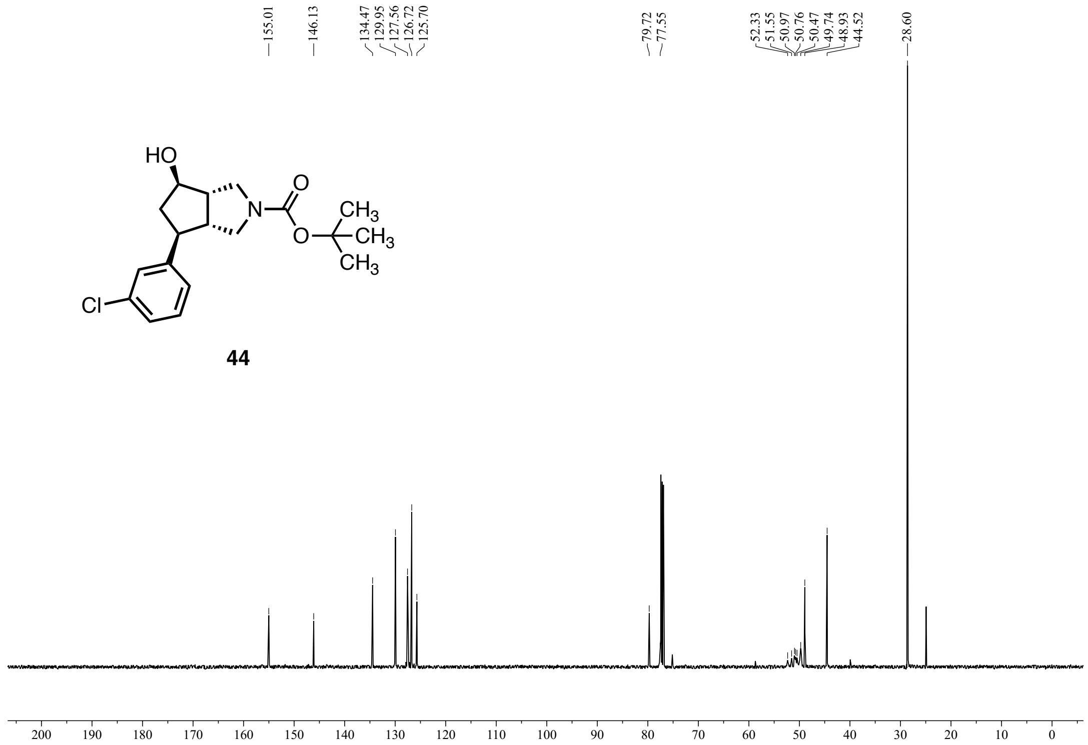


**Figure S46:** $^{13}$C NMR (126 MHz, $CDCl_3$) of **44**.

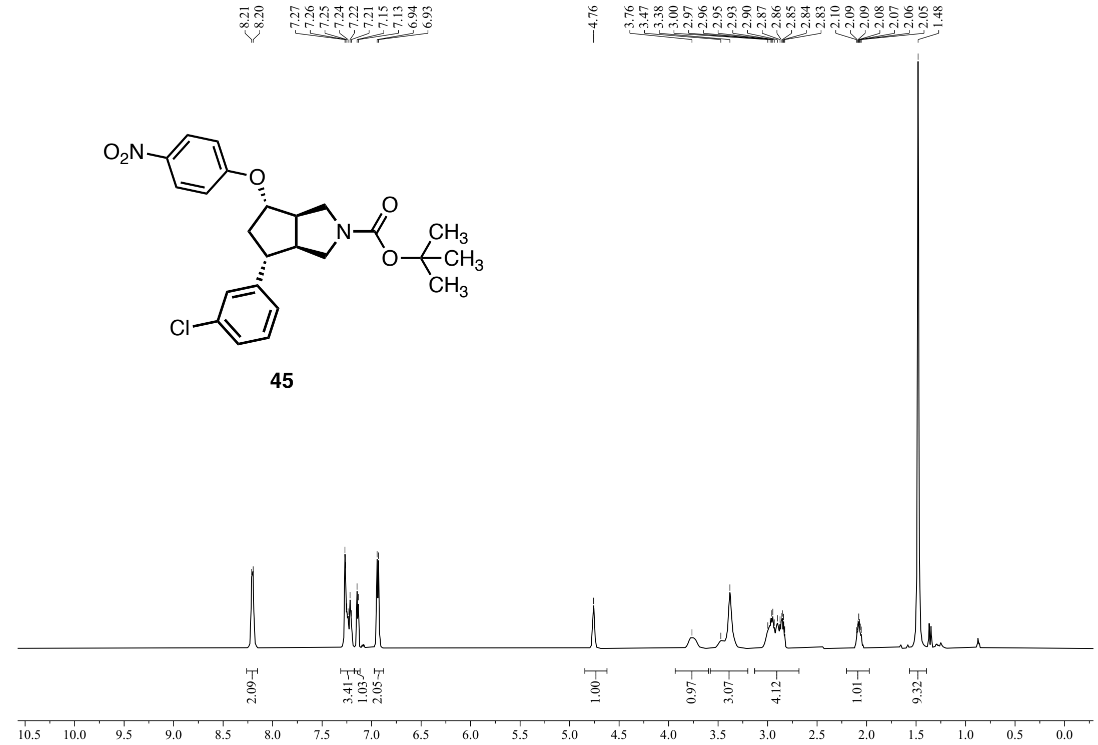


**Figure S47:** $^1$H NMR (600 MHz, $CDCl_3$) of **45**.

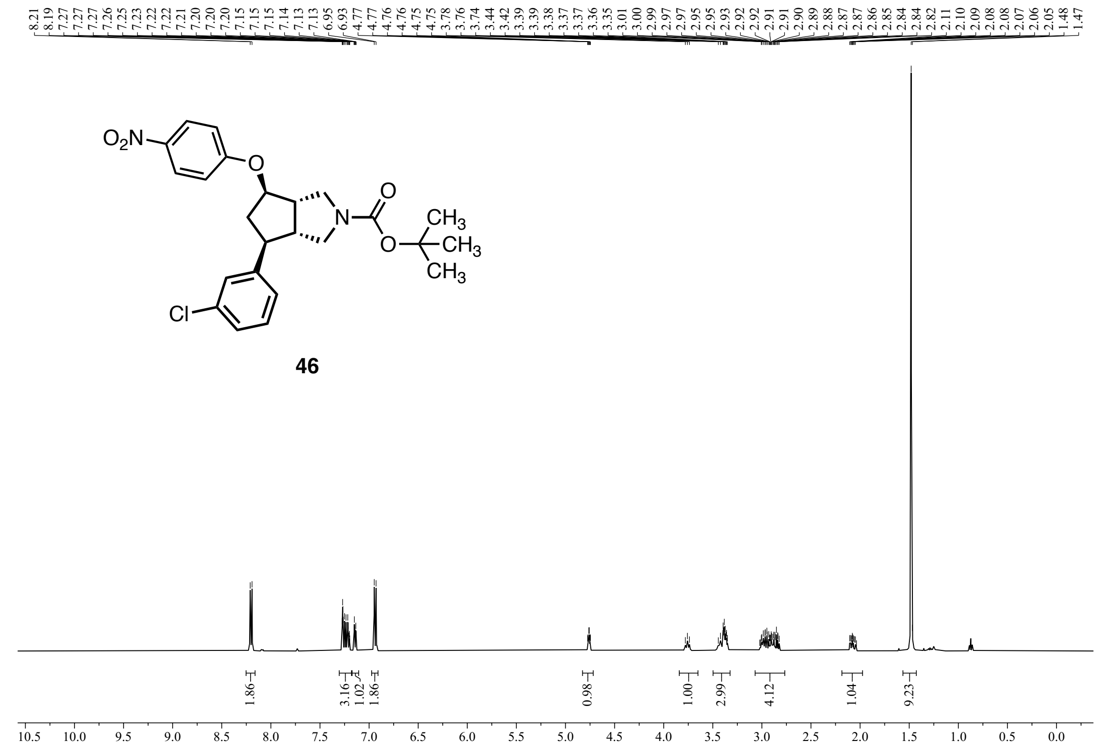


**Figure S48:** $^1$H NMR (500 MHz, $CDCl_3$) of **46**.

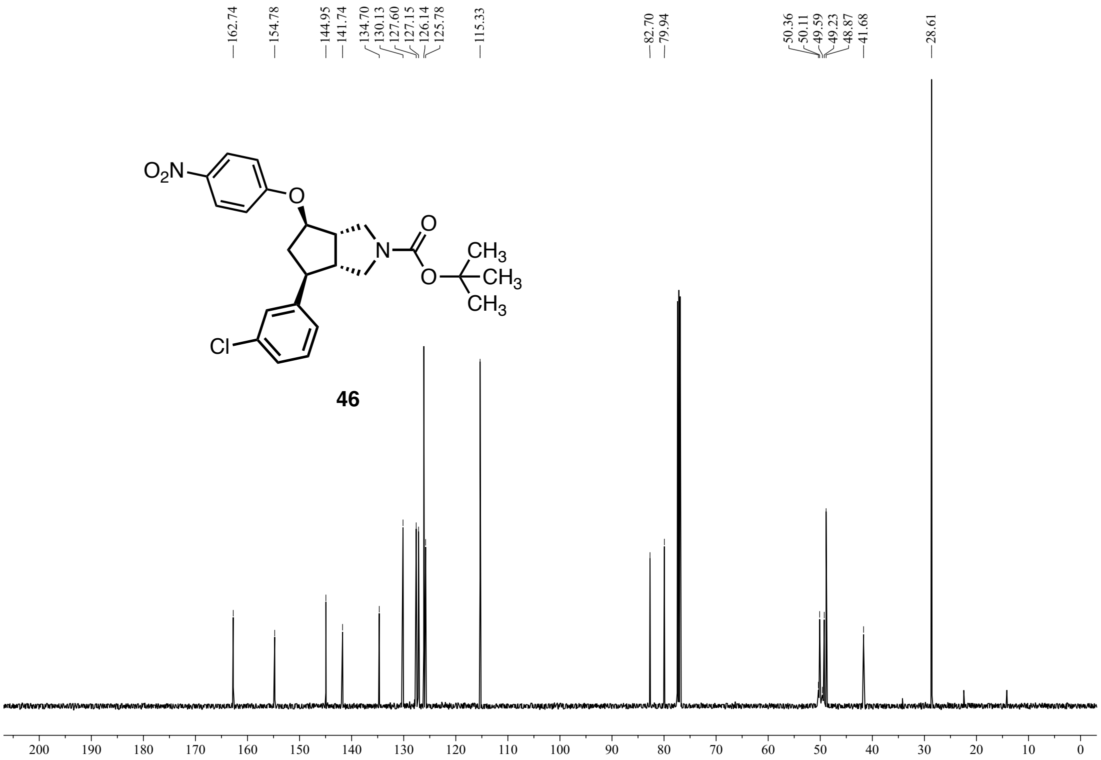


**Figure S49:** $^{13}$C NMR (126 MHz, $CDCl_3$) of **46**.

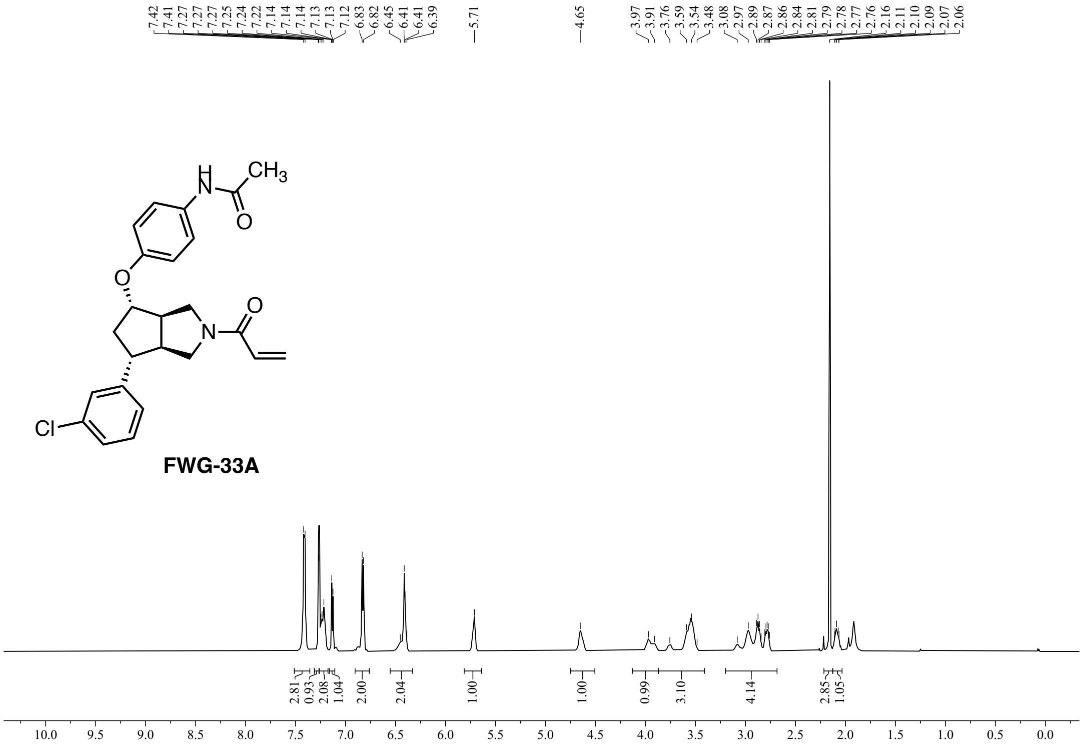


**Figure S50:** $^1$H NMR (600 MHz, $CDCl_3$) of **FWG-33A**.

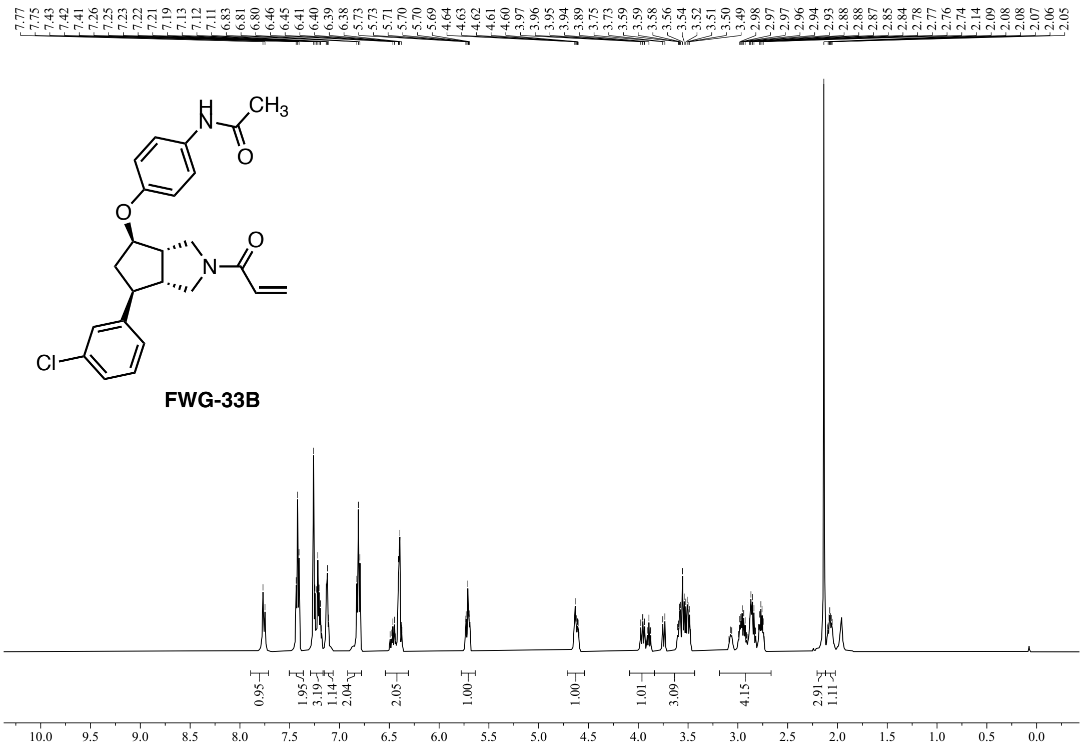


**Figure S51:** $^1$H NMR (600 MHz, $CDCl_3$) of **FWG-33B**.

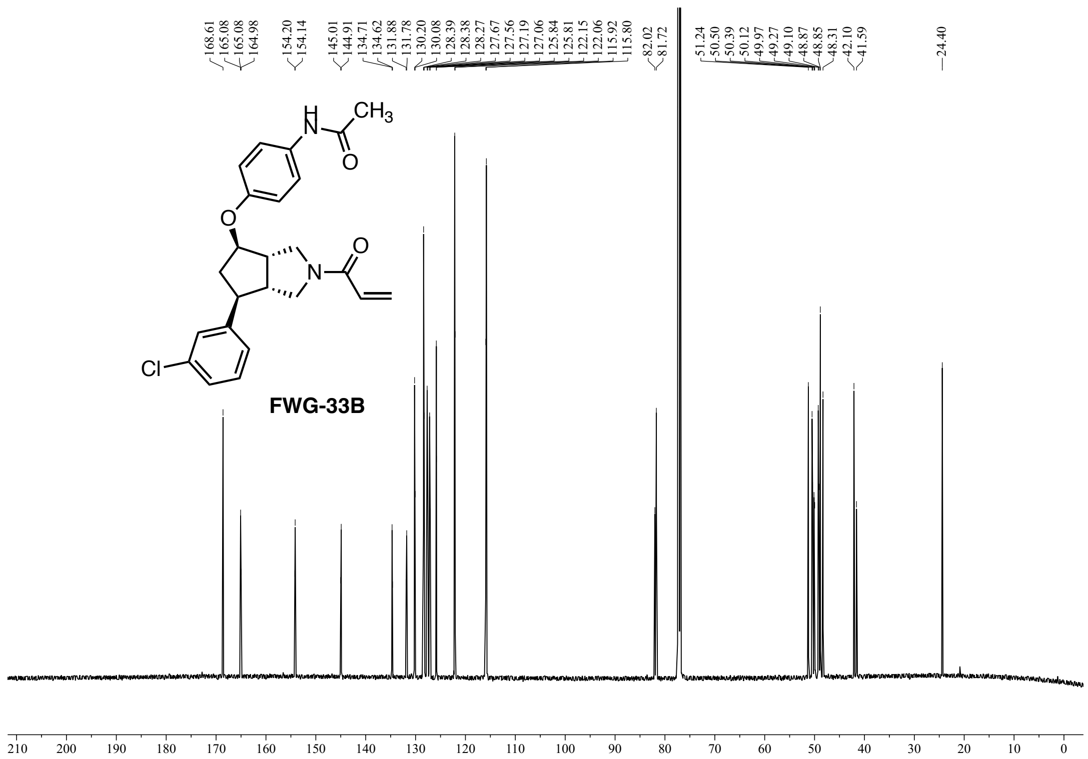


**Figure S52:** $^{13}$C NMR (151 MHz, $CDCl_3$) of **FWG-33B**.

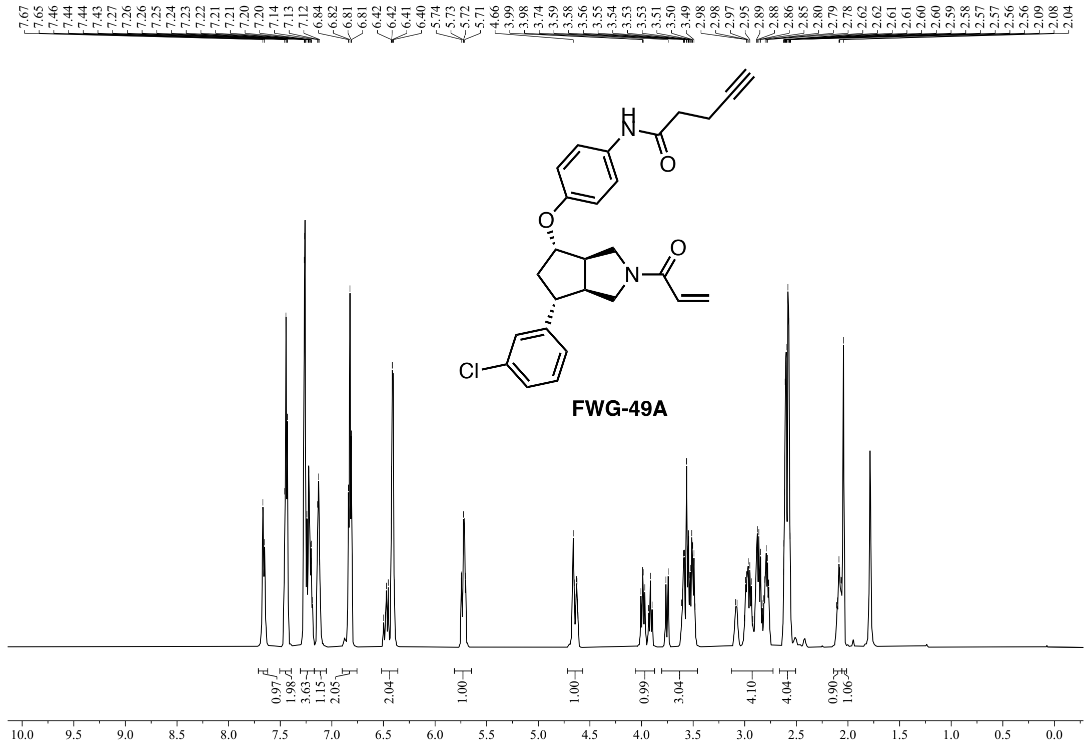


**Figure S53:** $^{1}$H NMR (600 MHz, $CDCl_3$) of **FWG-49A**.

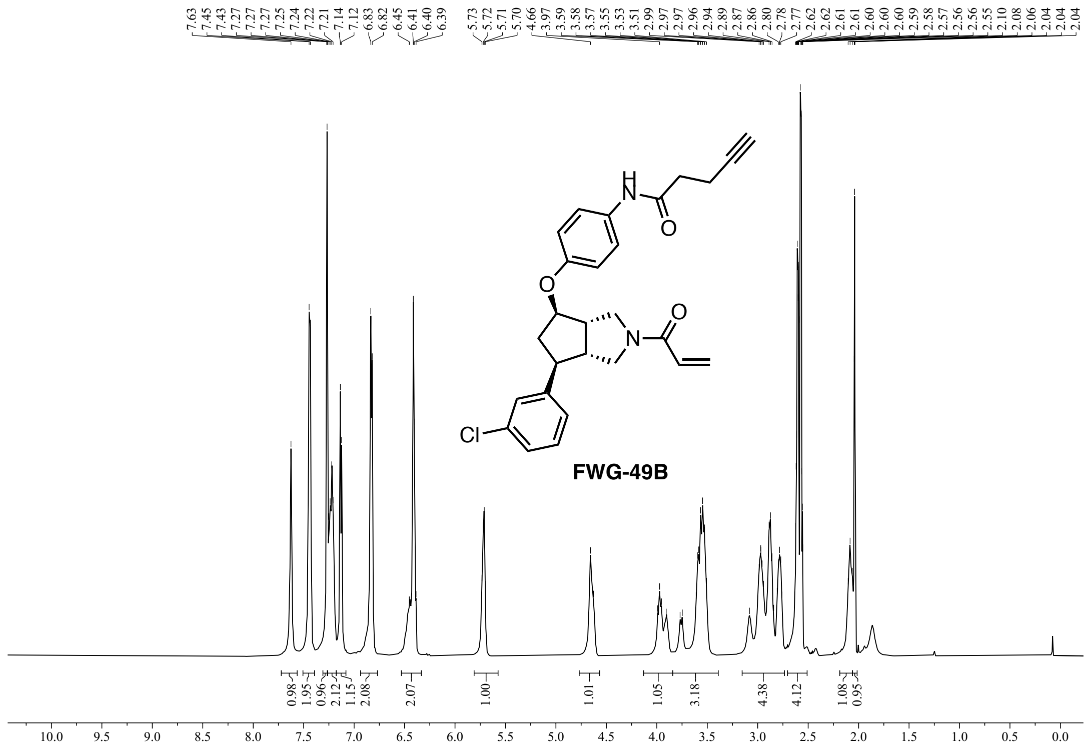


**Figure S54:** $^{1}$H NMR (600 MHz, $CDCl_3$) of **FWG-49B**.

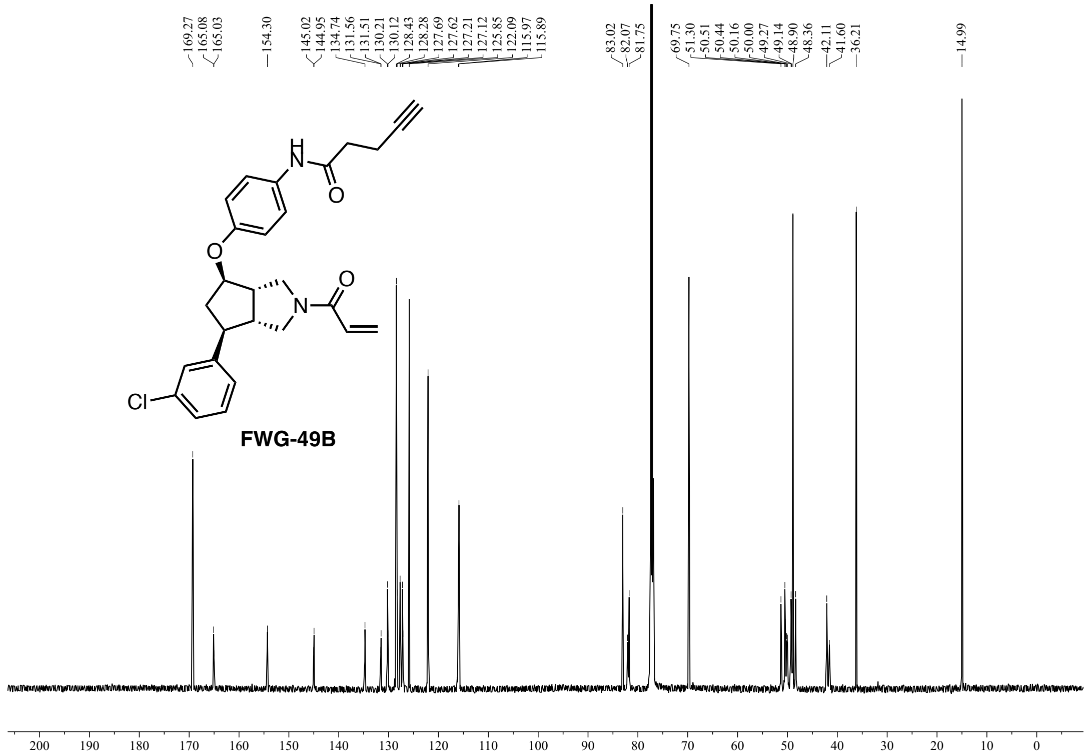

**Figure S55:** $^{13}C$ NMR (600 MHz, $CDCl_3$) of **FWG-49B**.

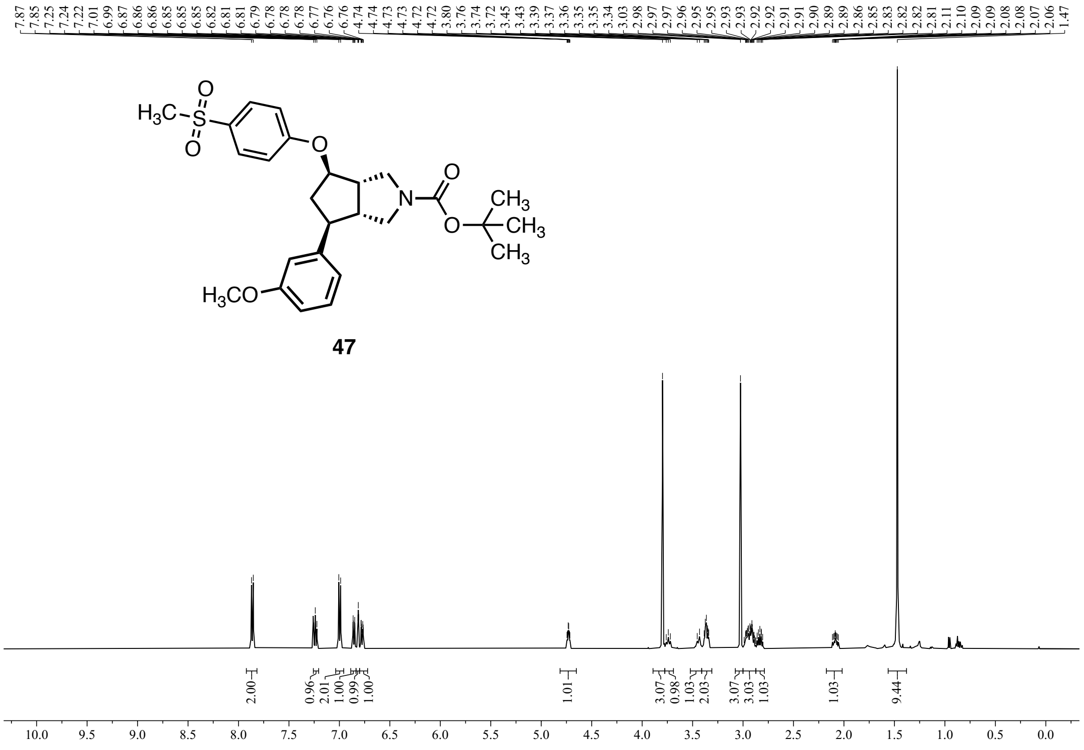


**Figure S56:** $^{1}$H NMR (500 MHz, $CDCl_3$) of **47**.

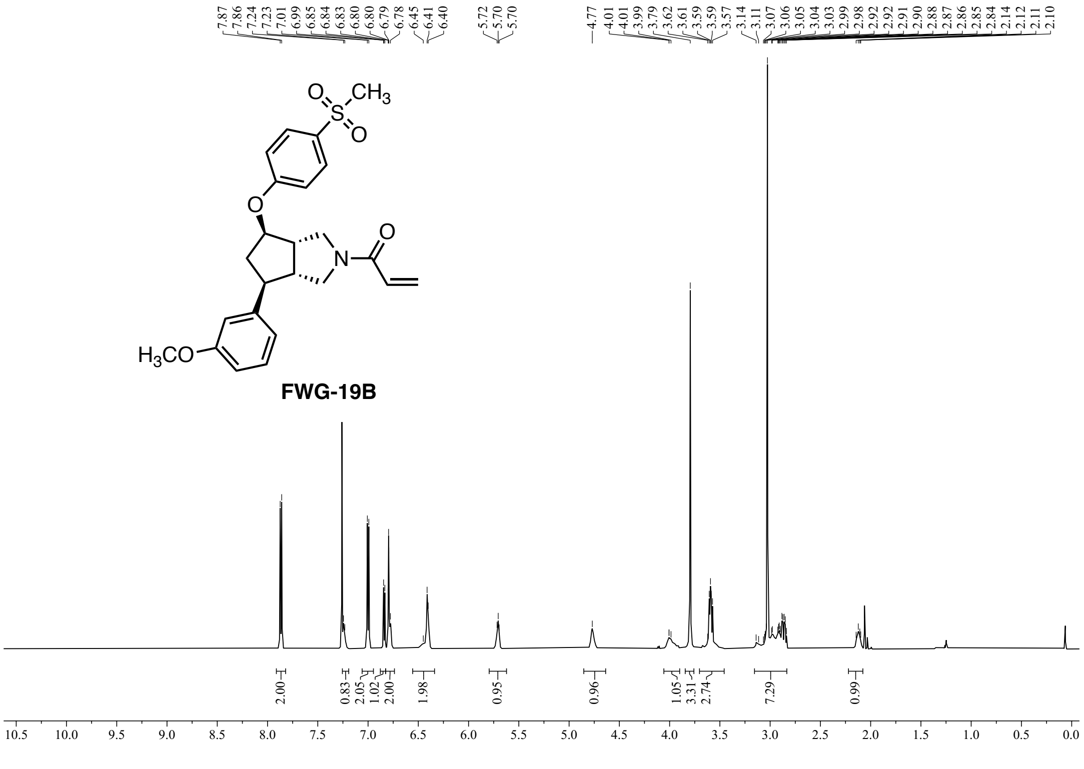


**Figure S57:** $^{1}$H NMR (500 MHz, $CDCl_3$) of **FWG-19B**.

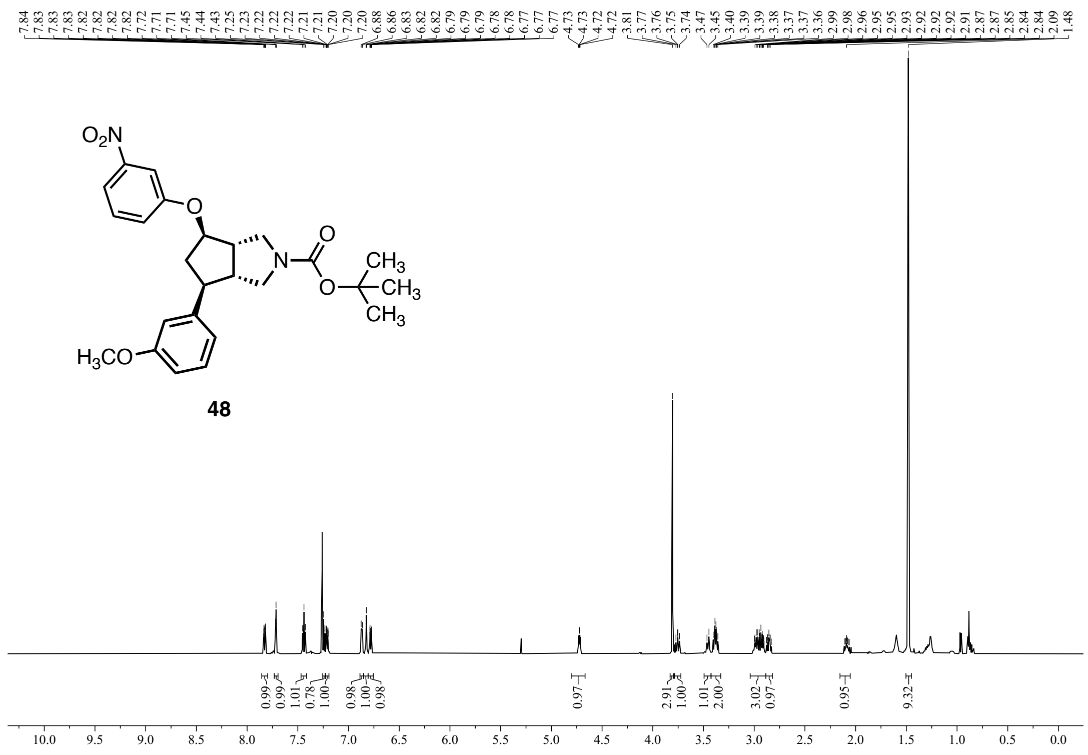

Figure S58: $^1$H NMR (600 MHz, $CDCl_3$) of **48**.

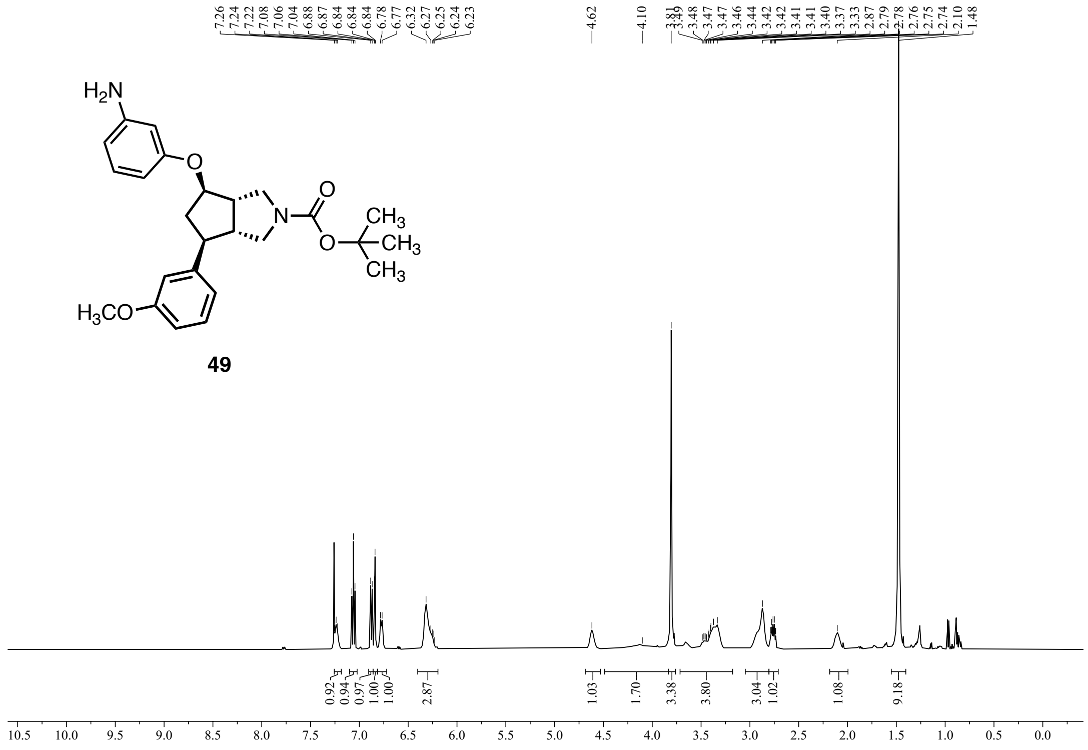

Figure S59: $^1$H NMR (500 MHz, $CDCl_3$) of **49**.

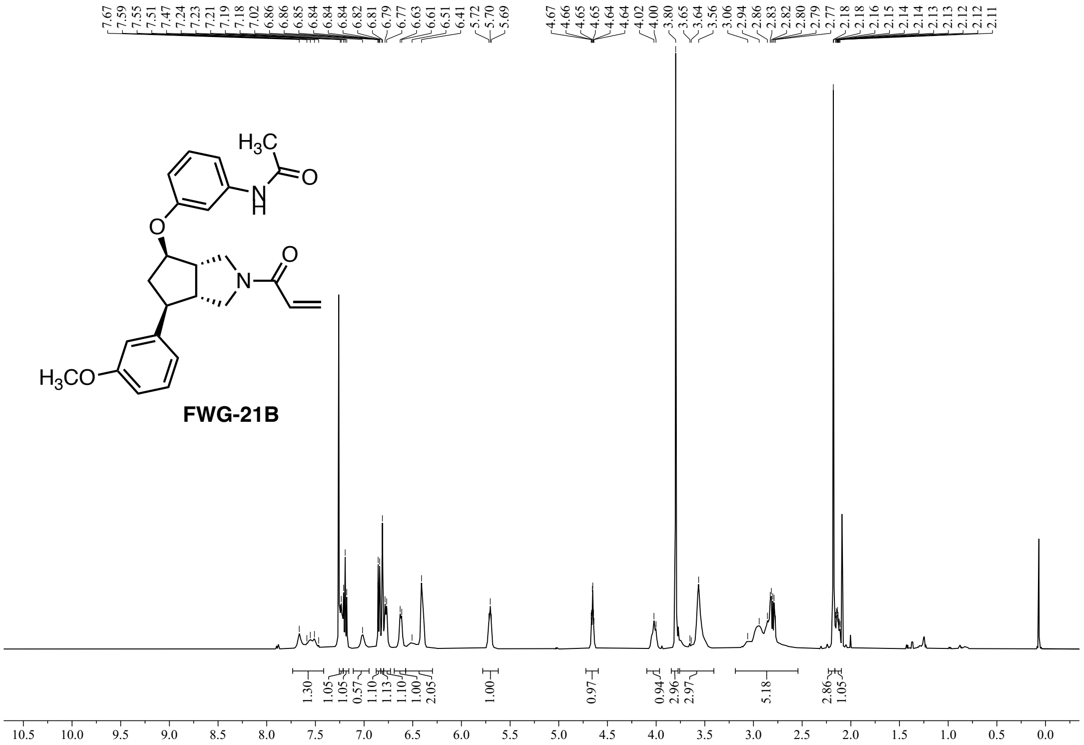


**Figure S60:** $^1$H NMR (500 MHz, $CDCl_3$) of **FWG-21B**.

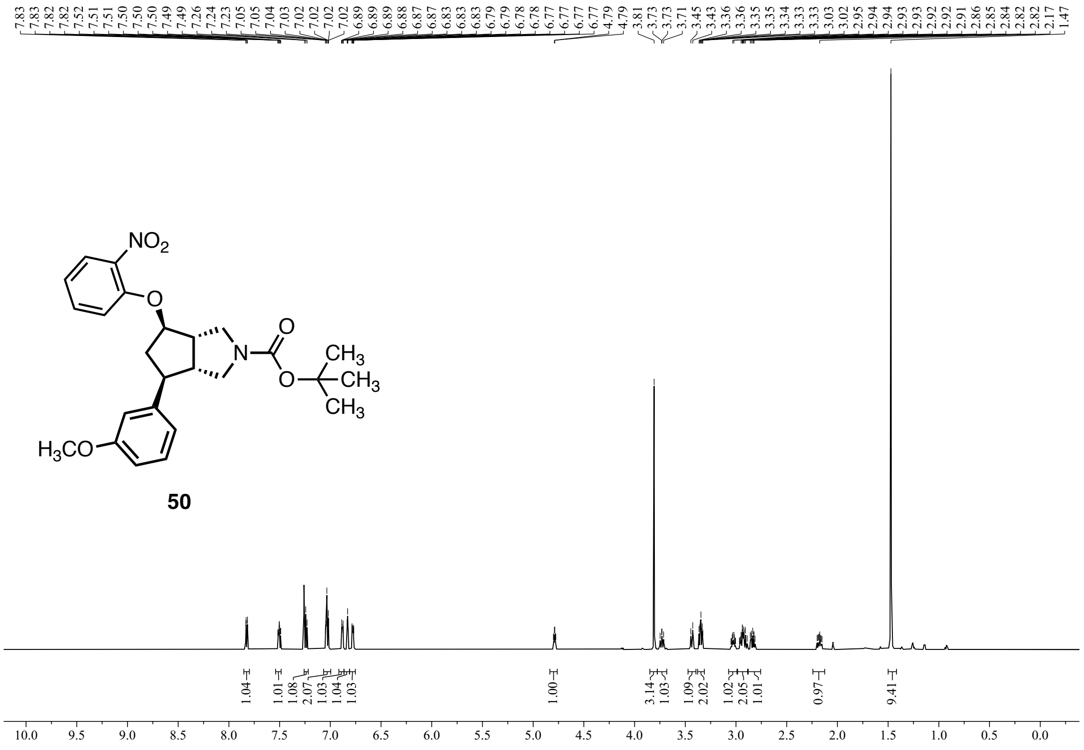


**Figure S61:** $^1$H NMR (600 MHz, $CDCl_3$) of **50**.

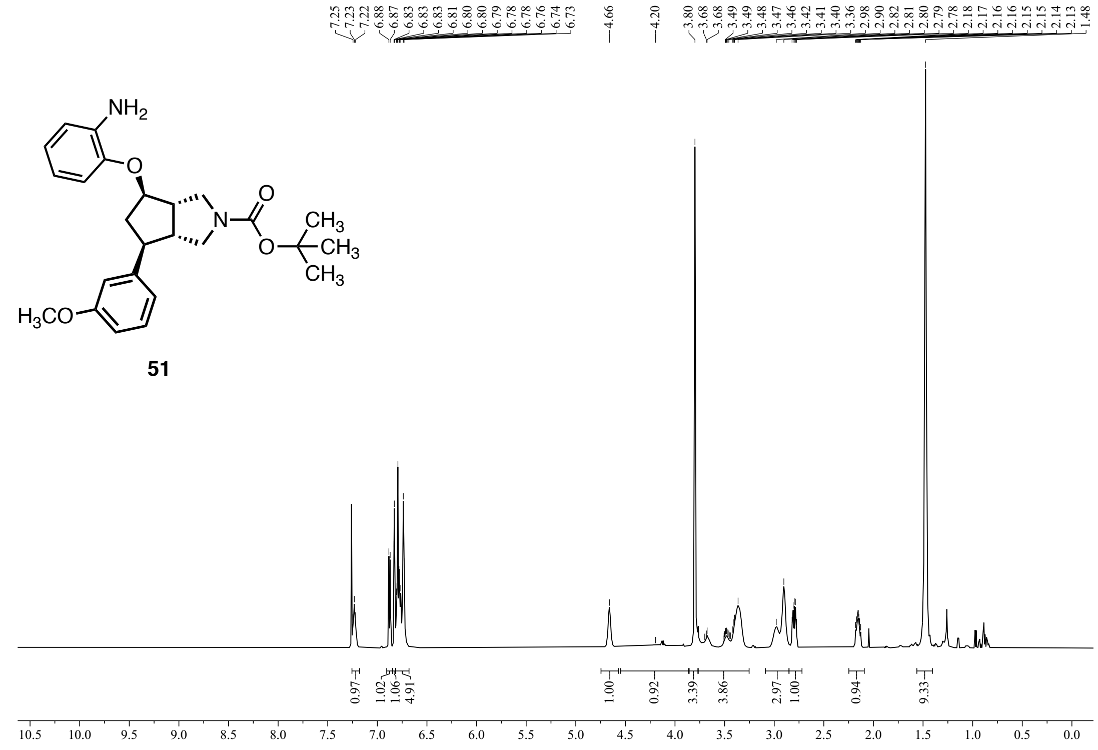

51

**Figure S62:** $^1$H NMR (600 MHz, $CDCl_3$) of **51**.

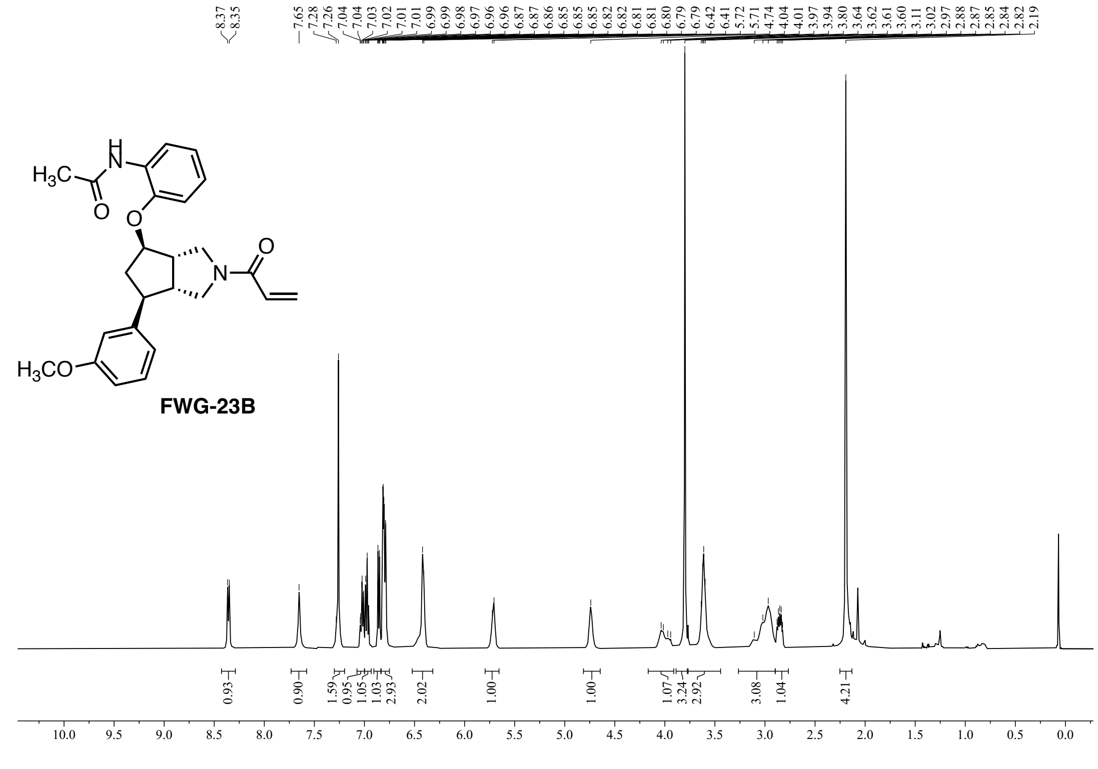

**Figure S63:** $^1$H NMR (500 MHz, $CDCl_3$) of **FWG-23B**.

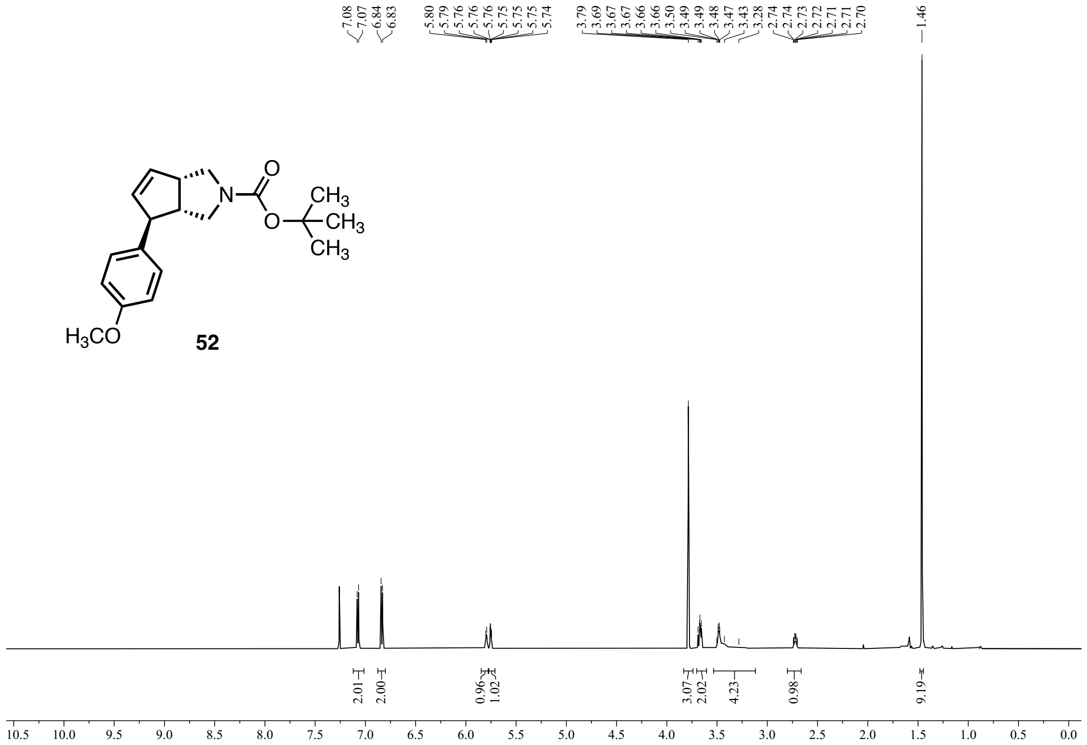


**Figure S64:** $^{1}$H NMR (600 MHz, $CDCl_3$) of **52**.

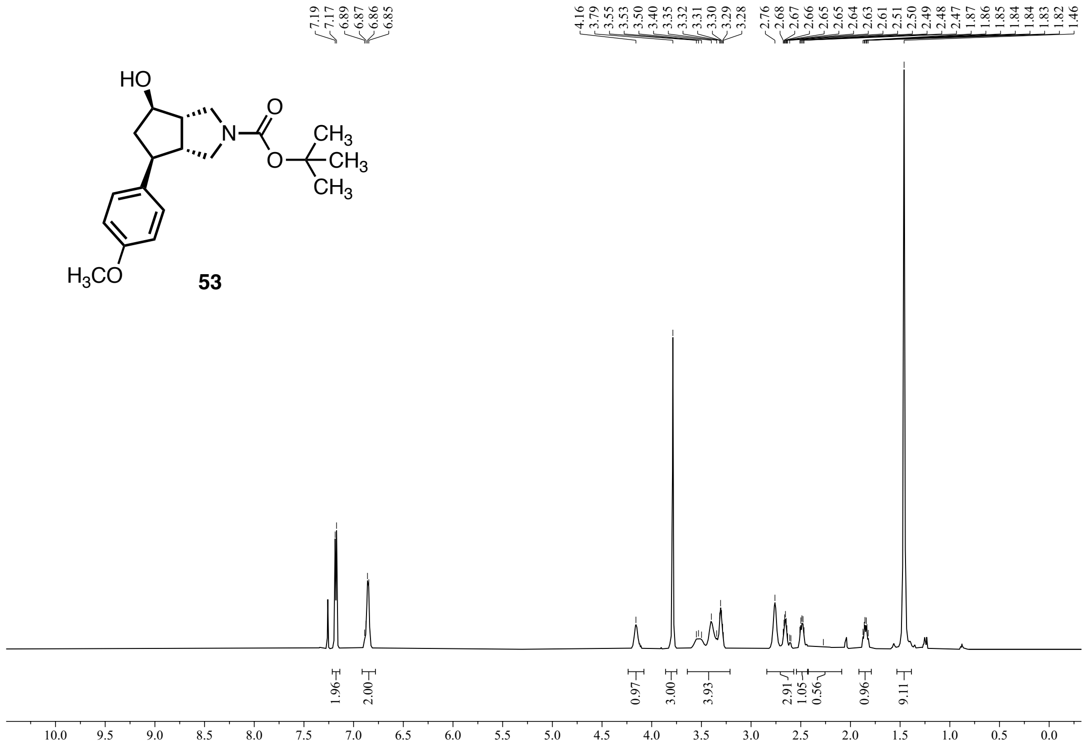


**Figure S65:** $^{1}$H NMR (600 MHz, $CDCl_3$) of **53**.

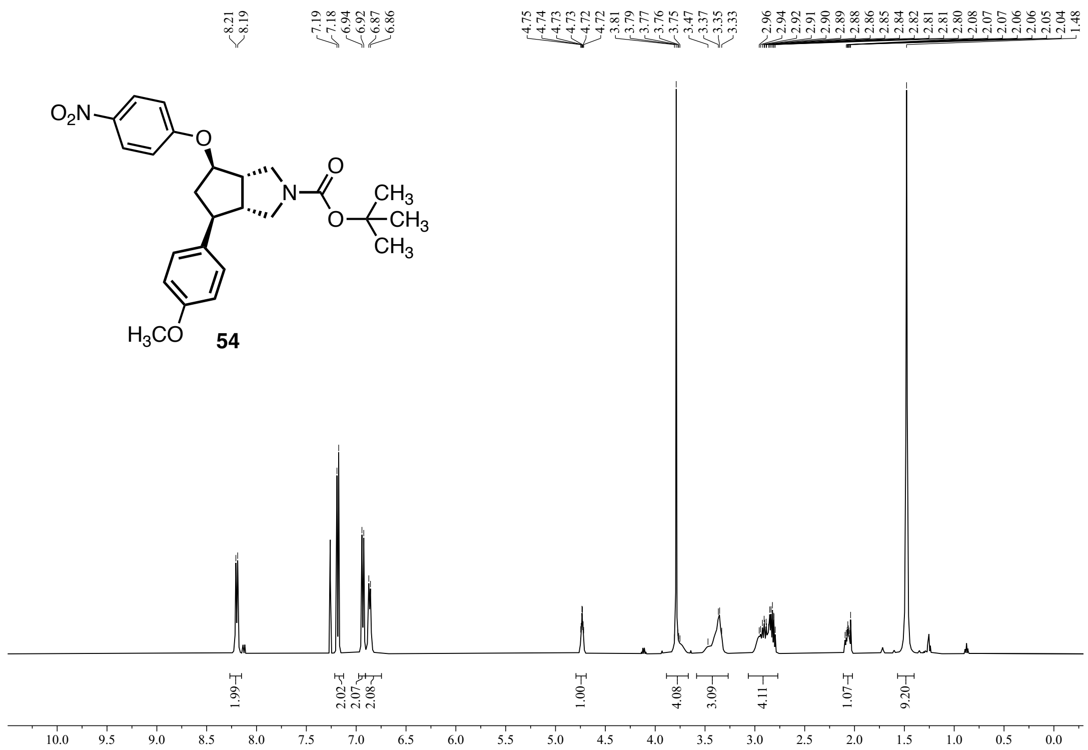

**Figure S66:** $^{1}$H NMR (500 MHz, $CDCl_3$) of **54**.

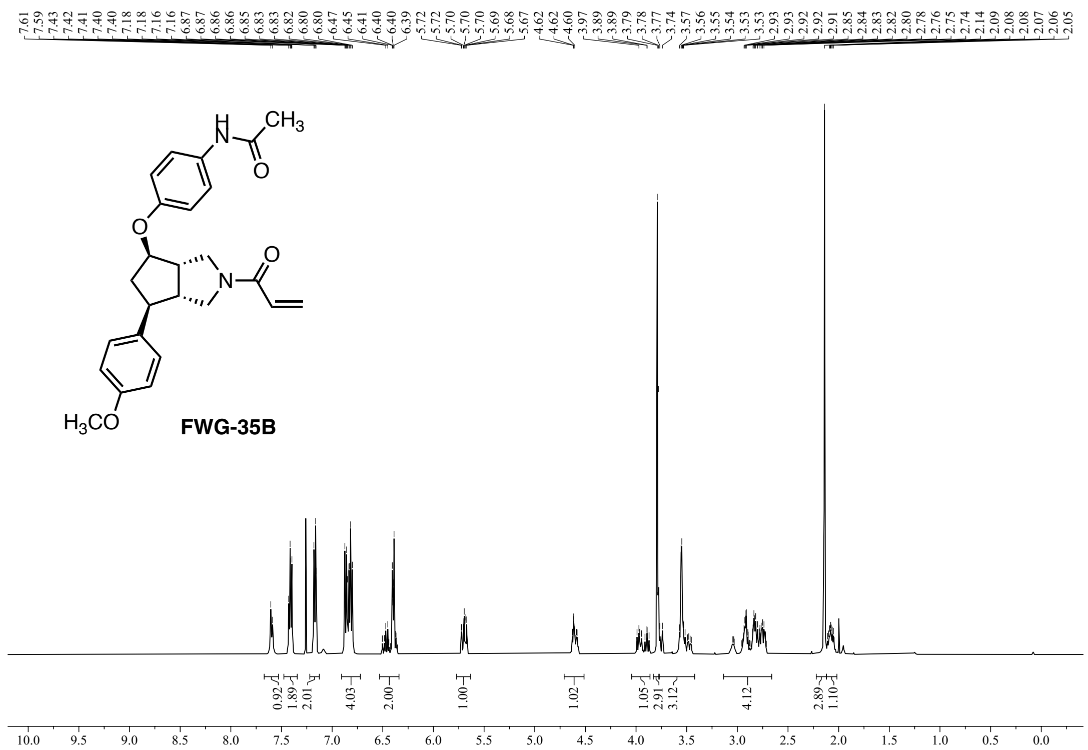

**Figure S67:** $^{1}$H NMR (500 MHz, $CDCl_3$) of **FWG-35B**.

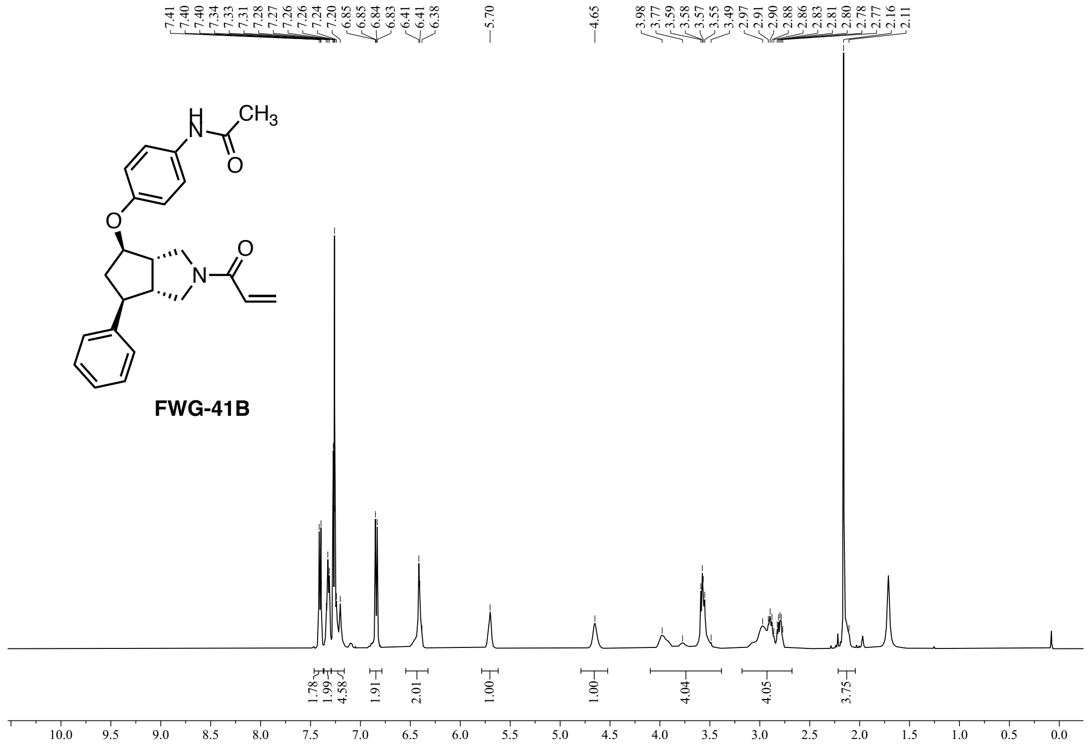


**Figure S68:** $^1$H NMR (500 MHz, $CDCl_3$) of **FWG-41B**.

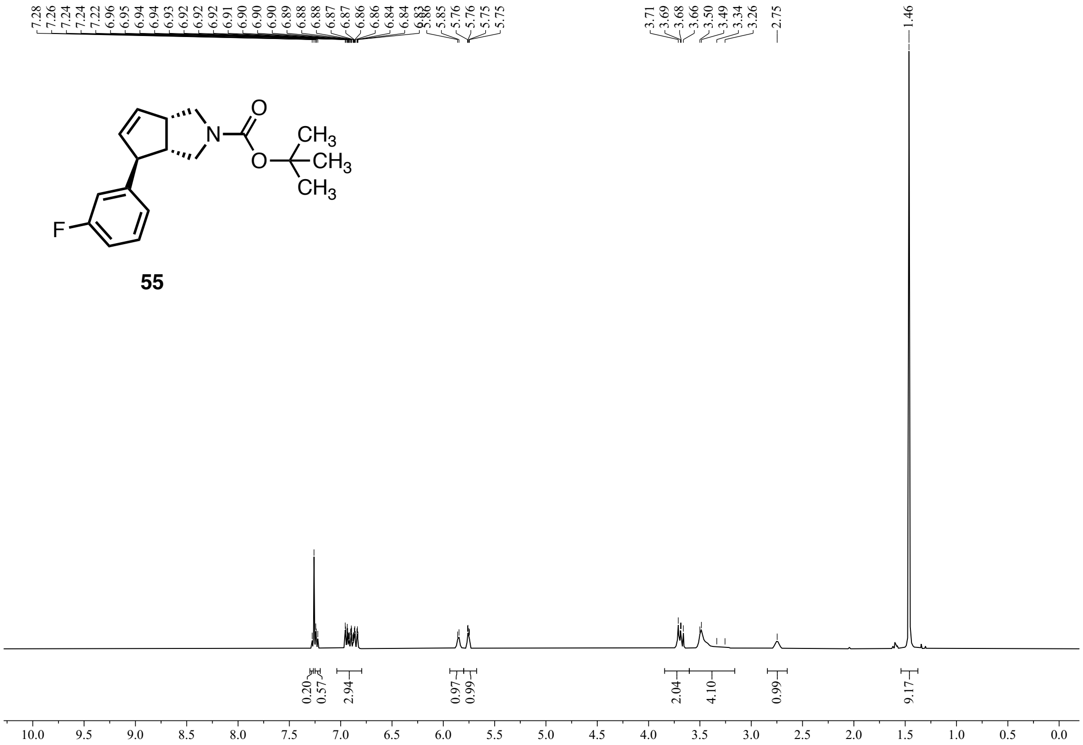

**Figure S69:** $^{1}H$ NMR (400 MHz, $CDCl_3$) of **55**.

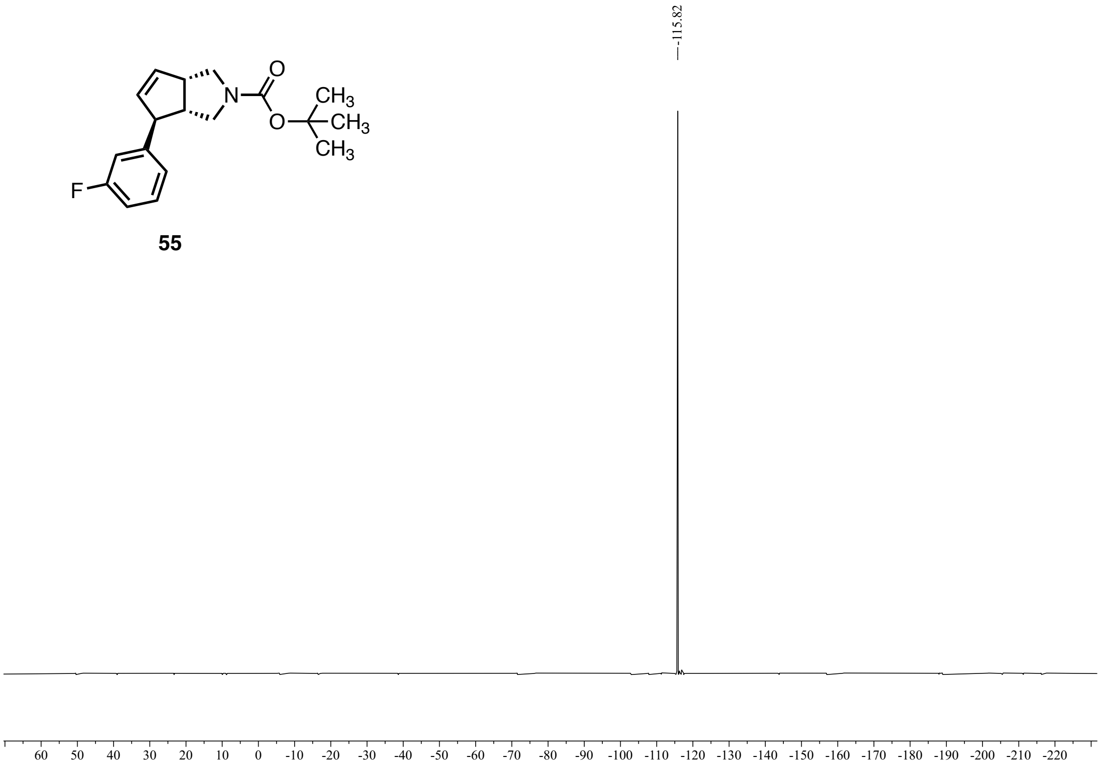

**Figure S70:** $^{19}F$ NMR {$^{1}H$} (376 MHz, $CDCl_3$) of **55**.

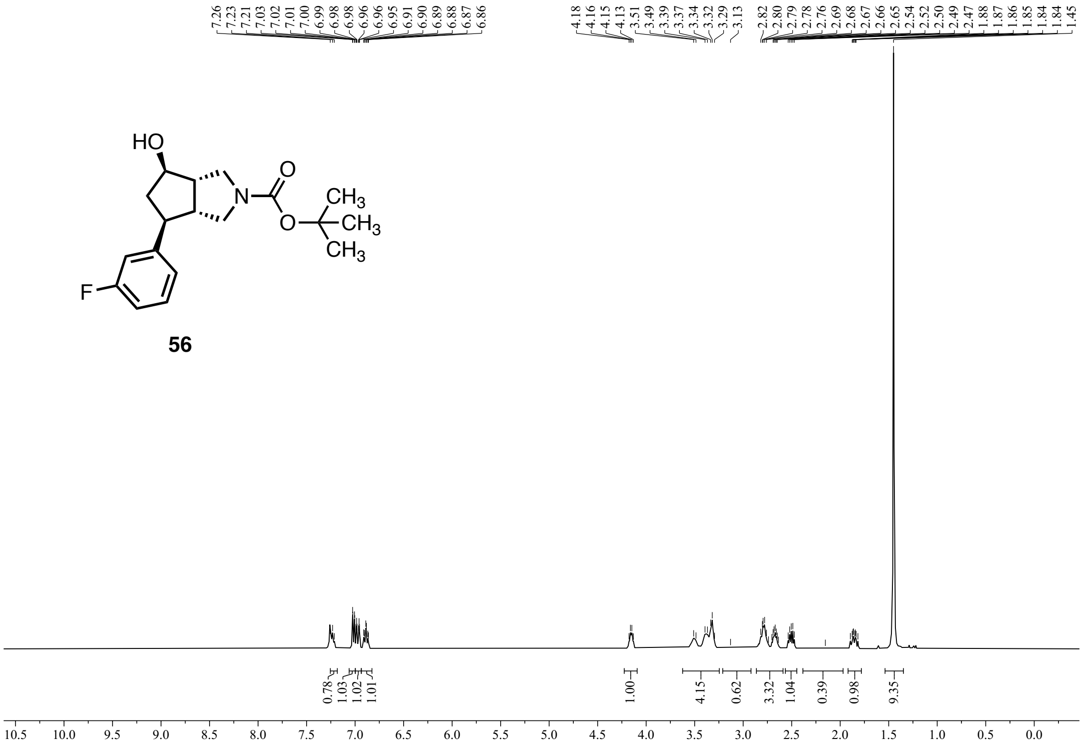

**Figure S71:** $^{1}H$ NMR (400 MHz, $CDCl_3$) of **56**.

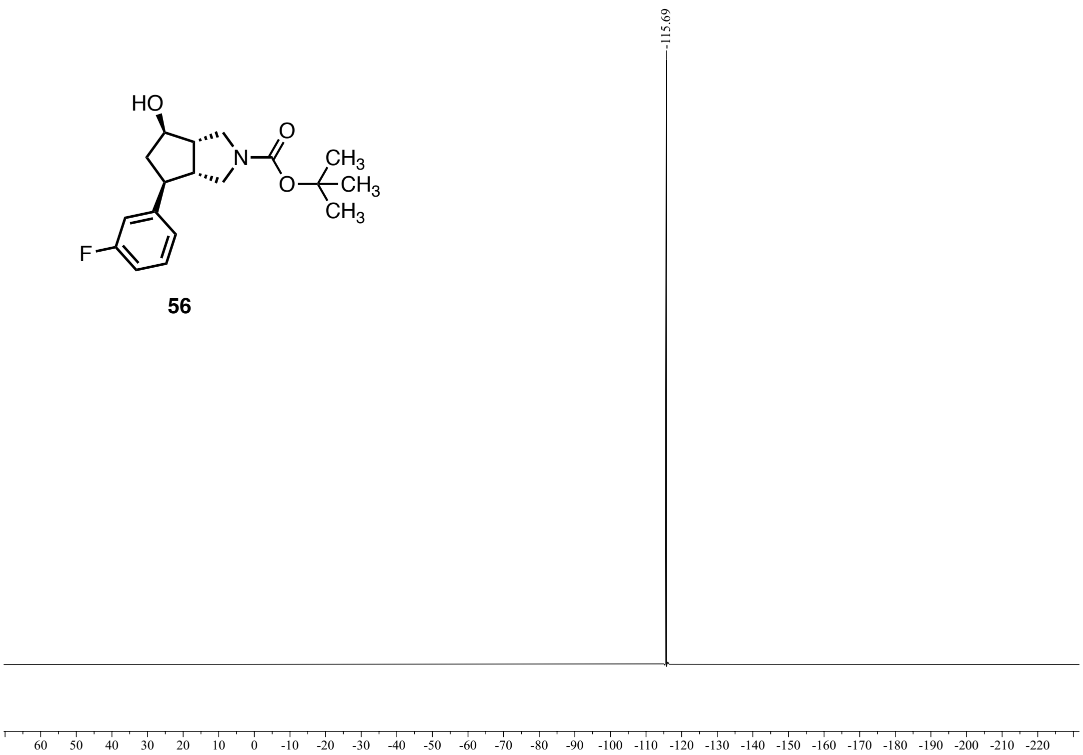

**Figure S72:** $^{19}F$ NMR {$^{1}H$} (376 MHz, $CDCl_3$) of **56**.

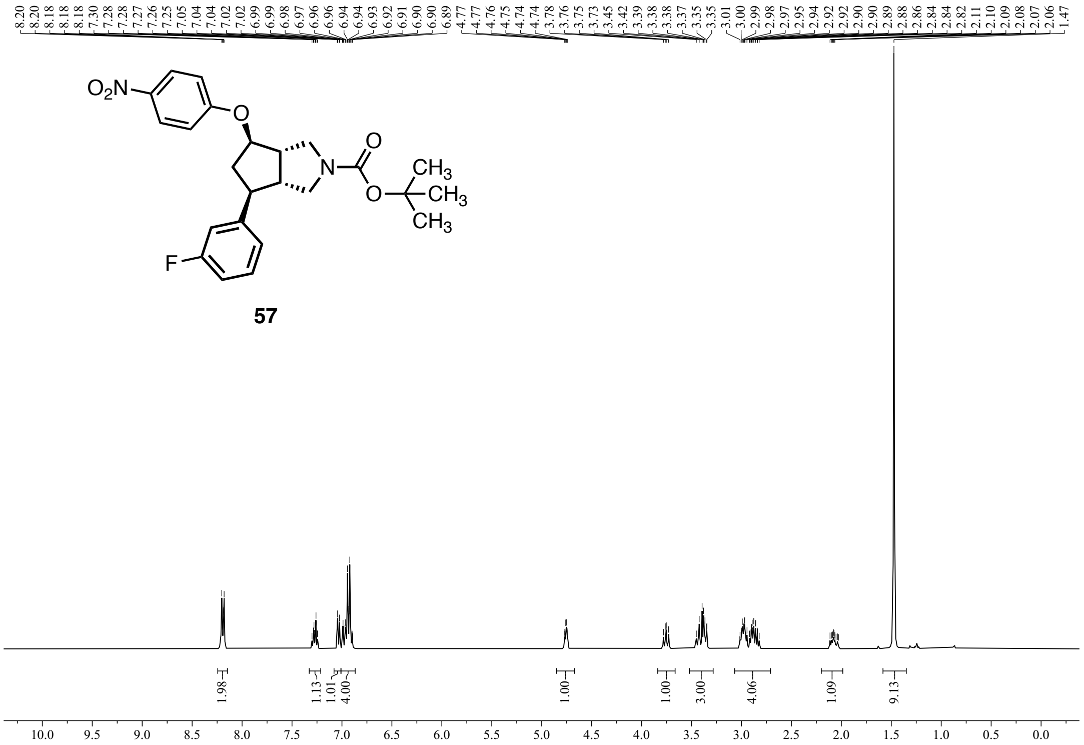


**Figure S73:** $^1H$ NMR (400 MHz, $CDCl_3$) of **57**.

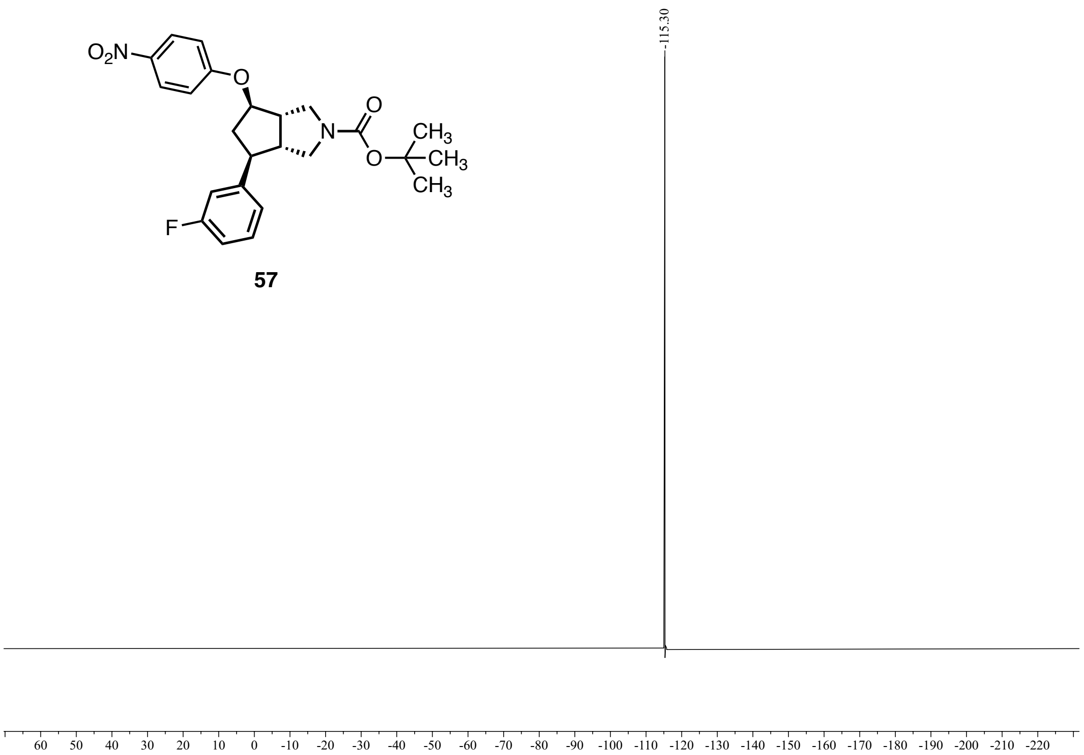


**Figure S74:** $^{19}F$ NMR {$^1H$} (376 MHz, $CDCl_3$) of **57**.

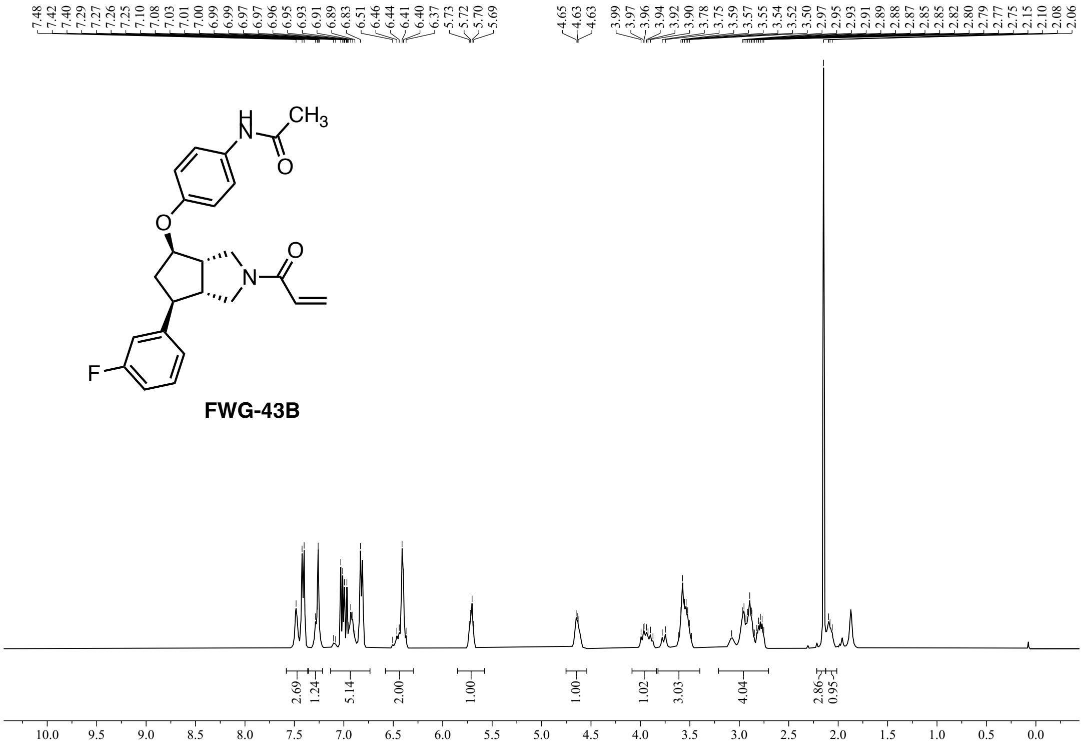


**Figure S75:** $^{1}H$ NMR (400 MHz, $CDCl_3$) of **FWG-43B**.

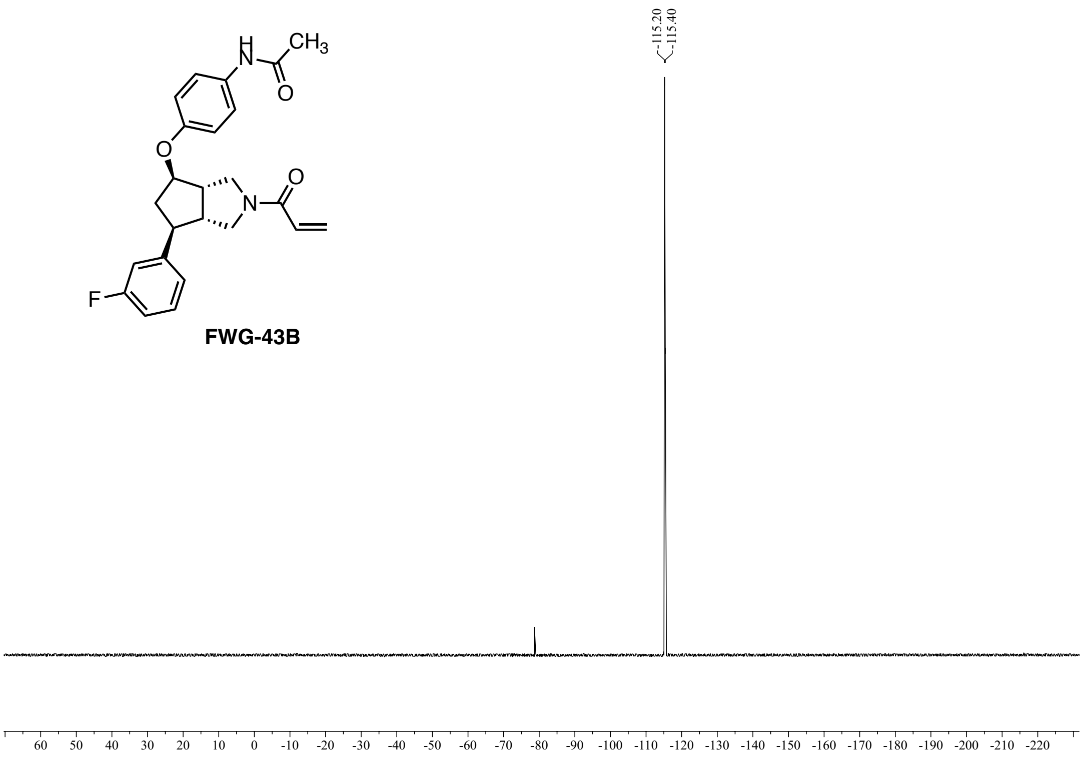


**Figure S76:** $^{19}F$ NMR {$^{1}H$} (376 MHz, $CDCl_3$) of **FWG-43B**.

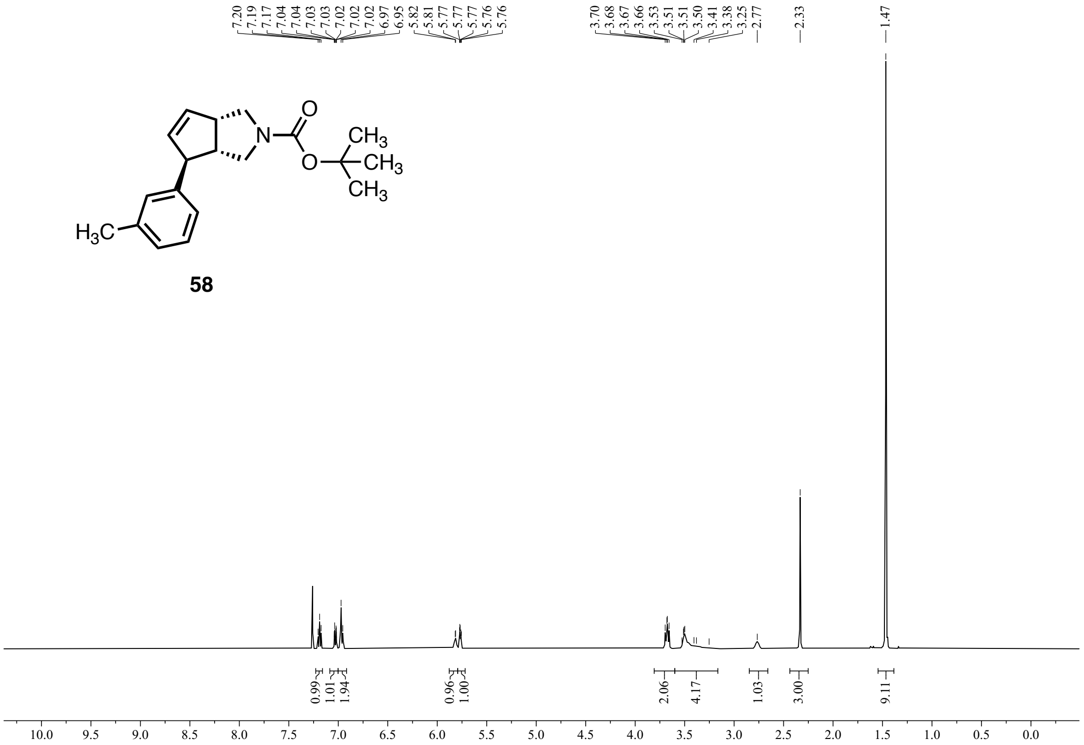


**Figure S77:** $^{1}H$ NMR (500 MHz, $CDCl_3$) of **58**.

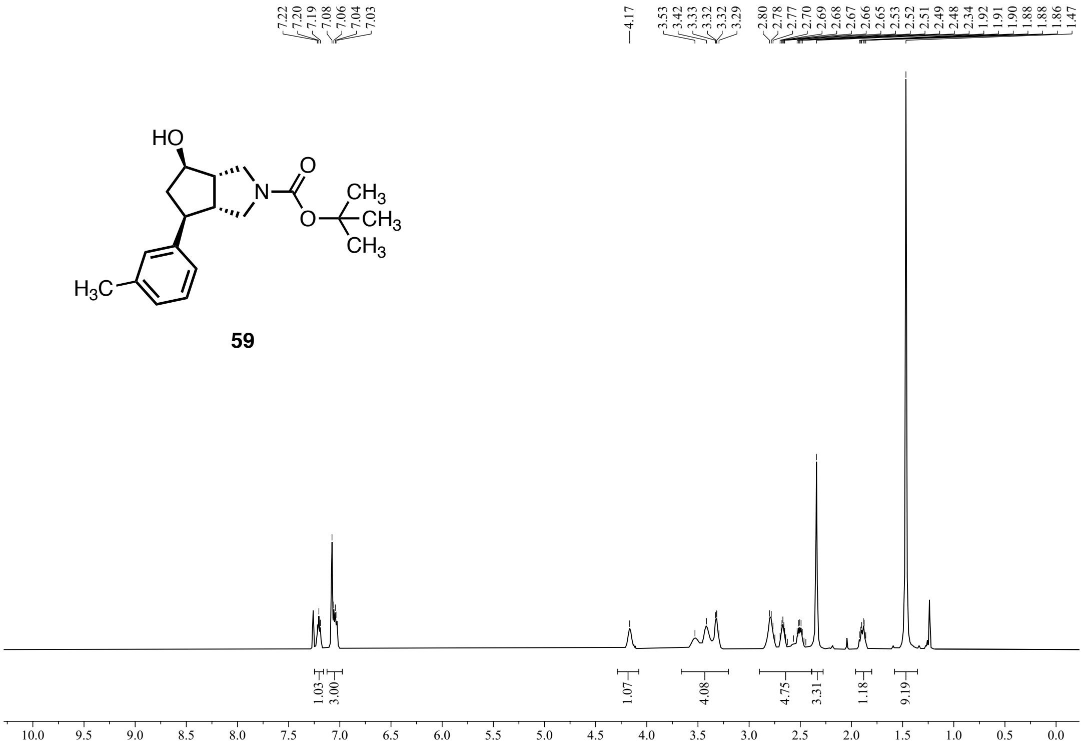


**Figure S78:** $^{1}H$ NMR (500 MHz, $CDCl_3$) of **59**.

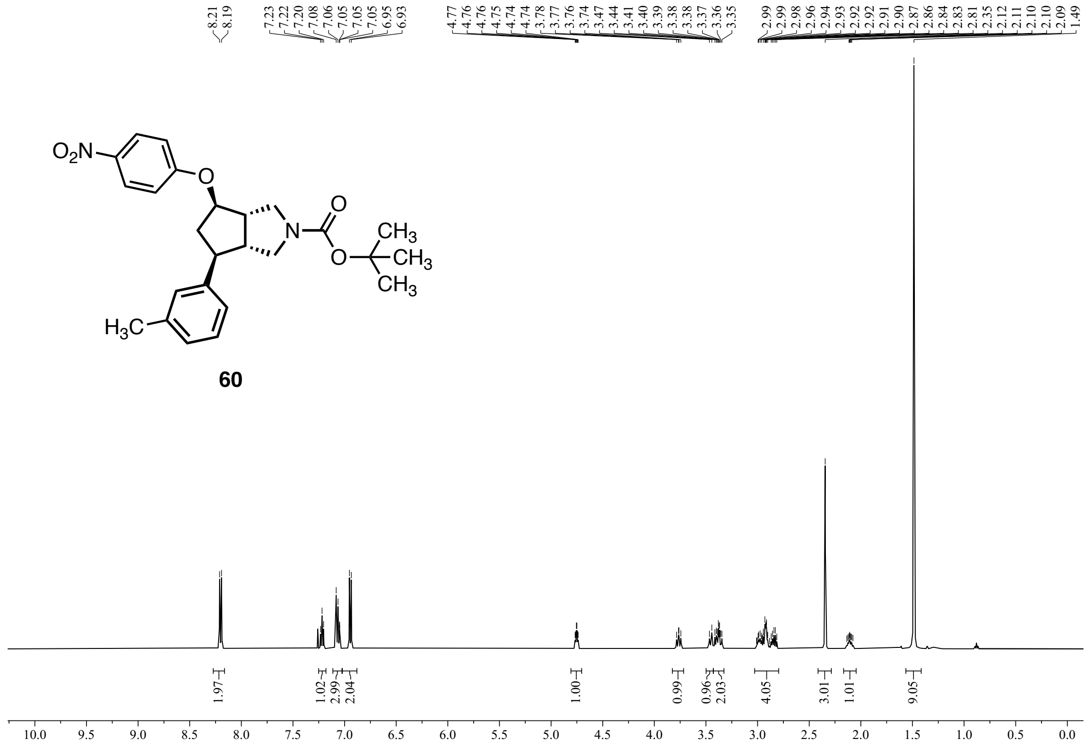

Figure S79: $^{1}H$ NMR (500 MHz, $CDCl_3$) of **60**.

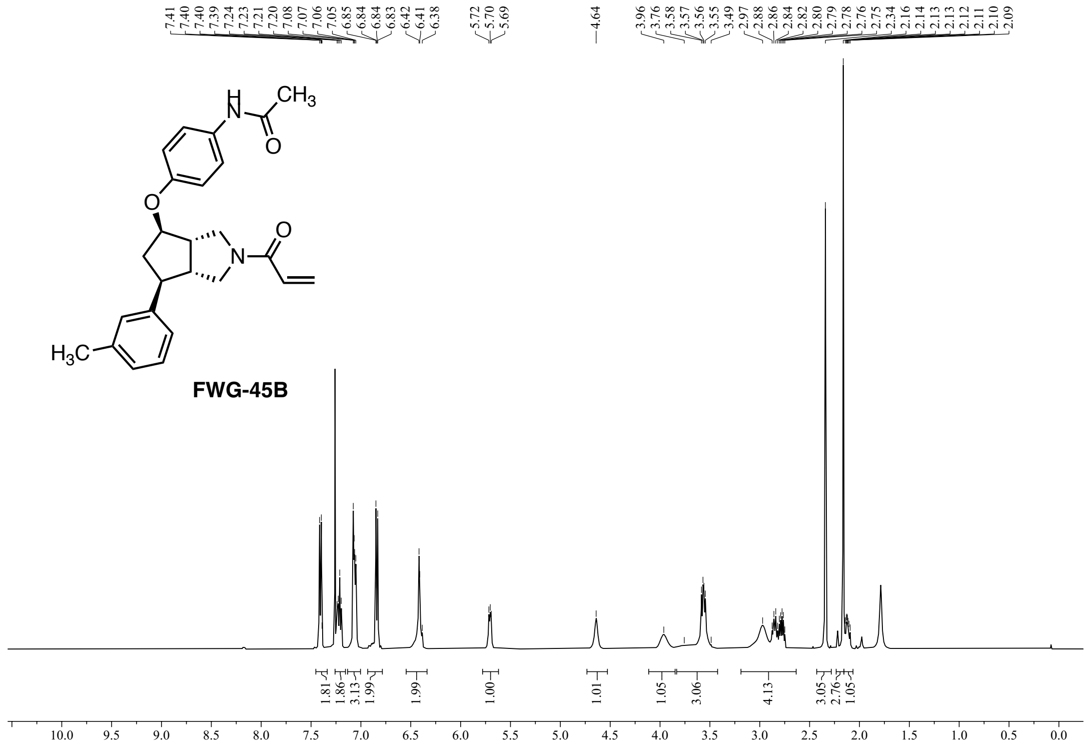

Figure S80: $^{1}H$ NMR (500 MHz, $CDCl_3$) of **FWG-45B**.

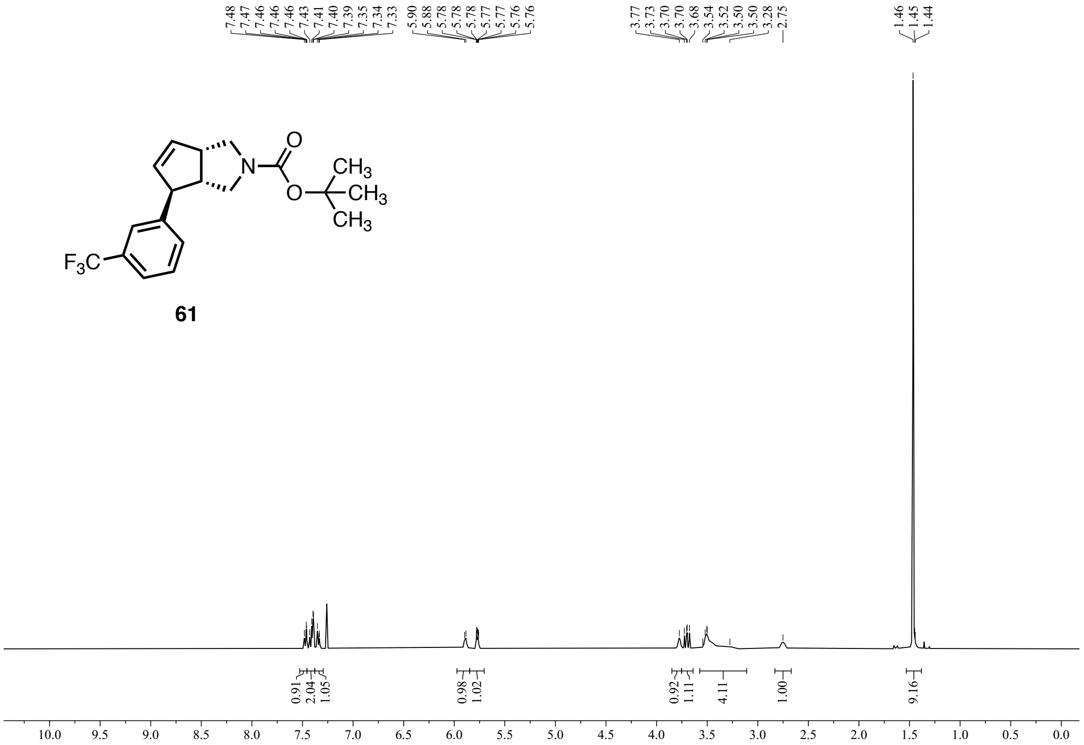


**Figure S81:** $^{1}H$ NMR (400 MHz, $CDCl_3$) of **61**.

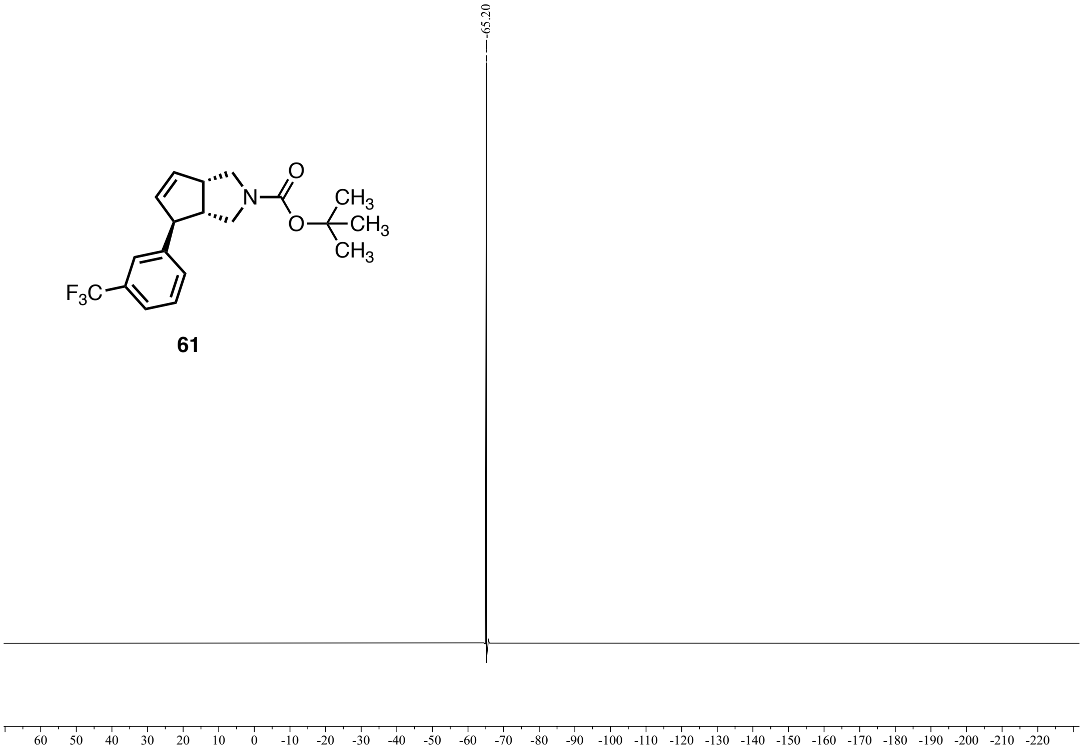


**Figure S82:** $^{19}F$ NMR {$^{1}H$} (376 MHz, $CDCl_3$) of **61**.

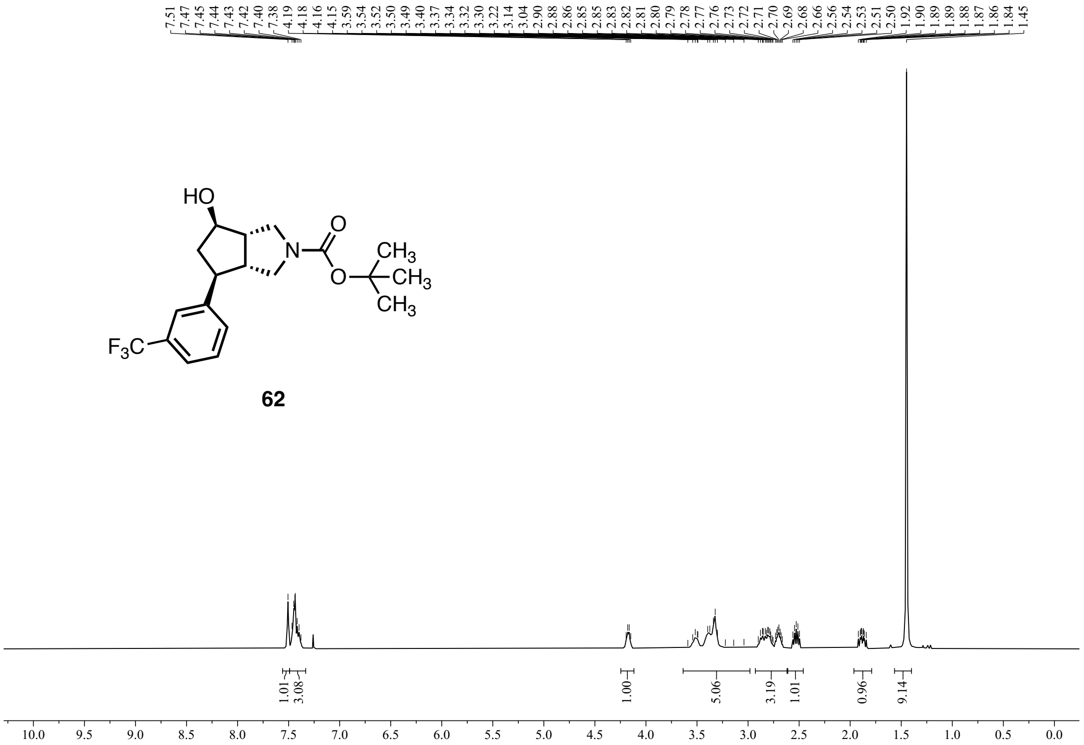


**Figure S83:** $^{1}$H NMR (400 MHz, $CDCl_3$) of **62**.

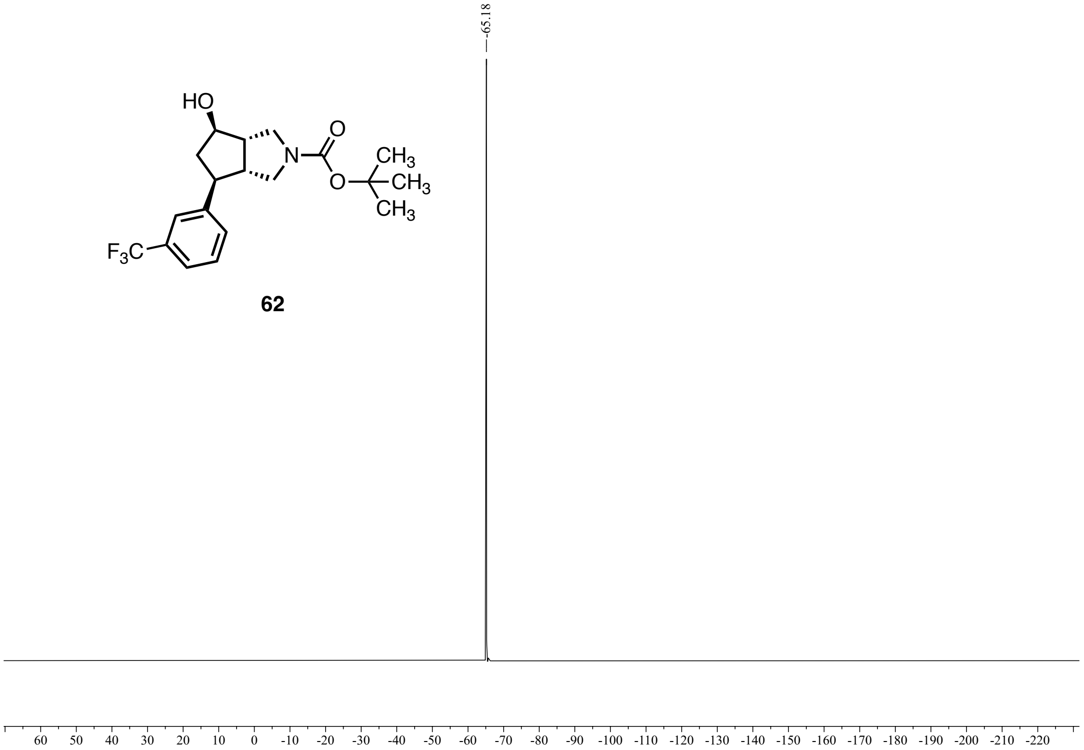


**Figure S84:** $^{19}$F NMR {$^{1}$H} (376 MHz, $CDCl_3$) of **62**.

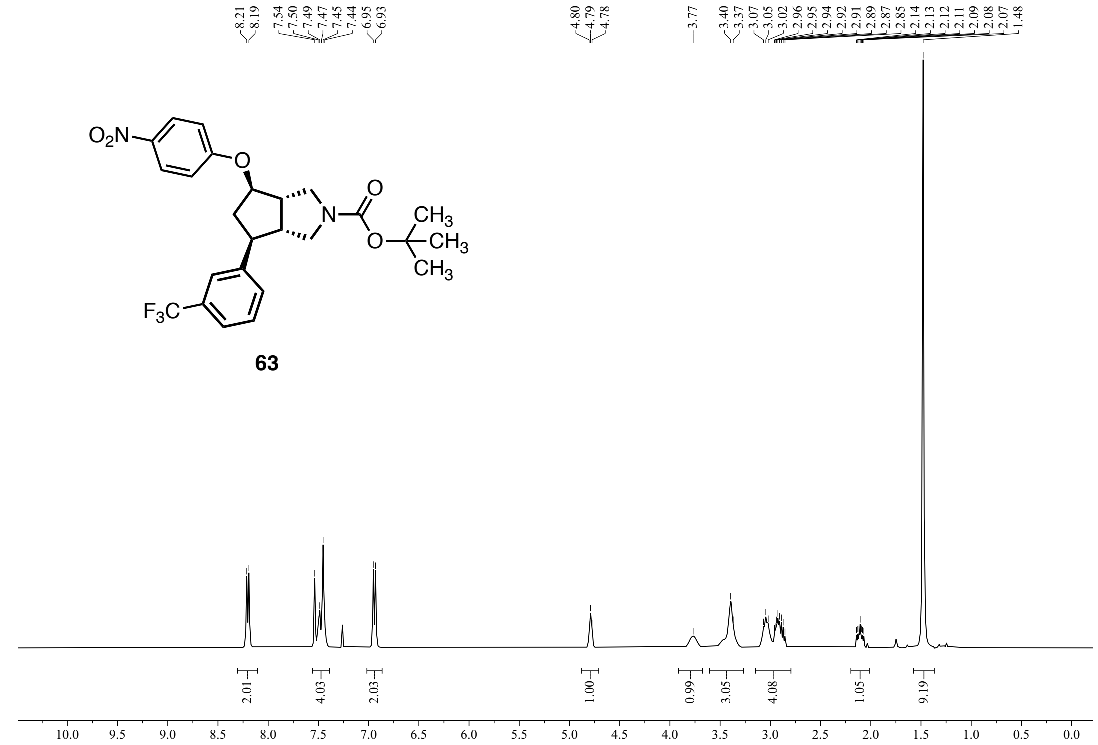


**Figure S85:** $^1$H NMR (400 MHz, $CDCl_3$) of **63**.

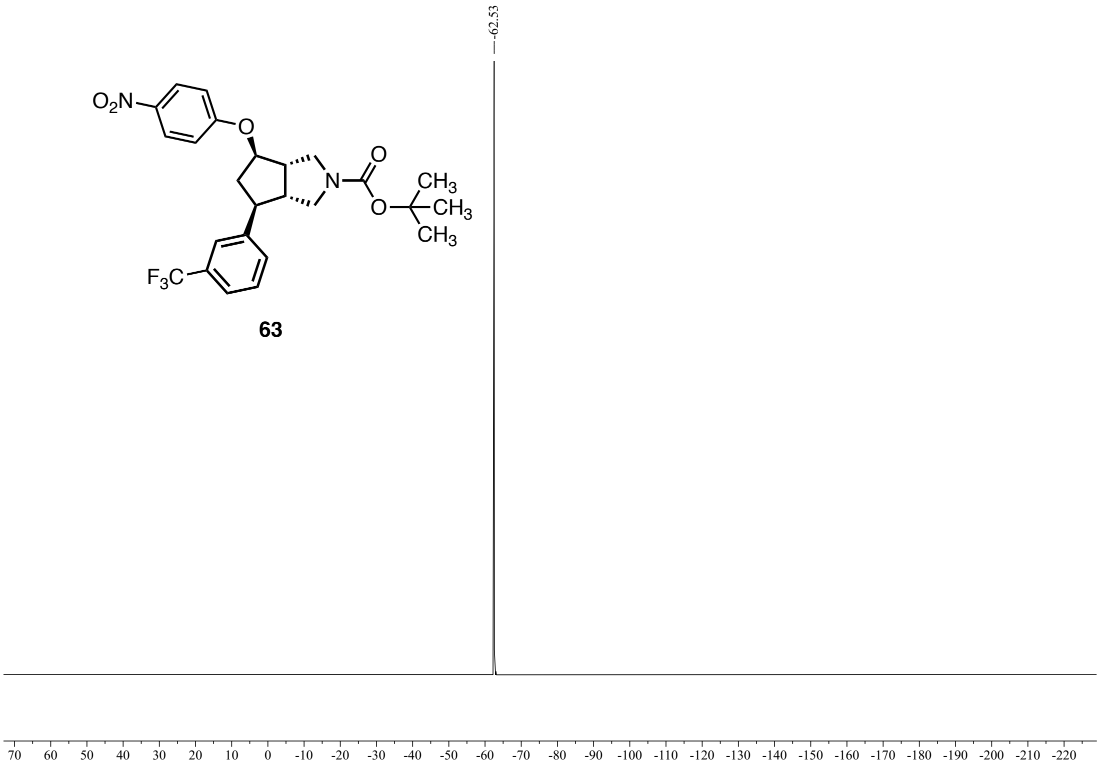


**Figure S86:** $^{19}$F NMR {$^1$H} (376 MHz, $CDCl_3$) of **63**.

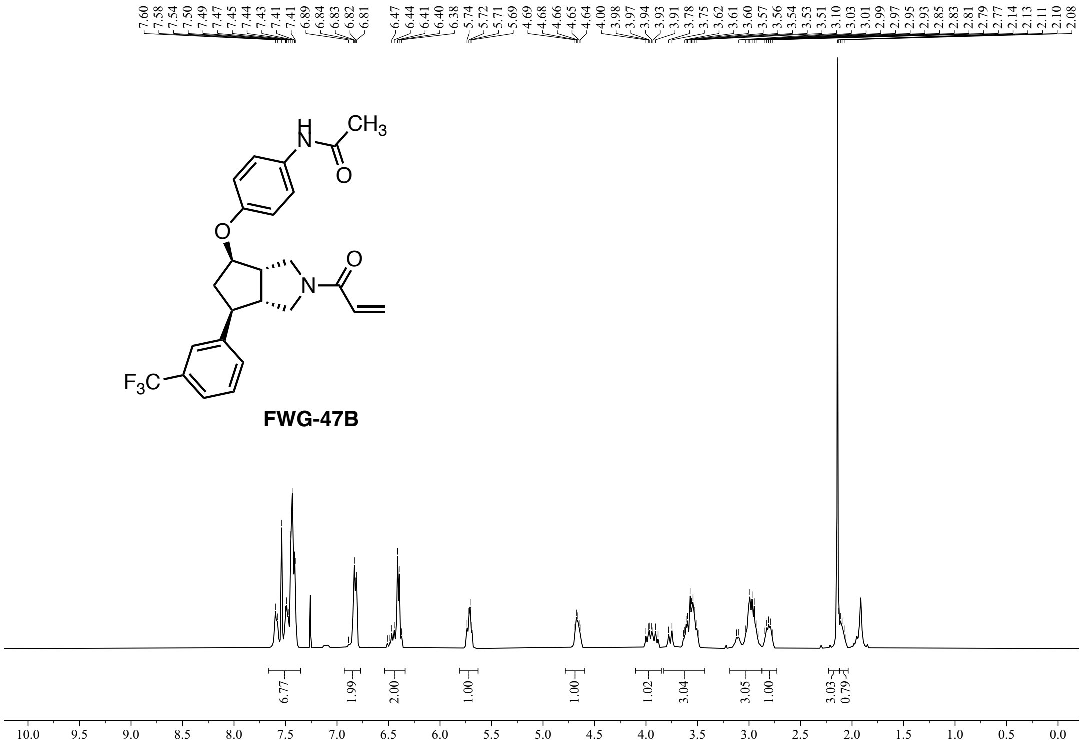

FWG-47B

Figure S87: $^{1}$H NMR (400 MHz, $CDCl_3$) of **FWG-47B.**

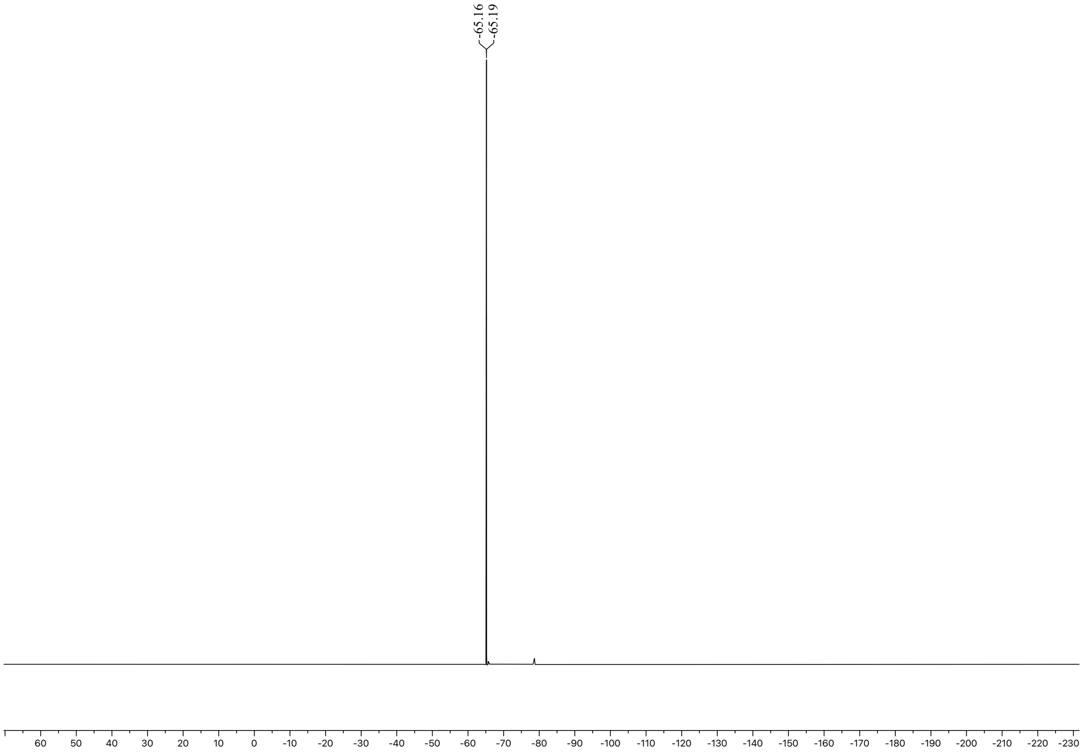

**Figure S88:** $^{19}$F NMR {$^{1}$H} (376 MHz, $CDCl_3$) of **FWG-47B.**

#### SFC

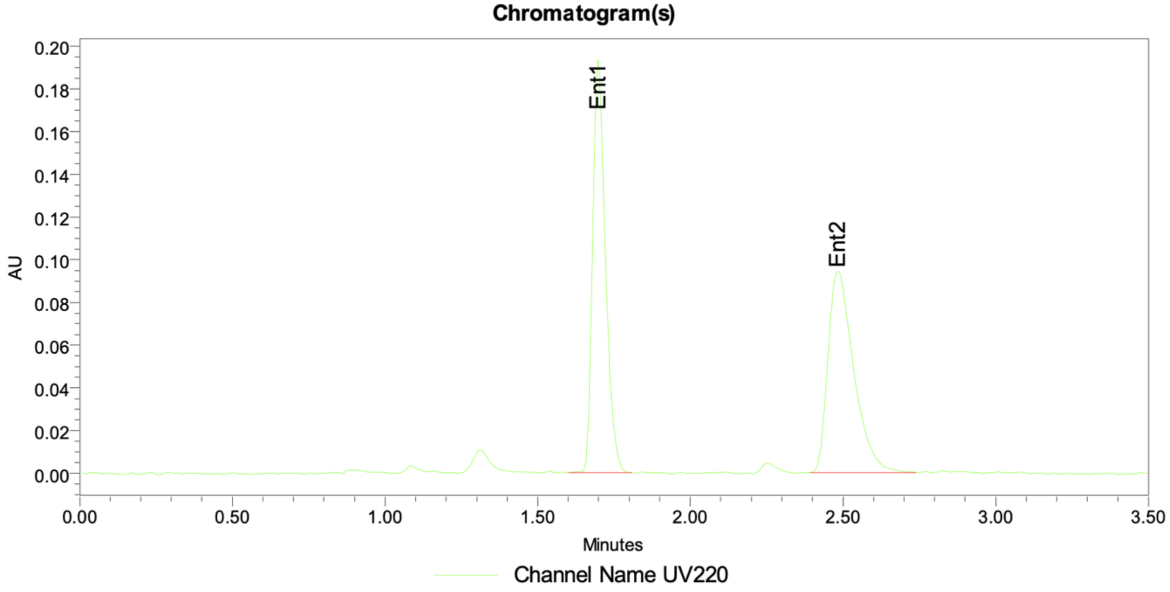


Peak Info

| | Channel Name | Name | RT | Area | Height (µV) | ent1 | ent2 | ee |
|---|---|---|---|---|---|---|---|---|
| 1 | UV220 | Ent1 | 1.70 | 565007 | 194168 | 50.41 | 49.59 | 0.83 |
| 2 | UV220 | Ent2 | 2.48 | 555733 | 94420 | 50.41 | 49.59 | 0.83 |

**Figure S89:** SFC trace of (±)-**3**.

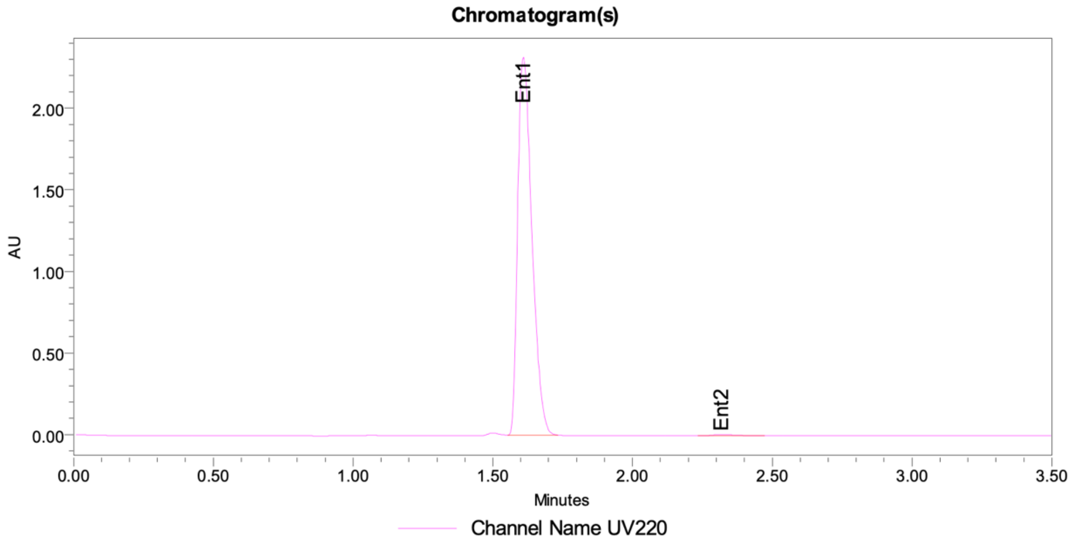


Peak Info

| | Channel Name | Name | RT | Area | Height (µV) | ent1 | ent2 | ee |
|---|---|---|---|---|---|---|---|---|
| 1 | UV220 | Ent1 | 1.61 | 8290512 | 2315949 | 99.52 | 0.48 | 99.03 |
| 2 | UV220 | Ent2 | 2.32 | 40355 | 6770 | 99.52 | 0.48 | 99.03 |

**Figure S90:** SFC trace of **3**.

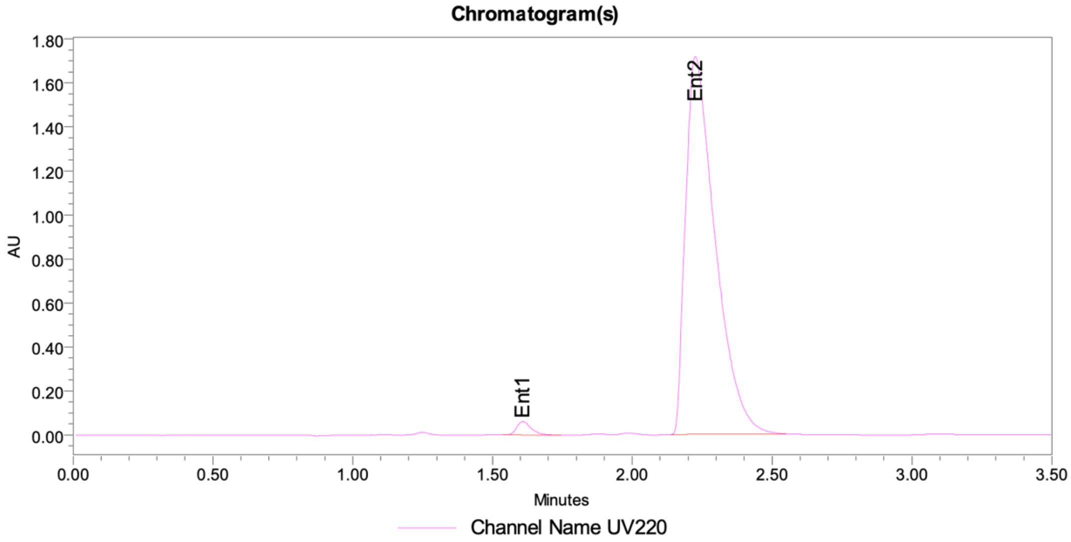


Peak Info

| | Channel Name | Name | RT | Area | Height (µV) | ent1 | ent2 | ee |
|---|---|---|---|---|---|---|---|---|
| 1 | UV220 | Ent1 | 1.61 | 215983 | 61260 | 1.68 | 98.32 | -96.64 |
| 2 | UV220 | Ent2 | 2.23 | 12639153 | 1717365 | 1.68 | 98.32 | -96.64 |

**Figure S91:** SFC trace of **4**.

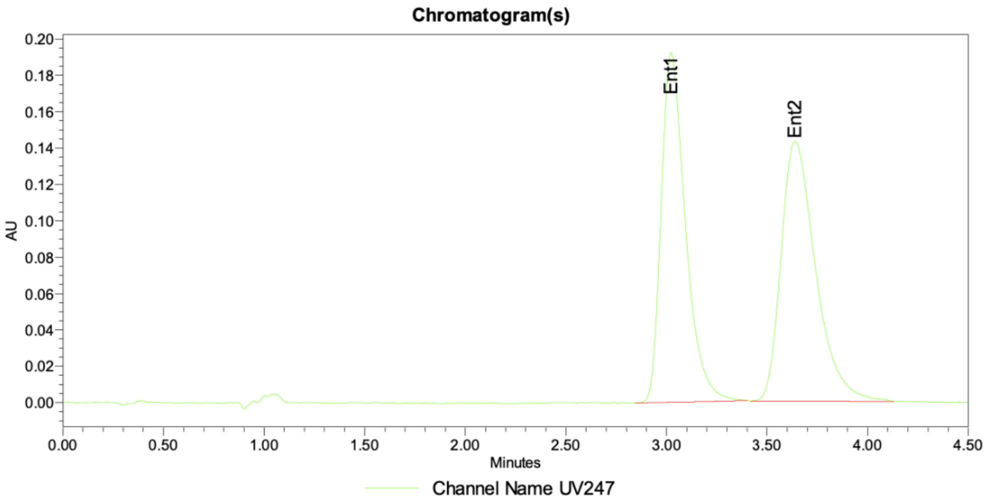


Peak Info

| | Channel Name | Name | RT | Area | Height (μV) | ent1 | ent2 | ee |
|---|---|---|---|---|---|---|---|---|
| 1 | UV247 | Ent1 | 3.02 | 1617203 | 192319 | 48.67 | 51.33 | -2.67 |
| 2 | UV247 | Ent2 | 3.64 | 1705807 | 142926 | 48.67 | 51.33 | -2.67 |

**Figure S92:** SFC trace of an equimolar mixture of **FWG-1A** and **FWG-1B.**

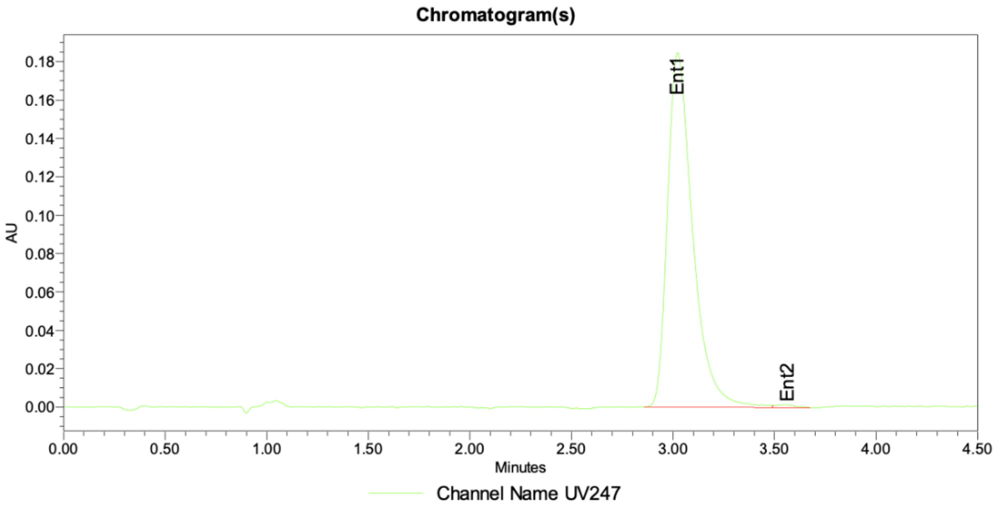


Peak Info

| | Channel Name | Name | RT | Area | Height (μV) | ent1 | ent2 | ee |
|---|---|---|---|---|---|---|---|---|
| 1 | UV247 | Ent1 | 3.02 | 1599391 | 184797 | 99.38 | 0.62 | 98.76 |
| 2 | UV247 | Ent2 | 3.57 | 9959 | 1468 | 99.38 | 0.62 | 98.76 |

**Figure S93:** SFC trace of **FWG-1A**.

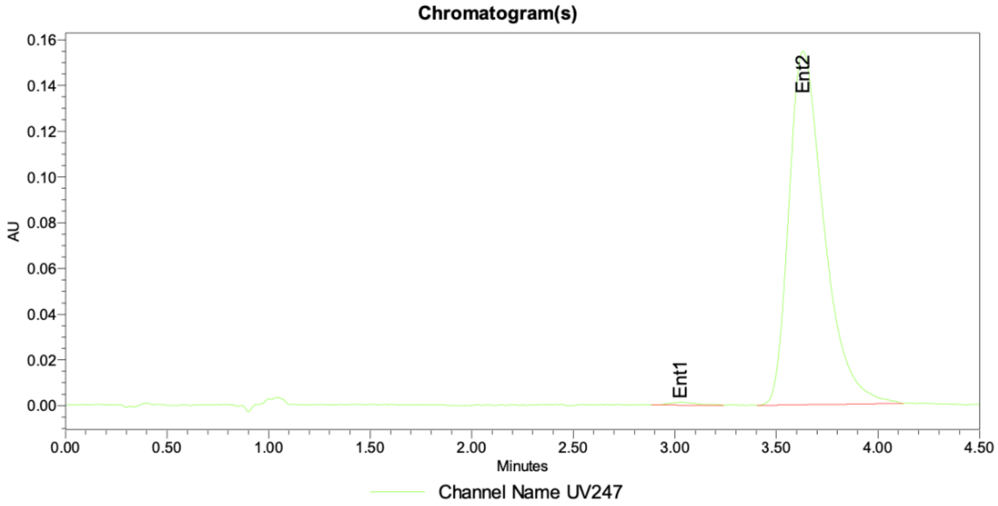


Peak Info

| | Channel Name | Name | RT | Area | Height (μV) | ent1 | ent2 | ee |
|---|---|---|---|---|---|---|---|---|
| 1 | UV247 | Ent1 | 3.03 | 12120 | 1381 | 0.65 | 99.35 | -98.71 |
| 2 | UV247 | Ent2 | 3.63 | 1863267 | 154661 | 0.65 | 99.35 | -98.71 |

**Figure S94:** SFC trace of **FWG-1B**.

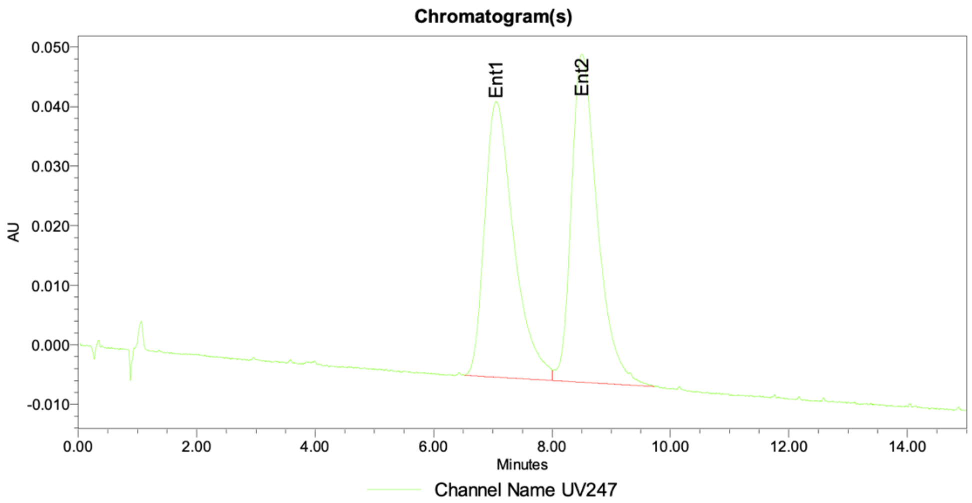


Peak Info

| | Channel Name | Name | RT | Area | Height (μV) | ent1 | ent2 | ee |
|---|---|---|---|---|---|---|---|---|
| 1 | UV247 | Ent1 | 7.05 | 1582516 | 46267 | 48.40 | 51.60 | -3.20 |
| 2 | UV247 | Ent2 | 8.51 | 1686977 | 55090 | 48.40 | 51.60 | -3.20 |

**Figure S95:** SFC trace of an equimolar mixture of **FWG-2A** and **FWG-2B**.

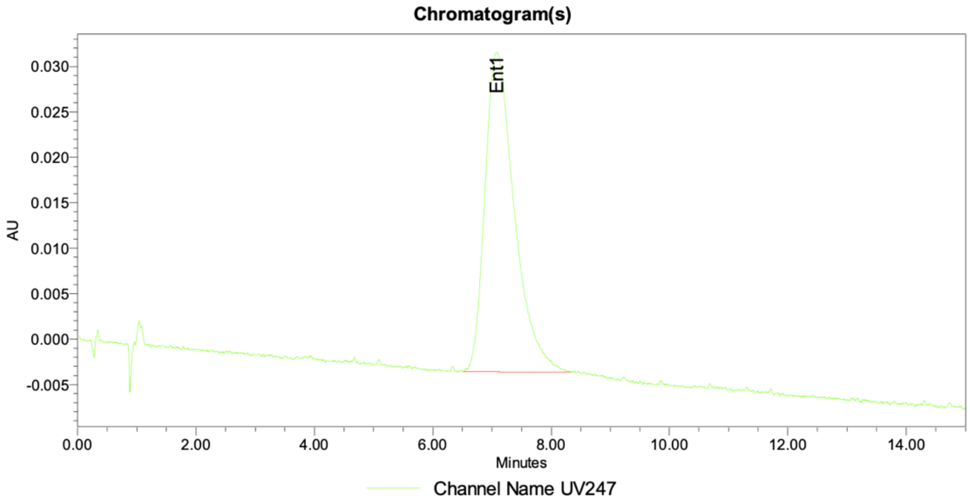


Peak Info

| | Channel Name | Name | RT | Area | Height (μV) | ent1 | ent2 | ee |
|---|---|---|---|---|---|---|---|---|
| 1 | UV247 | Ent1 | 7.08 | 1213865 | 35178 | 100.00 | | 100.00 |
| 2 | UV247 | Ent2 | 8.51 | | | 100.00 | | 100.00 |

**Figure S96:** SFC trace of **FWG-2A**.

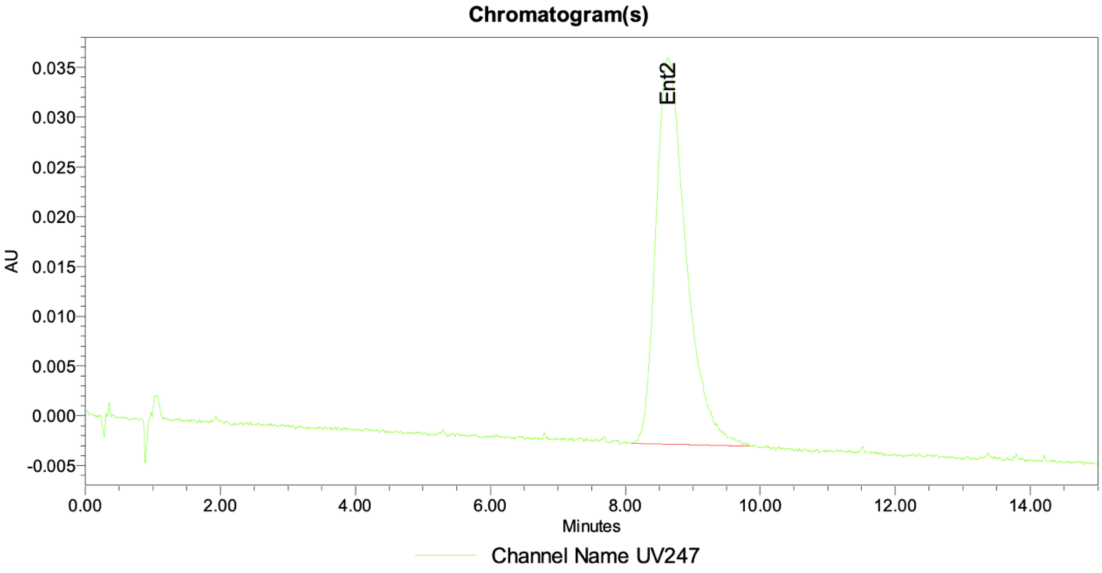


Peak Info

| | Channel Name | Name | RT | Area | Height (μV) | ent1 | ent2 | ee |
|---|---|---|---|---|---|---|---|---|
| 1 | UV247 | Ent1 | 7.05 | | | | | |
| 2 | UV247 | Ent2 | 8.63 | 1201315 | 38801 | | | |

**Figure S97:** SFC trace of **FWG-2B**.

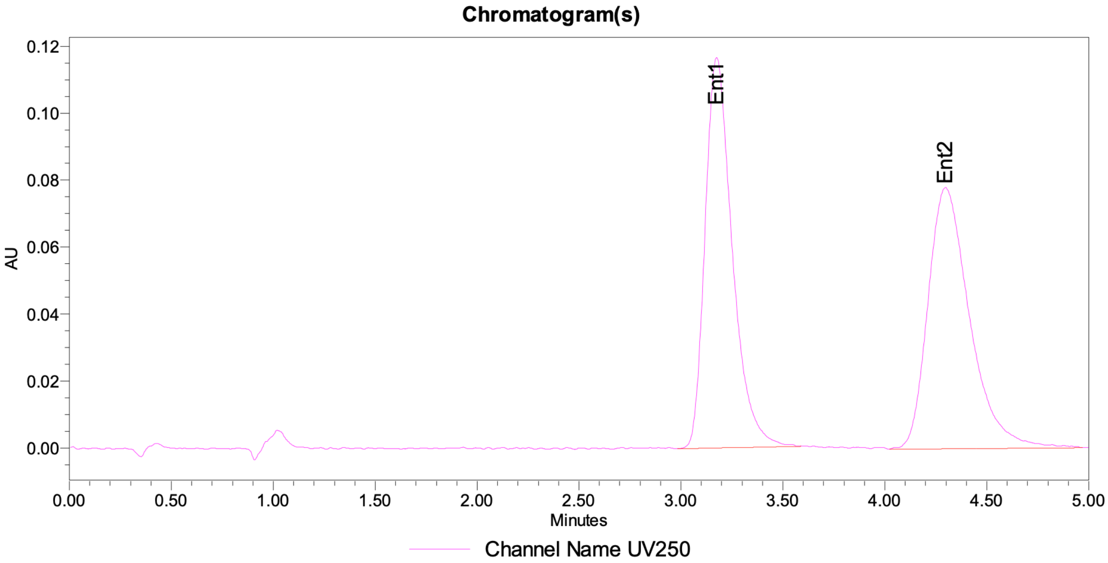


Peak Info

| | Channel Name | Name | RT | Area | Height (μV) | ent1 | ent2 | ee |
|---|---|---|---|---|---|---|---|---|
| 1 | UV250 | Ent1 | 3.18 | 1084502 | 116635 | 49.38 | 50.62 | -1.24 |
| 2 | UV250 | Ent2 | 4.30 | 1111702 | 77997 | 49.38 | 50.62 | -1.24 |

**Figure S98:** SFC trace of an equimolar mixture of **FWG-3A** and **FWG-3B.**

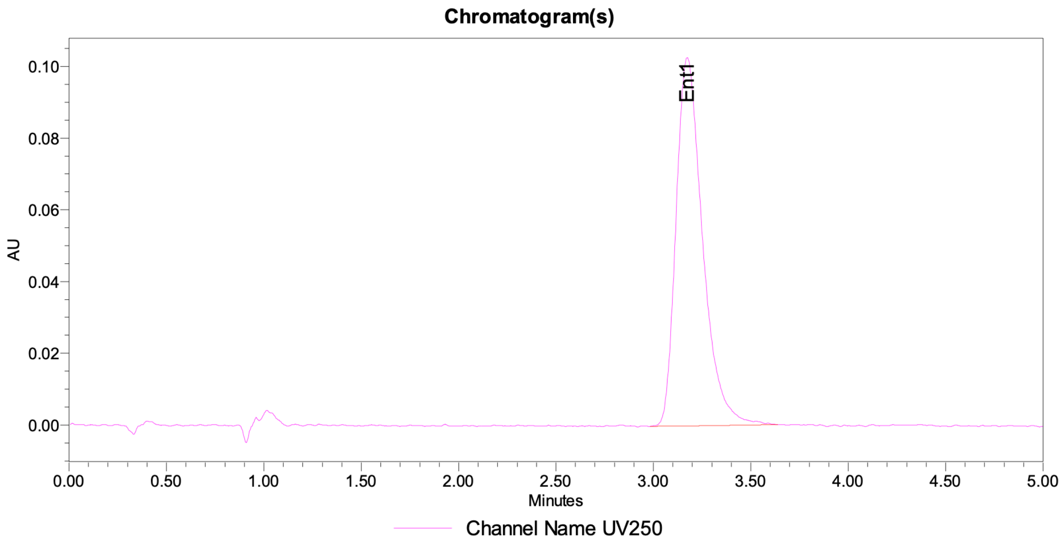


Peak Info

| | Channel Name | Name | RT | Area | Height (μV) | ent1 | ent2 | ee |
|---|---|---|---|---|---|---|---|---|
| 1 | UV250 | Ent1 | 3.17 | 976114 | 102753 | 100.00 | | 100.00 |
| 2 | UV250 | Ent2 | 4.45 | | | 100.00 | | 100.00 |

**Figure S99:** SFC trace of **FWG-3A**.

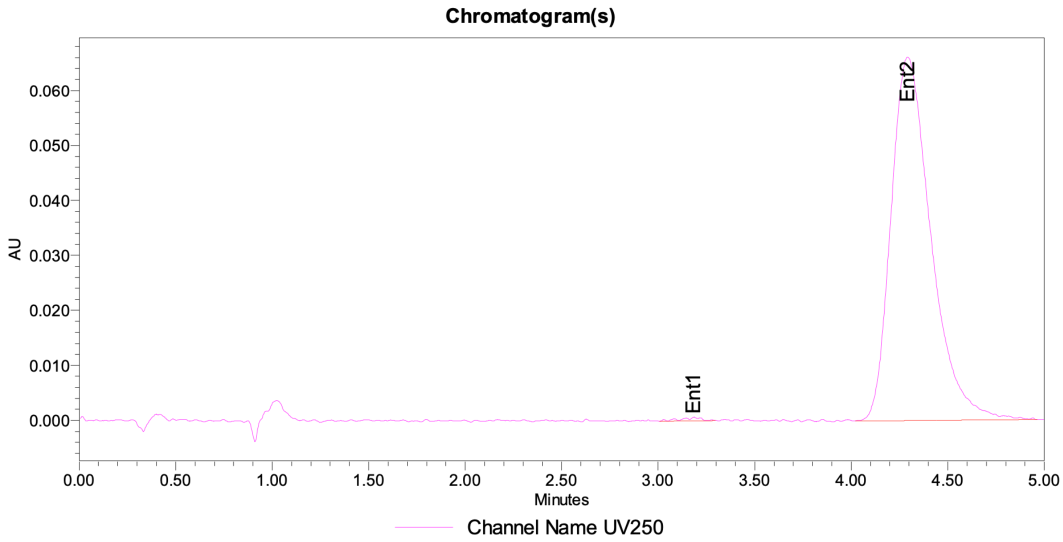


Peak Info

| | Channel Name | Name | RT | Area | Height (μV) | ent1 | ent2 | ee |
|---|---|---|---|---|---|---|---|---|
| 1 | UV250 | Ent1 | 3.18 | 4707 | 660 | 0.49 | 99.51 | -99.02 |
| 2 | UV250 | Ent2 | 4.29 | 952400 | 66135 | 0.49 | 99.51 | -99.02 |

**Figure S100:** SFC trace of **FWG-3B**.

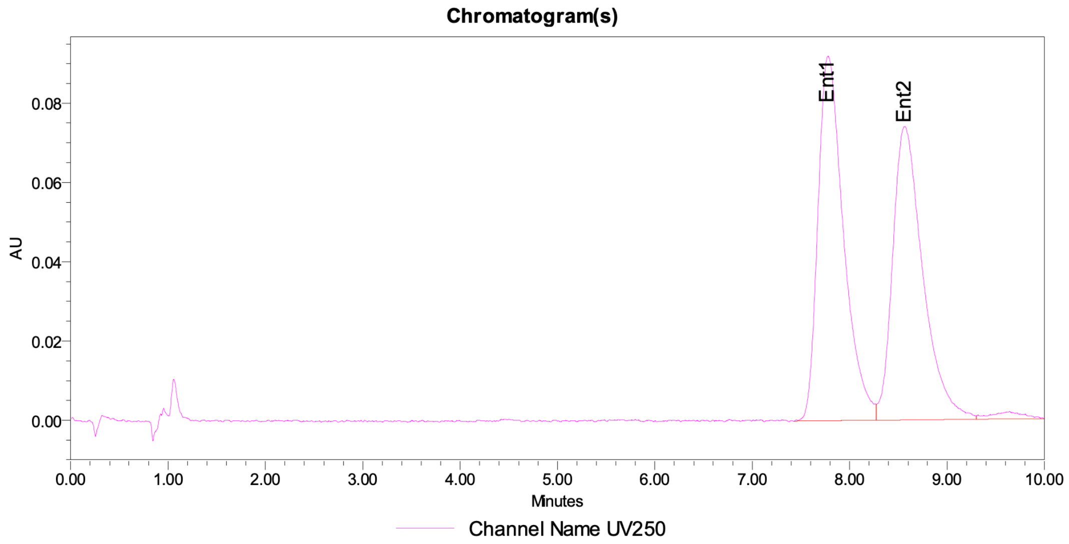


Peak Info

| | Channel Name | Name | RT | Area | Height (µV) | ent1 | ent2 | ee |
|---|---|---|---|---|---|---|---|---|
| 1 | UV250 | Ent1 | 7.78 | 1688279 | 92065 | 51.42 | 48.58 | 2.85 |
| 2 | UV250 | Ent2 | 8.56 | 1594802 | 74040 | 51.42 | 48.58 | 2.85 |
| 3 | UV250 | | 9.64 | 43319 | 1823 | 51.42 | 48.58 | 2.85 |

**Figure S101:** SFC trace of an equimolar mixture of **FWG-4A** and **FWG-4B.**

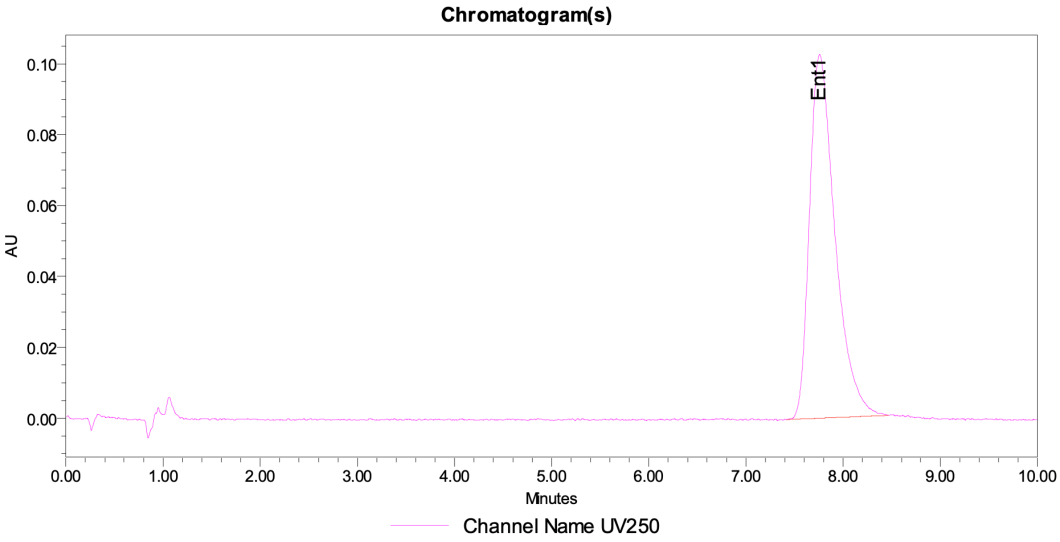


Peak Info

| | Channel Name | Name | RT | Area | Height (μV) | ent1 | ent2 | ee |
|---|---|---|---|---|---|---|---|---|
| 1 | UV250 | Ent1 | 7.76 | 1875385 | 102678 | 100.00 | | 100.00 |
| 2 | UV250 | Ent2 | 8.56 | | | 100.00 | | 100.00 |

**Figure S102:** SFC trace of **FWG-4A**.

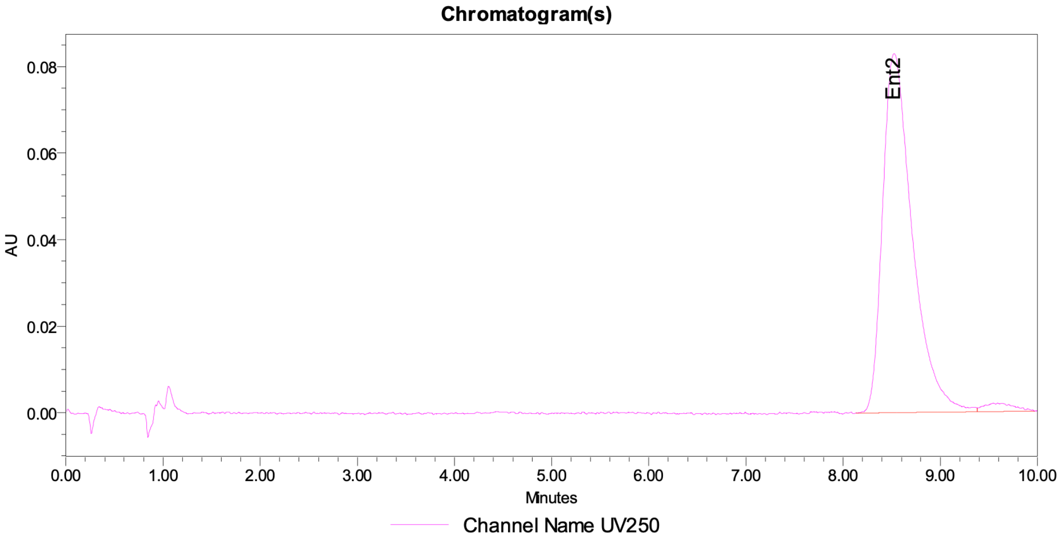


Peak Info

| | Channel Name | Name | RT | Area | Height (μV) | ent1 | ent2 | ee |
|---|---|---|---|---|---|---|---|---|
| 1 | UV250 | Ent1 | 7.78 | | | | | |
| 2 | UV250 | Ent2 | 8.52 | 1765933 | 82963 | | | |
| 3 | UV250 | | 9.54 | 42165 | 1952 | | | |

**Figure S103:** SFC trace of **FWG-4B**.

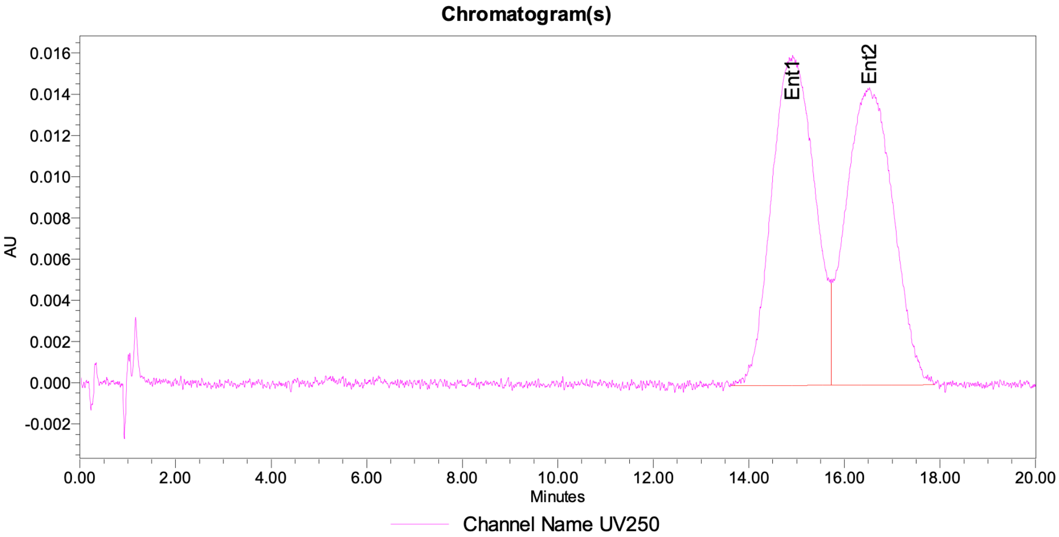


Peak Info

| | Channel Name | Name | RT | Area | Height (μV) | ent1 | ent2 | ee |
|---|---|---|---|---|---|---|---|---|
| 1 | UV250 | Ent1 | 14.92 | 999468 | 16011 | 50.08 | 49.92 | 0.15 |
| 2 | UV250 | Ent2 | 16.52 | 996426 | 14415 | 50.08 | 49.92 | 0.15 |

**Figure S104:** SFC trace of SFC trace of an equimolar mixture of **FWG-9A** and **FWG-9B.**

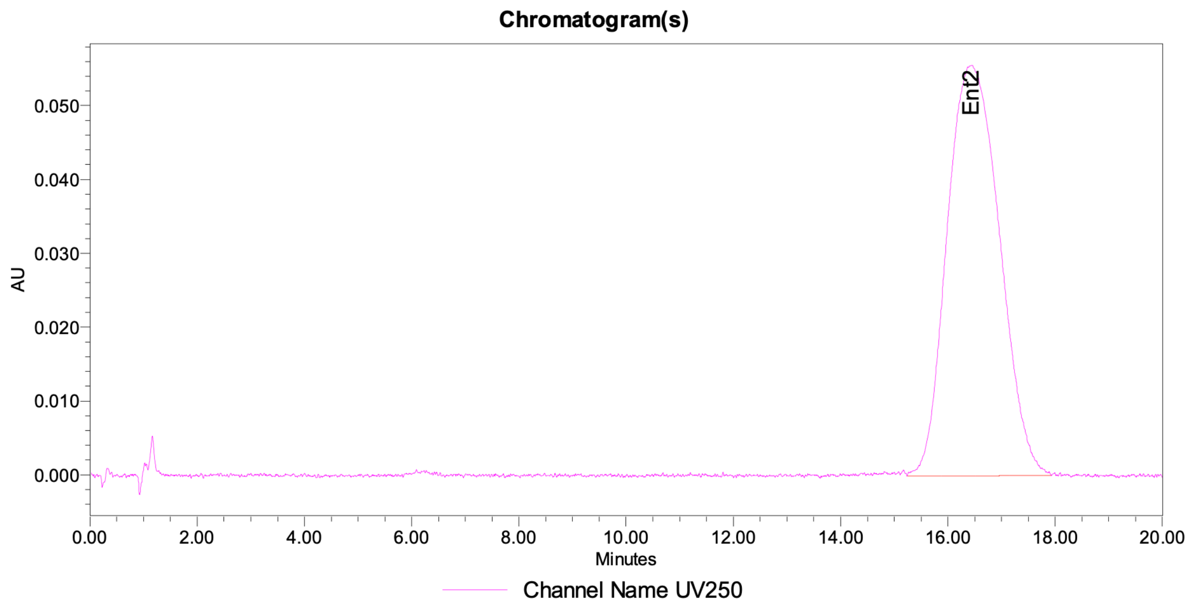


Peak Info

| | Channel Name | Name | RT | Area | Height (µV) | ent1 | ent2 | ee |
|---|---|---|---|---|---|---|---|---|
| 1 | UV250 | Ent1 | 14.92 | | | | | |
| 2 | UV250 | Ent2 | 16.43 | 3778272 | 55611 | | | |

**Figure S105:** SFC trace of **FWG-9A**.

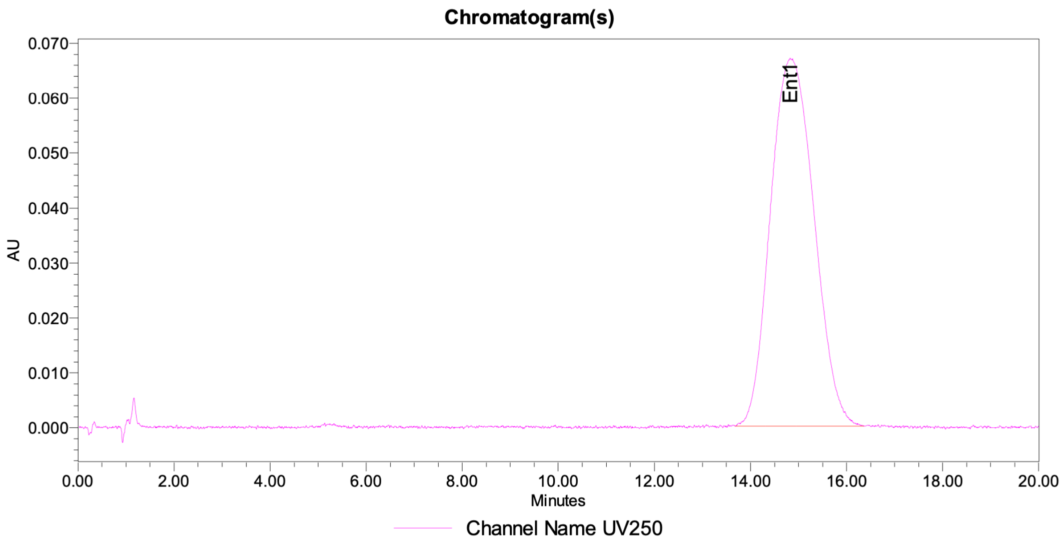


Peak Info

| | Channel Name | Name | RT | Area | Height (µV) | ent1 | ent2 | ee |
|---|---|---|---|---|---|---|---|---|
| 1 | UV250 | Ent1 | 14.83 | 4224412 | 66988 | 100.00 | | 100.00 |
| 2 | UV250 | Ent2 | 16.52 | | | 100.00 | | 100.00 |

**Figure S106:** SFC trace of **FWG-9B**.

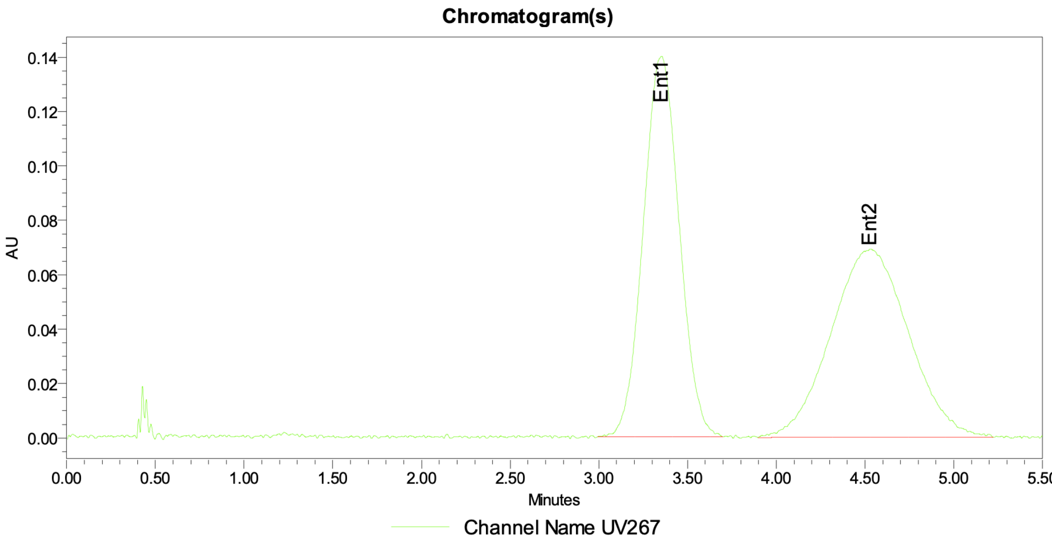


Peak Info

| | Channel Name | Name | RT | Area | Height (µV) | ent1 | ent2 | ee |
|---|---|---|---|---|---|---|---|---|
| 1 | UV267 | Ent1 | 3.35 | 1973254 | 139975 | 48.42 | 51.58 | -3.16 |
| 2 | UV267 | Ent2 | 4.53 | 2101964 | 69220 | 48.42 | 51.58 | -3.16 |

**Figure S107:** SFC trace of SFC trace of an equimolar mixture of **FWG-11A** and **FWG-11B.**

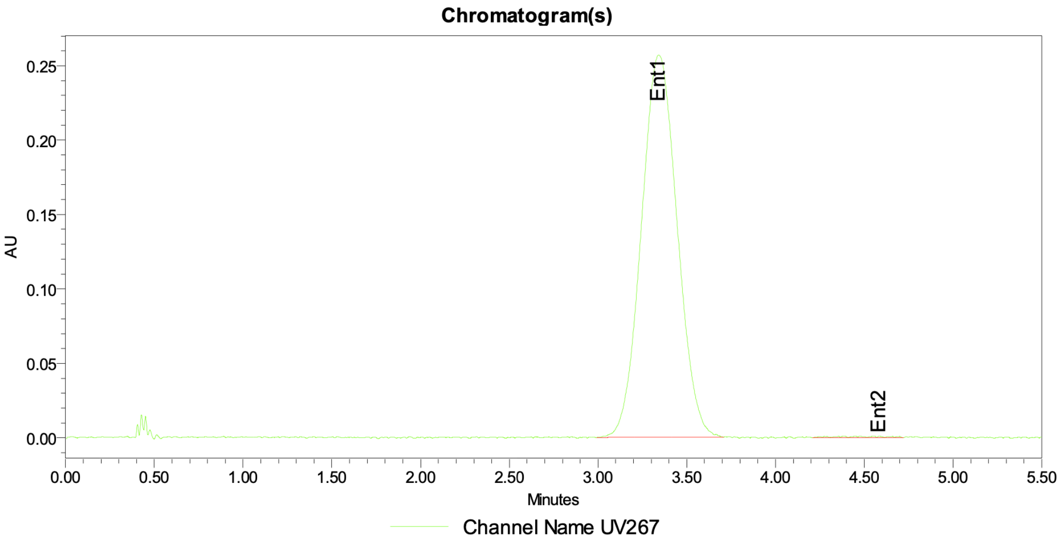


Peak Info

| | Channel Name | Name | RT | Area | Height (µV) | ent1 | ent2 | ee |
|---|---|---|---|---|---|---|---|---|
| 1 | UV267 | Ent1 | 3.34 | 3599994 | 256808 | 99.48 | 0.52 | 98.97 |
| 2 | UV267 | Ent2 | 4.58 | 18672 | 1118 | 99.48 | 0.52 | 98.97 |

**Figure S108:** SFC trace of **FWG-11A**.

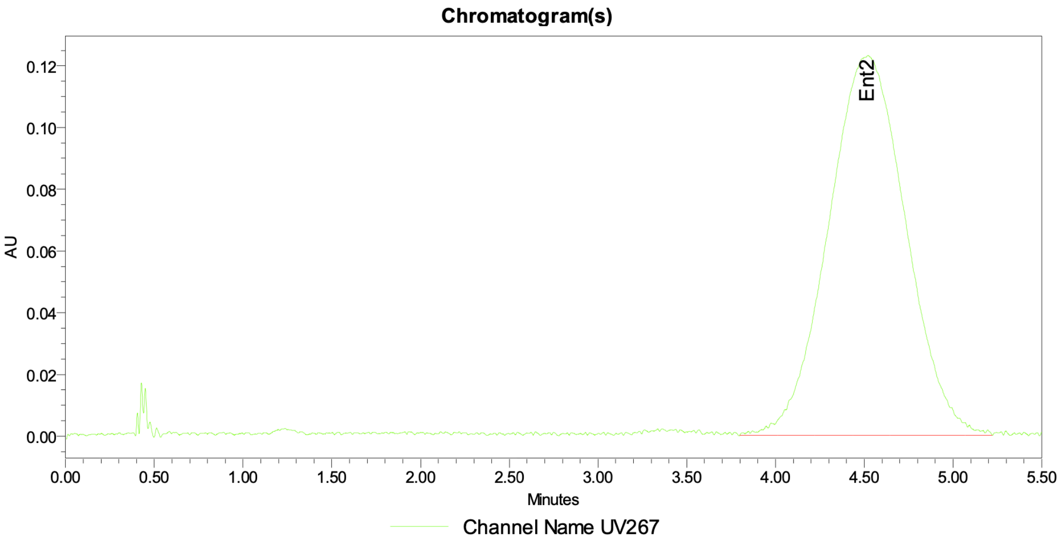


Peak Info

| | Channel Name | Name | RT | Area | Height (µV) | ent1 | ent2 | ee |
|---|---|---|---|---|---|---|---|---|
| 1 | UV267 | Ent1 | 3.36 | | | | 100.00 | -100.00 |
| 2 | UV267 | Ent2 | 4.52 | 3729354 | 123022 | | 100.00 | -100.00 |

**Figure S109:** SFC trace of **FWG-11B**.

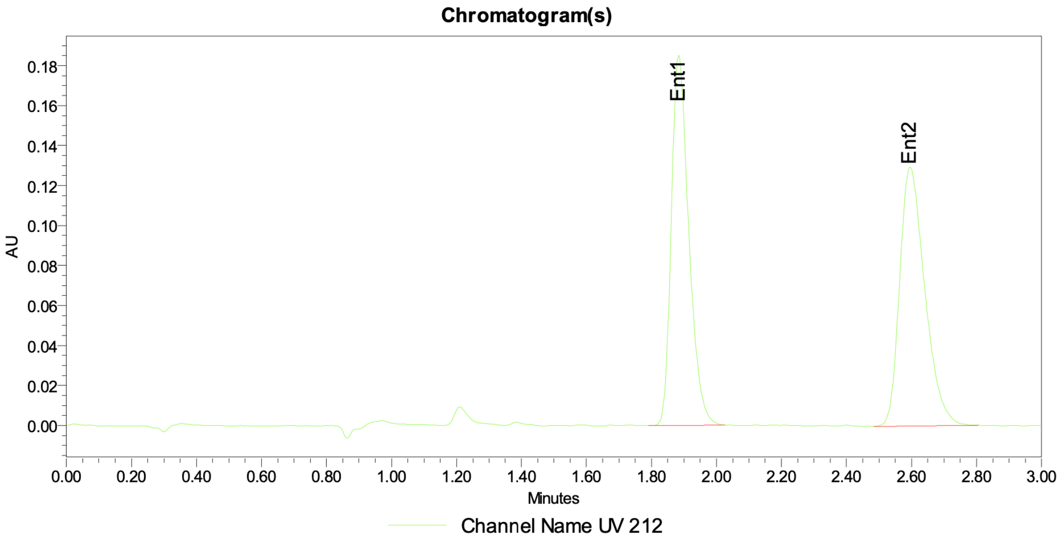


Peak Info

| | Channel Name | Name | RT | Area | Height (µV) | ent1 | ent2 | ee |
|---|---|---|---|---|---|---|---|---|
| 1 | UV 212 | Ent1 | 1.88 | 694154 | 185279 | 50.02 | 49.98 | 0.03 |
| 2 | UV 212 | Ent2 | 2.60 | 693725 | 129470 | 50.02 | 49.98 | 0.03 |

**Figure S110:** SFC trace of (±)-**41**.

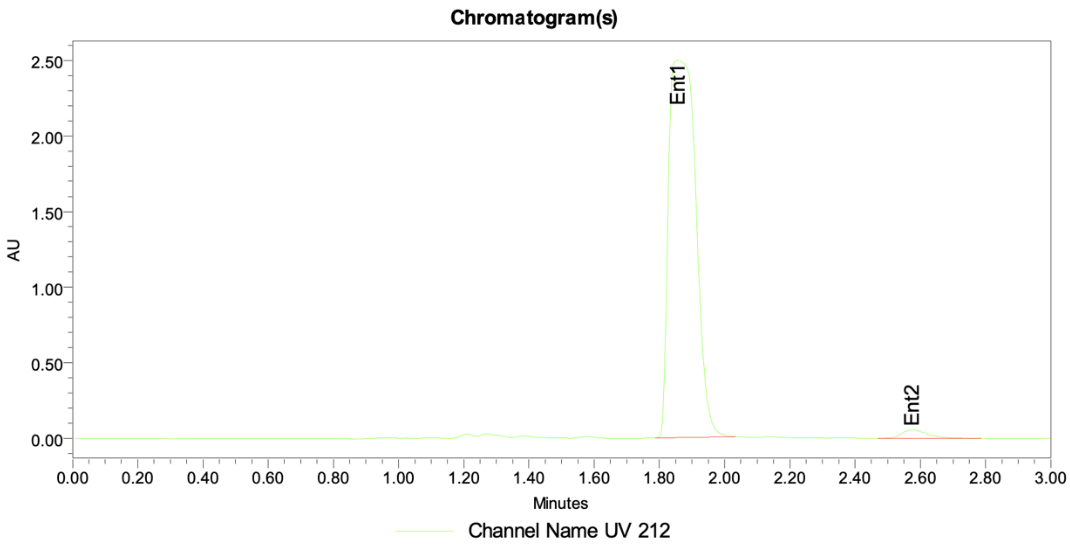


Peak Info

| | Channel Name | Name | RT | Area | Height (μV) | ent1 | ent2 | ee |
|---|---|---|---|---|---|---|---|---|
| 1 | UV 212 | Ent1 | 1.86 | 14451947 | 2497262 | 98.01 | 1.99 | 96.02 |
| 2 | UV 212 | Ent2 | 2.58 | 293148 | 55169 | 98.01 | 1.99 | 96.02 |

**Figure S111:** SFC trace of **41**.

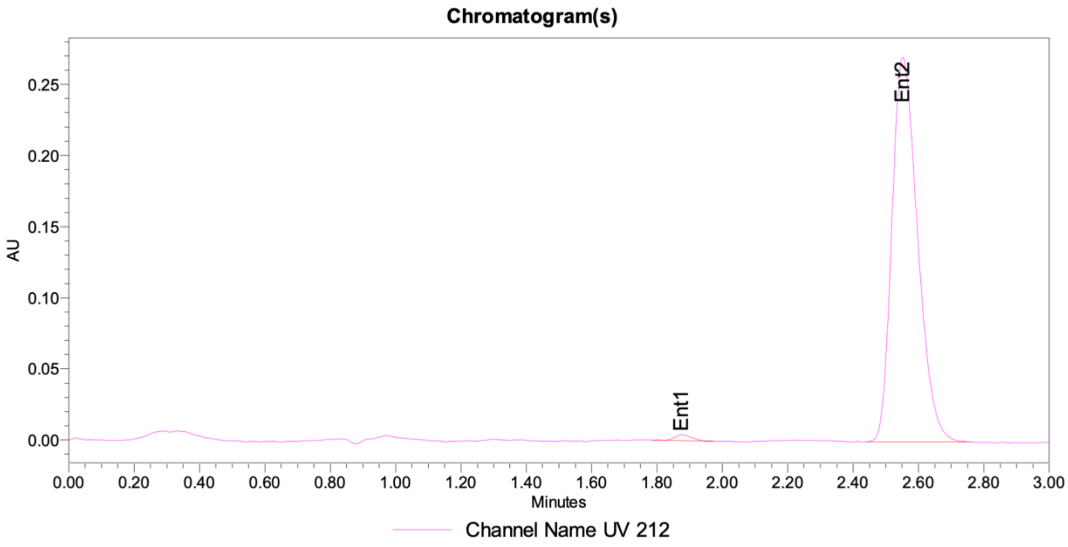


Peak Info

| | Channel Name | Name | RT | Area | Height (μV) | ent1 | ent2 | ee |
|---|---|---|---|---|---|---|---|---|
| 1 | UV 212 | Ent1 | 1.88 | 15300 | 4207 | 1.02 | 98.98 | -97.95 |
| 2 | UV 212 | Ent2 | 2.55 | 1479189 | 270567 | 1.02 | 98.98 | -97.95 |

**Figure S112:** SFC trace of **42**.

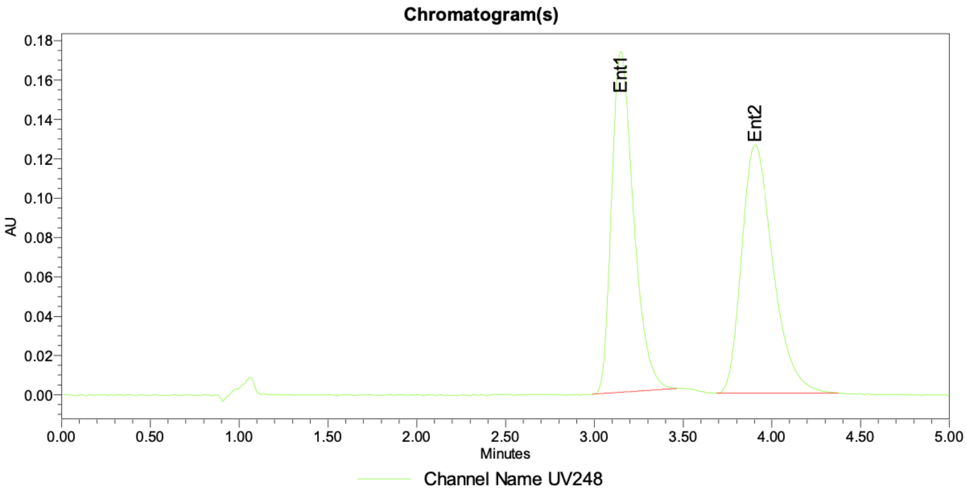


Peak Info

| | Channel Name | Name | RT | Area | Height (µV) | ent1 | ent2 | ee |
|---|---|---|---|---|---|---|---|---|
| 1 | UV248 | Ent1 | 3.15 | 1495766 | 173275 | 48.86 | 51.14 | -2.29 |
| 2 | UV248 | Ent2 | 3.90 | 1565734 | 126386 | 48.86 | 51.14 | -2.29 |

**Figure S113:** SFC trace of SFC trace of equimolar mixture of **FWG-33A** and **FWG-33B.**

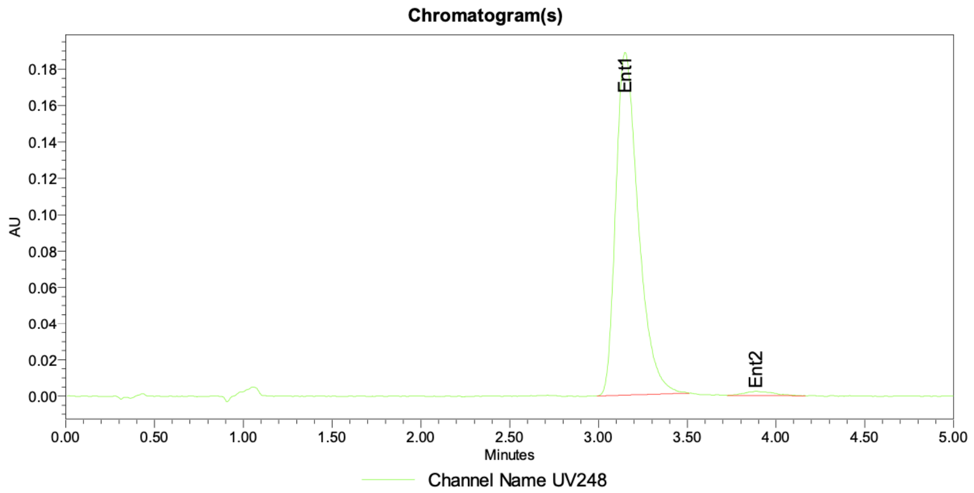


Peak Info

| | Channel Name | Name | RT | Area | Height (µV) | ent1 | ent2 | ee |
|---|---|---|---|---|---|---|---|---|
| 1 | UV248 | Ent1 | 3.15 | 1671637 | 188848 | 98.40 | 1.60 | 96.80 |
| 2 | UV248 | Ent2 | 3.88 | 27164 | 2366 | 98.40 | 1.60 | 96.80 |

**Figure S114:** SFC trace of **FWG-33A**.

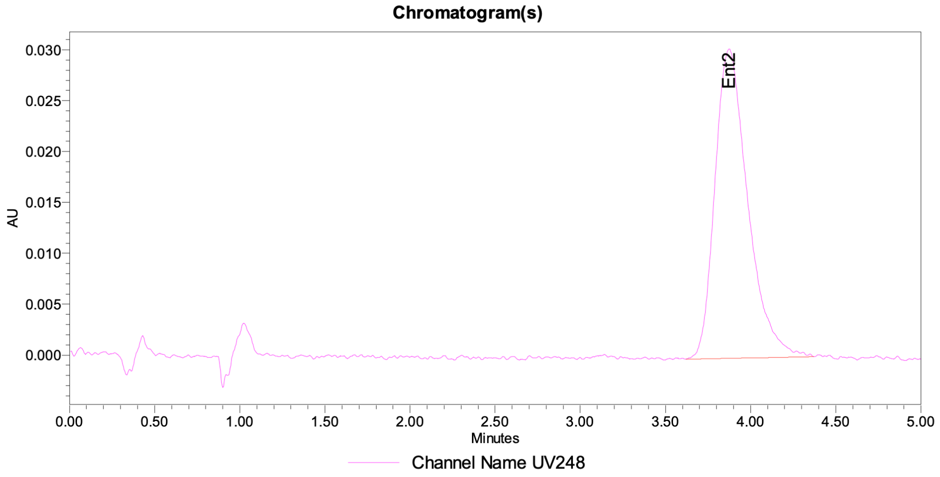


Peak Info

| | Channel Name | Name | RT | Area | Height (µV) | ent1 | ent2 | ee |
|---|---|---|---|---|---|---|---|---|
| 1 | UV248 | Ent1 | 3.19 | | | | | |
| 2 | UV248 | Ent2 | 3.87 | 408086 | 30411 | | | |

**Figure S115:** SFC trace of **FWG-33B**.

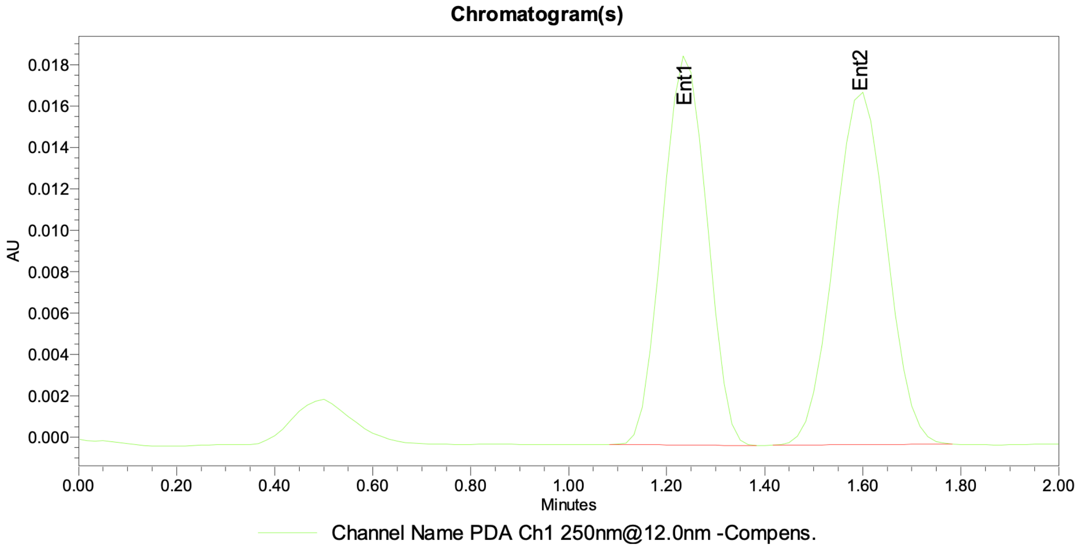


Peak Info

| | Channel Name | Name | RT | Area | Height (µV) | ent1 | ent2 | ee |
|---|---|---|---|---|---|---|---|---|
| 1 | PDA Ch1 250nm@12.0nm -Compens. | Ent1 | 1.24 | 118046 | 18715 | 48.03 | 51.97 | -3.95 |
| 2 | PDA Ch1 250nm@12.0nm -Compens. | Ent2 | 1.60 | 127747 | 17026 | 48.03 | 51.97 | -3.95 |

**Figure S116:** SFC trace of SFC trace of equimolar mixture of **FWG-49A** and **FWG-49B.**

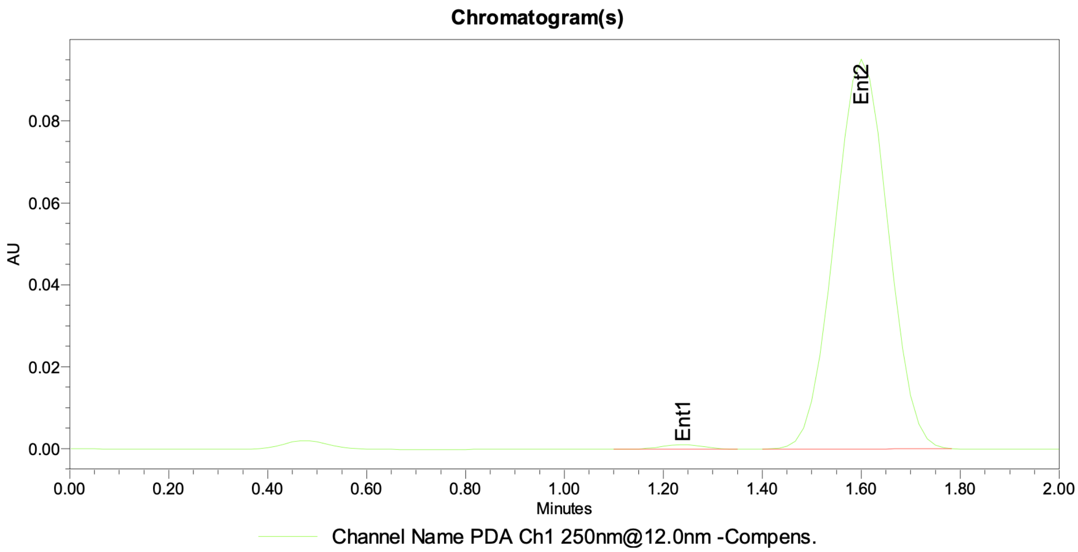


Peak Info

| | Channel Name | Name | RT | Area | Height (µV) | ent1 | ent2 | ee |
|---|---|---|---|---|---|---|---|---|
| 1 | PDA Ch1 250nm@12.0nm -Compens. | Ent1 | 1.24 | 7086 | 1129 | 0.98 | 99.02 | -98.04 |
| 2 | PDA Ch1 250nm@12.0nm -Compens. | Ent2 | 1.60 | 714950 | 94976 | 0.98 | 99.02 | -98.04 |

**Figure S117:** SFC trace of **FWG-49A**.

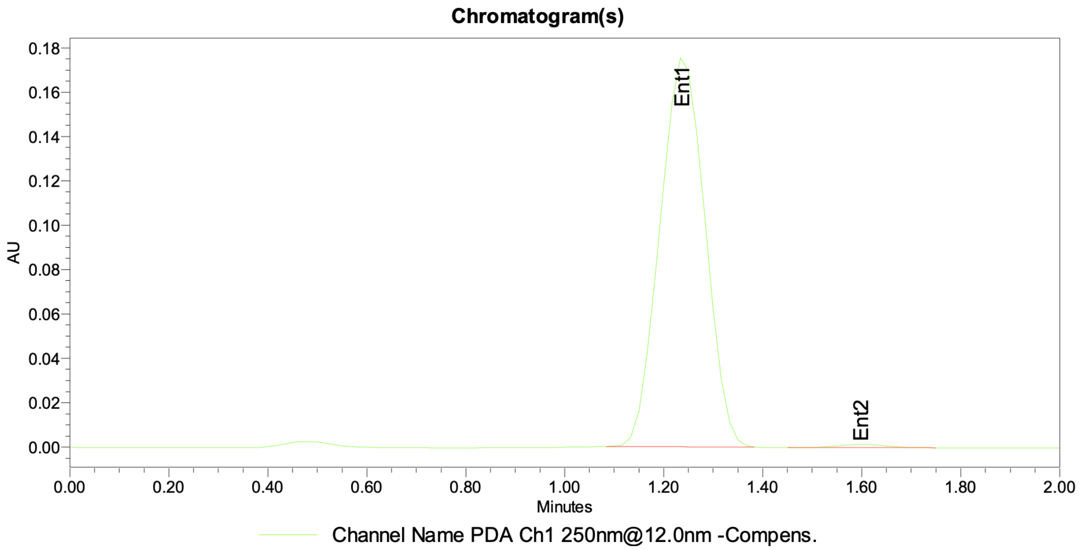


Peak Info

| | Channel Name | Name | RT | Area | Height (µV) | ent1 | ent2 | ee |
|---|---|---|---|---|---|---|---|---|
| 1 | PDA Ch1 250nm@12.0nm -Compens. | Ent1 | 1.24 | 1109681 | 175059 | 99.12 | 0.88 | 98.23 |
| 2 | PDA Ch1 250nm@12.0nm -Compens. | Ent2 | 1.60 | 9886 | 1365 | 99.12 | 0.88 | 98.23 |

**Figure S118:** SFC trace of **FWG-49B**.

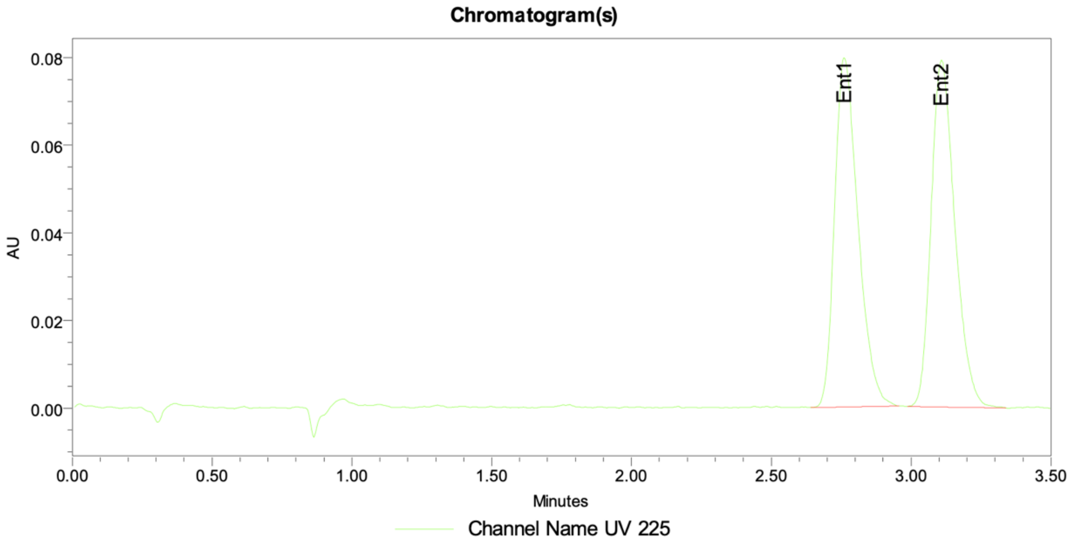


Peak Info

| | Channel Name | Name | RT | Area | Height (μV) | ent1 | ent2 | ee |
|---|---|---|---|---|---|---|---|---|
| 1 | UV 225 | Ent1 | 2.76 | 464750 | 79732 | 49.85 | 50.15 | -0.29 |
| 2 | UV 225 | Ent2 | 3.11 | 467479 | 79179 | 49.85 | 50.15 | -0.29 |

**Figure S119:** SFC trace of (±)-**52**.

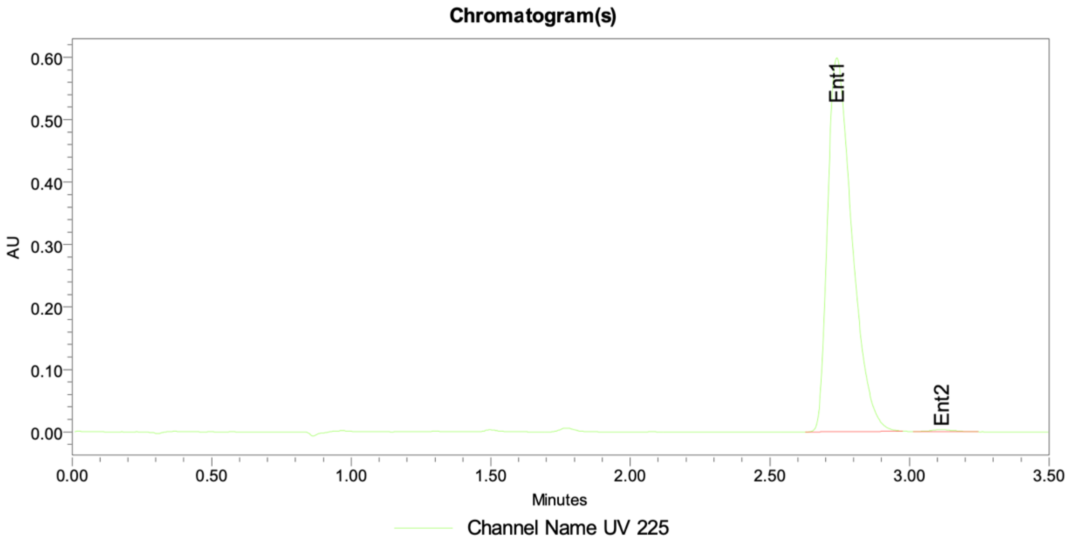


Peak Info

| | Channel Name | Name | RT | Area | Height (μV) | ent1 | ent2 | ee |
|---|---|---|---|---|---|---|---|---|
| 1 | UV 225 | Ent1 | 2.74 | 3642586 | 599048 | 99.53 | 0.47 | 99.06 |
| 2 | UV 225 | Ent2 | 3.12 | 17151 | 3115 | 99.53 | 0.47 | 99.06 |

**Figure S120:** SFC trace of **52**.

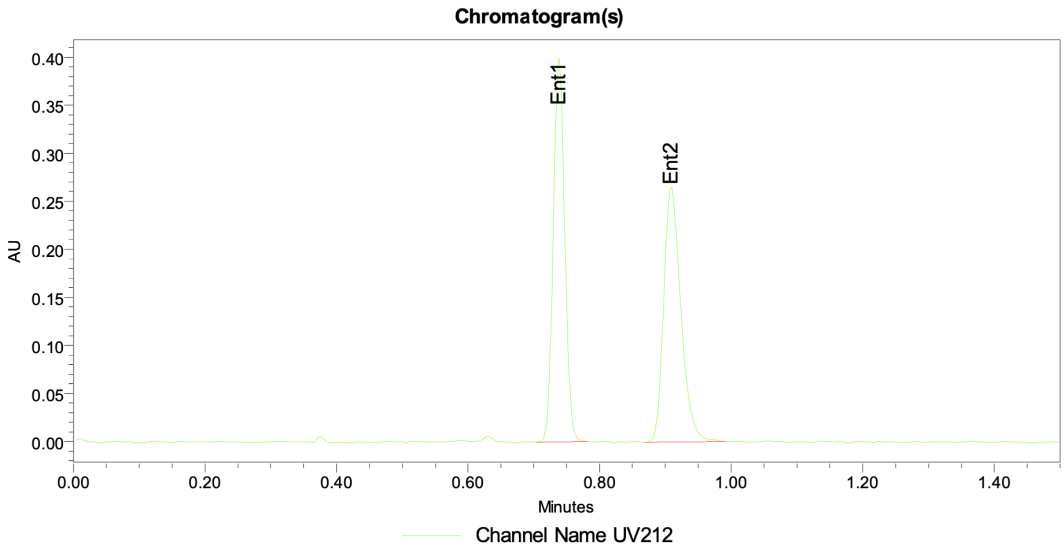


Peak Info

| | Channel Name | Name | RT | Area | Height (μV) | ent1 | ent2 | ee |
|---|---|---|---|---|---|---|---|---|
| 1 | UV212 | Ent1 | 0.74 | 481464 | 398384 | 50.07 | 49.93 | 0.13 |
| 2 | UV212 | Ent2 | 0.91 | 480191 | 264996 | 50.07 | 49.93 | 0.13 |

**Figure S121:** SFC trace of **55**.

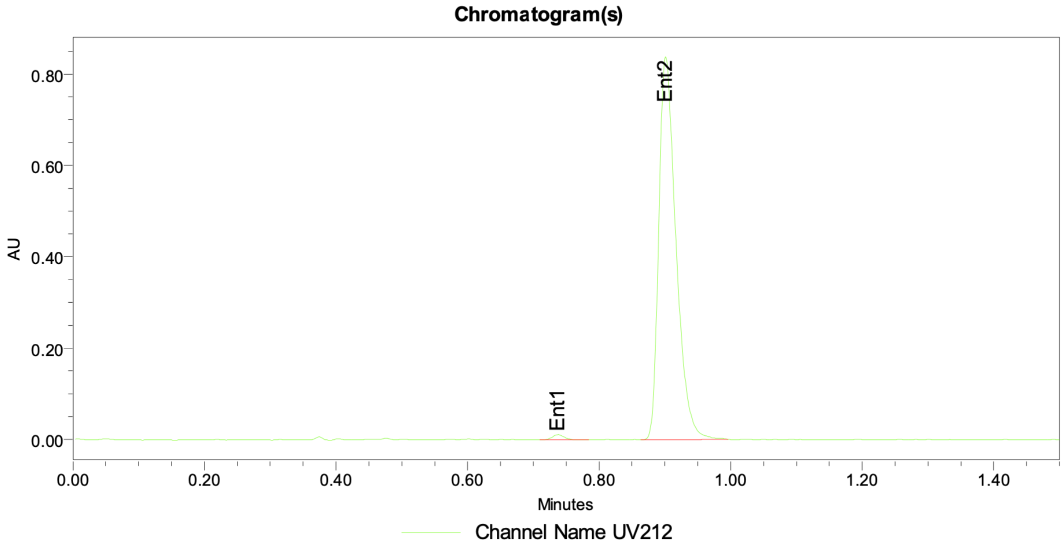


Peak Info

| | Channel Name | Name | RT | Area | Height (μV) | ent1 | ent2 | ee |
|---|---|---|---|---|---|---|---|---|
| 1 | UV212 | Ent1 | 0.74 | 15092 | 11612 | 0.98 | 99.02 | -98.03 |
| 2 | UV212 | Ent2 | 0.90 | 1517881 | 839311 | 0.98 | 99.02 | -98.03 |

**Figure S122:** SFC trace of **55**.

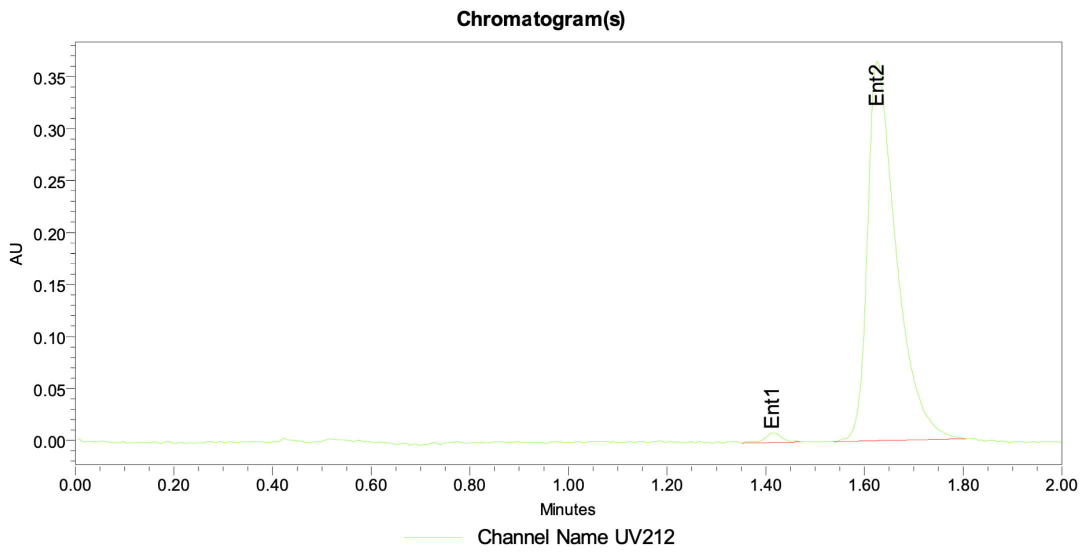


Peak Info

| | Channel Name | Name | RT | Area | Height (µV) | ent1 | ent2 | ee |
|---|---|---|---|---|---|---|---|---|
| 1 | UV212 | Ent1 | 1.42 | 22405 | 9341 | 1.49 | 98.51 | -97.02 |
| 2 | UV212 | Ent2 | 1.63 | 1483700 | 365076 | 1.49 | 98.51 | -97.02 |

**Figure S123:** SFC trace of (±)-**58**.

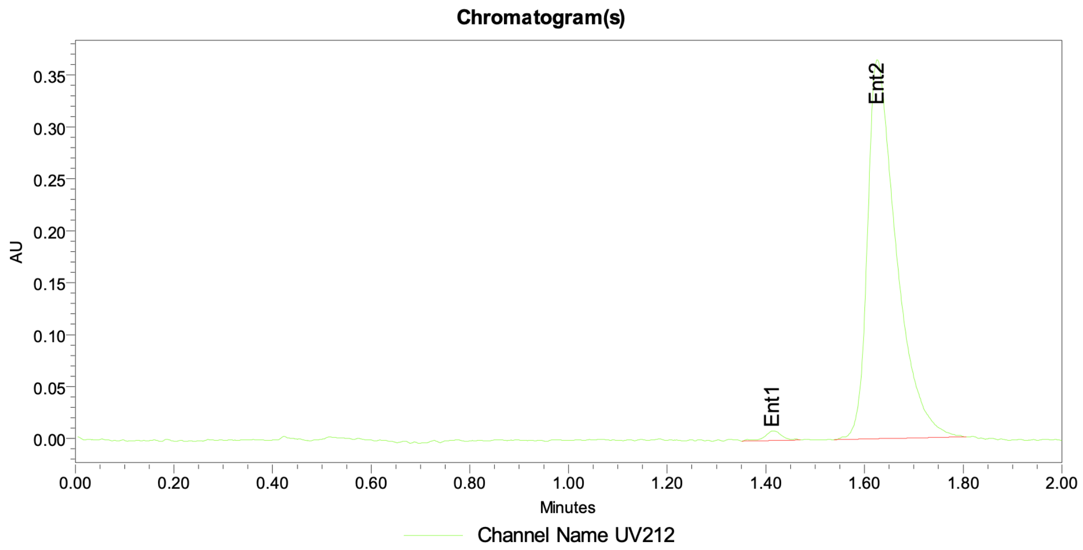


Peak Info

| | Channel Name | Name | RT | Area | Height (µV) | ent1 | ent2 | ee |
|---|---|---|---|---|---|---|---|---|
| 1 | UV212 | Ent1 | 1.42 | 22405 | 9341 | 1.49 | 98.51 | -97.02 |
| 2 | UV212 | Ent2 | 1.63 | 1483700 | 365076 | 1.49 | 98.51 | -97.02 |

**Figure S124:** SFC trace of **58**.

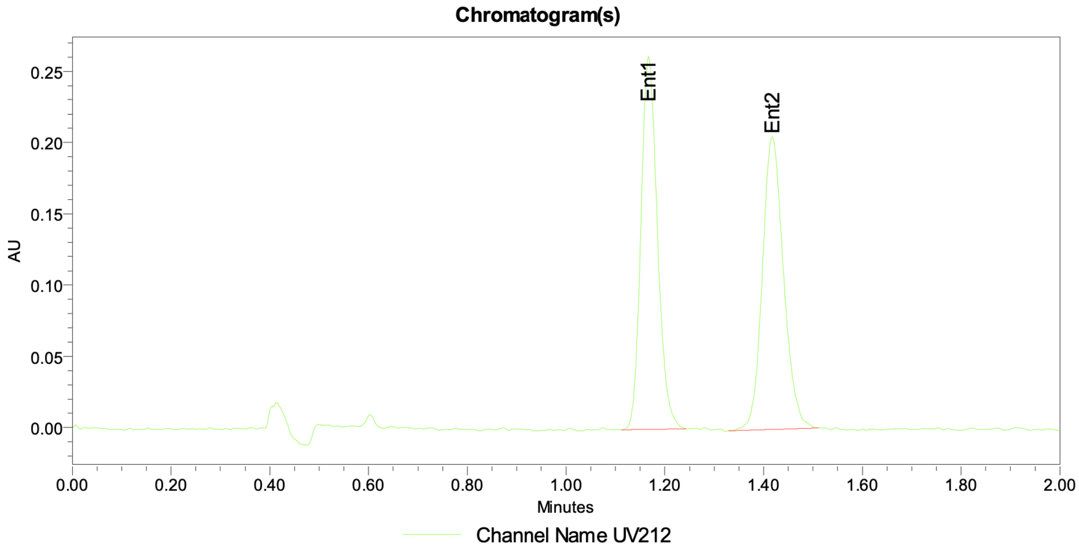


Peak Info

| | Channel Name | Name | RT | Area | Height (µV) | ent1 | ent2 | ee |
|---|---|---|---|---|---|---|---|---|
| 1 | UV212 | Ent1 | 1.17 | 606916 | 261793 | 49.75 | 50.25 | -0.50 |
| 2 | UV212 | Ent2 | 1.42 | 613025 | 205324 | 49.75 | 50.25 | -0.50 |

**Figure S125:** SFC trace of (±)-**61**.

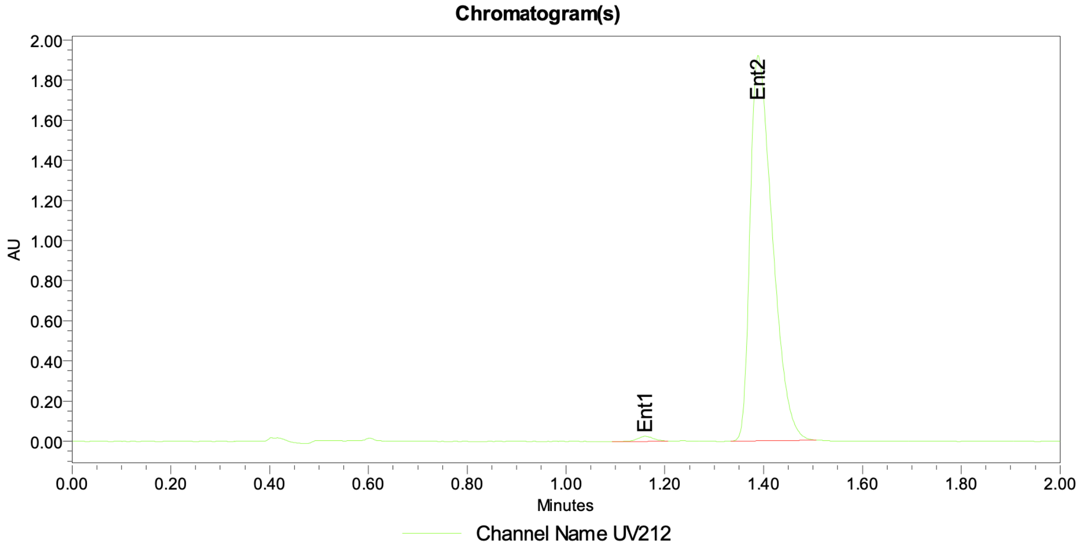


Peak Info

| | Channel Name | Name | RT | Area | Height (µV) | ent1 | ent2 | ee |
|---|---|---|---|---|---|---|---|---|
| 1 | UV212 | Ent1 | 1.16 | 56898 | 24684 | 0.94 | 99.06 | -98.12 |
| 2 | UV212 | Ent2 | 1.39 | 6010575 | 1921909 | 0.94 | 99.06 | -98.12 |

**Figure S126:** SFC trace of **61**.

#### References

1. Goetzke, F. W., Mortimore, M. & Fletcher, S. P. Enantio- and Diastereoselective Suzuki–Miyaura Coupling with Racemic Bicycles. *Angew. Chem. Int. Ed.* **58**, 12128–12132 (2019).
2. Mishra, S., Modicom, F. C. T., Dean, C. L. & Fletcher, S. P. Catalytic asymmetric synthesis of carbocyclic C-nucleosides. *Commun. Chem.* **5**, 154 (2022).
