## Supplementary Tables and Figures for "Complexoform-restricted covalent TRMT112 ligands that allosterically agonize METTL5"

**Supplementary Table 1**: Data collection and refinement statistics (molecular replacement).

| **TRMT112-METTL5 bound to SAM and FWG-33B** | |
|---|---|
| **PDB ID 9OHL** | |
| **Data collection** | |
| Wavelength (Å) | 1.00005 |
| Resolution (Å) | 1.3 |
| Space group | P212121 |
| Unit cell (Å) | 56.7 70.6 84.6 |
| Unique reflections | 801777 (5110) |
| Multiplicity | 5.8 (3.1) |
| Completeness | 93.2 (60.3) |
| Mean I/sigma(I) | 9.6 (0.9) |
| Wilson B-factor | 12 |
| Rsym | 0.10 (0.94) |
| Rpim | 0.04 (0.60) |
| CC1/2 | 0.99 (0.33) |
| **Refinement** | |
| Reflections using in refinement | 80172 (5109) |
| Reflections used for Rfree | 4107 (245) |
| R-work | 0.171 |
| R-free | 0.205 |
| RMS (bonds, Å) | 0.24 |
| RMS (angles, °) | 4.31 |
| Ramachandran allowed (%) | 100 |
| Ramachandran outliers (%) | 0 |
| Number of non-hydrogen atoms | 3381 |
| Protein | 2596 |
| ligand | 63 |
| solvent | 722 |
| Average B-factor | 18 |
| Protein | 14 |
| Ligand | 14 |
| Water | 32 |

*values in parenthesis include only reflections from the highest resolution shell

**Supplementary Table 2**: ORFs used in this manuscript.

| | |
|---|---|
| WT-TRMT112-FLAG<br>(*H. sapiens*,<br>codon optimized for mammalian expression) | ATGAAACTGCTGACCCACAACCTGCTCAGCAGCCACG<br>TGCGGGGCGTGGGCAGCAGAGGATTTCCACTGAGACT<br>GCAGGCTACAGAGGTGCGGATCTGCCCCGTCGAGTTC<br>AACCCCAACTTCGTGGCCAGAATGATCCCCAAGGTGG<br>AATGGTCCGCCTTCCTGGAGGCCGCCGACAACCTGAG<br>GCTGATCCAGGTGCCTAAGGGCCCTGTGGAAGGCTAC<br>GAGGAAAACGAGGAATTCCTGCGGACCATGCATCACC<br>TGCTGCTGGAAGTTGAGGTGATCGAGGGAACACTGCA<br>GTGTCCTGAGAGCGGCAGAATGTTCCCTATTTCTAGAG<br>GCATCCCTAATATGCTGCTGAGCGAGGAAGAAACCGA<br>GTCTGGCTCGGGCTCGGGTGATTACAAAGACGATGAC<br>GATAAGTGA |
| WT-HA-N6AMT1<br>(*H. sapiens*) | ATGTACCCATACGATGTTCCAGATTACGCTGGTACCAT<br>GGCAGGGGAGAACTTCGCTACGCCGTTCCACGGGCA<br>CGTGGGCCGCGGCGCCTTCAGCGACGTGTACGAGCC<br>CGCGGAGGACACGTTTCTGCTTTTGAACGCGCTGGAG<br>GCAGCGGCTGCCGAACTGGCAGGAGTGGAAATATGCC<br>TGGAAGTAGGGTCAGGGTCTGGTGTAGTATCTGCATT<br>CCTAGCCTCTATGATAGGCCCTCAGGCTTTGTACATGT<br>GCACTGATATCAACCCTGAGGCAGCAGCTTGTACCCTA<br>GAGACAGCACGCTGTAACAAAGTTCACATTCAACCAGT<br>TATTACAGATTTGGTCAAAGGCTTGCTACCAAGATTGA<br>CCGAAAAAGTTGATCTTCTGGTGTTTAATCCCCCCTAT<br>GTAGTGACTCCACCTCAAGAGGTAGGAAGTCACGGAA<br>TAGAGGCAGCTTGGGCTGGTGGCAAAAATGGTCGGGA<br>AGTCATGGACAGGTTTTTTCCCCTGGTTCCAGATCTCC<br>TTTCACCAAAAGGATTATTCTATTTAGTTACCATTAAAG<br>AAAACAACCCAGAAGAAATTTTGAAAATAATGAAGACA<br>AAAGGTCTGCAAGGAACCACTGCACTTTCCAGACAAG |

| | |
|---|---|
| | CAGGCCAAGAAACTCTTTCAGTCCTCAAGTTCACCAAG TCTTAG |
| WT-HA-METTL5 (*H. sapiens*) | ATGTACCCATACGATGTTCCAGATTACGCTGGTACCAT GAAGAAAGTAAGGCTTAAGGAACTAGAGAGTCGCCTG CAACAAGTGGATGGATTTGAAAAGCCCAAGCTACTTCT GGAACAGTATCCTACCAGGCCGCACATTGCAGCATGT ATGCTCTATACAATCCATAACACTTATGATGACATTGAA AATAAAGTCGTTGCAGATCTAGGATGTGGTTGTGGAGT ACTTAGCATCGGAACTGCAATGTTAGGAGCAGGGTTGT GTGTTGGATTTGACATAGATGAAGACGCATTGGAAATA TTTAATAGGAATGCAGAAGAGTTTGAGTTAACAAATATT GACATGGTTCAATGTGATGTGTGCTTATTATCTAACAG AATGTCCAAGTCATTCGATACAGTAATTATGAATCCTCC CTTTGGGACCAAAAATAATAAAGGGACAGATATGGCTT TTCTAAAGACTGCTTTGGAAATGGCAAGAACAGCAGTA TATTCCTTACACAAATCCTCAACTAGAGAACATGTTCAA AAGAAAGCTGCAGAATGGAAAATCAAGATAGATATTAT AGCAGAACTTCGATATGACCTGCCAGCATCATACAAGT TTCACAAAAAGAAATCAGTGGACATTGAAGTGGACCTA ATTCGGTTTTCCTTTTGA |
| WT-HA-BUD23 (*H. sapiens*) | ATGTACCCATACGATGTTCCAGATTACGCTGGTACCAT GGCGTCCCGCGGCCGGCGTCCGGAGCATGGCGGACC CCCAGAGCTGTTTTATGACGAGACAGAAGCCCGGAAA TACGTTCGCAACTCACGGATGATTGATATCCAGACCAG GATGGCTGGGCGAGCATTGGAGCTTCTTTATCTGCCA GAGAATAAGCCCTGTTACCTGCTGGATATTGGCTGTGG CACTGGGCTGAGTGGAAGTTATCTGTCAGATGAAGGG CACTATTGGGTGGGCCTGGATATCAGCCCTGCCATGC TGGATGAGGCTGTGGACCGAGAGATAGAGGGAGACCT |

| | |
|---|---|
| | GCTGCTGGGGGATATGGGCCAGGGCATCCCATTCAAG CCAGGCACATTTGATGGTTGCATCAGCATTTCTGCTGT GCAGTGGCTCTGTAATGCTAACAAGAAGTCTGAAAACC CTGCCAAGCGCCTGTACTGCTTTTTTGCTTCTCTTTTTT CTGTTCTCGTCCGGGGATCCCGAGCTGTCCTGCAGCT GTACCCTGAGAACTCAGAGCAGTTGGAGCTGATCACA ACCCAGGCCACAAAGGCAGGCTTCTCCGGTGGCATGG TGGTAGACTACCCTAACAGTGCCAAAGCAAAGAAATTC TACCTCTGCTTGTTTTCTGGGCCTTCGACCTTTATACCA GAGGGGCTGAGTGAAAATCAGGATGAAGTTGAACCCA GGGAGTCTGTGTTCACCAATGAGAGGTTCCCATTAAG GATGTCGAGGCGGGGAATGGTGAGGAAGAGTCGGGC ATGGGTGCTGGAGAAGAAGGAGCGGCACAGGCGCCA GGGCAGGGAAGTCAGACCTGACACCCAGTACACCGG CCGCAAGCGCAAGCCCCGCTTCTAA |
| His-METTL5 (*H. sapiens*, codon optimized for *E. coli* expression) | ATGCACCACCACCACCACCACGAAAATTTGTATTTCCA GTCAATGAAGAAGGTACGTCTGAAGGAGCTTGAATCC CGTCTTCAGCAAGTGGACGGTTTTGAGAAACCGAAGC TGTTGCTGGAGCAGTATCCAACGCGCCCGCATATCGC AGCTTGTATGCTGTATACTATCCACAACACCTATGACG ACATTGAGAATAAAGTAGTCGCAGATTTGGGTTGTGGA TGCGGAGTTCTGAGTATCGGGACGGCGATGCTTGGAG CCGGTTTGTGTGTGGGGTTCGATATCGATGAGGACGC ACTGGAAATTTTTAACCGCAACGCGGAAGAGTTTGAAT TGACTAACATTGACATGGTGCAATGCGACGTTTGTCTT TTAAGTAATCGTATGAGCAAGTCATTTGACACTGTTATT ATGAACCCACCTTTCGGAACTAAAAATAACAAAGGGAC CGATATGGCGTTCCTGAAGACGGCCCTTGAGATGGCC CGCACAGCCGTATATTCCCTTCACAAGAGCTCTACCCG TGAACACGTTCAGAAGAAAGCCGCTGAGTGGAAGATT |

| | |
|---|---|
| | AAGATTGACATTATCGCTGAGCTTCGTTATGACCTGCC<br>AGCCTCATATAAATTTCATAAAAAGAAGTCCGTGGATAT<br>TGAAGTTGATTTGATCCGCTTCAGTTTTTGA |
| TRMT112<br>(*H. sapiens*,<br>codon optimized for *E. coli* expression) | ATGAAGCTGCTGACGCACAATTTACTTTCATCACATGT<br>CCGCGGAGTCGGCTCCCGTGGGTTTCCGCTGCGCTTA<br>CAAGCAACTGAAGTCCGCATCTGTCCTGTCGAATTTAA<br>TCCTAATTTCGTCGCTCGTATGATCCCAAAAGTTGAGT<br>GGAGTGCATTTCTGGAAGCAGCGGACAACCTGCGTCT<br>GATTCAGGTCCCTAAAGGACCTGTCGAAGGATACGAA<br>GAAAACGAAGAATTTTTACGCACCATGCACCATCTGTT<br>ACTGGAAGTCGAGGTGATCGAGGGAACTTTGCAGTGC<br>CCAGAATCTGGGCGCATGTTCCCCATTTCCCGCGGCA<br>TCCCTAACATGCTTCTTTCCGAAGAGGAGACCGAATCT<br>TGA |

**Supplementary Table 3**: Antibodies used in this study.

| Primary antibodies | Species | Supplier | Dilution |
|---|---|---|---|
| FLAG® (HRP) (M2) | N/A, monoclonal | Sigma-Aldrich: A8592 | 1:5,000 |
| β-Actin Antibody (C4) | N/A, monoclonal | Santa Cruz: sc-47778 | 1:5,000 |
| GAPDH (HRP) (D16H11) | N/A, monoclonal | Cell signaling: 8884 | 1:2,000 |
| HA (HRP) (6E2) | N/A, monoclonal | Cell signaling: 2999 | 1:2,000 |
| Vinculin (E1E9V) (HRP) | N/A, monoclonal | Cell signaling: 18799 | 1:2,000 |
| TRMT112 (F-7) | mouse, monoclonal | Santa Cruz: sc-398481 | 1:1,000 |
| METTL5 | rabbit, polyclonal | Proteintech: 16791-1-AP | 1:2,000 |
| BUD23 | rabbit, polyclonal | Proteintech: 28192-1-AP | 1:3,000 |
| THUMPD3 | rabbit, polyclonal | Proteintech: 19807-1-AP | 1:2,000 |
| N6AMT1 | rabbit, polyclonal | Proteintech: 16211-1-AP | 1:1,000 |
| TRMT11 | rabbit, polyclonal | Proteintech: 17555-1-AP | 1:1,000 |
| THUMPD2 (D1) | mouse, monoclonal | Santa Cruz: sc-393018 | 1:500 |
| ALKBH8 | rabbit, polyclonal | Sigma-Aldrich: HPA061514 | 1:500 |

| Secondary antibodies | Species | Supplier | Dilution |
|---|---|---|---|
| anti-mouse IgG (HRP) | N/A, monoclonal | Cell signaling: 7076 | 1:2,000 |
| anti-rabbit IgG (HRP) | N/A, monoclonal | Santa Cruz: sc-2357 | 1:5,000 |

**Supplementary Table 4:** Fractionation of TMT$^{16}$-plex into 10 fractions.

| Fraction Number | Acetonitrile (%) | Combined into Fraction |
|---|---|---|
| 1 | 7.5 | 1 |
| 2 | 10.0 | 2 |
| 3 | 12.5 | 3 |
| 4 | 15.0 | 4 |
| 5 | 17.5 | 5 |
| 6 | 20.0 | 6 |
| 7 | 22.5 | 7 |
| 8 | 25.0 | 8 |
| 9 | 27.5 | 9 |
| 10 | 30.0 | 10 |
| 11 | 32.5 | 1 |
| 12 | 35.0 | 2 |
| 13 | 37.5 | 3 |
| 14 | 40.0 | 4 |
| 15 | 42.5 | 5 |
| 16 | 45.0 | 6 |
| 17 | 47.5 | 7 |
| 18 | 50.0 | 8 |
| 19 | 52.5 | 9 |
| 20 | 55.0 | 10 |
| 21 | 57.5 | 1 |
| 22 | 60.0 | 2 |
| 23 | 62.5 | 3 |
| 24 | 65.0 | 4 |
| 25 | 67.5 | 5 |
| 26 | 70.0 | 6 |
| 27 | 72.5 | 7 |

| | | |
|---|---|---|
| 28 | 75.0 | 8 |
| 29 | 80.0 | 9 |
| 30 | 95.0 | 10 |

**Supplementary Table 5:** Fractionation of TMT$^{10}$-plex into 5 fractions.

| Fraction Number | Acetonitrile (%) | Combined into Fraction |
|---|---|---|
| 1 | 7.5 | 1 |
| 2 | 10.0 | 2 |
| 3 | 12.5 | 3 |
| 4 | 15.0 | 4 |
| 5 | 17.5 | 5 |
| 6 | 20.0 | 1 |
| 7 | 25.0 | 2 |
| 8 | 30.0 | 3 |
| 9 | 35.0 | 4 |
| 10 | 40.0 | 5 |
| 11 | 45.0 | 1 |
| 12 | 50.0 | 2 |
| 13 | 55.0 | 3 |
| 14 | 80.0 | 4 |
| 15 | 95.0 | 5 |

**Supplementary Table 6:** Fractionation of TMT$^{16}$-plex into 3 fractions.

| Fraction Number | Acetonitrile (%) | Combined into Fraction |
|---|---|---|
| 1 | 7.5 | 1 |
| 2 | 12.5 | 2 |
| 3 | 15.0 | 3 |
| 4 | 20.0 | 1 |
| 5 | 25.0 | 2 |
| 6 | 30.0 | 3 |
| 7 | 35.0 | 1 |
| 8 | 60.0 | 2 |
| 9 | 95.0 | 3 |

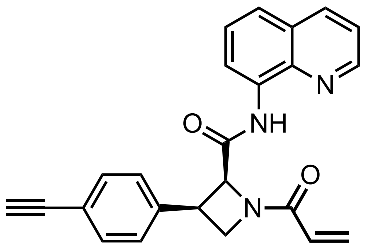

**MY-11A**

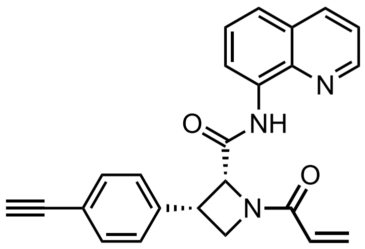

**MY-11B**

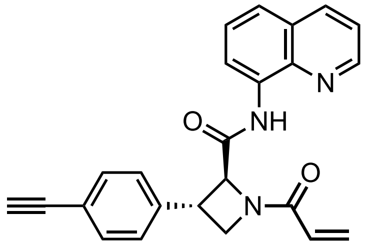

**MY-12A**

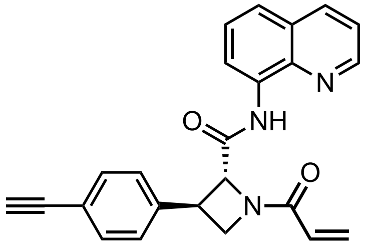

**MY-12B**

**Supplementary Fig. 1:** Structures of alkyne (FWG-MY-11A, 11B, 12A, and 12B) azetidine acrylamide stereoprobes.

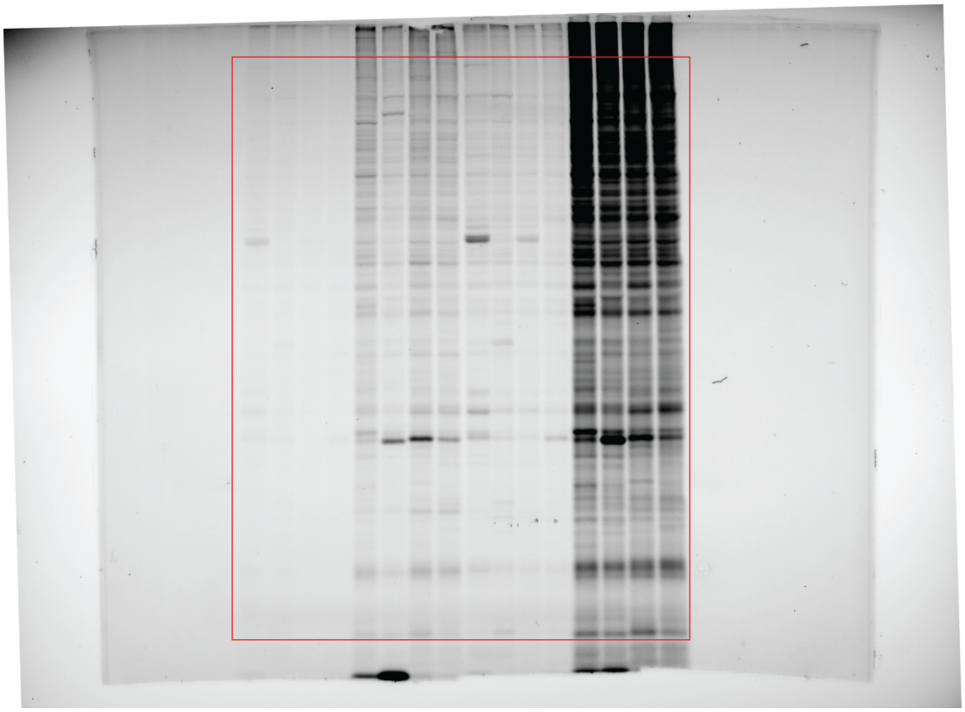

**Supplementary Fig. 2a:** Uncropped image of Rhodamine scan in reference to Fig. 1b (ABPP).

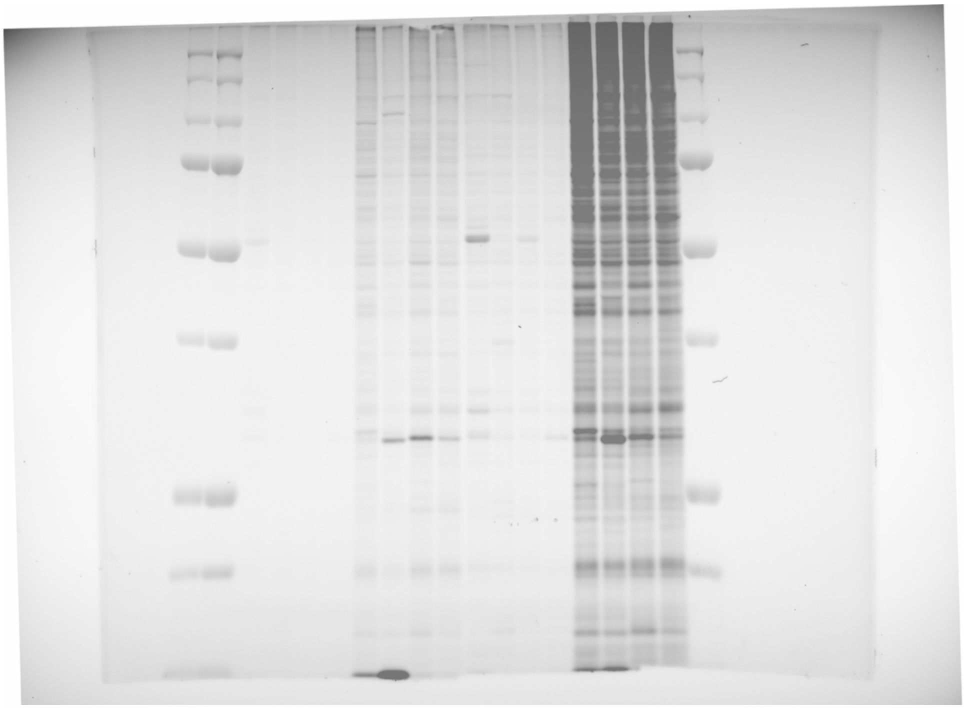

**Supplementary Fig. 2b:** Uncropped composite image of Cy5 and Rhodamine scan in reference to Fig. 1b (ABPP).

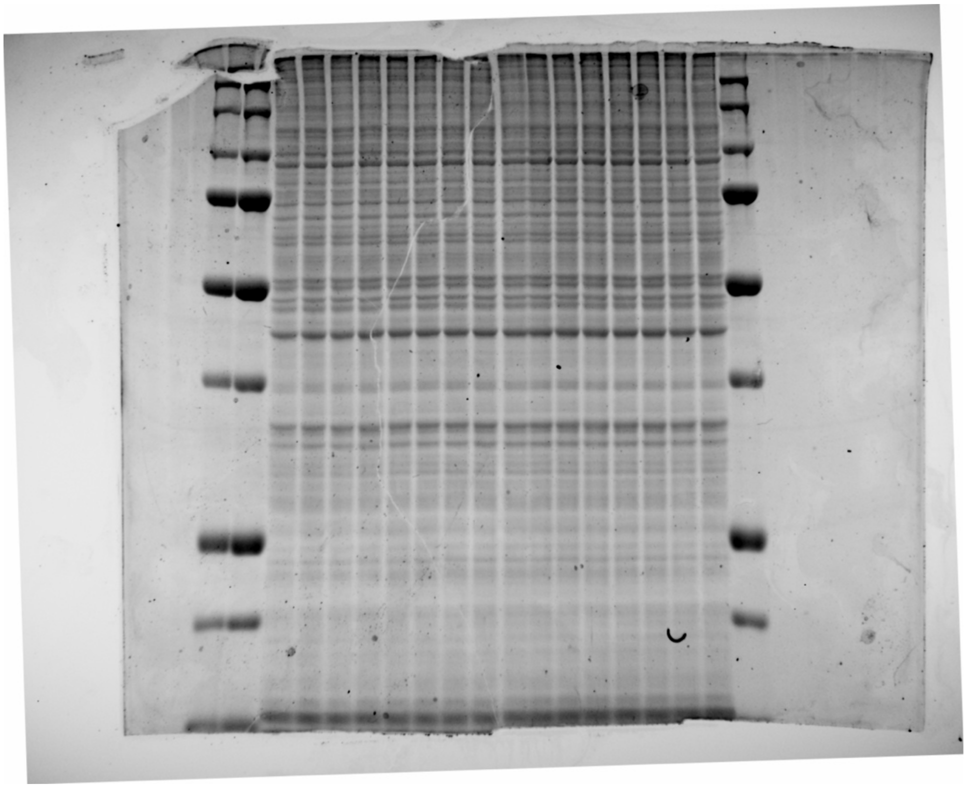

**Supplementary Fig. 3:** Uncropped image of Coomassie scan in reference to Fig. 1b (Coomassie).

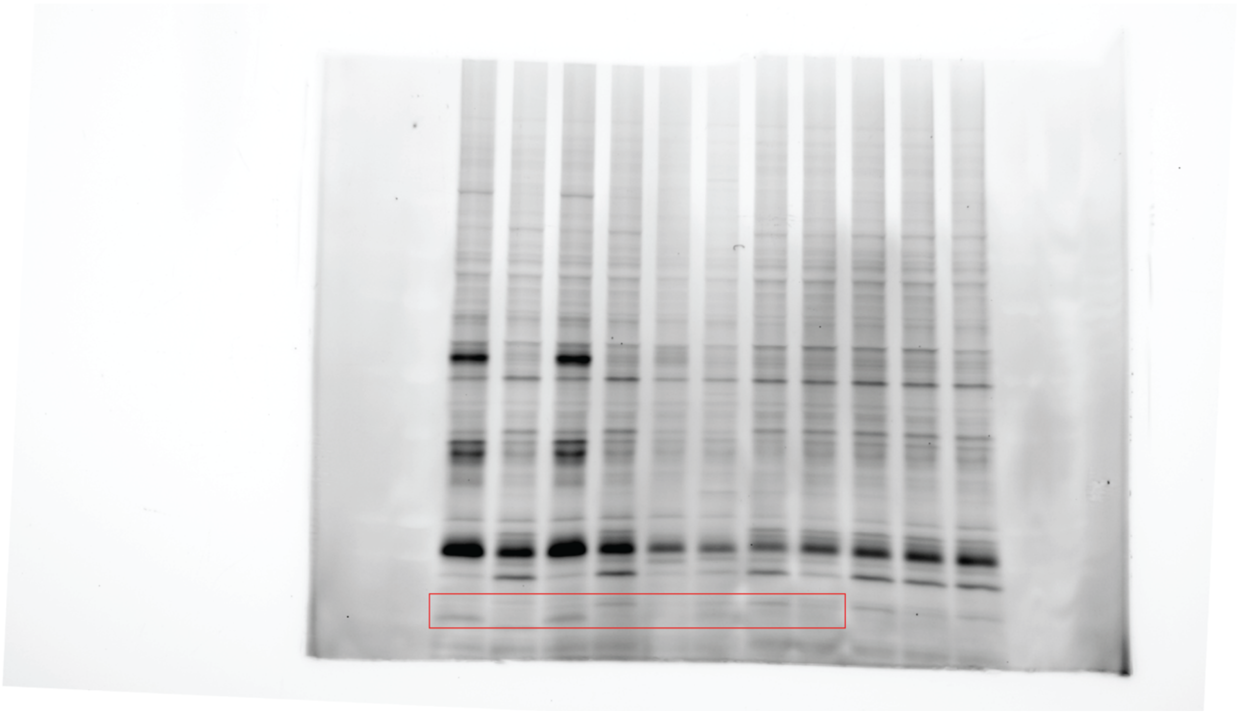

**Supplementary Fig. 4a:** Uncropped image of Rhodamine scan in reference to Fig. 2b (ABPP, input).

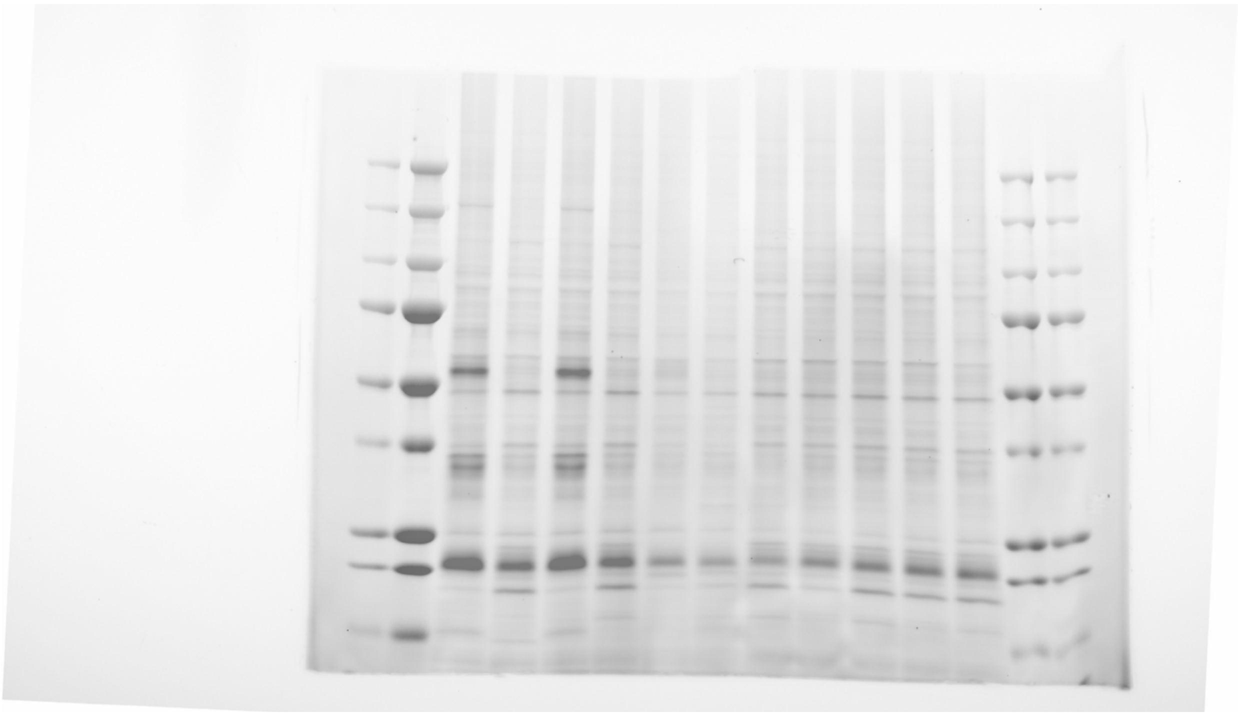

**Supplementary Fig. 4b:** Uncropped composite image of Cy5 and Rhodamine scan in reference to Fig. 2b (input, ABPP).

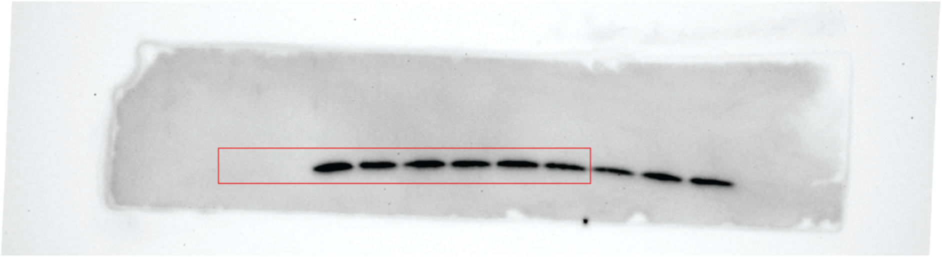

**Supplementary Fig. 5a:** Uncropped image of Chemiluminescence scan in reference to Fig. 2b (FLAG, input).

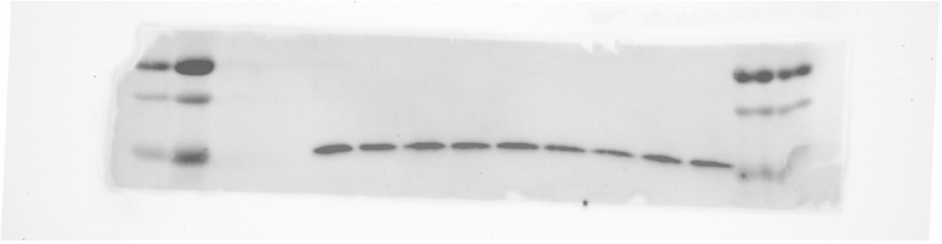

**Supplementary Fig. 5b:** Uncropped composite image of Cy5 and Chemiluminescence scan in reference to Fig. 2b (FLAG, input).

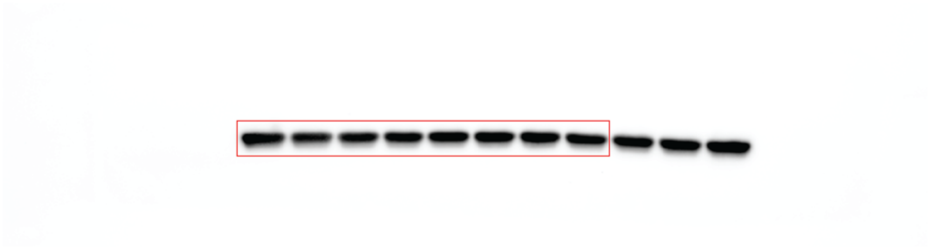

**Supplementary Fig. 6a:** Uncropped image of Chemiluminescence scan in reference to Fig. 2b (β-actin, input).

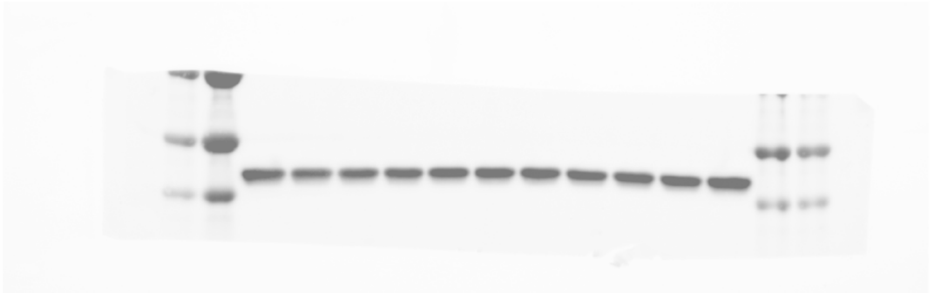

**Supplementary Fig. 6b:** Uncropped composite image of Cy5 and Chemiluminescence scan in reference to Fig. 2b (β-actin, input).

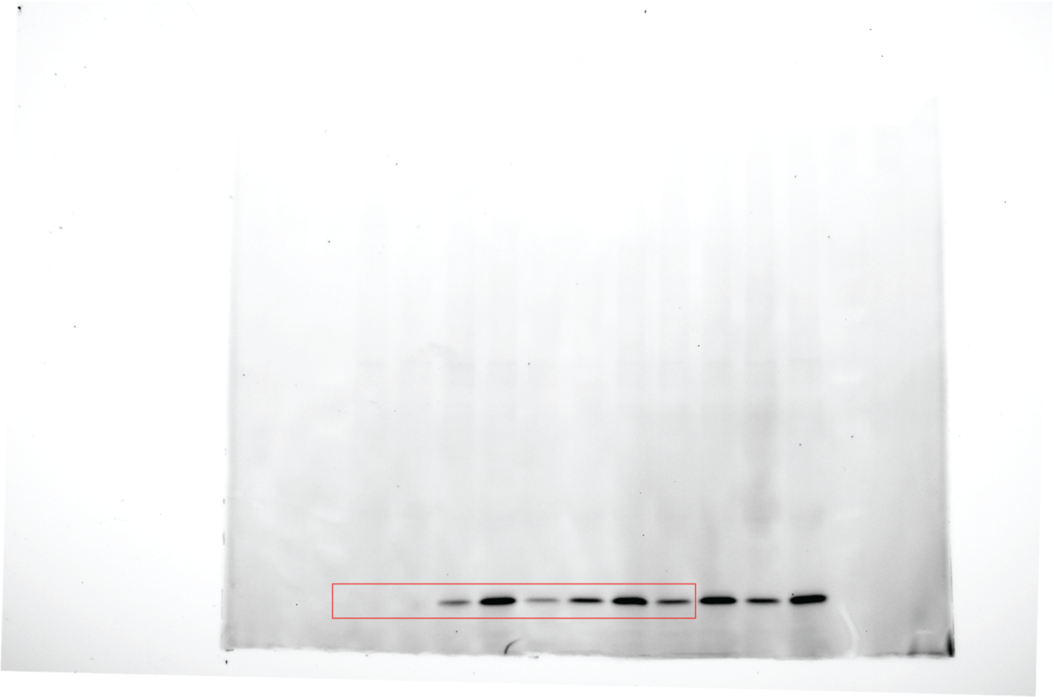

**Supplementary Fig. 7a:** Uncropped image of Rhodamine scan in reference to Fig. 2b (ABPP, IP).

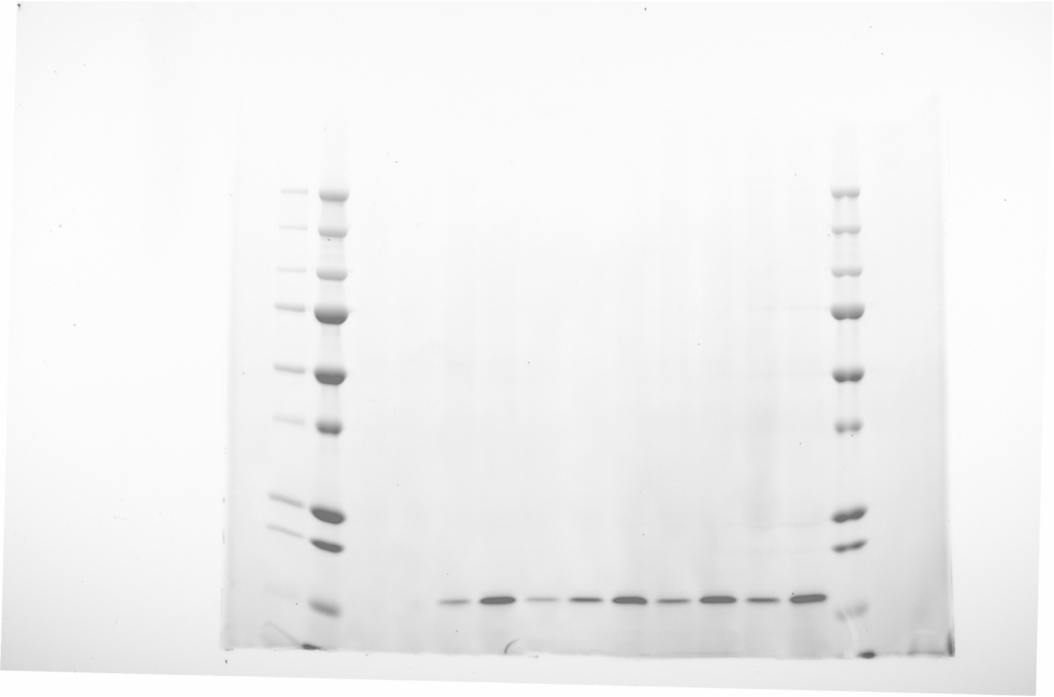

**Supplementary Fig. 7b:** Uncropped composite image of Cy5 and Rhodamine scan in reference to Fig. 2b (ABPP, IP).

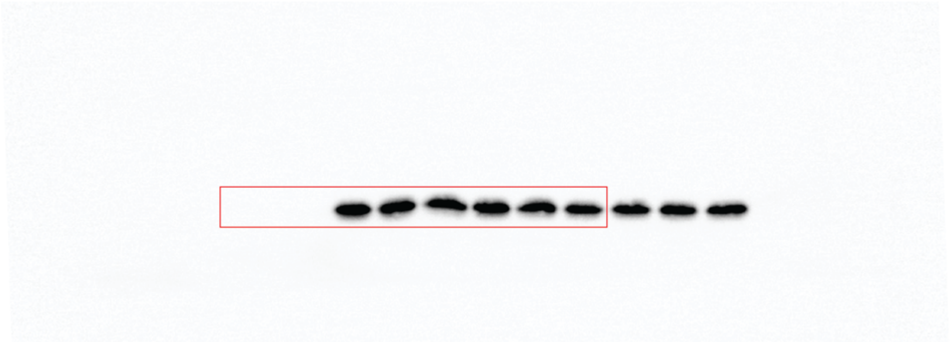

**Supplementary Fig. 8a:** Uncropped image of Chemiluminescence scan in reference to Fig. 2b (FLAG, IP).

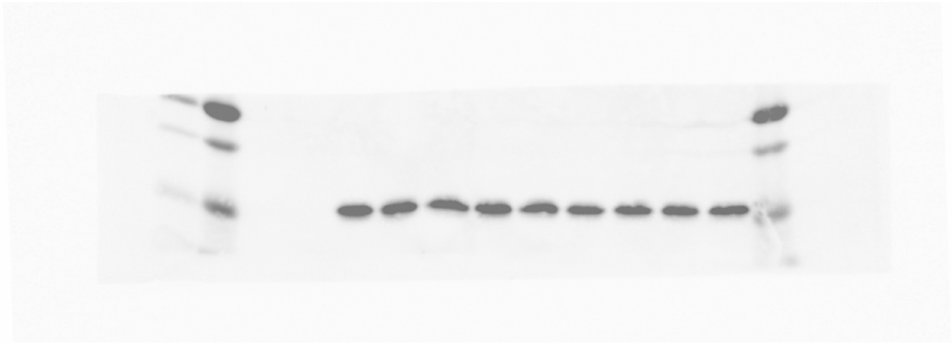

**Supplementary Fig. 8b:** Uncropped composite image of Cy5 and Chemiluminescence scan in reference to Fig. 2b (FLAG, IP).

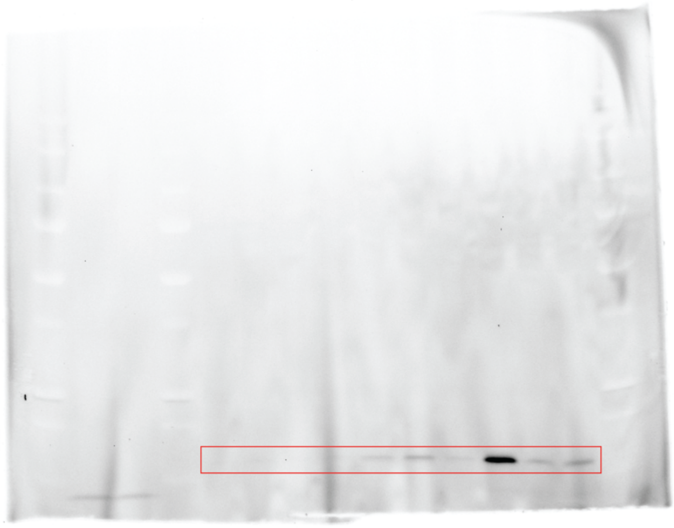

**Supplementary Fig. 9a:** Uncropped image of Rhodamine scan in reference to Fig. 2e (ABPP).

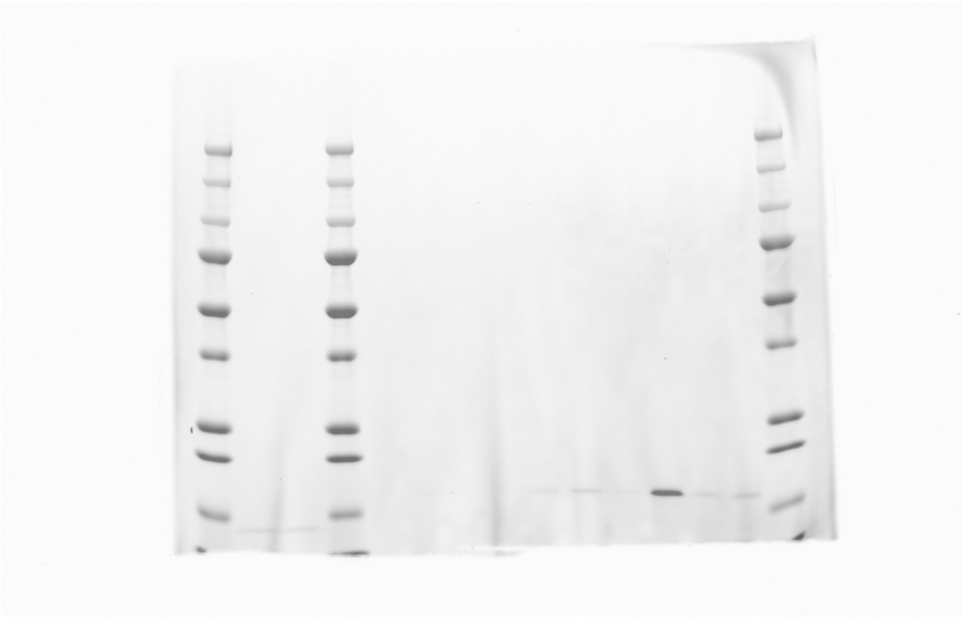

**Supplementary Fig. 9b:** Uncropped composite image of Cy5 and Rhodamine scan in reference to Fig. 2e (ABPP).

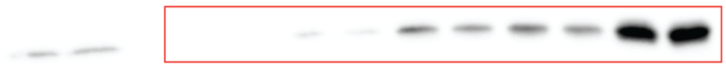

**Supplementary Fig. 10a:** Uncropped image of Chemiluminescence scan in reference to Fig. 2e (FLAG, short exposure).

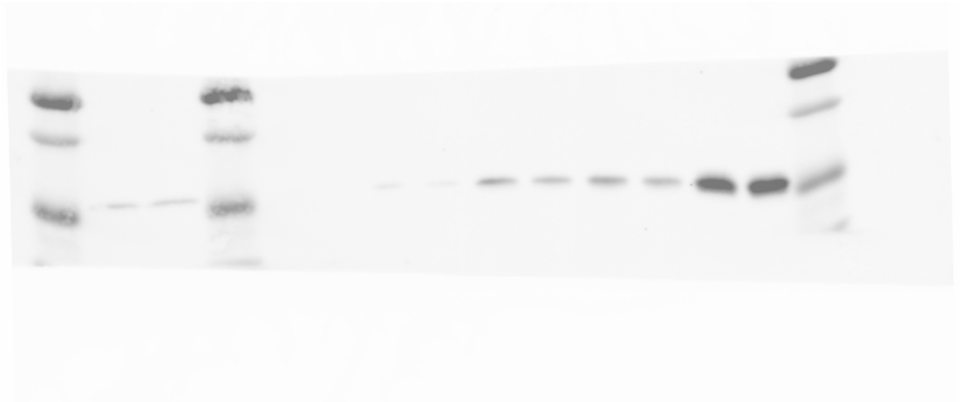

**Supplementary Fig. 10b:** Uncropped composite image of Cy5 and Chemiluminescence scan in reference to Fig. 2e (FLAG, short exposure).

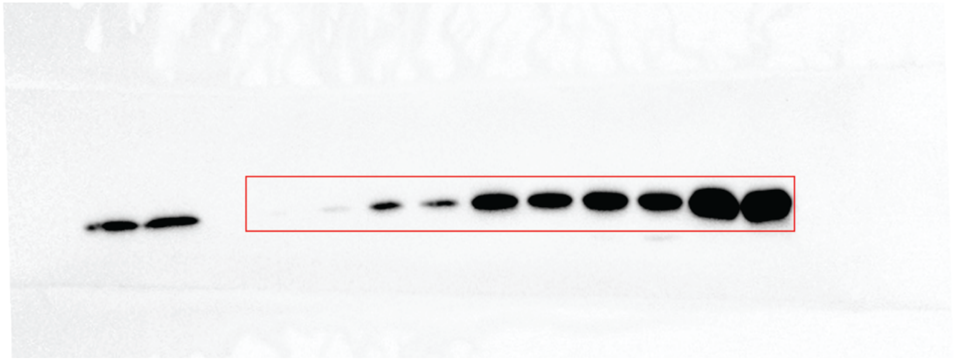

**Supplementary Fig. 11a:** Uncropped image of Chemiluminescence scan in reference to Fig. 2e (FLAG, long exposure).

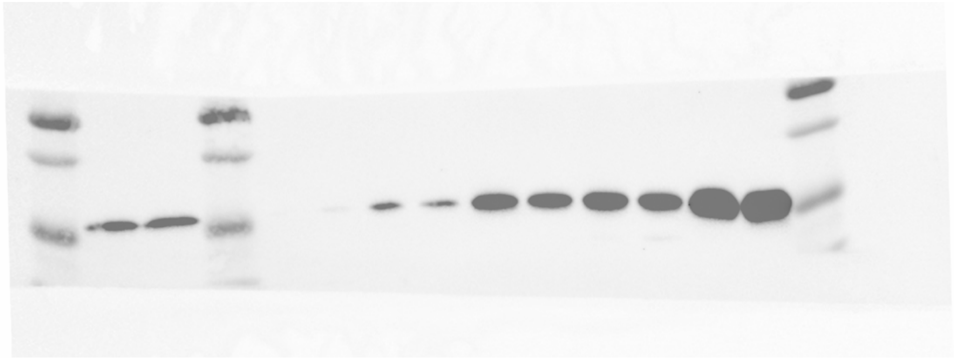

**Supplementary Fig. 11b:** Uncropped composite image of Cy5 and Chemiluminescence scan in reference to Fig. 2e (FLAG, long exposure).

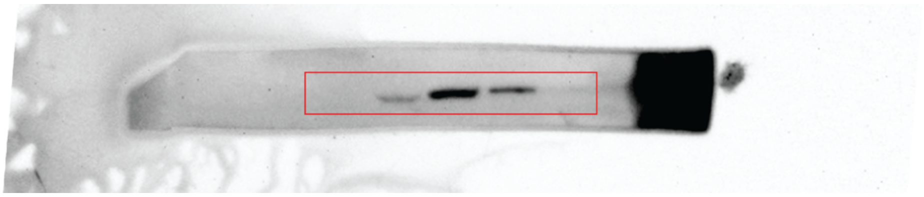

**Supplementary Fig. 12a:** Uncropped image of Chemiluminescence scan in reference to Fig. 2f (ALKBH8).

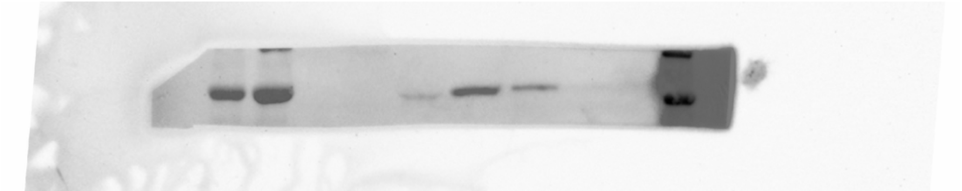

**Supplementary Fig. 12b:** Uncropped composite image of Cy5 and Chemiluminescence scan in reference to Fig. 2e (ALKBH8).

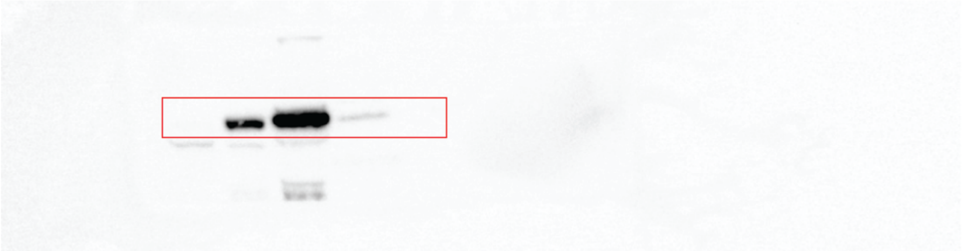

**Supplementary Fig. 13a:** Uncropped image of Chemiluminescence scan in reference to Fig. 2f (THUMPD3).

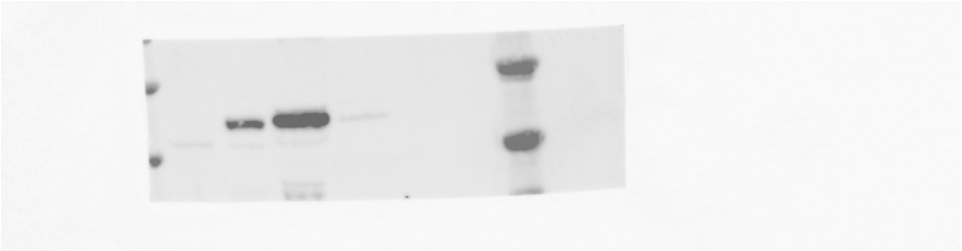

**Supplementary Fig. 13b:** Uncropped composite image of Cy5 and Chemiluminescence scan in reference to Fig. 2e (THUMPD3).

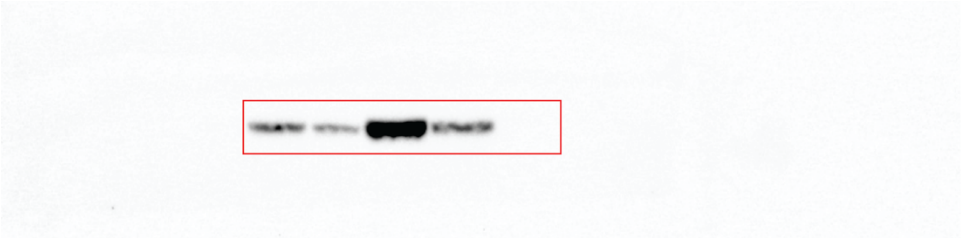

**Supplementary Fig. 14a:** Uncropped image of Chemiluminescence scan in reference to Fig. 2f (THUMPD2).

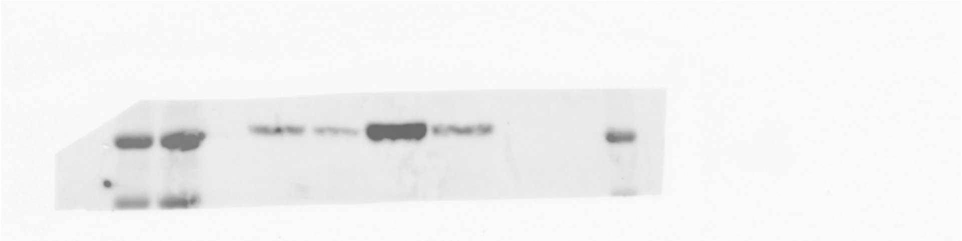

**Supplementary Fig. 14b:** Uncropped composite image of Cy5 and Chemiluminescence scan in reference to Fig. 2f (THUMPD2).

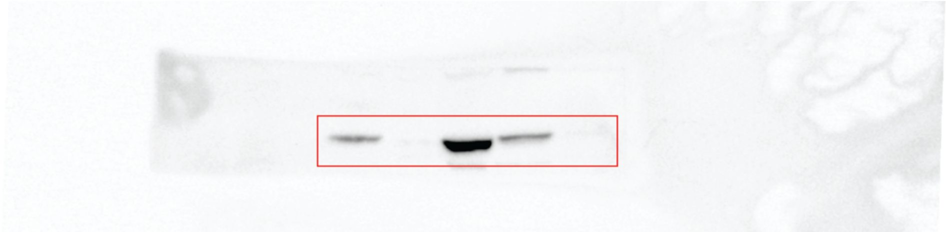

**Supplementary Fig. 15a:** Uncropped image of Chemiluminescence scan in reference to Fig. 2f (TRMT11).

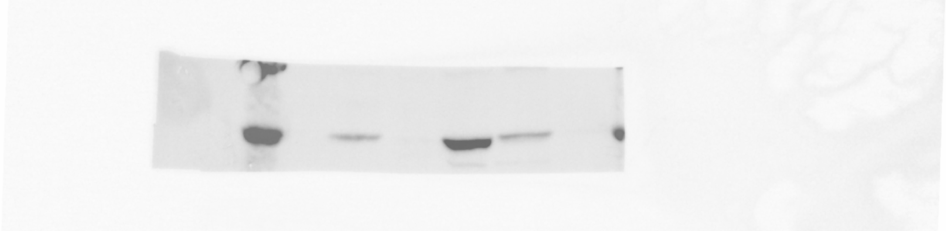

**Supplementary Fig. 15b:** Uncropped composite image of Cy5 and Chemiluminescence scan in reference to Fig. 2f (TRMT11).

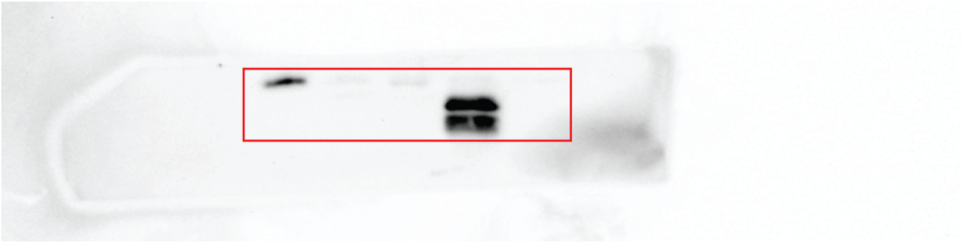

**Supplementary Fig. 16a:** Uncropped image of Chemiluminescence scan in reference to Fig. 2f (BUD23).

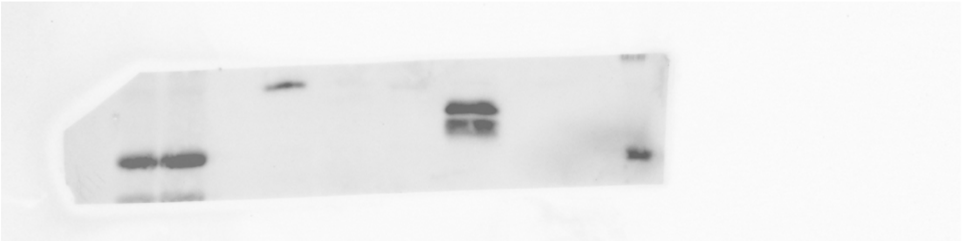

**Supplementary Fig. 16b:** Uncropped composite image of Cy5 and Chemiluminescence scan in reference to Fig. 2f (BUD23).

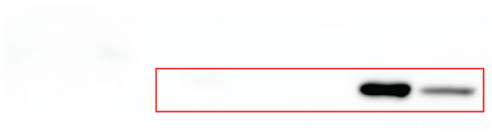

**Supplementary Fig. 17a:** Uncropped image of Chemiluminescence scan in reference to Fig. 2f (METTL5).

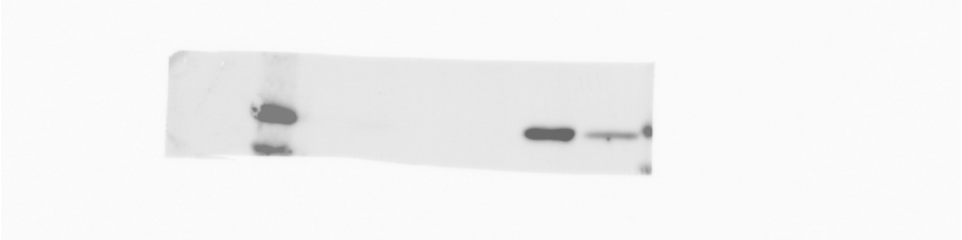

**Supplementary Fig. 17b:** Uncropped composite image of Cy5 and Chemiluminescence scan in reference to Fig. 2f (METTL5).

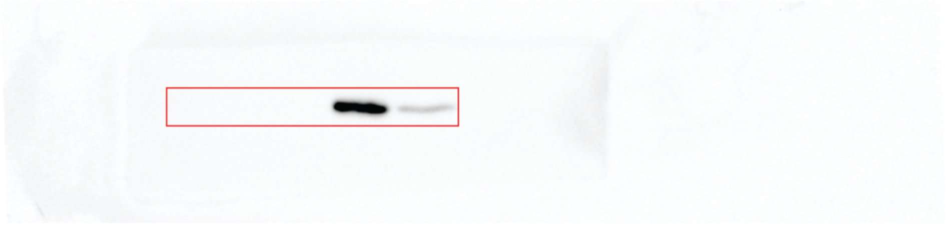

**Supplementary Fig. 18a:** Uncropped image of Chemiluminescence scan in reference to Fig. 2f (N6AMT1).

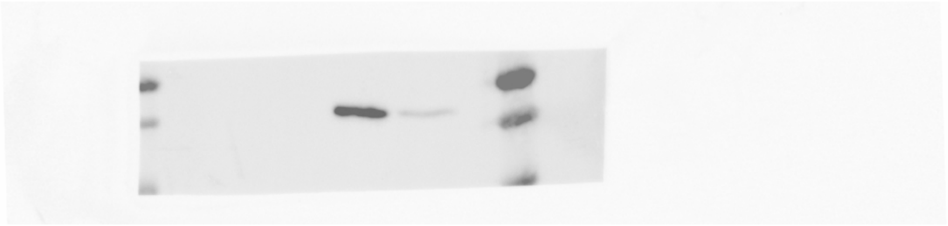

**Supplementary Fig. 18b:** Uncropped composite image of Cy5 and Chemiluminescence scan in reference to Fig. 2f (N6AMT1).

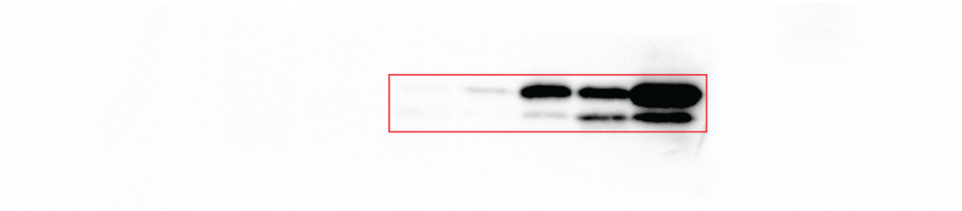

**Supplementary Fig. 19a:** Uncropped image of Chemiluminescence scan in reference to Fig. 2f (TRMT112).

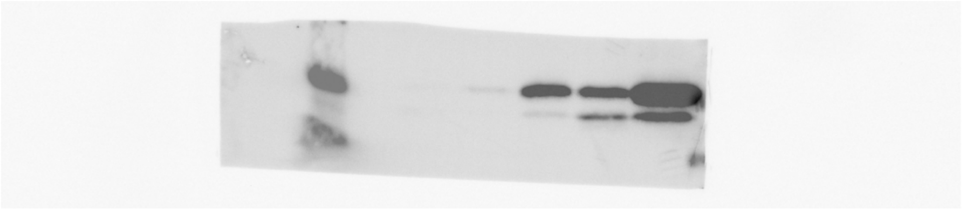

**Supplementary Fig. 19b:** Uncropped composite image of Cy5 and Chemiluminescence scan in reference to Fig. 2f (TRMT112).

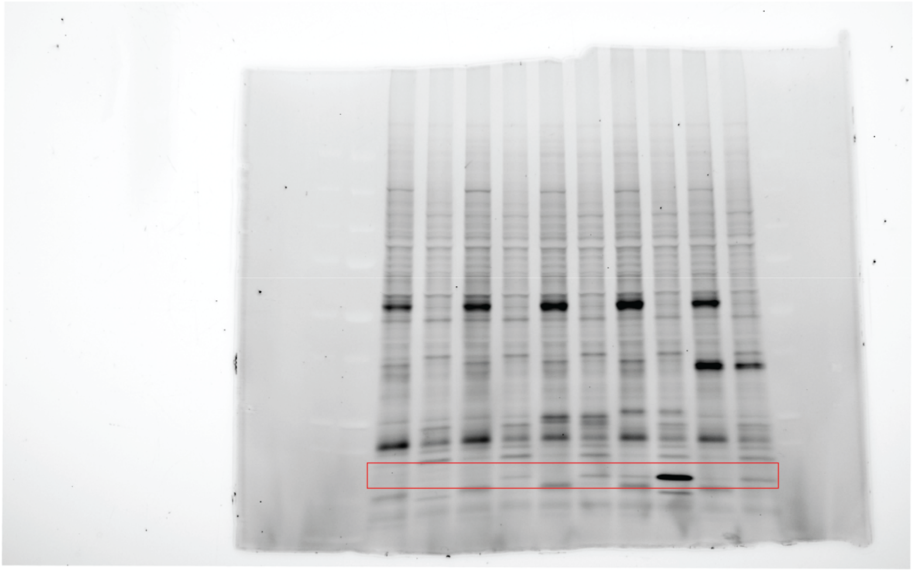

**Supplementary Fig. 20a:** Uncropped image of Rhodamine scan in reference to Fig. 2g (ABPP).

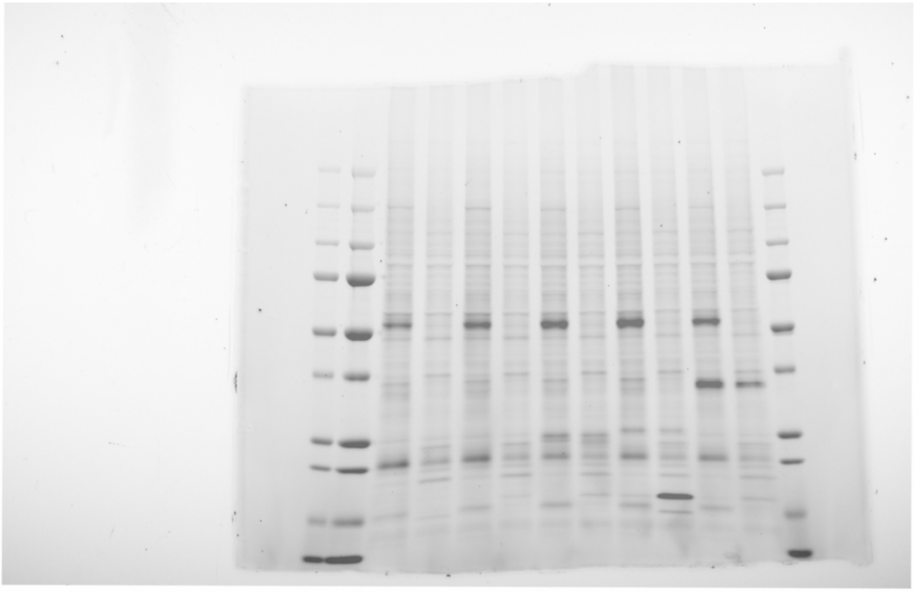

**Supplementary Fig. 20b:** Uncropped composite image of Cy5 and Rhodamine scan in reference to Fig. 2g (ABPP).

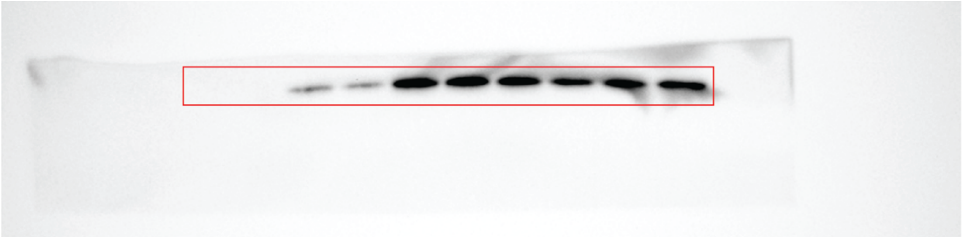

**Supplementary Fig. 21a:** Uncropped image of Chemiluminescence scan in reference to Fig. 2g (FLAG).

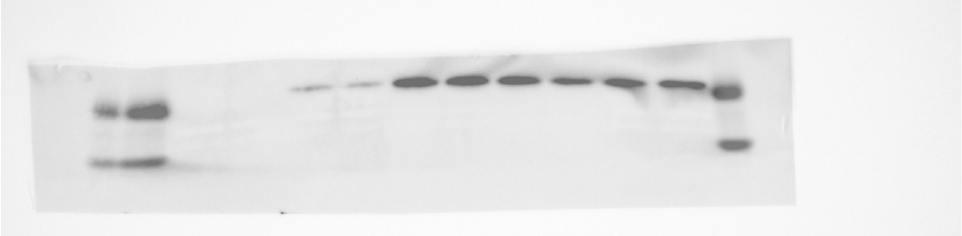

**Supplementary Fig. 21b:** Uncropped composite image of Cy5 and Chemiluminescence scan in reference to Fig. 2g (FLAG).

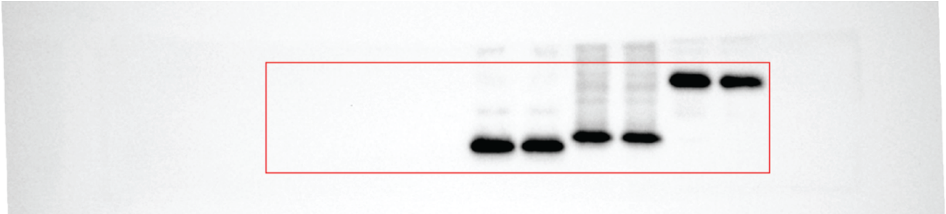

**Supplementary Fig. 22a:** Uncropped image of Chemiluminescence scan in reference to Fig. 2g (HA).

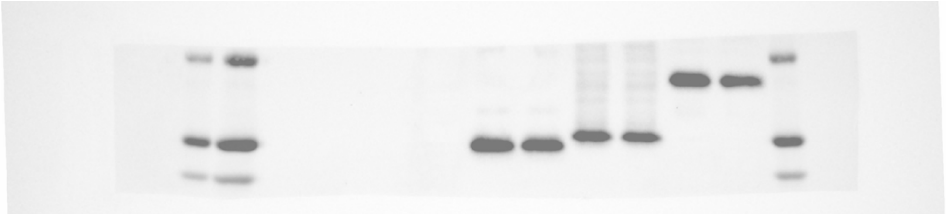

**Supplementary Fig. 22b:** Uncropped composite image of Cy5 and Chemiluminescence scan in reference to Fig. 2g (HA).

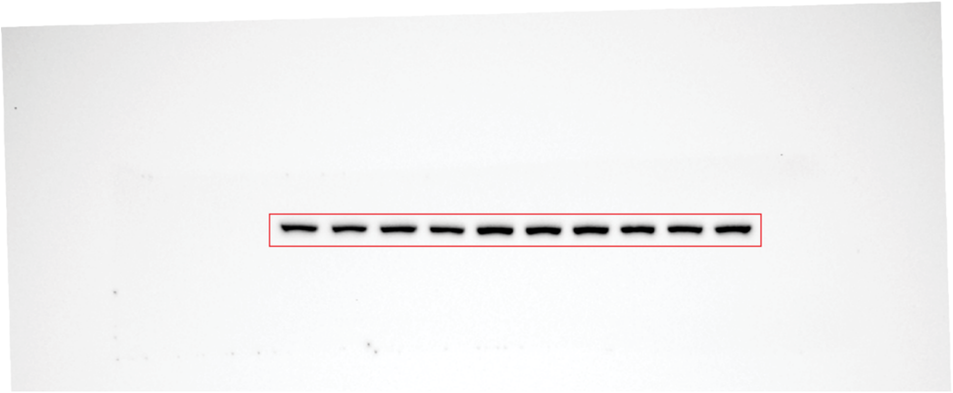

**Supplementary Fig. 23a:** Uncropped image of Chemiluminescence scan in reference to Fig. 2g (vinculin).

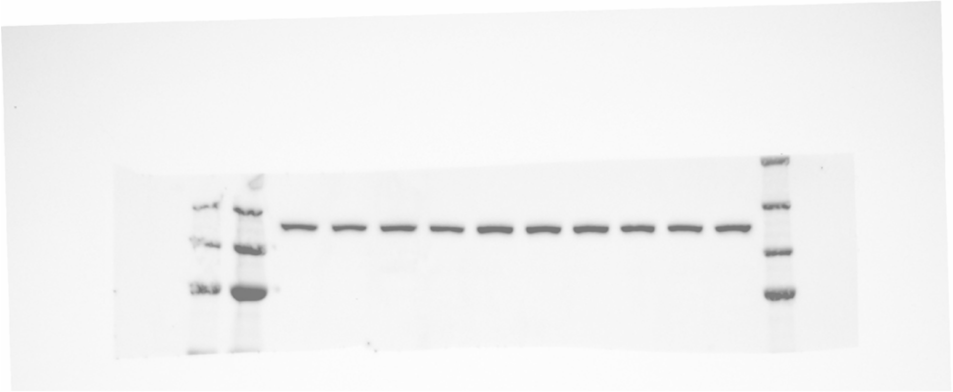

**Supplementary Fig. 23b:** Uncropped composite image of Cy5 and Chemiluminescence scan in reference to Fig. 2g (vinculin).

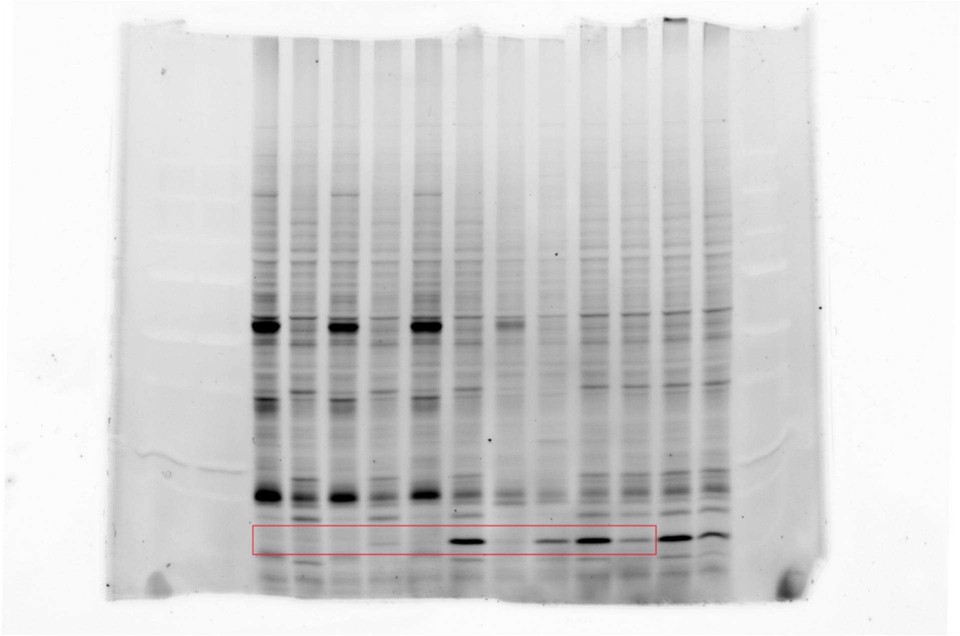

**Supplementary Fig. 24a:** Uncropped image of Rhodamine scan in reference to Fig. 2h (ABPP).

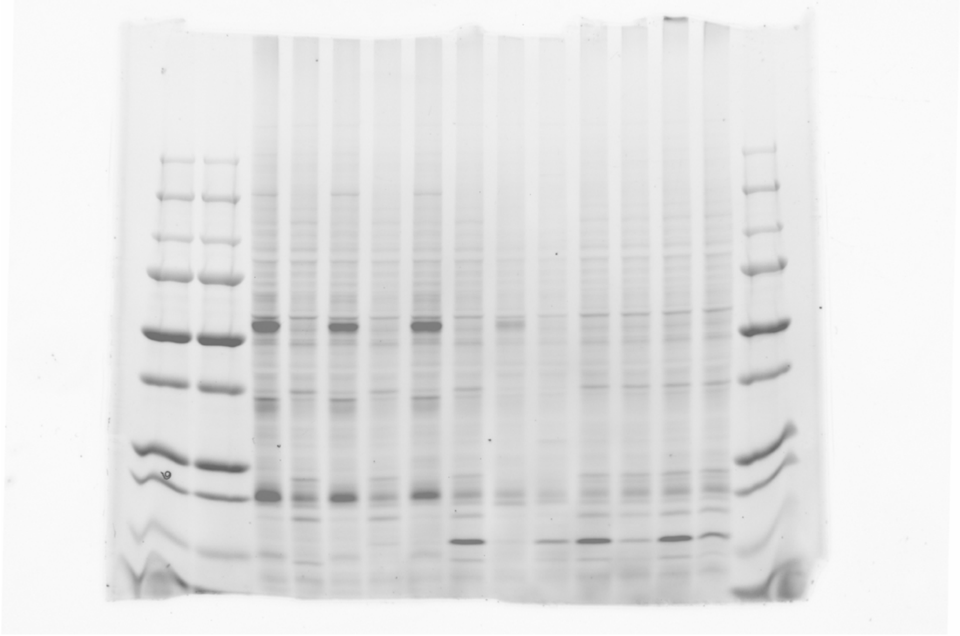

**Supplementary Fig. 24b:** Uncropped composite image of Cy5 and Rhodamine scan in reference to Fig. 2h (ABPP).

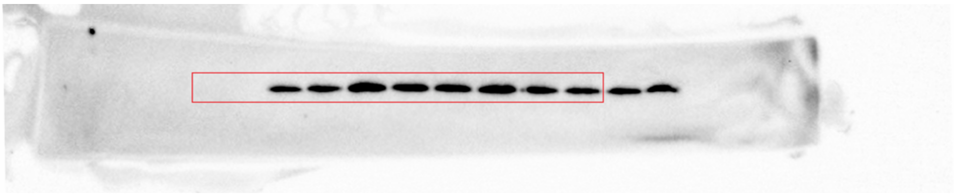

**Supplementary Fig. 25a:** Uncropped image of Chemiluminescence scan in reference to Fig. 2h (FLAG).

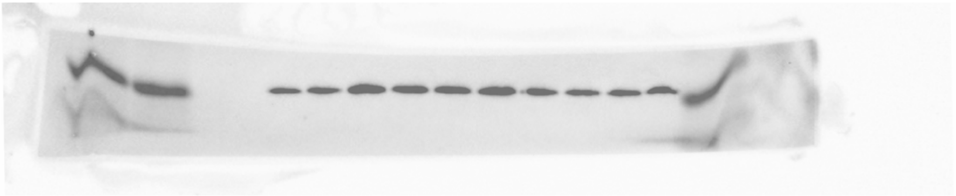

**Supplementary Fig. 25b:** Uncropped composite image of Cy5 and Chemiluminescence scan in reference to Fig. 2h (FLAG).

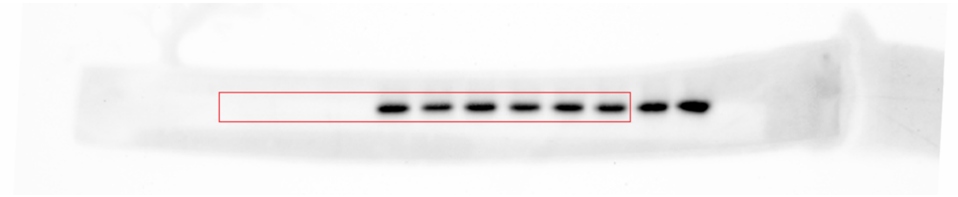

**Supplementary Fig. 26a:** Uncropped image of Chemiluminescence scan in reference to Fig. 2h (HA).

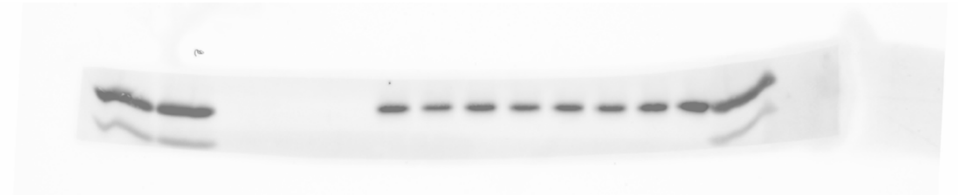

**Supplementary Fig. 26b:** Uncropped composite image of Cy5 and Chemiluminescence scan in reference to Fig. 2h (HA).

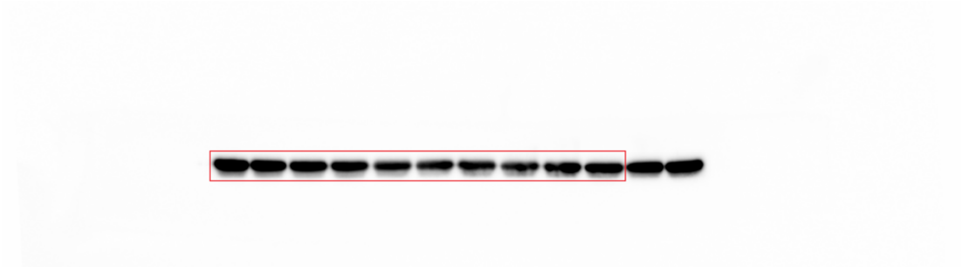

**Supplementary Fig. 27a:** Uncropped image of Chemiluminescence scan in reference to Fig. 2h (β-actin).

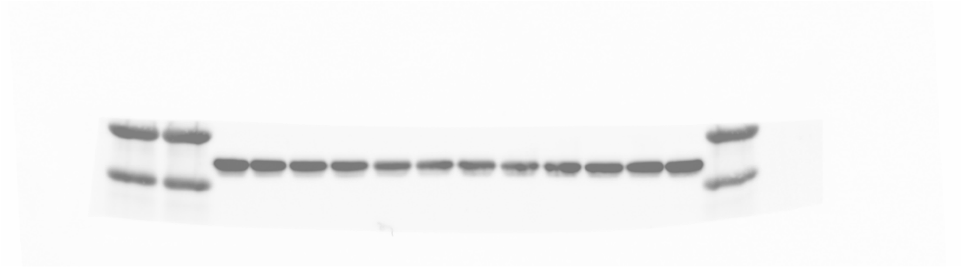

**Supplementary Fig. 27b:** Uncropped composite image of Cy5 and Chemiluminescence scan in reference to Fig. 2h (β-actin).

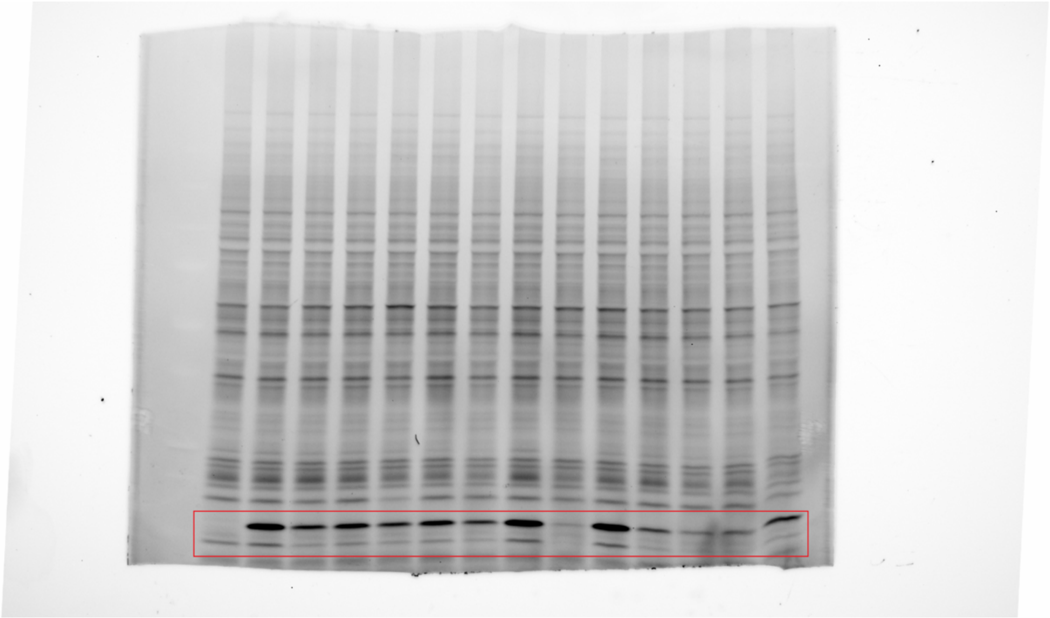

**Supplementary Fig. 28a:** Uncropped image of Rhodamine scan in reference to Fig. 3c (ABPP).

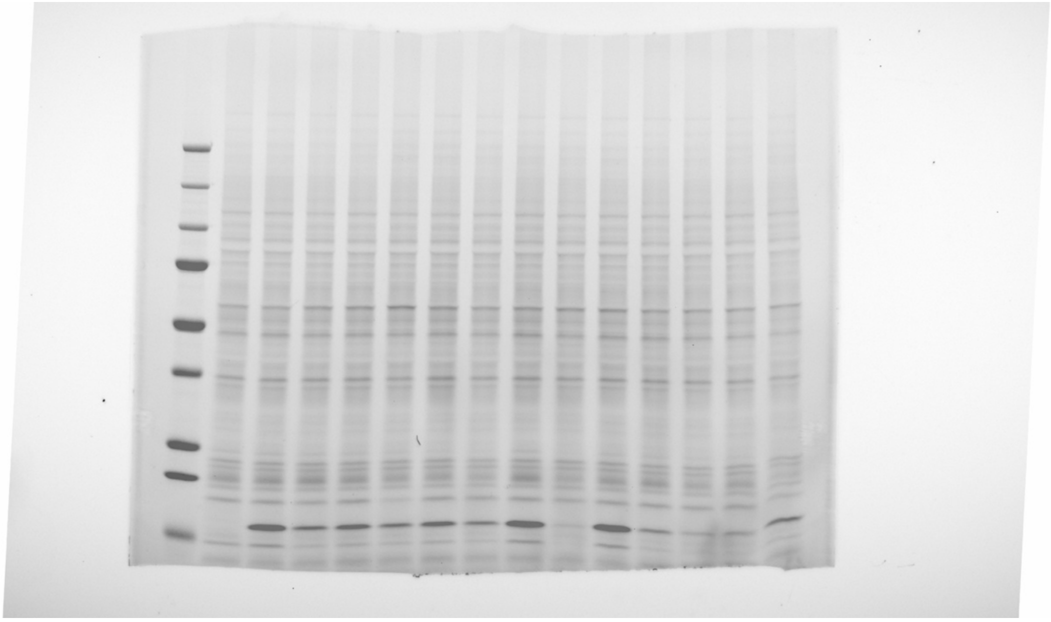

**Supplementary Fig. 28b:** Uncropped composite image of Cy5 and Rhodamine scan in reference to Fig. 3c (ABPP).

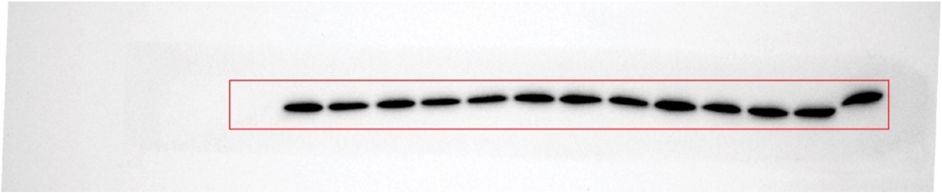

**Supplementary Fig. 29a:** Uncropped image of Chemiluminescence scan in reference to Fig. 3c (FLAG).

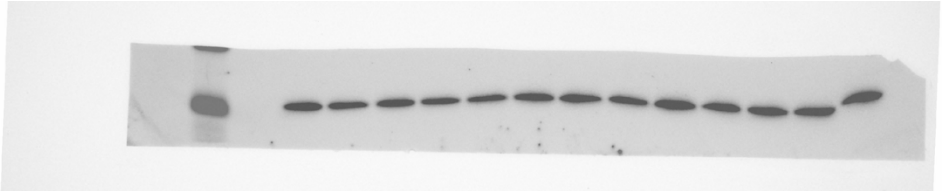

**Supplementary Fig. 29b:** Uncropped composite image of Cy5 and Chemiluminescence scan in reference to Fig. 3c (FLAG).

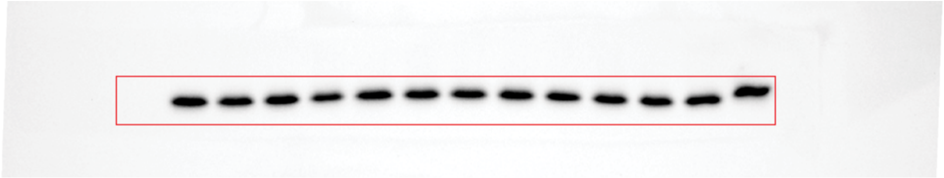

**Supplementary Fig. 30a:** Uncropped image of Chemiluminescence scan in reference to Fig. 3c (HA).

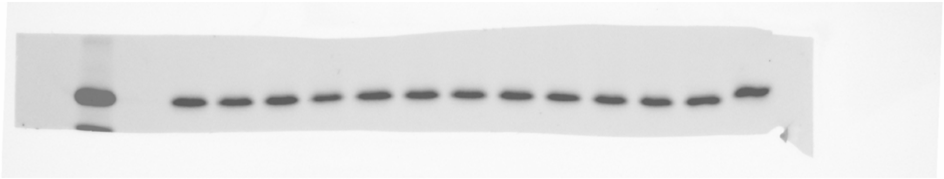

**Supplementary Fig. 30b:** Uncropped composite image of Cy5 and Chemiluminescence scan in reference to Fig. 3c (HA).

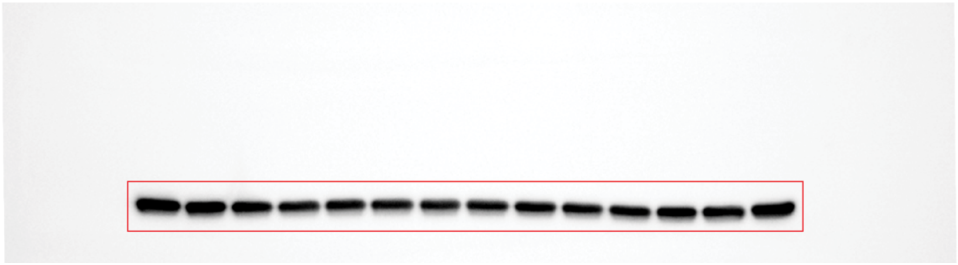

**Supplementary Fig. 31a:** Uncropped image of Chemiluminescence scan in reference to Fig. 3c (β-actin).

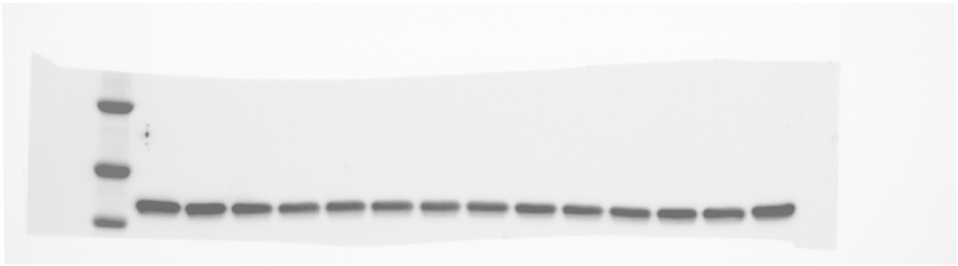

**Supplementary Fig. 31b:** Uncropped composite image of Cy5 and Chemiluminescence scan in reference to Fig. 3c (β-actin).

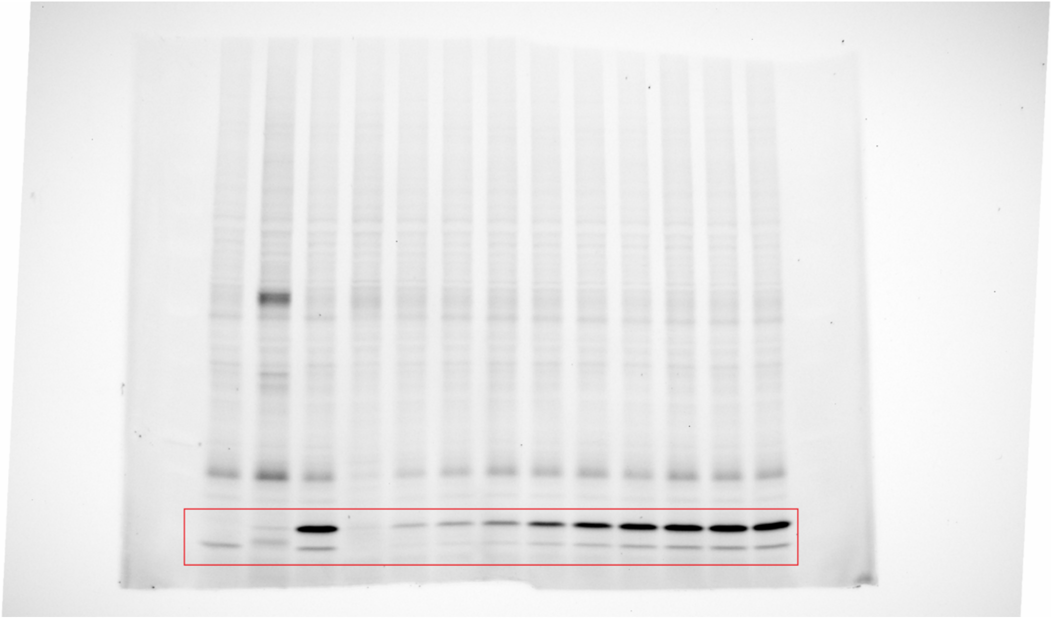

**Supplementary Fig. 32a:** Uncropped image of Rhodamine scan in reference to Fig. 3d (ABPP).

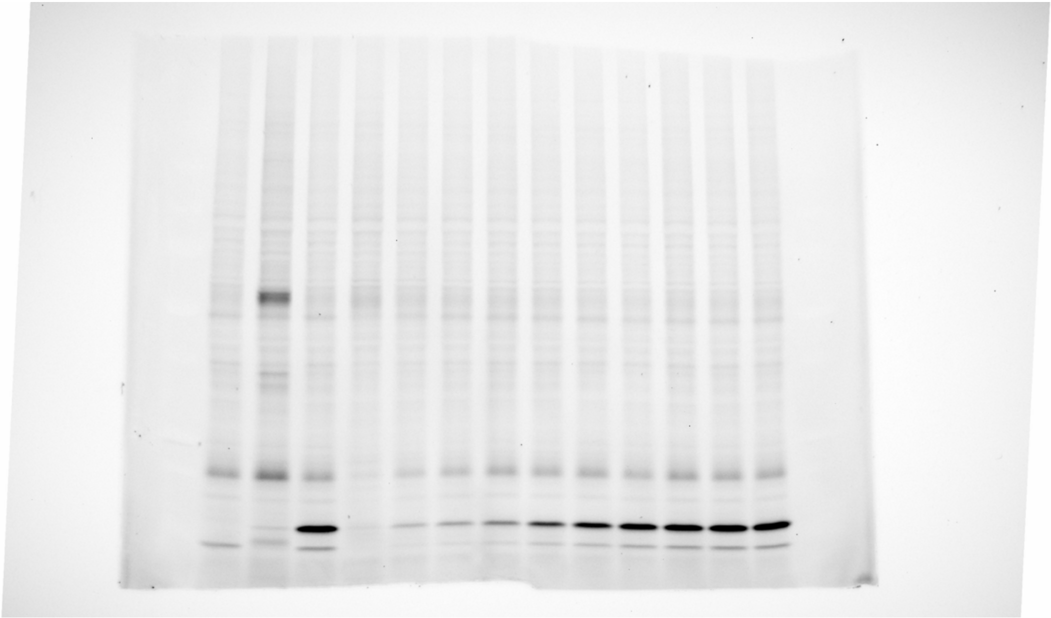

**Supplementary Fig. 32b:** Uncropped composite image of Cy5 and Rhodamine scan in reference to Fig. 3d (ABPP).

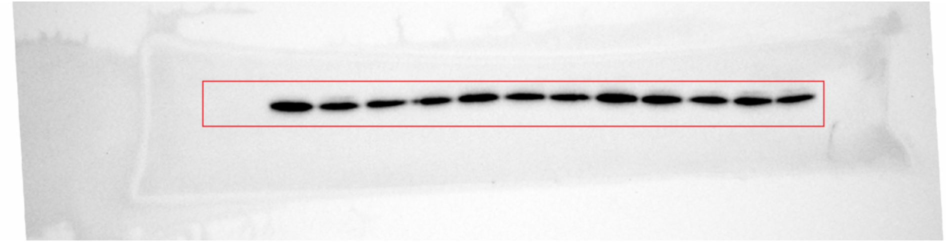

**Supplementary Fig. 33a:** Uncropped image of Chemiluminescence scan in reference to Fig. 3d (FLAG).

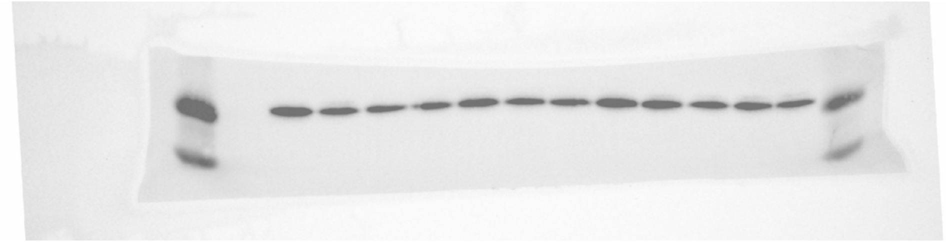

**Supplementary Fig. 33b:** Uncropped composite image of Cy5 and Chemiluminescence scan in reference to Fig. 3d (FLAG).

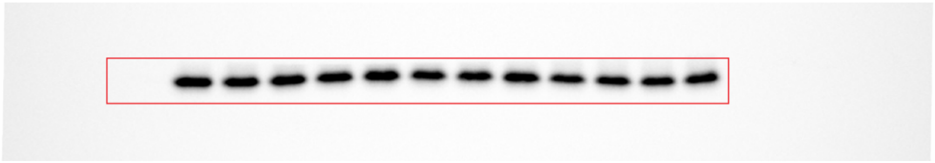

**Supplementary Fig. 34a:** Uncropped image of Chemiluminescence scan in reference to Fig. 3d (HA).

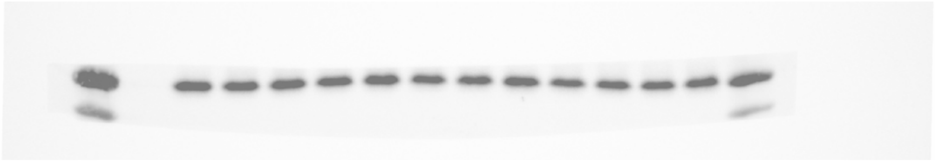

**Supplementary Fig. 34b:** Uncropped composite image of Cy5 and Chemiluminescence scan in reference to Fig. 3d (HA).

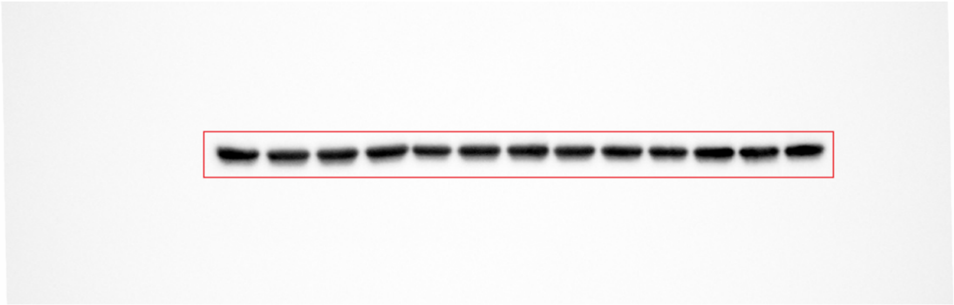

**Supplementary Fig. 35a:** Uncropped image of Chemiluminescence scan in reference to Fig. 3d (β-actin).

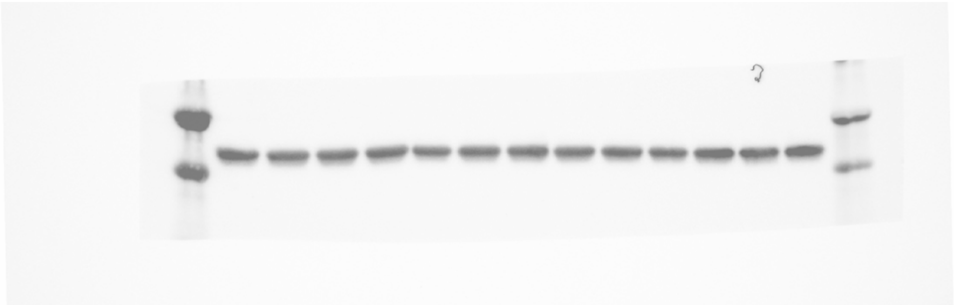

**Supplementary Fig. 35b:** Uncropped composite image of Cy5 and Chemiluminescence scan in reference to Fig. 3d (β-actin).

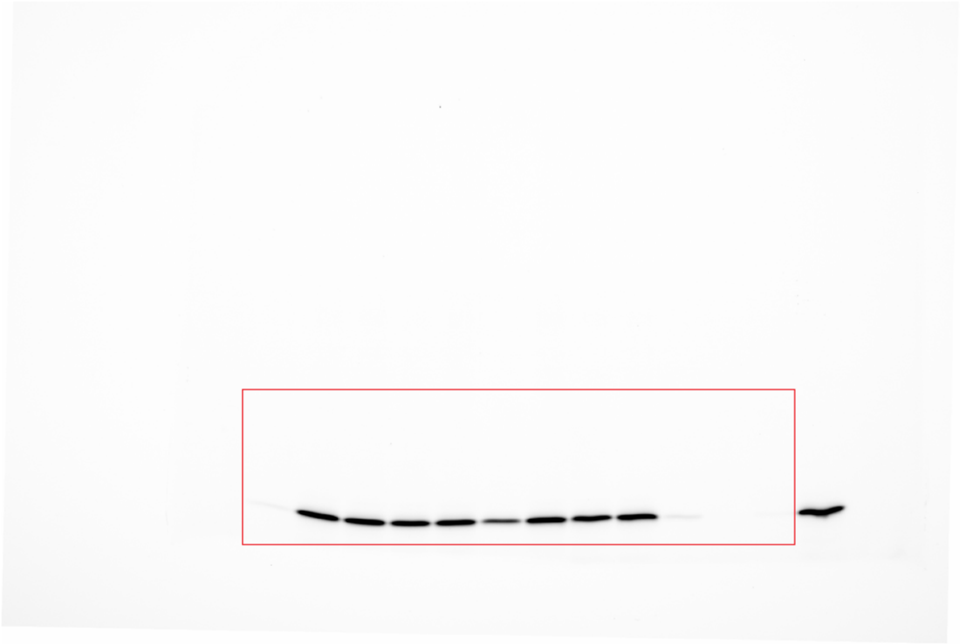

**Supplementary Fig. 36a:** Uncropped image of Rhodamine scan in reference to Fig. 4a (ABPP).

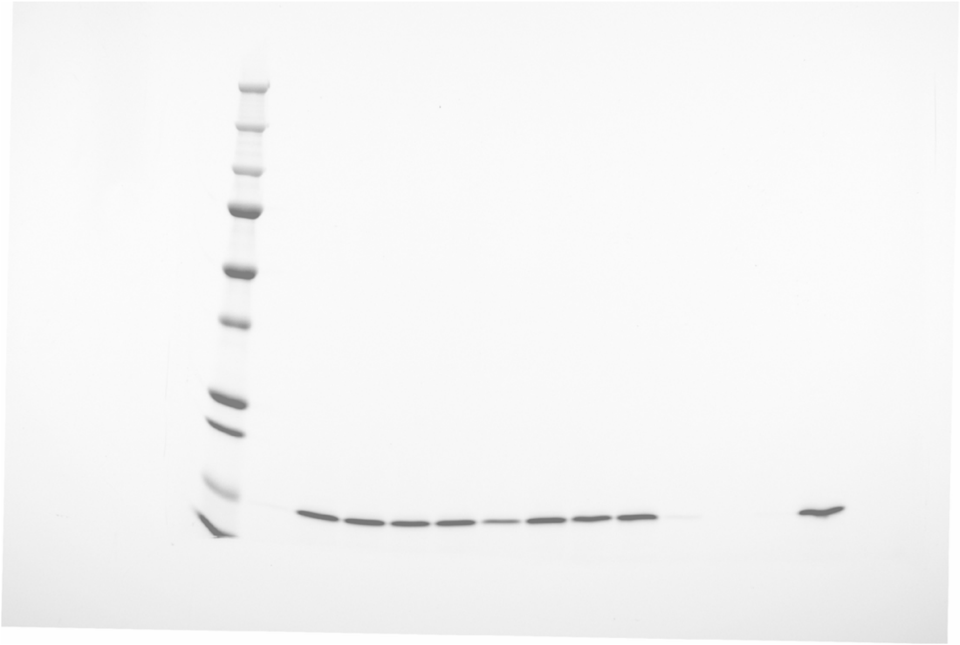

**Supplementary Fig. 36b:** Uncropped composite image of Cy5 and Rhodamine scan in reference to Fig. 4a (ABPP).

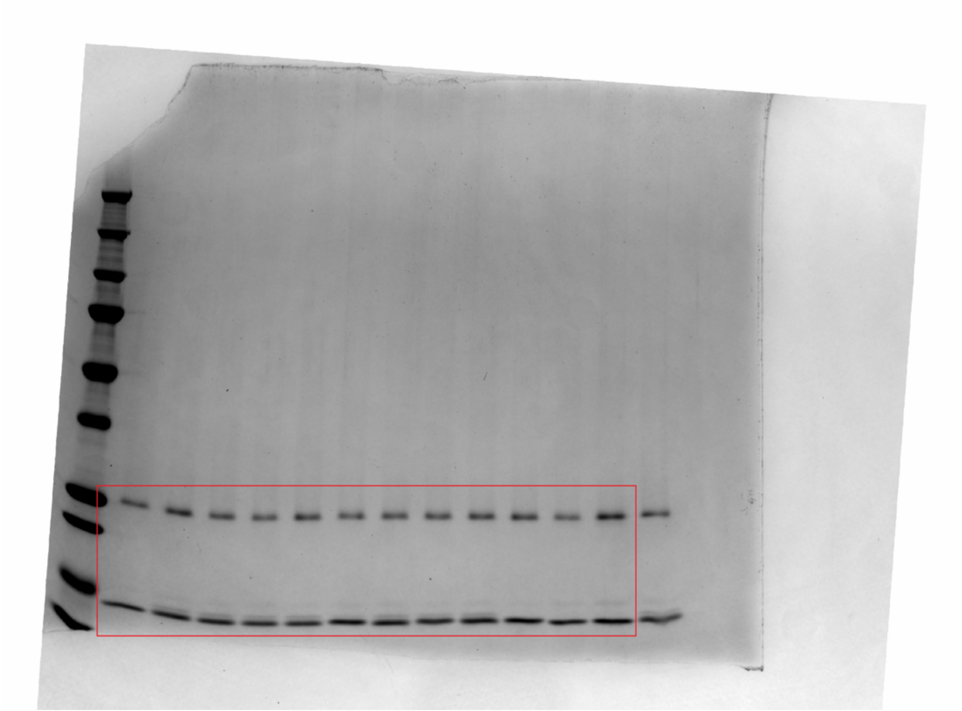

**Supplementary Fig. 37:** Uncropped image of Coomassie scan in reference to Fig. 4a (Coomassie).

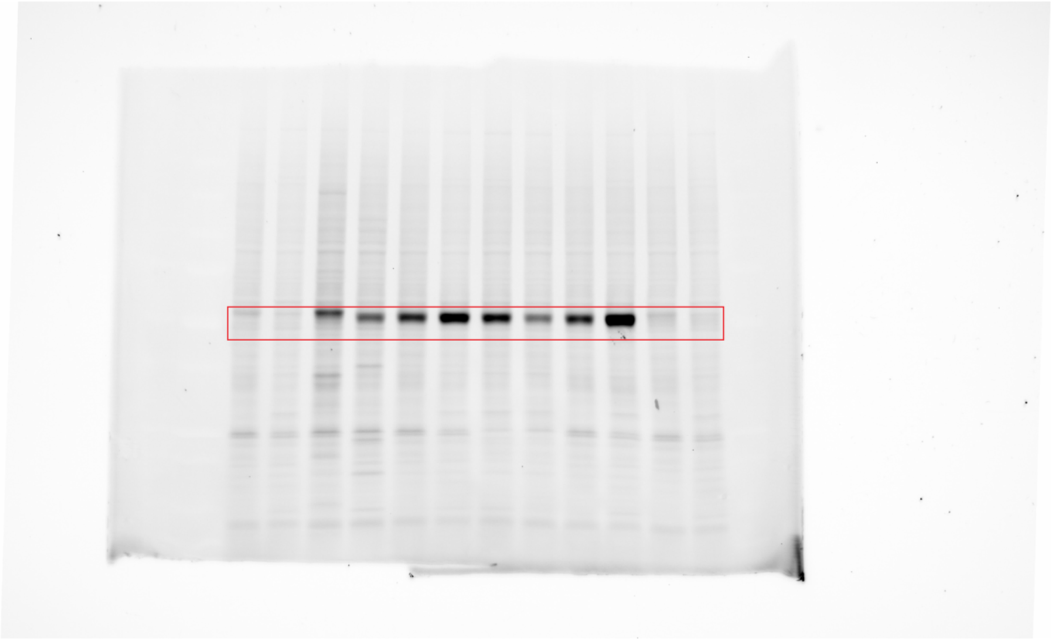

**Supplementary Fig. 38a:** Uncropped image of Rhodamine scan in reference to Extended Data Fig. 4f (ABPP).

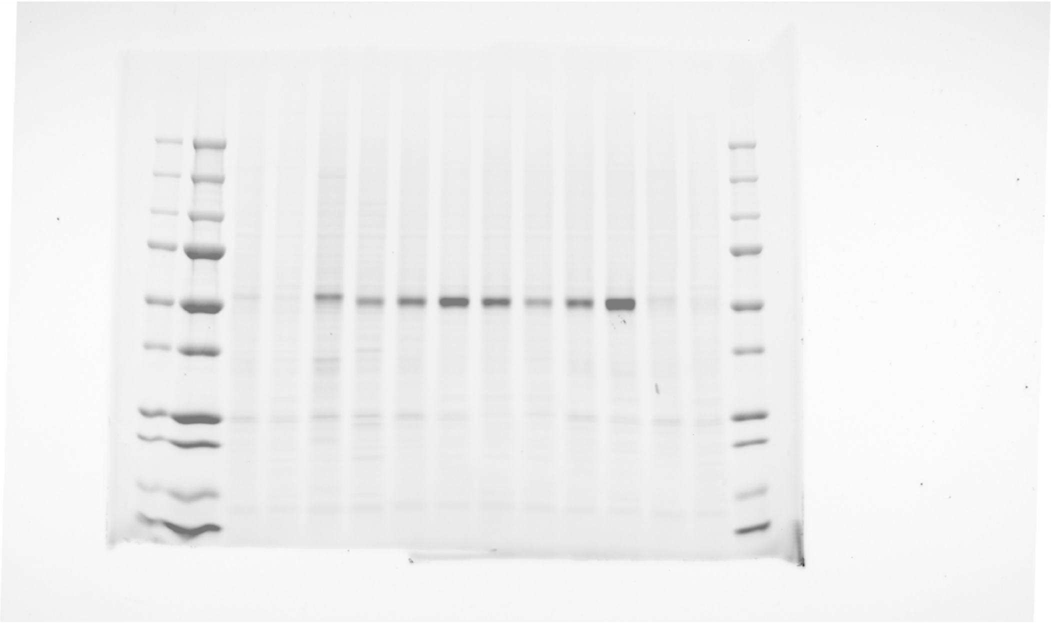

**Supplementary Fig. 38b:** Uncropped composite image of Cy5 and Rhodamine scan in reference to Extended Data Fig. 4f (ABPP).

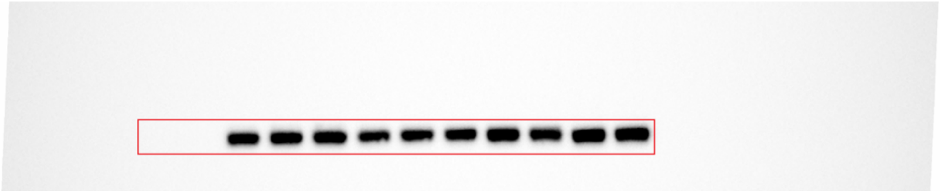

**Supplementary Fig. 39a:** Uncropped image of Chemiluminescence scan in reference to Extended Data Fig. 4f (FLAG).

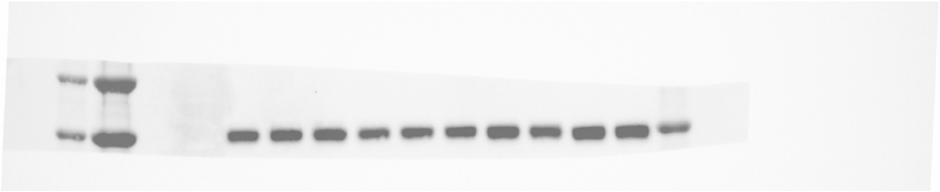

**Supplementary Fig. 39b:** Uncropped composite image of Cy5 and Chemiluminescence scan in reference to Extended Data Fig. 4f (FLAG).

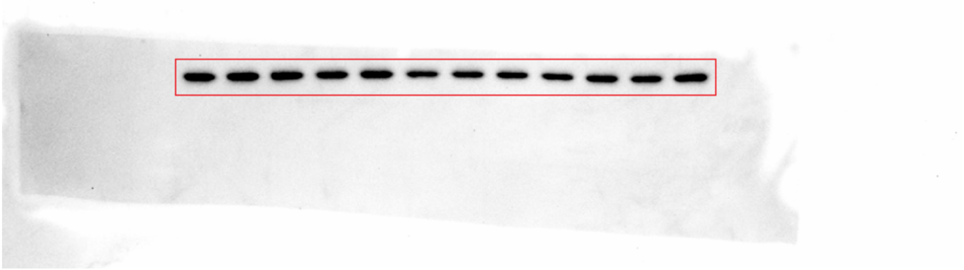

**Supplementary Fig. 40a:** Uncropped image of Chemiluminescence scan in reference to Extended Data Fig. 4f (GAPDH).

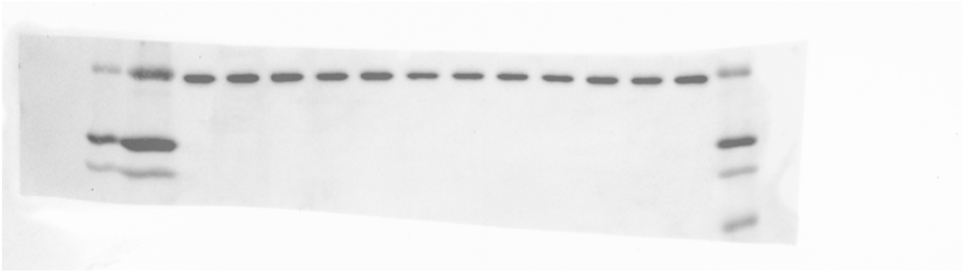

**Supplementary Fig. 40b:** Uncropped composite image of Cy5 and Chemiluminescence scan in reference to Extended Data Fig. 4f (GAPDH).

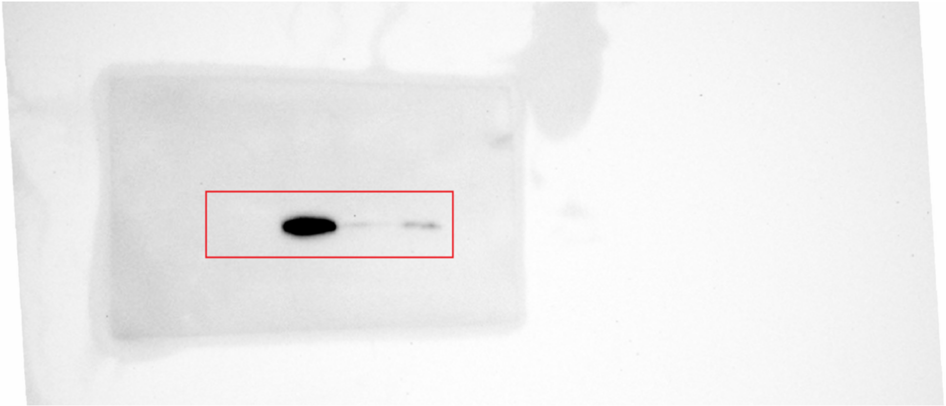

**Supplementary Fig. 41a:** Uncropped image of Chemiluminescence scan in reference to Extended Data Fig. 5f (FLAG).

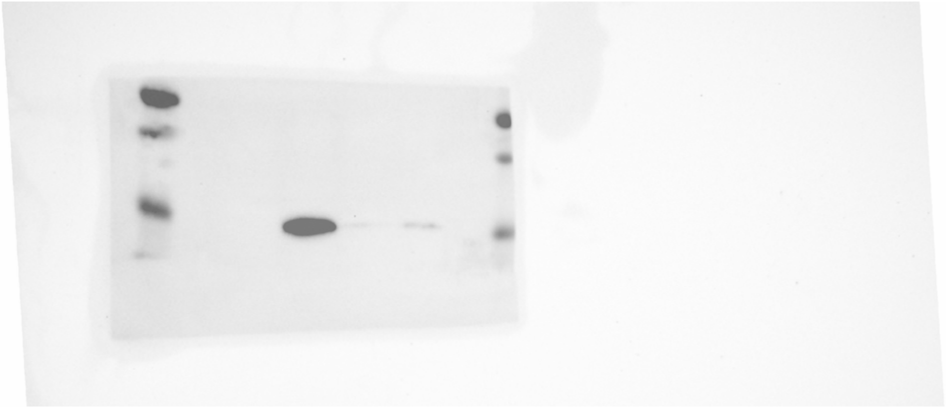

**Supplementary Fig. 41b:** Uncropped composite image of Cy5 and Chemiluminescence scan in reference to Extended Data Fig. 5f (FLAG).

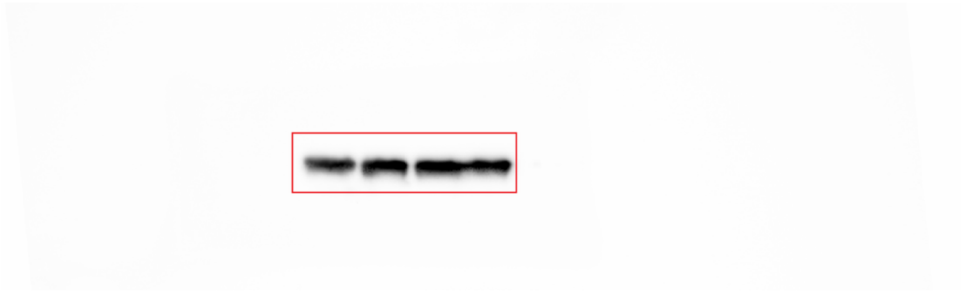

**Supplementary Fig. 42a:** Uncropped image of Chemiluminescence scan in reference to Extended Data Fig. 5f (β-actin).

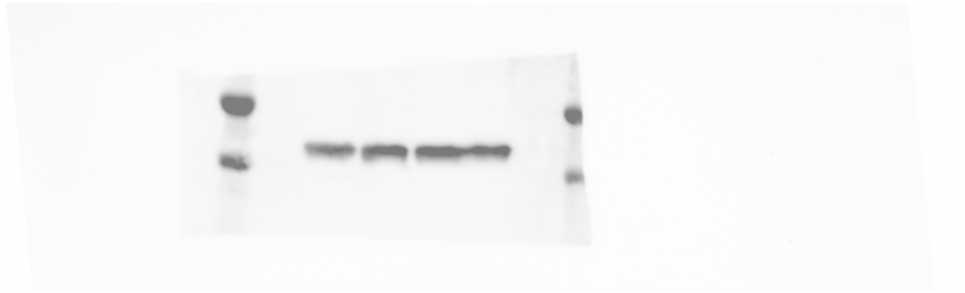

**Supplementary Fig. 42b:** Uncropped composite image of Cy5 and Chemiluminescence scan in reference to Extended Data Fig. 5f (β-actin).

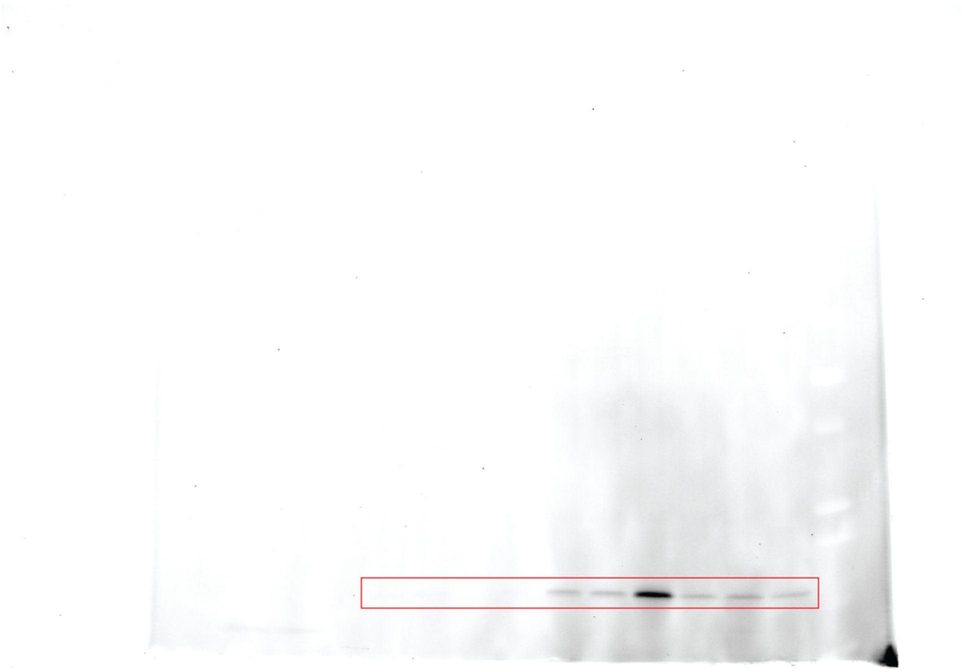

**Supplementary Fig. 43a:** Uncropped image of Rhodamine scan in reference to Extended Data Fig. 4g (ABPP).

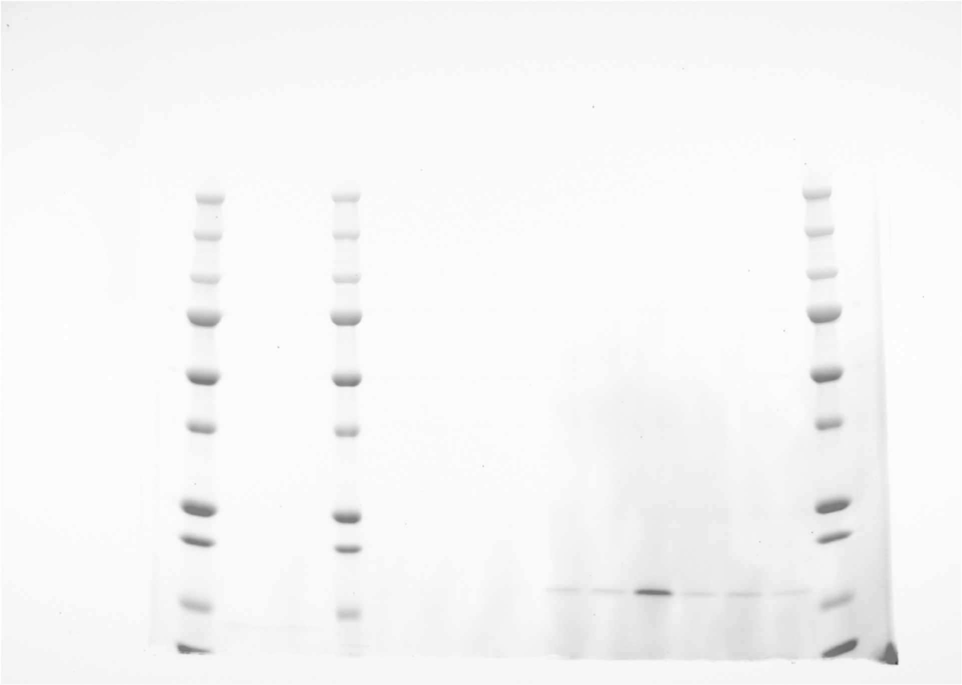

**Supplementary Fig. 43b:** Uncropped composite image of Cy5 and Rhodamine scan in reference to Extended Data Fig. 4g (ABPP).

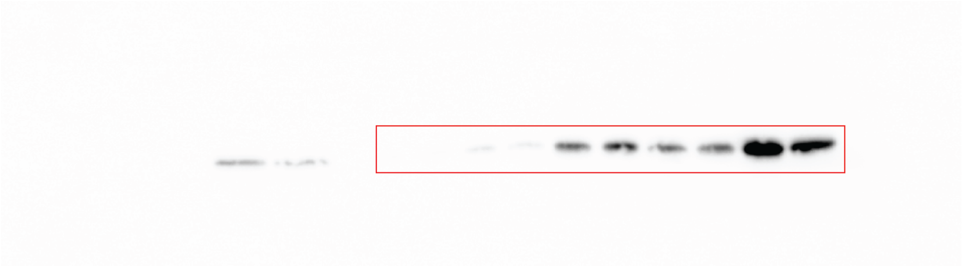

**Supplementary Fig. 44a:** Uncropped image of Chemiluminescence scan in reference to Extended Data Fig. 5g (FLAG, short exposure).

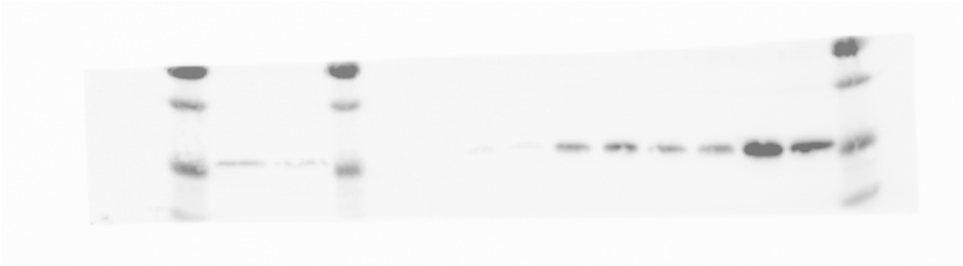

**Supplementary Fig. 44b:** Uncropped composite image of Cy5 and Chemiluminescence scan in reference to Extended Data Fig. 5g (FLAG, short exposure).

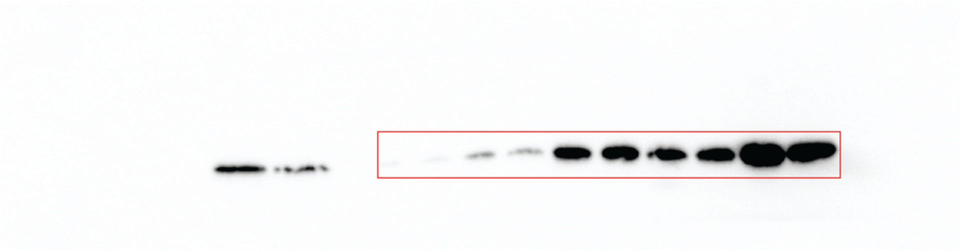

**Supplementary Fig. 45a:** Uncropped image of Chemiluminescence scan in reference to Extended Data Fig. 5g (FLAG, long exposure).

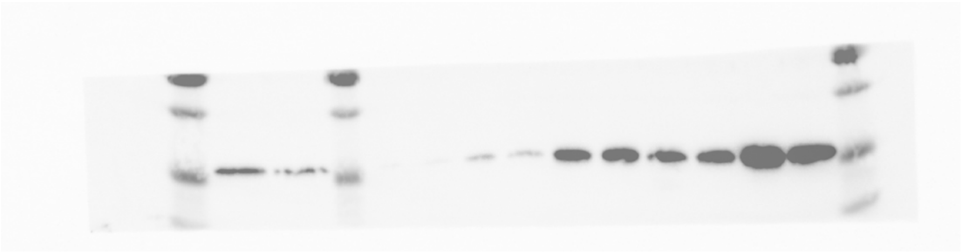

**Supplementary Fig. 45b:** Uncropped composite image of Cy5 and Chemiluminescence scan in reference to Extended Data Fig. 5g (FLAG, long exposure).

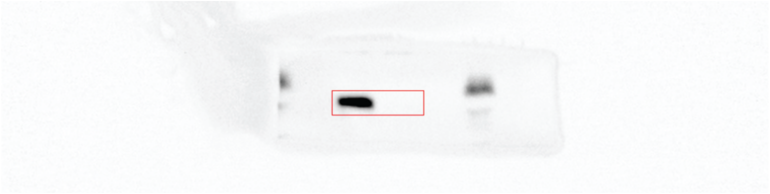

**Supplementary Fig. 46a:** Uncropped image of Chemiluminescence scan in reference to Extended Data Fig. 5h (METTL5).

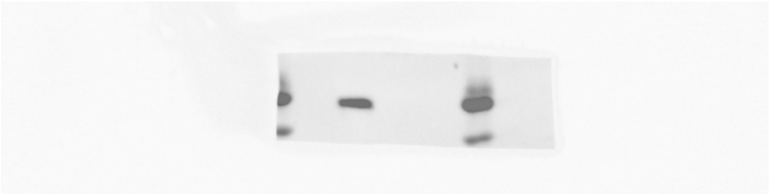

**Supplementary Fig. 46b:** Uncropped composite image of Cy5 and Chemiluminescence scan in reference to Extended Data Fig. 5h (METTL5).

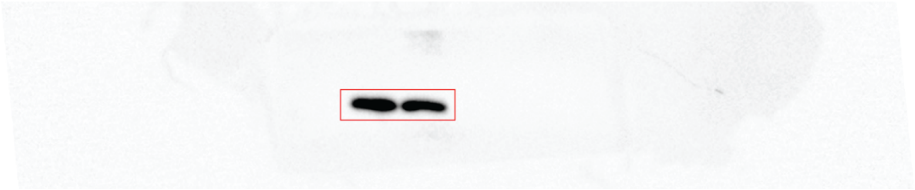

**Supplementary Fig. 47a:** Uncropped image of Chemiluminescence scan in reference to Extended Data Fig. 5h (TRMT112).

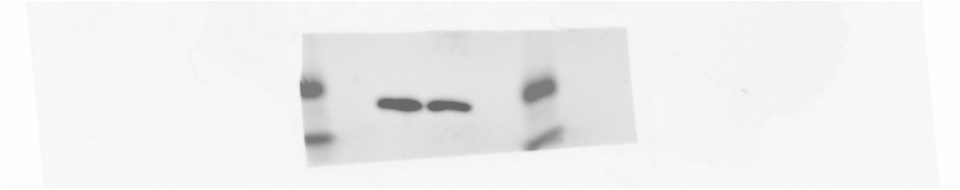

**Supplementary Fig. 47b:** Uncropped composite image of Cy5 and Chemiluminescence scan in reference to Extended Data Fig. 5h (TRMT112).

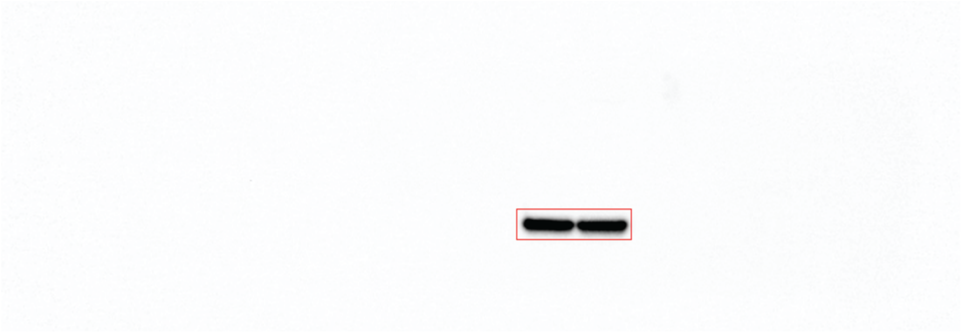

**Supplementary Fig. 48a:** Uncropped image of Chemiluminescence scan in reference to Extended Data Fig. 5h (β-actin).

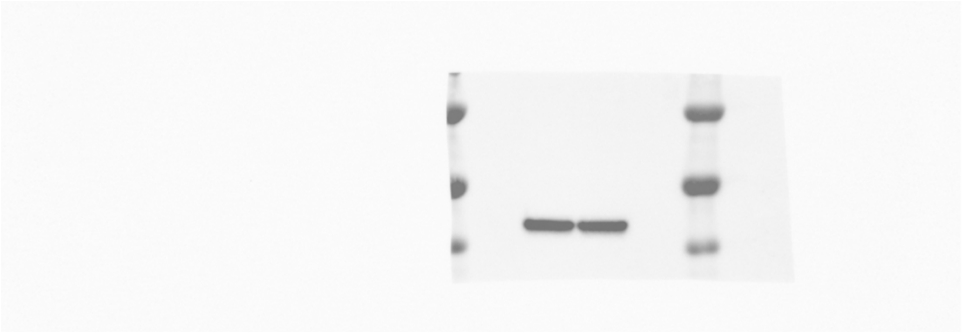

**Supplementary Fig. 48b:** Uncropped composite image of Cy5 and Chemiluminescence scan in reference to Extended Data Fig. 5h (β-actin).

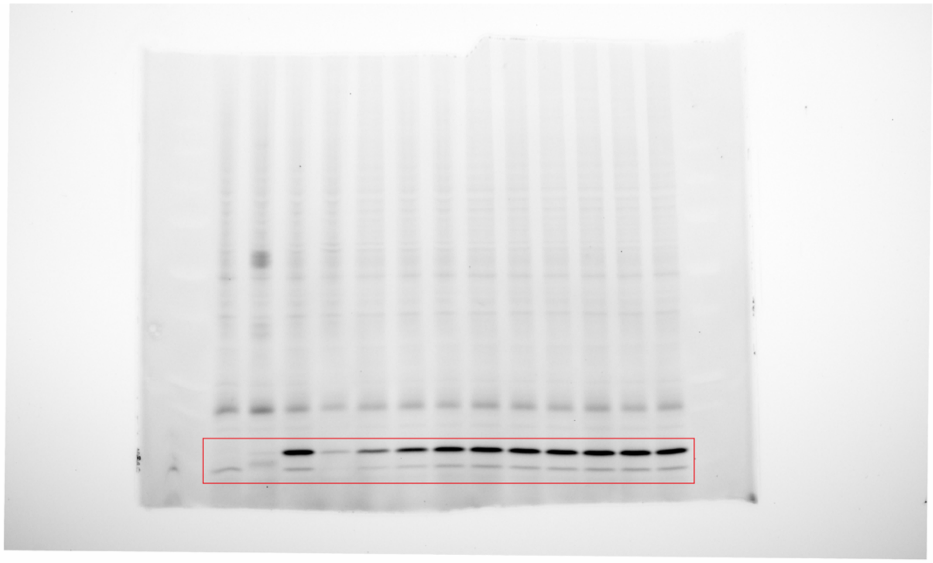

**Supplementary Fig. 49a:** Uncropped image of Rhodamine scan in reference to Extended Data Fig. 6a (ABPP).

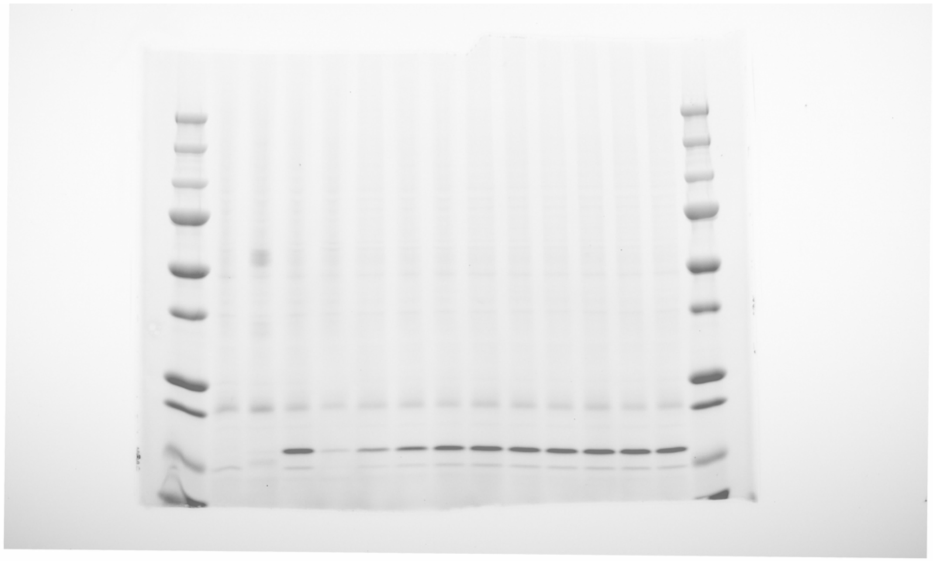

**Supplementary Fig. 49b:** Uncropped composite image of Cy5 and Rhodamine scan in reference to Extended Data Fig. 6a (ABPP).

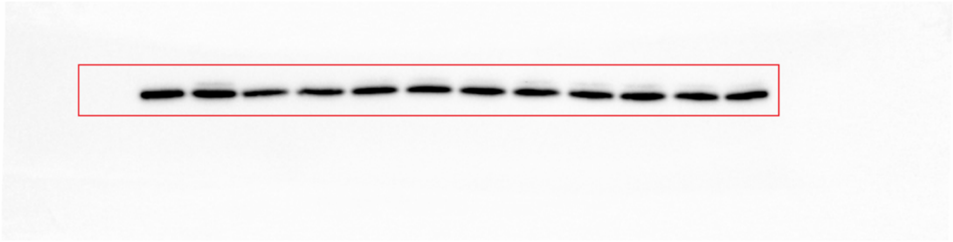

**Supplementary Fig. 50a:** Uncropped image of Chemiluminescence scan in reference to Extended Data Fig. 6a (FLAG).

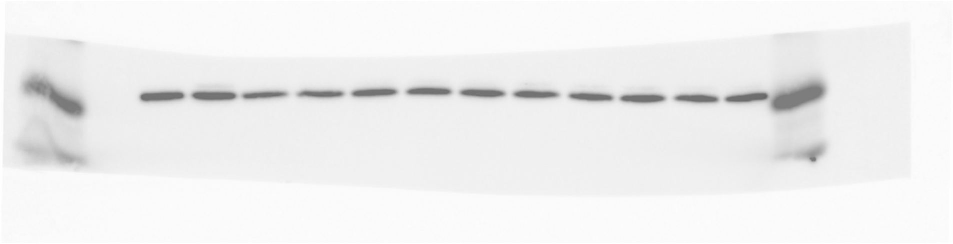

**Supplementary Fig. 50b:** Uncropped composite image of Cy5 and Chemiluminescence scan in reference to Extended Data Fig. 6a (FLAG).

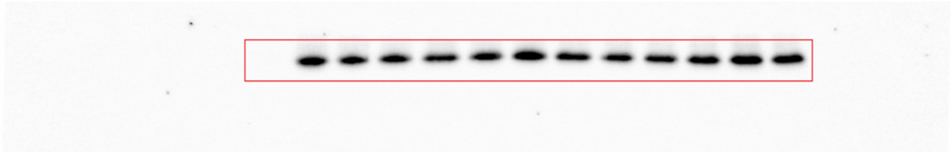

**Supplementary Fig. 51a:** Uncropped image of Chemiluminescence scan in reference to Extended Data Fig. 6a (HA).

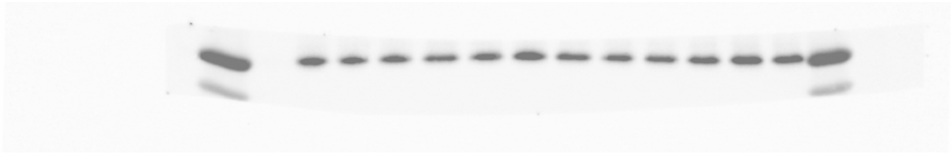

**Supplementary Fig. 51b:** Uncropped composite image of Cy5 and Chemiluminescence scan in reference to Extended Data Fig. 6a (HA).

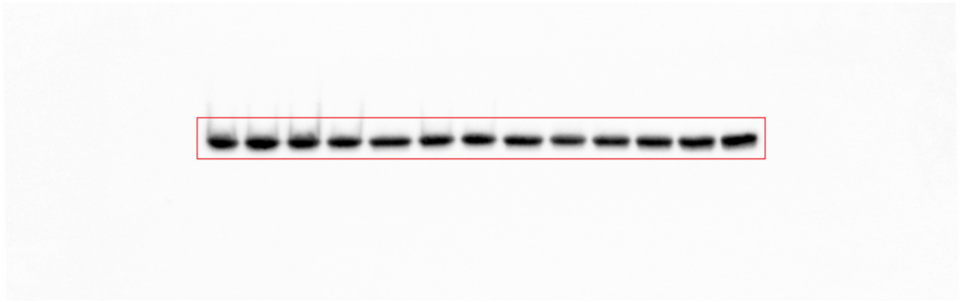

**Supplementary Fig. 52a:** Uncropped image of Chemiluminescence scan in reference to Extended Data Fig. 6a (β-actin).

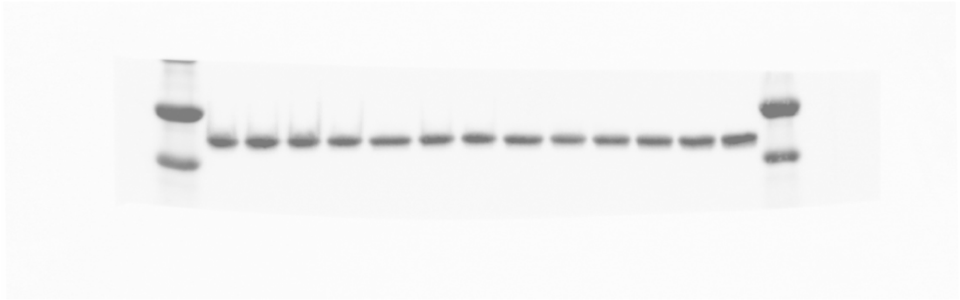

**Supplementary Fig. 52b:** Uncropped composite image of Cy5 and Chemiluminescence scan in reference to Extended Data Fig. 6a (β-actin).

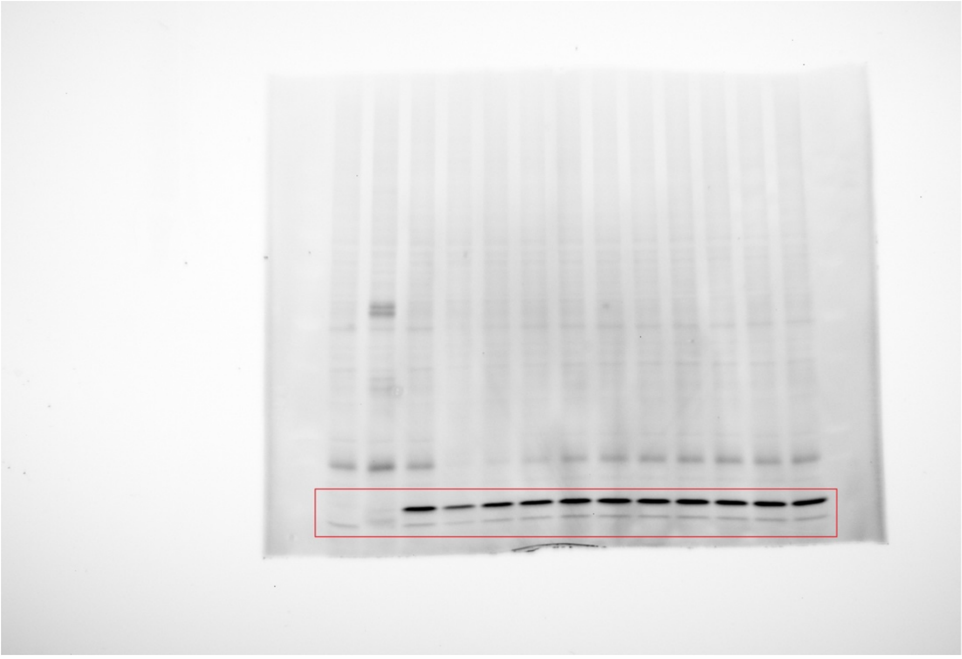

**Supplementary Fig. 53a:** Uncropped image of Rhodamine scan in reference to Extended Data Fig. 6d (ABPP).

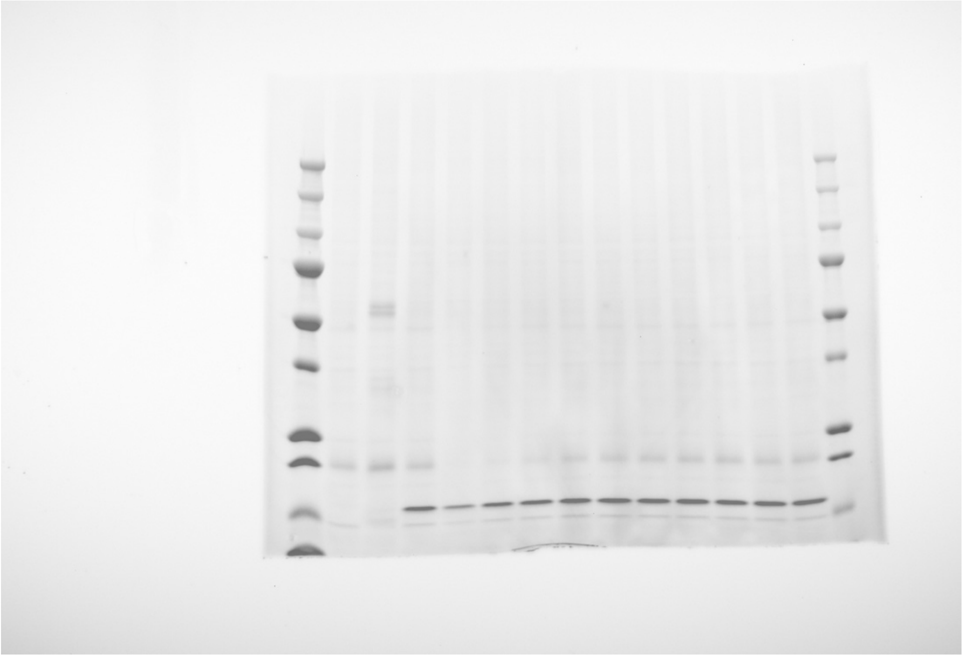

**Supplementary Fig. 53b:** Uncropped composite image of Cy5 and Rhodamine scan in reference to Extended Data Fig. 6d (ABPP).

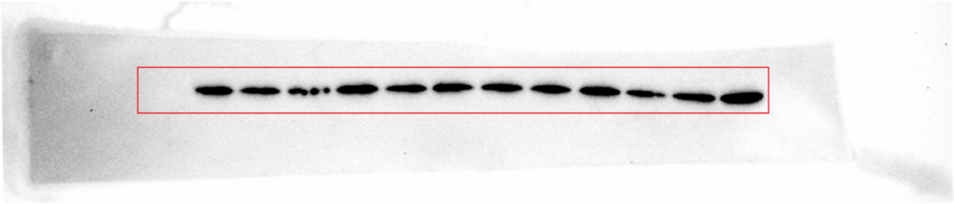

**Supplementary Fig. 54a:** Uncropped image of Chemiluminescence scan in reference to Extended Data Fig. 6d (FLAG).

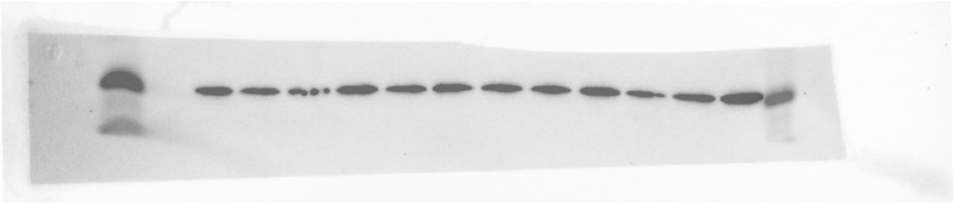

**Supplementary Fig. 54b:** Uncropped composite image of Cy5 and Chemiluminescence scan in reference to Extended Data Fig. 6d (FLAG).

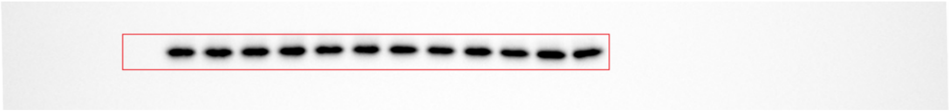

**Supplementary Fig. 55a:** Uncropped image of Chemiluminescence scan in reference to Extended Data Fig. 6d (HA).

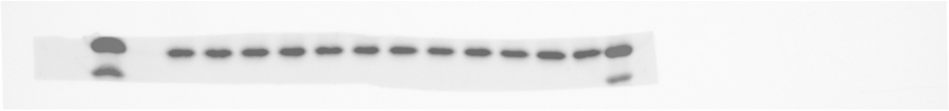

**Supplementary Fig. 55b:** Uncropped composite image of Cy5 and Chemiluminescence scan in reference to Extended Data Fig. 6d (HA).

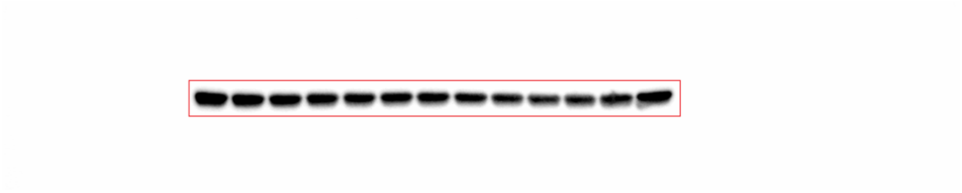

**Supplementary Fig. 56a:** Uncropped image of Chemiluminescence scan in reference to Extended Data Fig. 6d (β-actin).

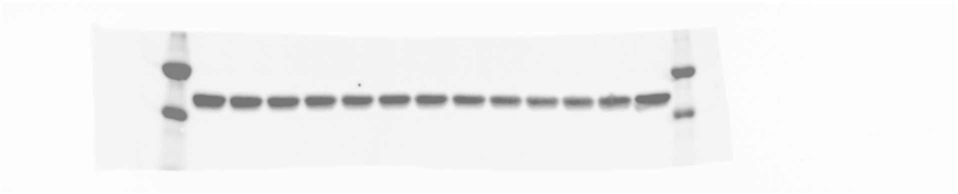

**Supplementary Fig. 56b:** Uncropped composite image of Cy5 and Chemiluminescence scan in reference to Extended Data Fig. 6d (β-actin).

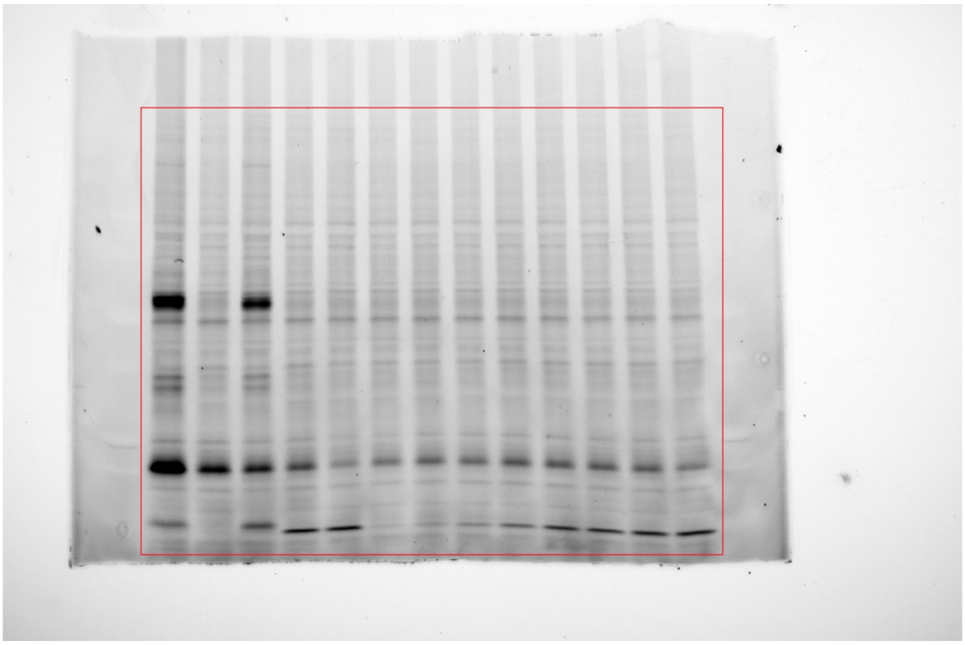

**Supplementary Fig. 57a:** Uncropped image of Rhodamine scan in reference to Extended Data Fig. 6d (ABPP).

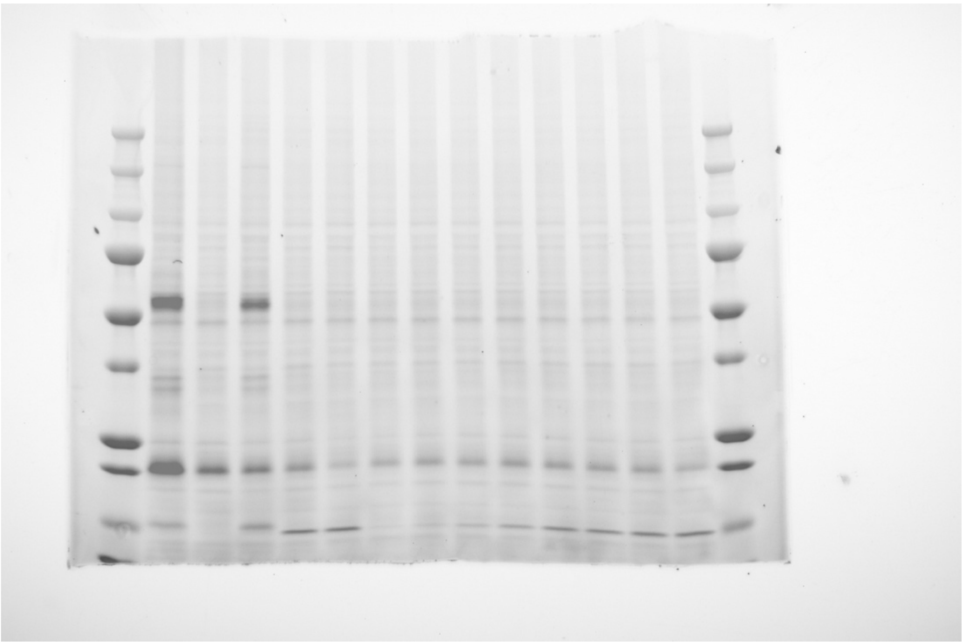

**Supplementary Fig. 57b:** Uncropped composite image of Cy5 and Rhodamine scan in reference to Extended Data Fig. 6d (ABPP).

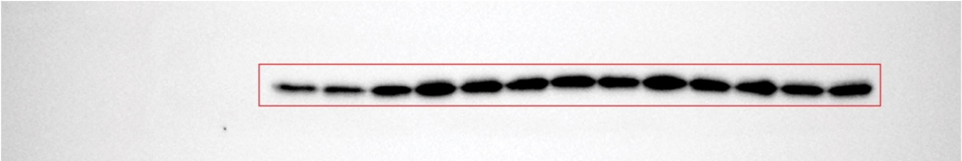

**Supplementary Fig. 58a:** Uncropped image of Chemiluminescence scan in reference to Extended Data Fig. 6d (TRMT112).

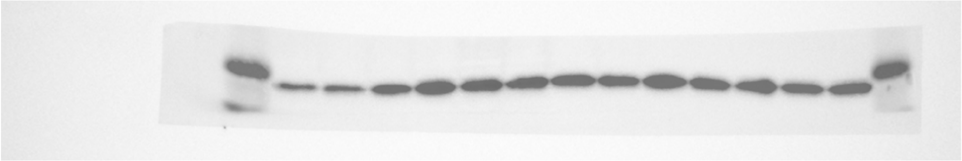

**Supplementary Fig. 58b:** Uncropped composite image of Cy5 and Chemiluminescence scan in reference to Extended Data Fig. 6d (TRMT112).

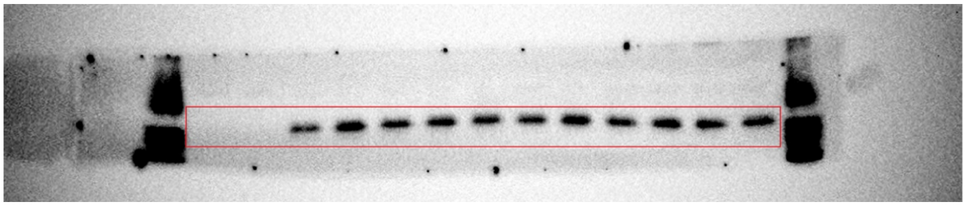

**Supplementary Fig. 59a:** Uncropped image of Chemiluminescence scan in reference to Extended Data Fig. 6d (METTL5).

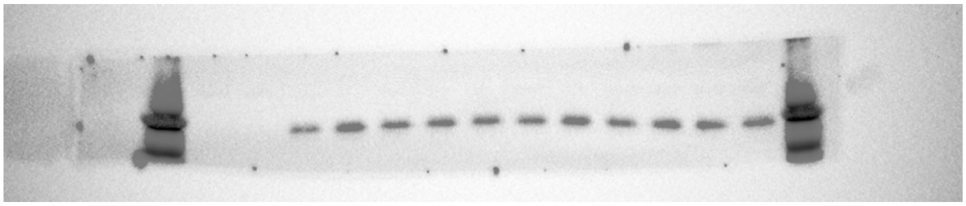

**Supplementary Fig. 59b:** Uncropped composite image of Cy5 and Chemiluminescence scan in reference to Extended Data Fig. 6d (METTL5).

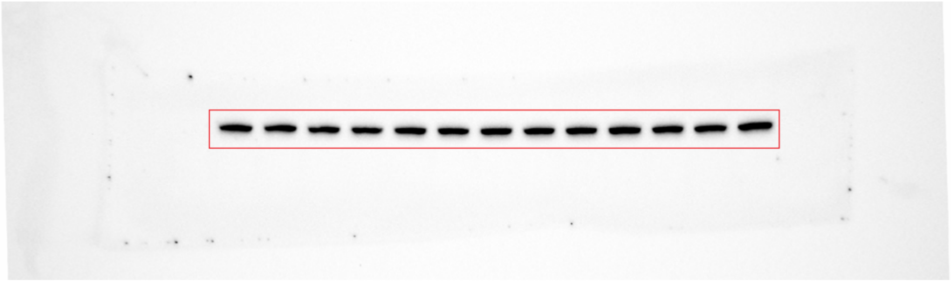

**Supplementary Fig. 60a:** Uncropped image of Chemiluminescence scan in reference to Extended Data Fig. 6d (vinculin).

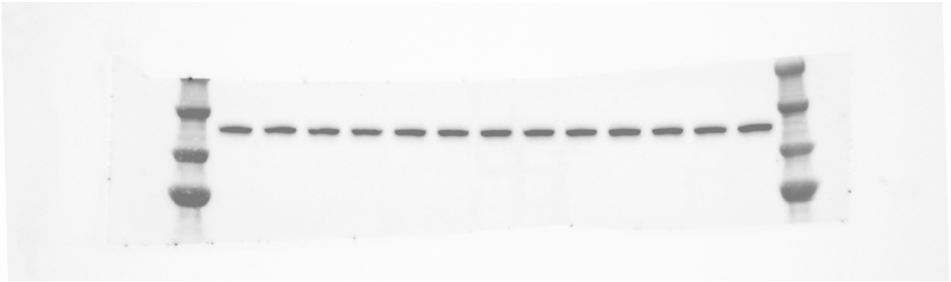

**Supplementary Fig. 60b:** Uncropped composite image of Cy5 and Chemiluminescence scan in reference to Extended Data Fig. 6d (vinculin).
